# That’s not a Hybrid: How to Distinguish Patterns of Admixture and Isolation-by-Distance

**DOI:** 10.1101/2024.04.15.589658

**Authors:** Ben J. Wiens, Jocelyn P. Colella

## Abstract

Describing naturally occurring genetic variation is a fundamental goal of molecular phylogeography and population genetics. Popular methods for this task include *STRUCTURE*, a model-based algorithm that assigns individuals to genetic clusters, and Principal Component Analysis (PCA), a parameter-free method. The ability of *STRUCTURE* to infer mixed ancestry makes it popular for documenting natural hybridization, which is of considerable interest to evolutionary biologists, given that such systems provide a window into the speciation process. Yet *STRUCTURE* can produce misleading results when its underlying assumptions are violated, like when genetic variation is distributed continuously. To test the ability of *STRUCTURE* and PCA to accurately distinguish admixture from continuous variation, we use forward-time simulations to generate population genetic data under three demographic scenarios: two involving admixture and one with isolation-by-distance (IBD). *STRUCTURE* and PCA alone cannot distinguish admixture from IBD, but complementing these analyses with triangle plots, which visualize hybrid index against interclass heterozygosity, provides more accurate inference of demographic history. We demonstrate that triangle plots are robust to missing data, while *STRUCTURE* and PCA are not, and show that setting a low allele frequency difference threshold for AIM identification can accurately characterize the relationship between hybrid index and interclass heterozygosity across demographic histories of admixture and range expansion. While *STRUCTURE* and PCA provide useful summaries of genetic variation, results should be paired with triangle plots before admixture is inferred.

## Introduction

Inferring the number of genetic clusters and ascribing ancestry proportions to individuals is a near-ubiquitous first step when confronted with genetic data sourced from wild populations (Novembre, 2016; Lawson et al., 2018). The program *STRUCTURE* was among the first to address this task, by implementing model-based clustering under a Bayesian framework (Pritchard et al., 2000; Novembre, 2016). While similar Bayesian models (Alexander et al., 2009; Anderson & Thompson, 2002; Corander & Marttinen, 2006; Gao et al., 2007; Guillot et al., 2005; Huelsenbeck & Andolfatto, 2007; Wilson & Rannala, 2003) and alternative approaches have since been introduced (Beugin et al., 2018; Jombart et al., 2010; Novembre & Stephens, 2008; Pickrell & Pritchard, 2012; Tang et al., 2005) *STRUCTURE* remains one of the most widely used approaches to describing genetic variation in natural populations. To that point, in 2023 alone, the original *STRUCTURE* paper (Pritchard et al., 2000) received over 2,000 citations.

*STRUCTURE* excels at identifying genetic clusters when the underlying assumptions of the model are met, but it can provide misleading results when those assumptions are violated (Bradburd et al., 2018; Latch et al., 2006; Lawson et al., 2018; Novembre, 2016; Puechmaille, 2016; Wang, 2017). Indeed, it is often the case that empirical datasets do not meet the assumptions of *STRUCTURE* (Lawson et al., 2018). For example, *STRUCTURE* assumes that individual genotypes are randomly drawn from a set of K discrete populations, each with their own set of allele frequencies (Bradburd et al., 2018). When that assumption is not met, such as when genetic variation is continuous across the landscape, population structure is hierarchical, or populations have experienced different amounts of genetic drift, *STRUCTURE* results do not reflect the true demographic history (Lawson et al., 2018). Numerous strategies have been suggested to overcome these limitations, including less literal interpretation of results (Lawson et al., 2018), use of alternative clustering methods (Corander et al., 2008; Jombart et al., 2010), and implementation of models that account for specific demographic histories (Bradburd et al., 2018; Chen et al., 2007; François & Durand, 2010). Use of multiple data analysis strategies, including parametric and nonparametric approaches and testing multiple models with different underlying assumptions, is currently the best practice, but comes at the cost of ever-increasing computational burden.

A common (mis)interpretation of *STRUCTURE* plots is that assignment of one individual to multiple clusters is indicative of admixture (Kong & Kubatko, 2021). While it is true that mixed ancestry is expected in the case of hybridizing taxa, other demographic histories can also lead to assignment to multiple clusters. Notably, a history of isolation-by-distance (IBD) creates a pattern of continuous variation across populations, which violates the assumption that individuals can be sorted into K discrete populations (Frantz et al., 2009; Lawson et al., 2018). In the case of IBD, relatedness decays with increasing geographic distance. Intuitively, IBD is common in nature, as individuals are more likely to mate with individuals that are geographically proximal compared to those farther away (Meirmans, 2012). When IBD is a main feature of the data, clustering algorithms often assign individuals at either end of the sampled range to distinct clusters and infer individuals in the geographic center to have some degree of mixed ancestry (Frantz et al., 2009; Bradburd et al., 2018).

Discriminating IBD from admixture is an important first step for identifying and investigating hybridization in natural systems, which is a popular area of research within evolutionary biology as hybrid zones provide a window into the speciation process (Mallet et al., 2016; Payseur & Rieseberg, 2016). A number of models for hybrid zone dynamics have been proposed, but a common model for hybrid zone formation is secondary contact after allopatric divergence (Hewitt, 1988). In temperate zones, this pattern can be explained by isolation and divergence in separate refugia during ice ages, when glaciers covered much of the northern hemisphere, followed by range expansion and secondary contact when glaciers receded (Hewitt, 1988, 2000, 2004). This process is credited with hybrid zone formation across a wide range of taxa (Harrison & Larson, 2016; Toews & Brelsford, 2012). The outcome of hybridization after secondary contact depends on the fitness of recombinant genotypes in an environment (Moran et al., 2021). If there is no selection against admixed individuals, alleles can freely introgress into parental populations and the width of the zone is expected to increase with time, a model often referred to as *neutral diffusio*n (Hewitt, 1988). In contrast, selection against recombinant genotypes is common, and when balanced by dispersal of parentals into the zone, a stable, narrow cline referred to as a *tension zone* is formed (Barton & Hewitt, 1985). Tension zones are stable through time, but can shift geographically, as has been documented in many cases (Barton & Hewitt, 1985; Ryan et al., 2018; Taylor et al., 2014). In a special case of the tension zone model, hybridization occurs at an ecotone and admixed individuals are only fit at the ecotone, resulting in a contact zone that is stable both spatially and temporally (Hewitt, 1988). While secondary contact after range expansion from isolated refugia is a common biogeographic scenario, this process also generates continuous genetic variation within each expanding taxon. If there is no secondary contact, either because there is not another taxon to contact or because reproductive barriers are complete, genetic variation will be continuous and best described by IBD. Since IBD can result in genetic clustering patterns that resemble those expected under admixture, investigators must first rule out IBD before analyzing hybrid zones in more detail.

There are a few strategies for discriminating IBD from admixture. One approach is to incorporate geographic data during cluster inference, with the assumption that genetic relatedness decays with geographic distance. This strategy is implemented in *conStruct*, which accurately describes individuals that evolved under a history of IBD as belonging to one genetic cluster (Bradburd et al., 2018). A different strategy is to incorporate genotype data when identifying genetic clusters. At sites with fixed differences in diploid parental populations, a first filial generation (F1) hybrid inherits a different allele from each parent, and is thus heterozygous at each such site. In the event of backcrossing between an F1 and a parental individual, offspring are expected to be heterozygous at half of those sites. Inclusion of genotype frequencies during clustering is implemented in software such as *newhybrids*, which can distinguish between six genotypic classes (Anderson & Thompson, 2002).

Visualizations of hybrid index with interclass heterozygosity are called triangle plots, and can be used to identify hybrid classes and provide evidence for admixture (Fitzpatrick, 2012; Wiens & Colella, 2024). Triangle plots provide a simple, intuitive method for validating whether patterns of mixed ancestry inferred by *STRUCTURE* conform to expectations under a demographic history of admixture. Under a history of admixture, individuals inferred as having 50-50 ancestry by *STRUCTURE* (when K=2) should have a hybrid index of 0.5 and elevated interclass heterozygosity (1.0 in F1s, 0.5 in F2s and later filial generations) relative to parentals. In fact, under Hardy-Weinberg Equilibrium (HWE), it is impossible for admixture to produce an individual of 50-50 ancestry that has <0.5 interclass heterozygosity (Wiens & Colella, 2024). In contrast, there should be no relationship between hybrid index and interclass heterozygosity when admixture has not occurred and IBD is the main feature of the data (under such a history, “hybrid index” takes on a slightly different meaning, as it no longer represents actual admixture, but simply the proportion of ancestry matching one “parental” population). Range expansion out of a refugial population is a biogeographic scenario which generates a pattern of IBD, and in this special case, a negative linear relationship between “hybrid index” and interclass heterozygosity is expected, with interclass heterozygosity lower in more recently founded populations (Excoffier et al., 2009; Milá et al., 2000). If expansion is not recent, and barring differences in population sizes or selection regimes, interclass heterozygosity is expected to be constant across populations (Eckert et al., 2008).

Here, we examine the relationship between hybrid index and interclass heterozygosity under three demographic scenarios, and illustrate the utility of triangle plots for distinguishing between histories of admixture and IBD. We analyze simulated data using two popular methods for initial characterization of genetic variation, *STRUCTURE* and Principal Component Analysis (PCA). We show that these methods yield nearly identical results for genetic data simulated under different demographic histories, while triangle plots effectively distinguish between admixture and IBD. We then examine how *STRUCTURE*, PCA, and triangle plots change with increasing time since contact (measured in terms of generations) and how missing data can skew results. We also investigate how the allele frequency difference threshold (δ) for identifying ancestry-informative markers (AIMs) affects the relationship between hybrid index and interclass heterozygosity. We then incorporate geographic information during the genetic clustering process, as implemented in *conStruct*, and compare results. Finally, we discuss how triangle plots can be paired with clustering analyses for distinguishing between histories of admixture and IBD.

## Methods

### Simulations & Models

Forward-time genetic simulations were performed under a non-Wright-Fisher model in SLiM 3 (Haller & Messer, 2019). Three biogeographic scenarios were modeled: (1) neutral diffusion upon secondary contact (hereafter “ND”); (2) a parapatric contact zone upon secondary contact (i.e. geographically-bounded contact, hereafter “GBC”); and (3) isolation-by-distance due to geographic expansion from a single source population (hereafter “IBD”). Each model consisted of three phases (Fig 1). Phase I was the same for each, and modeled a common ancestor prior to divergence. Phase I lasted 10,000 generations and consisted of two populations with a high migration rate, such that on average 20% of individuals switched populations every generation. Phase II modeled allopatric divergence between two populations for 10,000 generations, during which no migration occurred. At this point in the IBD simulation, only one of the two populations was retained. Phase III lasted 20,000 generations and modeled range expansion under a linear, stepping-stone model of 21 populations, which provided enough time and space to capture the relevant outcomes of each simulation. The two (or one, in the IBD model) parental populations were positioned at the extreme ends of the “landscape”, and migration could only occur only between adjacent populations. Two migrants were randomly selected from each population, each generation, to move into each adjacent population, and migration only occurred from a population if it contained at least 200 eligible migrants (see below). In the GBC simulation, migration could occur into but not out of the central-most population, serving as a model of hybridization without introgression into parental populations, which is thought to be common in nature (Abbott et al., 2013; Barton & Hewitt, 1985, 1989; Buggs, 2007). With this approach, selection against hybrids outside of the ecotone is modeled without the addition of multiple parameters (e.g. selection coefficients, DMIs, additive/dominance effects of alleles, etc.) that would be required to explicitly simulate a tension zone or environmentally-dependent selection at an ecotone, and which would make comparison across models difficult.

**Figure 1.**
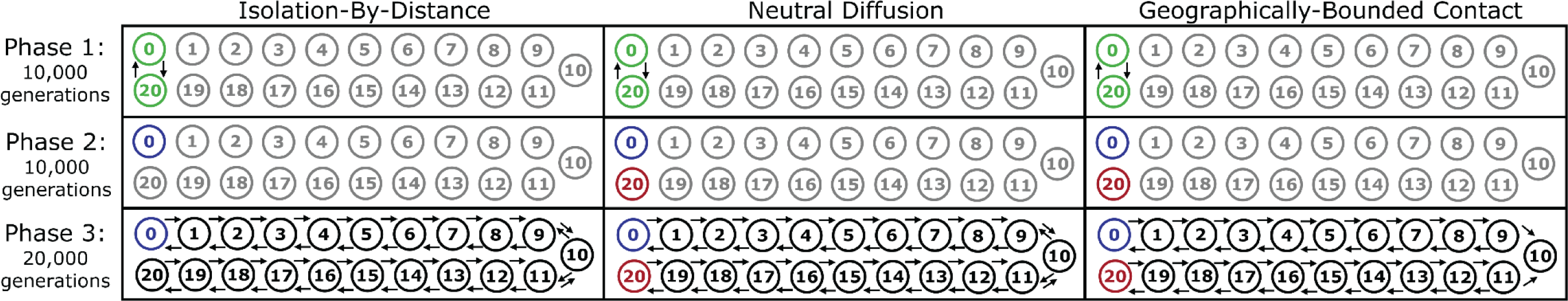
Schematic of the three phases of each simulation. In Phase I, individuals evolve in the green populations (p0 and p20) with a high migration rate and no individuals exist in the gray populations. In Phase II, migration between the parental populations p0 (blue) and p20 (red) ceases, and they evolve independently. In Phase III, migration occurs between all populations under a stepping-stone model.

Individuals had a lifespan of two generations and populations had a carrying capacity of 500 individuals. Only individuals that existed in the previous generation could reproduce, and all were hermaphroditic, such that every eligible individual reproduced with another randomly chosen individual in the same population. The number of offspring per mating followed a Poisson distribution (λ = 1.04). After all eligible individuals had a chance to reproduce, they were removed from the simulation, so that only offspring remained. Then, migrants were selected from among the offspring and moved to an adjacent population. Since individuals existed for two generations and the carrying capacity per population was 500, the maximum number of eligible migrants per population fluctuated around 250.

Each individual had a 100,000 bp diploid genome with a mutation rate of 10^-7^ and a uniform recombination rate of 10^-5^. All mutations were selectively neutral. Alleles that were fixed in a parental population at the end of Phase II, but which were not fixed at the end of Phase I, were identified and tracked through the end of the simulation, at which point their average frequency in each population was calculated.

During Phase III, 20 individuals were randomly sampled from each population every 200 generations. For sites which were polymorphic in the sample, each sampled individual’s genotype was output to a VCF file for downstream analyses. In the ND and GBC simulations, the first sample occurred when 20 of the 21 populations had at least 210 individuals, which effectively guaranteed that the first sample would occur within one to five generations after secondary contact. In the IBD simulation, the first sample occurred when every population had at least 20 individuals, which represented the earliest time point at which every population had enough individuals to take a sample. Due to the inherent stochasticity of the simulations, sampling did not occur in the exact same generation for each simulation, but sampled generations reflect equivalent evolutionary time points across simulations. This approach essentially standardizes sampling across simulations by taking the first sample as soon as each population has at least 20 individuals to sample. Subsequently, the generation at which each sample is recorded is based on the number of generations since the first sample.

To validate consistency of results, each simulation (ND, GBC, and IBD) was replicated 10 times. During each replicate, 10 separate, random samples were taken at generations 200, 1,000, and 10,000. For each subsample in each replicate, average hybrid index and interclass heterozygosity was computed for each population and visualized to describe the distribution of possible outcomes for each simulation. At each benchmark generation, we verified that average hybrid index and interclass heterozygosity in our full simulations followed expected distributions.

### Missing data

Unless otherwise mentioned, downstream analyses were performed with complete genotype matrices output from SLiM. Empirical datasets, however, usually contain some degree of missing data. Therefore, to characterize the effect of missing data on results, we randomly converted genotypes to missing data in the generation 1,000 sample of each simulation to create datasets with 10%, 30%, and 50% overall missing data. Missing data in empirical datasets is usually not distributed evenly across individuals. Often a few samples have more missing data than others, due to lower quality or quantity of input DNA, among other variables, leading to a right-skewed distribution of missing data across individuals (Rivera-Colón et al., 2021).

Therefore, we drew the percent of missing data per individual from a Beta distribution with α= 1.5 and β such that the chosen overall percent of missing data across individuals was met (Fig S5).

### Clustering approaches

For each sampled generation of each simulation, we ran five replicates of *STRUCTURE* at K=2 for 200,000 MCMC repetitions with a burn-in of 50,000 (Pritchard et al., 2000). We used the admixture model and did not use prior population information. We randomly selected one of the five replicates of each run for visualization. We conducted PCAs using the R package *adegenet*, retaining the first 10 principal components (PCs) and visualizing the first two (Jombart, 2008).

### Triangle plots

We used *triangulaR* to calculate hybrid index and interclass heterozygosity, and visualize triangle plots (Wiens & Colella, 2024). This approach relies on using samples from parental populations to identify ancestry-informative markers (AIMs), defined as loci with allele frequency differences above a chosen threshold between the parental populations (Rosenberg et al., 2003). For each sampled generation of each simulation, we calculated the hybrid index and interclass heterozygosity of each individual using AIMs with an allele frequency difference of δ=0.5 in the parental populations (or, in the IBD simulation, the parental population and the population furthest from it). To investigate how the allele frequency difference threshold influenced results, we also built triangle plots for generations 1,000 and 10,000 of each simulation by identifying AIMs with thresholds of 0.5, 0.75, and 1.

### Incorporating geography in cluster identification

Incorporating geographic information during genetic cluster inference provides a more accurate description of systems with continuous variation (Bradburd et al., 2018). We applied this method, as implemented in the R package *conStruct*, to the data from each simulation at generations 0, 200, 1,000, 10,000, and 19,000. Running *conStruct* using individual allele frequencies for the 420 sampled individuals (20 individuals from 21 populations) is not computationally feasible, therefore, we pooled individuals to estimate allele frequencies in each population. Geographic distance was calculated as the number of steps across the stepping-stone model (e.g. p10 is one step from p11 and two steps from p8). We cross validated models that included geography (spatial) against those that did not (nonspatial), using 80% of the dataset for training and 20% testing, for K=1-3 with eight replicates each for 1,000 MCMC iterations and the default burn-in. The spatial model performed best, so we also ran a longer MCMC chain (200,000 iterations with the default burn-in) for this model with five replicates at K=2.

## Results

### Simulation replicates and validation

Hybrid index and interclass heterozygosity of individuals in each population in the full ND, GBC, and IBD simulations followed the general trends based on ten replications each with ten random subsamples (Figs S1-S3). Almost all (97%) values of hybrid index and interclass heterozygosity estimated from the full simulations fell within the observed values from the replicate simulations. Concordance of these statistics at the sampled generations suggests that the full simulations, which were analyzed in more detail, are representative of the general outcomes of each model (ND, GBC, and IBD). Results of the full simulations are reported for three generations: 0 (first sample), 1,000, 10,000. The number of SNPs per sampled generation ranged from 767 to 1,934. Results are shown for every sampled generation in Supplementary Material 2.

### Clustering approaches

For every sampled generation in each simulation, the *STRUCTURE* algorithm inferred peripheral populations (i.e. p0 and p20) as separate, distinct clusters, with a gradient of mixed ancestry found in interim populations (Fig 2). Similarly, the first PC from each PCA explained differentiation between the peripheral populations, while the second PC explained differentiation of the central most populations from the peripheral populations (Fig 2). In every generation of the GBC model and in generation 0 of the ND model, individuals with mixed ancestry were restricted to the central-most population (p10). In the early stages of the IBD and ND models, the transition in inferred ancestry occurs over a few (∼1-6) populations, before widening into a broader cline involving more (∼9-12) populations, as seen in the *STRUCTURE* and PCAs plots (Fig 2).

**Figure 2.**
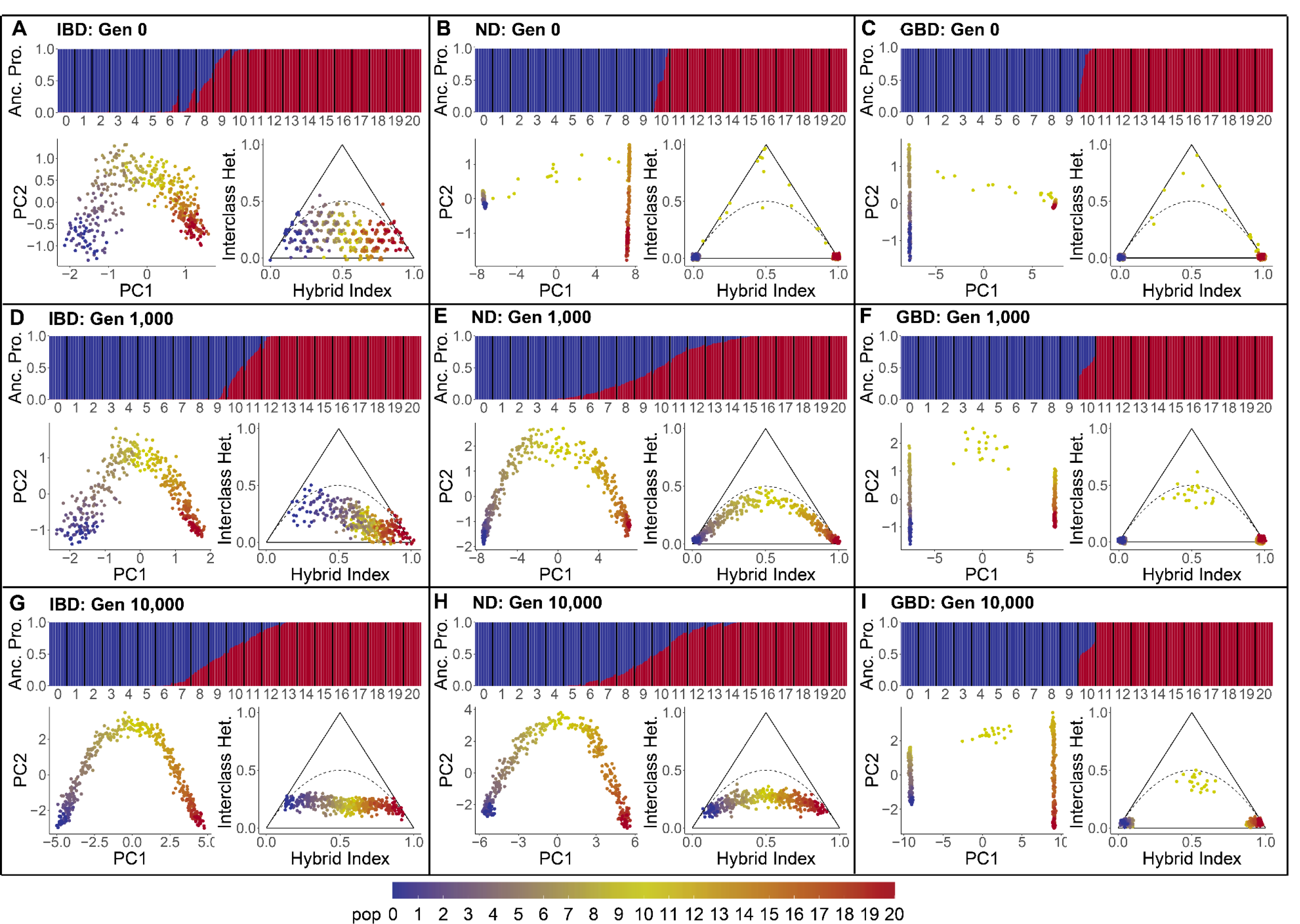
*STRUCTURE*, PCA, and triangle plots (allele frequency difference of δ=0.5) for simulations of Isolation-by-Distance (IBD), Neutral Diffusion (ND), and Geographically-Bounded Contact (GBC). In each panel, the *STRUCTURE* plot is on top, PCA bottom left, and triangle plot bottom right. Plots are shown for generations 0 **(A-C)**, 1,000, **(D-F)**, and 10,000 **(G-I)**. Color indicates the population of each sample on the PCA and triangle plots, and inferred ancestry proportions under K=2 on the *STRUCTURE* plot.

### Triangle plots

At generation 0 in the ND and GBC simulations, only individuals in p10 showed elevated interclass heterozygosity, with most individuals falling along the outer edges of the triangle, indicative of F1s and backcrossed individuals (Fig 2). In later generations of the GBC simulation, only individuals in p10 had elevated interclass heterozygosity. In later generations of the ND simulation, individuals in populations further from p10 display elevated interclass heterozygosity, with the peak in p10 decreasing over time. Notably, there were no individuals with intermediate ancestry (e.g. between 0.25 and 0.75) and low interclass heterozygosity (e.g. < 0.1) in any generation of the GBC or ND simulation. In contrast, there are many individuals in the IBD simulation with intermediate ancestry and low interclass heterozygosity through generation 10,000, and in those early generations there is a significant negative linear relationship between hybrid index and interclass heterozygosity (Table S1). Interclass heterozygosity in the IBD simulation is not elevated in the central populations (i.e. p9, p10, p11) until late in the simulation, sometime between generation 10,000 and 19,000 (Supplementary Material 2).

### Missing data

As expected, missing data strongly influenced PCA results (Fig 3). As the amount of missing data increased, individuals were plotted closer to the origin. This is evident across all models, especially when overall missing data was high (e.g. 50%). *STRUCTURE* plots were less affected by missing data, particularly in the ND and GBC simulations, as inferred individual ancestry proportions changed little as more missing data was added (Fig 3). Inferred ancestry proportions in the IBD simulation, however, were affected by missing data. Most notably, missing data sometimes shifted ancestry proportions in peripheral populations towards the genetic cluster of the other peripheral population. Missing data had little effect on triangle plots for the ND and GBC models, and a slight effect on the IBD model only when there was high overall missing data (Fig 3).

**Figure 3.**
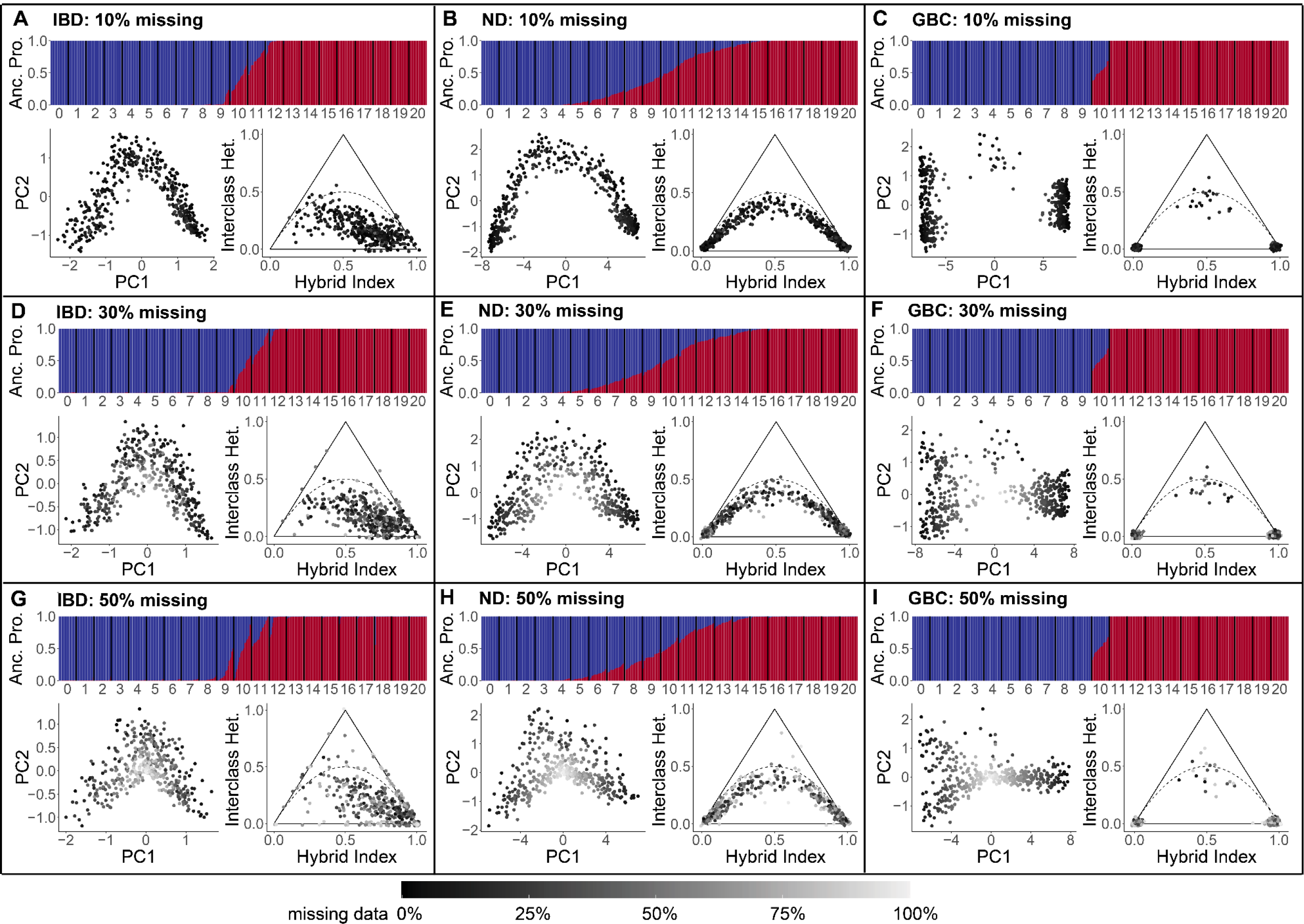
*STRUCTURE*, PCA, and triangle plots (allele frequency difference of δ=0.5) for simulations of Isolation-by-Distance (IBD), Neutral Diffusion (ND), and Geographically-Bounded Contact (GBC). In each panel, the *STRUCTURE* plot is on top, PCA bottom left, and triangle plot bottom right. All plots are for generation 1,000. Plots are shown for overall missing data percentages of 10% **(A-C)**, 30% **(D-F),** 50% **(G-I)**. Color indicates the percentage of missing data in each sample on the PCA and triangle plots, and inferred ancestry proportions under K=2 on the *STRUCTURE* plot.

### Allele frequency difference threshold and hybrid index

Increasing the allele frequency difference threshold between parental populations decreased the number of identified AIMs in every model (Figs 4 & 5). For the ND and GBC models, despite restricting calculation of hybrid index to fewer AIMs of more extreme allele frequency difference, the hybrid index and interclass heterozygosity of each individual changed little as the threshold increased (Fig 4 & 5). In contrast, the shape of the triangle plot for the IBD model changed dramatically as the threshold of allele frequency difference increased; not only did few sites pass higher thresholds, but individuals in parental populations were shifted toward the bottom corners of the plot and individuals in central populations were shifted toward the top middle.

**Figure 4.**
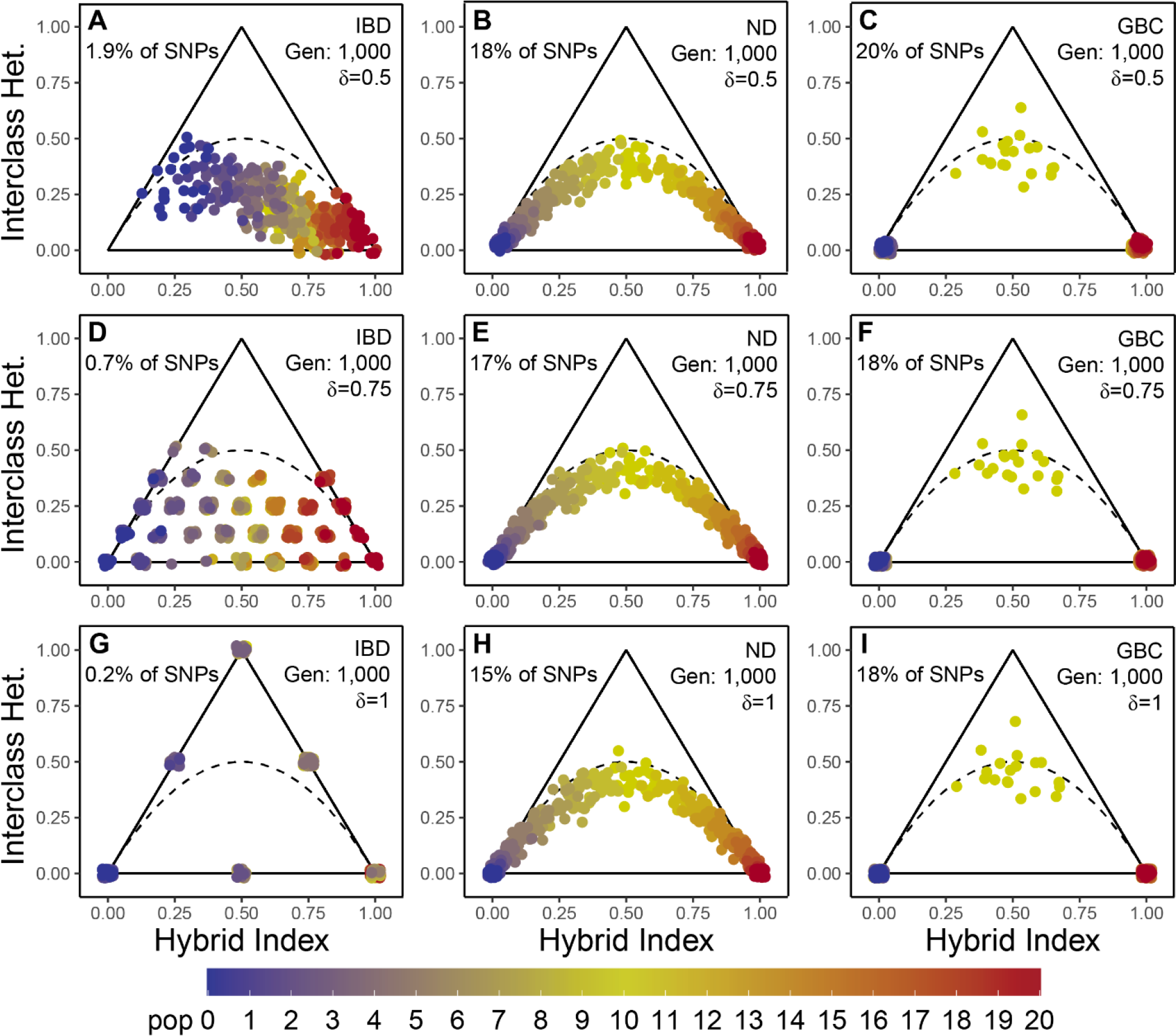
Triangle plots for simulations of Isolation-by-Distance (IBD), Neutral Diffusion (ND), and Geographically-Bounded Contact (GBC). All plots are for generation 1,000, and were built using AIMs identified under three allele frequency difference thresholds: δ=0.5 **(A-C)**, δ=0.75 **(D-F)**, and δ=1 **(G-I)**. Color indicates the population of each sample.

**Figure 5.**
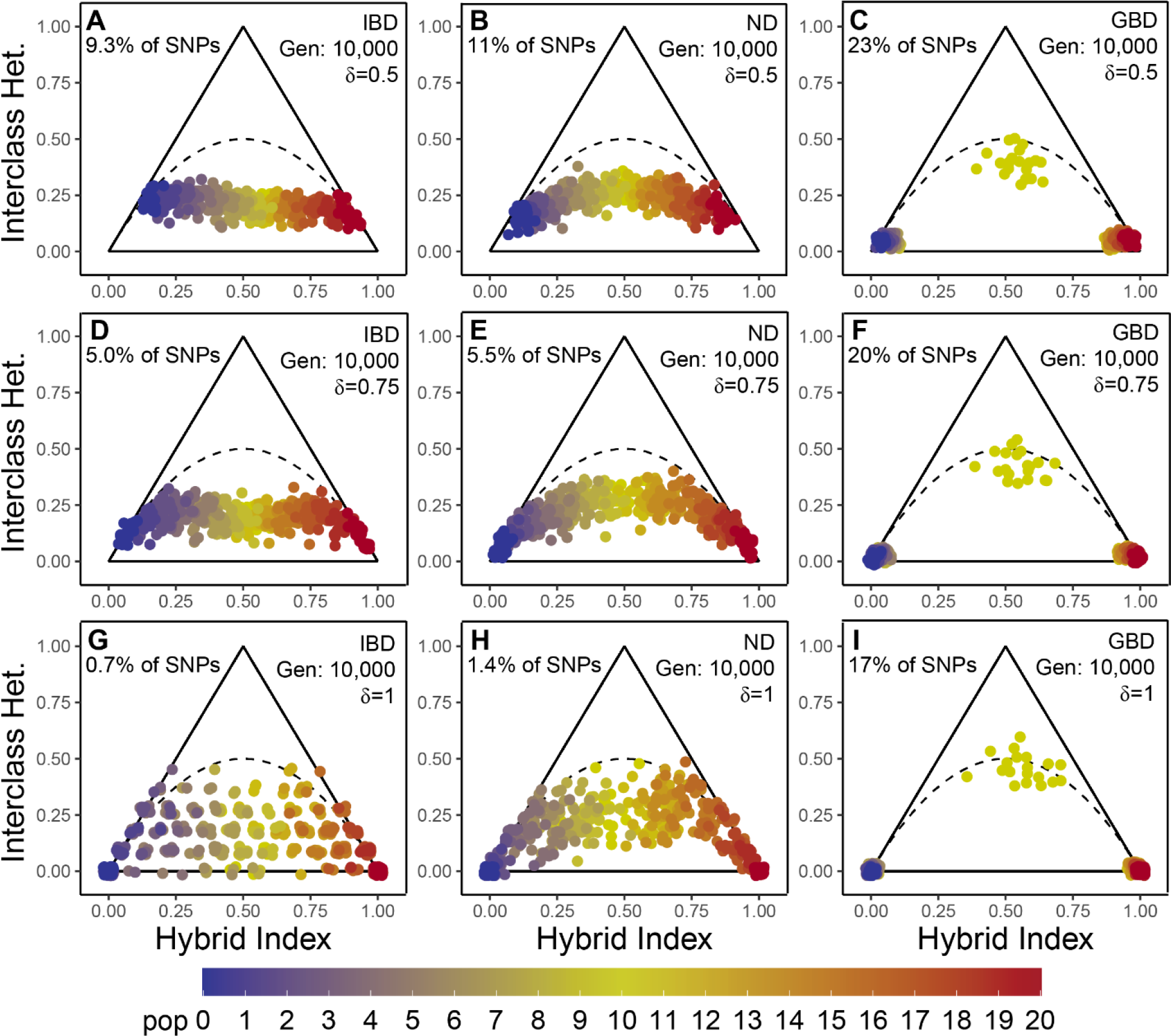
Triangle plots for simulations of Isolation-by-Distance (IBD), Neutral Diffusion (ND), and Geographically-Bounded Contact (GBC). All plots are for generation 10,000, and were built using AIMs identified under three allele frequency difference thresholds: δ=0.5 **(A-C)**, δ=0.75 **(D-F)**, and δ=1 **(G-I)**. Color indicates the population of each sample.

### Incorporating geography in cluster identification

Cross validation indicated that the spatial model was always preferred over the nonspatial model for every value of K in every generation and simulation (Fig S6). Under the spatial model at K=2 for every generation of the IBD simulation, all populations were assigned to the same cluster, indicating that the algorithm accurately infers no admixture (Figs S7 & S8). In contrast, admixture is accurately inferred in early generations of the ND simulation and all generations of the GBC simulation under the spatial model at K=2 in *conStruct* (Figs S7 & S8).

## Discussion

We use forward-time evolutionary simulations to show that histories of IBD and gene flow cannot be distinguished using common clustering approaches, such as those implemented by *STRUCTURE* or PCA. While more complex models exist for disentangling the relative roles of IBD and admixture in shaping genetic variation (Bradburd et al., 2018; François & Durand, 2010), many are not yet computationally feasible for large genomic datasets. Triangle plots, however, are a simple bioinformatic tool for analyzing population genetic datasets where admixture between two taxa is hypothesized. Building triangle plots as an initial step for exploratory analysis of population genetic data from potential hybrid zones will provide researchers with an intuitive visualization of genetic variation and guide further analyses and interpretation.

### STRUCTURE and PCA cannot distinguish admixture from IBD

*STRUCTURE* and PCA often comprise the first, exploratory steps for understanding genetic variation in a sample of individuals taken from wild populations, and influence the direction of further analyses (Bourgeois & Warren, 2021; Lou et al., 2021). Yet, such analyses fail to discriminate between two very different demographic histories: admixture versus IBD. At every sampled generation in our IBD simulation, *STRUCTURE* plots resemble what would be expected under a history of allopatric divergence and secondary contact (Fig 2). Further, PCA plots also exhibit similar patterns for data simulated under a history of IBD and neutral diffusion after secondary contact. Interpreting the results of *STRUCTURE* and PCA in isolation can lead to an incorrect understanding of the nature of genetic variation across geographic space when assignment to multiple clusters is taken as evidence for admixture.

### Triangle plots accurately distinguish admixture from IBD

Expanding exploratory analyses of genetic variation from natural populations to include triangle plots is a simple step that can distinguish between histories of IBD and gene flow (Fig 2). In cases of recent admixture, individuals at the geographic center of contact will always display elevated interclass heterozygosity, forming the expected signature triangle pattern (Fitzpatrick, 2012). Additionally, that pattern is robust to missing data (Fig 3), different allele frequency difference thresholds (Figs 4 & 5), and low sample sizes from the parental populations (Wiens & Colella, 2024).

A feature of triangle plots for data with a true history of admixture is that individuals with an intermediate hybrid index (≈0.5) do not have interclass heterozygosity lower than either parental population. Even using a relaxed allele frequency difference threshold of δ=0.5, individuals with a hybrid index of 0.5 maintained interclass heterozygosity around 0.5 for 1,000 generations in both admixture simulations, as expected under HWE (Fig 2, Wiens & Colella 2024). Reductions in interclass heterozygosity after 1,000 generations in the ND model can be explained by violations of HWE, such as genetic drift and accumulation of new mutations in the parental populations (discussed in more detail below). Triangle plots for data simulated under a history of IBD show a remarkably different pattern: interclass heterozygosity decreases with distance from the original source population, as expected for individuals at the leading edge of population expansions (Excoffier et al., 2009). With such few differences between p0 and p20 in early generations, this pattern is difficult to see, but notably, there are many individuals with intermediate ancestry and very low interclass heterozygosity, which is not expected in cases of true admixture.

### Incorporating geography improves cluster identification

Incorporating geography during genetic clustering using *conStruct* accurately classified populations in the IBD simulation as one cluster and identified two clusters and admixed individuals in the ND and GBC simulations. A major limitation of this method, however, is that it scales poorly with the number of individuals sequenced (Bradburd et al., 2018). Running *conStruct* on individual allele frequencies from 420 individuals was intractable, and thus we had to pool allele frequencies by population. This method still proved useful for distinguishing between continuous and discrete genetic variation, but limits investigation of ancestry proportions in individuals.

### Influence of the allele frequency difference threshold

Theoretically, the best allele frequency difference threshold for calculating hybrid index and interclass heterozygosity is δ=1, which equates to fixed differences between parental populations (Boecklen & Howard, 1997; Fitzpatrick, 2012). By using sites with fixed differences, there are concrete expectations for the relationship between hybrid index and interclass heterozygosity in various hybrid classes. In practice, however, lower allele frequency difference thresholds can provide estimations of hybrid index and interclass heterozygosity that are as accurate and more precise than those obtained with higher thresholds, because more AIMs pass lower thresholds (Wiens & Colella, 2024). Here, we find that early generation hybrids (e.g. F1s, backcrosses in generation 0 of the ND and GBC simulations) still appear as expected even with a relaxed allele frequency difference threshold of δ=0.5 (Fig 2). Further, there is virtually no difference in individual position within the triangle plots across different thresholds of allele frequency difference for the GBC simulations and the ND simulation at generation 1,000 (Fig 4).

Importantly, lowering the allele frequency difference threshold includes more sites compared to including only sites with fixed differences. In the IBD simulation at generation 1,000, for example, the expected pattern of decreasing interclass heterozygosity from the source population to newly founded populations (Excoffier et al., 2009) is apparent at a low allele frequency difference threshold of δ=0.5 (Fig 4). Since all of the simulated populations diverged relatively recently (1,000 generations ago), there are few sites with large differences in allele frequency between peripheral populations (p0 and p20), so increasing the threshold begins to obscure the true pattern. Further, by using only sites with fixed differences, parental populations by definition must have a interclass heterozygosity of 0, meaning that comparisons of interclass heterozygosity between those two populations is meaningless, and the expected signal of decreased interclass heterozygosity in recently expanded populations is lost. With progressively fewer sites meeting higher allele frequency difference thresholds, it also becomes more likely for individuals to have high interclass heterozygosity by chance alone, which could be misconstrued as evidence for an F1 hybrid. By relaxing the threshold of allele frequency difference between parental populations, the expected patterns of IBD can be distinguished from those of admixture without sacrificing identification of various hybrid classes.

### Inferring ancestry of parental populations

We used alleles which were fixed in each parental population at the end of Phase II as indicators of the true ancestry of each population. Calculating the average frequency of those alleles in each population at the end of the simulations (i.e. generation 19,000) shows that all populations are partially admixed as a result of neutral diffusion (Fig S4). Despite the presence of p0 alleles in p20 and vice versa, *STRUCTURE* does not infer admixture in the parental populations in generation 19,000 or any other generation (Supplementary Material 2). This highlights another shortcoming of *STRUCTURE*, which is that when alleles reach similar frequencies in all sampled populations, those alleles can no longer be used to infer admixture, even if they represent a history of introgression (Lawson et al., 2018). That limitation presents another reason to use a relaxed allele frequency difference threshold for triangle plots, because a low threshold does not restrict the hybrid index of the parental populations to zero or one. While literal interpretation of the *STRUCTURE* plot of generation 2,000 (and subsequent generations) of the ND simulation suggests no admixture in peripheral populations, the triangle plots shows that all populations are in fact partially admixed starting around generation 2,000 (Fig 2; Supplementary Material 2). Setting stricter thresholds for calculating hybrid index obscures this pattern, as it forces parental populations to have hybrid indices of zero or one, which may not reflect true ancestry proportions (Fig 5).

### How missing data affects clustering and triangle plots

Increasing missing data had the greatest effect on PCAs, compared to *STRUCTURE* and triangle plots (Fig 3). In general, missing data biased individuals towards a value of zero for both PC1 and PC2 (Fig 3), a result which has been observed elsewhere (Yi & Latch, 2022). In consequence, individuals with more missing data are pulled into the center of the plot, between p0 and p20. Admixed individuals are also expected to be intermediate between two populations on a PCA plot (Gompert & Buerkle, 2016; Patterson et al., 2006), so if missing data is not considered, that pattern may be mistaken as evidence for admixture.

*STRUCTURE* plots were less affected by high amounts of missing data, especially in the presence of admixture. Even with 50% overall missing data, inferred ancestry proportions are virtually the same as for the complete dataset (Fig 3). In contrast, *STRUCTURE* was more sensitive to high amounts of missing data in the IBD simulation, likely due to lesser divergence between peripheral populations. Ancestry proportions for individuals with high missing data in the IBD simulation could be dramatically different from ancestry proportions inferred with no missing data. Most notably, at 50% overall missing data, some individuals in the peripheral populations were identified as having a majority of ancestry from the wrong cluster (Fig 3).

While more robust to missing data than PCAs, our results highlight the need to interpret *STRUCTURE* output in light of missing data per individual.

Triangle plots proved robust to high degrees of missing data (Fig 3). The expected triangle pattern was consistent as missing data increased for admixture models. Missing data had a larger impact on hybrid indices and interclass heterozygosity for the IBD model, but even with the highest amount of missing data (50%), the pattern did not resemble expectations for a history of admixture. One reason that these triangle plots may be robust to missing data is the way that hybrid indices and heterozygosities are calculated in *triangulaR*. These metrics are calculated on a per-individual basis using only sites with non-missing data, and as such, they are not deflated by dividing by the total number of loci. It should be noted, however, that while triangle plots appear robust to missing data, this may only be true if missing data occurs randomly throughout the dataset. This is often not the case for empirical datasets, for which missing data may appear non-randomly for biological reasons, such as allele dropout in RADseq datasets, or as artifacts due to differences in sample quality or steps taken during quality filtering (Arnold et al., 2013; Huang & Knowles, 2016; Yi & Latch, 2022).

### With time, signatures of IBD and neutral diffusion become similar

Under Hardy-Weinberg equilibrium, it is not possible for an admixed individual to have a hybrid index of 0.5 and interclass heterozygosity less than 0.5 (Wiens & Colella, 2024). Yet, there is an observed decrease in interclass heterozygosity over time in the ND simulation (Fig 2).

Two mechanisms can account for this, both of which violate the assumptions of HWE. The first is random loss of alleles in heavily recombined populations through genetic drift, which is expected over time (Fitzpatrick, 2012; Milne & Abbott, 2008). If there is an equal chance of fixation of either parental allele in the recombined population, hybrid index will stay around 0.5 while interclass heterozygosity decreases. Another mechanism that could explain reduced interclass heterozygosity in intermediate individuals is mutation in the parental populations. If a new allele arises in one parental population after admixture begins at the contact zone, and that allele reaches high frequency or fixation, then that locus will be included in calculations of hybrid index and interclass heterozygosity. Until that allele has time to reach the center of the contact zone, admixed individuals will remain homozygous at that locus. Assuming equivalent mutation and substitution rates in the parental populations, hybrid index will stay around 0.5, but interclass heterozygosity will decrease.

For either of these mechanisms to noticeably decrease interclass heterozygosity in admixed individuals, a relatively large amount of time is required. For at least 1,000 generations after initial contact in the ND model, observed combinations of hybrid index and interclass heterozygosity follow HWE expectations. By 19,000 generations after contact, triangle plots for the ND and IBD simulations look remarkably similar, with only a small peak of interclass heterozygosity in the center populations (Supplementary Material 2). In the IBD simulation, that peak was not present in previous generations, which instead showed a pattern of decreasing interclass heterozygosity from population 0 to 20. This new peak can be explained by enough time for divergence of peripheral populations and for gene flow to bring newly arisen alleles to the center population.

While triangle plots are not able to distinguish histories of IBD and hybridization after 10,000 generations in these simulations, they do remarkably well up until that point. One caveat is that the number of generations for which these histories are distinguishable will depend on other factors, including mutation rate, migration rate, and natural selection. Higher migration rates will result in the observed joint distributions of hybrid index and interclass heterozygosity being reached in fewer generations, as well as fewer differences between the parental populations. Only one form of selection is simulated here, but other forms of selection against recombinants, such as accumulation of Dobzhansky-Muller incompatibilities (DMIs) in late generation hybrids, can also prevent the homogenization of parental populations (Christe et al., 2016; Cronemberger et al., 2020; Lindtke et al., 2012).

### Future directions

We simulated data under two simple models of hybrid zone dynamics, one in which there is no selection for or against admixture (ND) and another in which parentals can disperse into the contact zone but admixed individuals cannot disperse out of it (GBC). Further work is necessary to investigate the expected joint distribution of hybrid index and interclass heterozygosity under more complex models. For example, selection against late generation hybrids (possible through DMIs), pre-mating barriers to reproduction (e.g. conspecific recognition), heterosis, hybrid sterility/inviability in the heterogametic sex (i.e. Haldane’s rule), and unequal parental population sizes are expected to shift the observed joint distribution of hybrid index and interclass heterozygosity (Barton, 2001; Moran et al., 2021).

Triangle plots can distinguish between many classes of hybrids (F1s, backcrosses, etc.), but hybrid index and interclass heterozygosity alone cannot distinguish between F2s, F3s, and future such crosses under the assumptions of HWE (Gompert & Buerkle, 2016, Wiens & Colella, 2024). Development of a third metric to be used jointly with hybrid index and interclass heterozygosity for identifying such classes would be useful. For example, if recombination rates for focal taxa are known, it should be possible to use average ancestry tract lengths across the genome to infer the number of generations since hybridization between two parental individuals, and thus distinguish F2s from F3s and further crosses. Such an approach has been taken to model the timing and number of admixture pulses in contact zones (Corbett-Detig & Nielsen, 2017), and in theory could be extended to develop a metric that would help distinguish between hybrid classes.

## Acknowledgements

We thank Marlon Cobos, Lucas DeCicco, and Devon DeRaad for insightful discussions and early editorial feedback on the manuscript. This work was supported by the HPC facilities operated by the Center for Research Computing at the University of Kansas. This work was partially funded by National Science Foundation award to JPC (NSF#2100955).

## Data accessibility

SLiM, R, and Bash scripts used for simulating and analyzing genetic data, as well as raw SNP datasets for each simulation, are available on GitHub at https://github.com/omys-omics/IBDvsAdmixture.

## Author contributions

BJW conceptualized the study, simulated and analyzed the data, wrote the first version of the manuscript, and generated the figures. JPC funded the project and provided editorial feedback.

## References

1. Abbott, R., Albach, D., Ansell, S., Arntzen, J. W., Baird, S. J. E., Bierne, N., Boughman, J., Brelsford, A., Buerkle, C. A., Buggs, R., Butlin, R. K., Dieckmann, U., Eroukhmanoff, F., Grill, A., Cahan, S. H., Hermansen, J. S., Hewitt, G., Hudson, A. G., Jiggins, C., … Zinner, D. (2013). Hybridization and speciation. Journal of Evolutionary Biology, 26(2), 229–246. 10.1111/j.1420-9101.2012.02599.x

2. Alexander, D. H., Novembre, J., & Lange, K. (2009). Fast model-based estimation of ancestry in unrelated individuals. Genome Research, 19(9), 1655–1664. 10.1101/gr.094052.109

3. Anderson, E. C., & Thompson, E. A. (2002). A Model-Based Method for Identifying Species Hybrids Using Multilocus Genetic Data. Genetics, 160(3), 1217–1229. 10.1093/genetics/160.3.1217

4. Arnold, B., Corbett-Detig, R. B., Hartl, D., & Bomblies, K. (2013). RADseq underestimates diversity and introduces genealogical biases due to nonrandom haplotype sampling. Molecular Ecology, 22(11), 3179–3190. 10.1111/mec.12276

5. Barton, N. H. (2001). The role of hybridization in evolution. Molecular Ecology, 10, 551–568.

6. Barton, N. H., & Hewitt, G. M. (1985). Analysis of Hybrid Zones. Annual Review of Ecology and Systematics, 16(1), 113–148. 10.1146/annurev.es.16.110185.000553

7. Barton, N. H., & Hewitt, G. M. (1989). Adaptation, speciation and hybrid zones. Nature, 341(6242), Article 6242. 10.1038/341497a0

8. Beugin, M.-P., Gayet, T., Pontier, D., Devillard, S., & Jombart, T. (2018). A fast likelihood solution to the genetic clustering problem. Methods in Ecology and Evolution, 9(4), 1006–1016. 10.1111/2041-210X.12968

9. Boecklen, W. J., & Howard, D. J. (1997). Genetic Analysis of Hybrid Zones: Numbers of Markers and Power of Resolution. Ecology, 78(8), 2611–2616. 10.1890/0012-9658(1997)078[2611:GAOHZN]2.0.CO;2

10. Bourgeois, Y. X. C., & Warren, B. H. (2021). An overview of current population genomics methods for the analysis of whole-genome resequencing data in eukaryotes. Molecular Ecology, 30(23), 6036–6071. 10.1111/mec.15989

11. Bradburd, G. S., Coop, G. M., & Ralph, P. L. (2018). Inferring Continuous and Discrete Population Genetic Structure Across Space. Genetics, 210(1), 33–52. 10.1534/genetics.118.301333

12. Buggs, R. J. A. (2007). Empirical study of hybrid zone movement. Heredity, 99(3), Article 3. 10.1038/sj.hdy.6800997

13. Chen, C., Durand, E., Forbes, F., & François, O. (2007). Bayesian clustering algorithms ascertaining spatial population structure: A new computer program and a comparison study. Molecular Ecology Notes, 7(5), 747–756. 10.1111/j.1471-8286.2007.01769.x

14. Christe, C., Stölting, K. N., Bresadola, L., Fussi, B., Heinze, B., Wegmann, D., & Lexer, C. (2016). Selection against recombinant hybrids maintains reproductive isolation in hybridizing Populus species despite F1 fertility and recurrent gene flow. Molecular Ecology, 25(11), 2482–2498. 10.1111/mec.13587

15. Corander, J., & Marttinen, P. (2006). Bayesian identification of admixture events using multilocus molecular markers. Molecular Ecology, 15(10), 2833–2843. 10.1111/j.1365-294X.2006.02994.x

16. Corander, J., Marttinen, P., Sirén, J., & Tang, J. (2008). Enhanced Bayesian modelling in BAPS software for learning genetic structures of populations. BMC Bioinformatics, 9(1), 539. 10.1186/1471-2105-9-539

17. Corbett-Detig, R., & Nielsen, R. (2017). A Hidden Markov Model Approach for Simultaneously Estimating Local Ancestry and Admixture Time Using Next Generation Sequence Data in Samples of Arbitrary Ploidy. PLOS Genetics, 13(1), e1006529. 10.1371/journal.pgen.1006529

18. Cronemberger, Á. A., Aleixo, A., Mikkelsen, E. K., & Weir, J. T. (2020). Postzygotic isolation drives genomic speciation between highly cryptic Hypocnemis antbirds from Amazonia. Evolution, 74(11), 2512–2525. 10.1111/evo.14103

19. Eckert, C. G., Samis, K. E., & Lougheed, S. C. (2008). Genetic variation across species’ geographical ranges: The central–marginal hypothesis and beyond. Molecular Ecology, 17(5), 1170–1188. 10.1111/j.1365-294X.2007.03659.x

20. Excoffier, L., Foll, M., & Petit, R. J. (2009). Genetic Consequences of Range Expansions. Annual Review of Ecology, Evolution, and Systematics, 40(1), 481–501. 10.1146/annurev.ecolsys.39.110707.173414

21. Fitzpatrick, B. M. (2012). Estimating ancestry and heterozygosity of hybrids using molecular markers. BMC Evolutionary Biology, 12(1), 131. 10.1186/1471-2148-12-131

22. François, O., & Durand, E. (2010). Spatially explicit Bayesian clustering models in population genetics. Molecular Ecology Resources, 10(5), 773–784. 10.1111/j.1755-0998.2010.02868.x

23. Frantz, A. C., Cellina, S., Krier, A., Schley, L., & Burke, T. (2009). Using spatial Bayesian methods to determine the genetic structure of a continuously distributed population: Clusters or isolation by distance? Journal of Applied Ecology, 46(2), 493–505. 10.1111/j.1365-2664.2008.01606.x

24. Gao, H., Williamson, S., & Bustamante, C. D. (2007). A Markov Chain Monte Carlo Approach for Joint Inference of Population Structure and Inbreeding Rates From Multilocus Genotype Data. Genetics, 176(3), 1635–1651. 10.1534/genetics.107.072371

25. Gompert, Z., & Buerkle, C. A. (2016). What, if anything, are hybrids: Enduring truths and challenges associated with population structure and gene flow. Evolutionary Applications, 9(7), 909–923. 10.1111/eva.12380

26. Guillot, G., Estoup, A., Mortier, F., & Cosson, J. F. (2005). A Spatial Statistical Model for Landscape Genetics. Genetics, 170(3), 1261–1280. 10.1534/genetics.104.033803

27. Harrison, R. G., & Larson, E. L. (2016). Heterogeneous genome divergence, differential introgression, and the origin and structure of hybrid zones. Molecular Ecology, 25(11), 2454–2466. 10.1111/mec.13582

28. Hewitt, G. (2000). The genetic legacy of the Quaternary ice ages. Nature, 405(6789), Article 6789. 10.1038/35016000

29. Hewitt, G. M. (1988). Hybrid zones-natural laboratories for evolutionary studies. Trends in Ecology & Evolution, 3(7), 158–167. 10.1016/0169-5347(88)90033-X

30. Hewitt, G. M. (2004). Genetic consequences of climatic oscillations in the Quaternary. Philosophical Transactions of the Royal Society of London. Series B: Biological Sciences, 359(1442), 183–195. 10.1098/rstb.2003.1388

31. Huang, H., & Knowles, L. L. (2016). Unforeseen Consequences of Excluding Missing Data from Next-Generation Sequences: Simulation Study of RAD Sequences. Systematic Biology, 65(3), 357–365. 10.1093/sysbio/syu046

32. Huelsenbeck, J. P., & Andolfatto, P. (2007). Inference of Population Structure Under a Dirichlet Process Model. Genetics, 175(4), 1787–1802. 10.1534/genetics.106.061317

33. Jombart, T. (2008). adegenet: A R package for the multivariate analysis of genetic markers. Bioinformatics, 24(11), 1403–1405. 10.1093/bioinformatics/btn129

34. Jombart, T., Devillard, S., & Balloux, F. (2010). Discriminant analysis of principal components: A new method for the analysis of genetically structured populations. BMC Genetics, 11(1), 94. 10.1186/1471-2156-11-94

35. Kong, S., & Kubatko, L. S. (2021). Comparative Performance of Popular Methods for Hybrid Detection using Genomic Data. Systematic Biology, 70(5), 891–907. 10.1093/sysbio/syaa092

36. Latch, E. K., Dharmarajan, G., Glaubitz, J. C., & Rhodes, O. E. (2006). Relative performance of Bayesian clustering software for inferringpopulation substructure and individual assignment at low levels of population differentiation. Conservation Genetics, 7(2), 295–302. 10.1007/s10592-005-9098-1

37. Lawson, D. J., van Dorp, L., & Falush, D. (2018). A tutorial on how not to over-interpret STRUCTURE and ADMIXTURE bar plots. Nature Communications, 9(1), Article 1. 10.1038/s41467-018-05257-7

38. Lindtke, D., Buerkle, C. A., Barbará, T., Heinze, B., Castiglione, S., Bartha, D., & Lexer, C. (2012). Recombinant hybrids retain heterozygosity at many loci: New insights into the genomics of reproductive isolation in Populus. Molecular Ecology, 21(20), 5042–5058. 10.1111/j.1365-294X.2012.05744.x

39. Lou, R. N., Jacobs, A., Wilder, A. P., & Therkildsen, N. O. (2021). A beginner’s guide to low- coverage whole genome sequencing for population genomics. Molecular Ecology, 30(23), 5966–5993. 10.1111/mec.16077

40. Mallet, J., Besansky, N., & Hahn, M. W. (2016). How reticulated are species? BioEssays, 38(2), 140–149. 10.1002/bies.201500149

41. Meirmans, P. G. (2012). The trouble with isolation by distance. Molecular Ecology, 21(12), 2839–2846. 10.1111/j.1365-294X.2012.05578.x

42. Milá, B., Girman, D. J., Kimura, M., & Smith, T. B. (2000). Genetic evidence for the effect of a postglacial population expansion on the phylogeography of a North American songbird. Proceedings of the Royal Society B: Biological Sciences, 267(1447), 1033–1040.

43. Milne, R. I., & Abbott, R. J. (2008). Reproductive isolation among two interfertile Rhododendron species: Low frequency of post-F1 hybrid genotypes in alpine hybrid zones. Molecular Ecology, 17(4), 1108–1121. 10.1111/j.1365-294X.2007.03643.x

44. Moran, B. M., Payne, C., Langdon, Q., Powell, D. L., Brandvain, Y., & Schumer, M. (2021). The genomic consequences of hybridization. eLife, 10. 10.7554/ELIFE.69016

45. Novembre, J. (2016). Pritchard, Stephens, and Donnelly on Population Structure. Genetics, 204(2), 391–393. 10.1534/genetics.116.195164

46. Novembre, J., & Stephens, M. (2008). Interpreting principal component analyses of spatial population genetic variation. Nature Genetics, 40(5), Article 5. 10.1038/ng.139

47. Patterson, N., Price, A. L., & Reich, D. (2006). Population Structure and Eigenanalysis. PLOS Genetics, 2(12), e190. 10.1371/journal.pgen.0020190

48. Payseur, B. A., & Rieseberg, L. H. (2016). A genomic perspective on hybridization and speciation. Molecular Ecology, 25(11), 2337–2360. 10.1111/mec.13557

49. Pickrell, J., & Pritchard, J. (2012). Inference of population splits and mixtures from genome- wide allele frequency data. Nature Precedings, 1–1. 10.1038/npre.2012.6956.1

50. Pritchard, J. K., Stephens, M., & Donnelly, P. (2000). Inference of Population Structure Using Multilocus Genotype Data. Genetics, 155(2), 945–959.

51. Puechmaille, S. J. (2016). The program structure does not reliably recover the correct population structure when sampling is uneven: Subsampling and new estimators alleviate the problem. Molecular Ecology Resources, 16(3), 608–627. 10.1111/1755-0998.12512

52. Rivera-Colón, A. G., Rochette, N. C., & Catchen, J. M. (2021). Simulation with RADinitio improves RADseq experimental design and sheds light on sources of missing data. Molecular Ecology Resources, 21(2), 363–378. 10.1111/1755-0998.13163

53. Rosenberg, N. A., Li, L. M., Ward, R., & Pritchard, J. K. (2003). Informativeness of Genetic Markers for Inference of Ancestry*. The American Journal of Human Genetics, 73(6), 1402–1422. 10.1086/380416

54. Ryan, S. F., Deines, J. M., Scriber, J. M., Pfrender, M. E., Jones, S. E., Emrich, S. J., & Hellmann, J. J. (2018). Climate-mediated hybrid zone movement revealed with genomics, museum collection, and simulation modeling. Proceedings of the National Academy of Sciences, 115(10), E2284–E2291. 10.1073/pnas.1714950115

55. Tang, H., Peng, J., Wang, P., & Risch, N. J. (2005). Estimation of individual admixture: Analytical and study design considerations. Genetic Epidemiology, 28(4), 289–301. 10.1002/gepi.20064

56. Taylor, S. A., White, T. A., Hochachka, W. M., Ferretti, V., Curry, R. L., & Lovette, I. (2014). Climate-mediated movement of an avian hybrid zone. Current Biology: CB, 24(6), 671– 676. 10.1016/j.cub.2014.01.069

57. Toews, D. P. L., & Brelsford, A. (2012). The biogeography of mitochondrial and nuclear discordance in animals. Molecular Ecology, 21(16), 3907–3930. 10.1111/j.1365-294X.2012.05664.x

58. Wang, J. (2017). The computer program structure for assigning individuals to populations: Easy to use but easier to misuse. Molecular Ecology Resources, 17(5), 981–990. 10.1111/1755-0998.12650

59. Wiens, B. J., & Colella, J. P. (2024). triangulaR: an R package for identifying AIMs and building triangle plots using SNP data from hybrid zones. bioRxiv, 2024-03.

60. Wilson, G. A., & Rannala, B. (2003). Bayesian Inference of Recent Migration Rates Using Multilocus Genotypes. Genetics, 163(3), 1177–1191. 10.1093/genetics/163.3.1177

61. Yi, X., & Latch, E. K. (2022). Nonrandom missing data can bias Principal Component Analysis inference of population genetic structure. Molecular Ecology Resources, 22(2), 602–611. 10.1111/1755-0998.13498

