## Supplementary Material 1 for "That’s not a Hybrid: How to Distinguish Patterns of Admixture and Isolation-by-Distance"

Ben J. Wiens & Jocelyn P. Colella

**Table of Contents:**

| ***conStruct* analyses** | Page 2 |
| --- | --- |
| **Supplemental Figures** | Page 3-9 |
| **Supplemental Table** | Page 10 |
| **All structure and triangle plots** | Supplementary Material 2 |

*conStruct analyses*

Cross validation indicates that the spatial model in *conStruct* is preferred over the nonspatial model for every value of K in every generation of each simulation (Fig S6). Under the spatial model, K=1 generally has the highest likelihood for the IBD simulation, although the 95% confidence intervals for the likelihood of K=1 and K=2 overlap in each generation except generation 10,000. In early generations of ND simulation, K=1 has a much lower likelihood than K=2 until generation 10,000, at which point the likelihoods of these values of K are very similar. In the GBC simulation, the likelihood of K=1 is always much lower than the likelihood of K=2.

Comparing results across early generations of the simulations under a spatial model and assuming K=2 shows that *conStruct* is able to distinguish between patterns of continuous variation, a feature of the IBD simulation, and discrete variation, features of the ND and GBC models (Figs S7 & S8). The second genetic cluster (or “layer”, to use the terms of the authors of *conStruct*) contributes little to the overall covariance in the early generations of the IBD simulation, indicating that the algorithm accurately infers essentially zero no admixture (Fig S7). In contrast, both layers contribute equally to the overall covariance in early generations (0, 200, 1,000) of the ND model and all generations of the GBC model, indicating the presence of two discrete clusters, which can be seen to be mixing when viewing the plot of inferred ancestry proportions in each population (Fig S8). In late generations (10,000 and 19,000) of the ND model, only one layer contributes to the overall covariance (Fig S7).

Incorporating geographic data during genetic clustering has a strong advantage over methods which only use genetic data, especially when trying to discriminate between continuous variation, a pattern which can be generated by IBD, and discrete variation. This method consistently identifies continuous variation in our IBD simulation and describes the populations as belonging to a single cluster, a desirable result as it prevents misinterpretation of continuous variation as evidence of admixture. Despite the advantages of *conStruct* over *STRUCTURE* and similar genetic clustering methods, there are some drawbacks to *conStruct*. There is a practical drawback, as the algorithm currently scales poorly with increased sampling. Allele frequencies can be pooled for each population to facilitate computational efficacy, but this approach limits interpretation of results. Specifically, individual admixture proportions are unknown when data are pooled by population. As such, it would be not possible to distinguish a population containing only 50-50 hybrids from a population composed of an equal number of each parental taxa. When admixture is identified with *conStruct* for pooled populations, follow up analyses on individual allele frequencies will still be necessary. Another drawback is that *conStruct* does not identify admixture for late generations of the ND simulation. A likely reason for this is that with enough time, neutral diffusion of alleles across parental boundaries will result in continuous variation, reminiscent of IBD. This problem is not restricted to *conStruct* though, as *STRUCTURE*, PCA, and triangle plots for generation 19,000 of the IBD and ND simulations all look very similar (Fig 2). To discriminate IBD from neutral diffusion when contact is thought to have occurred long ago, more detailed analyses will be needed.

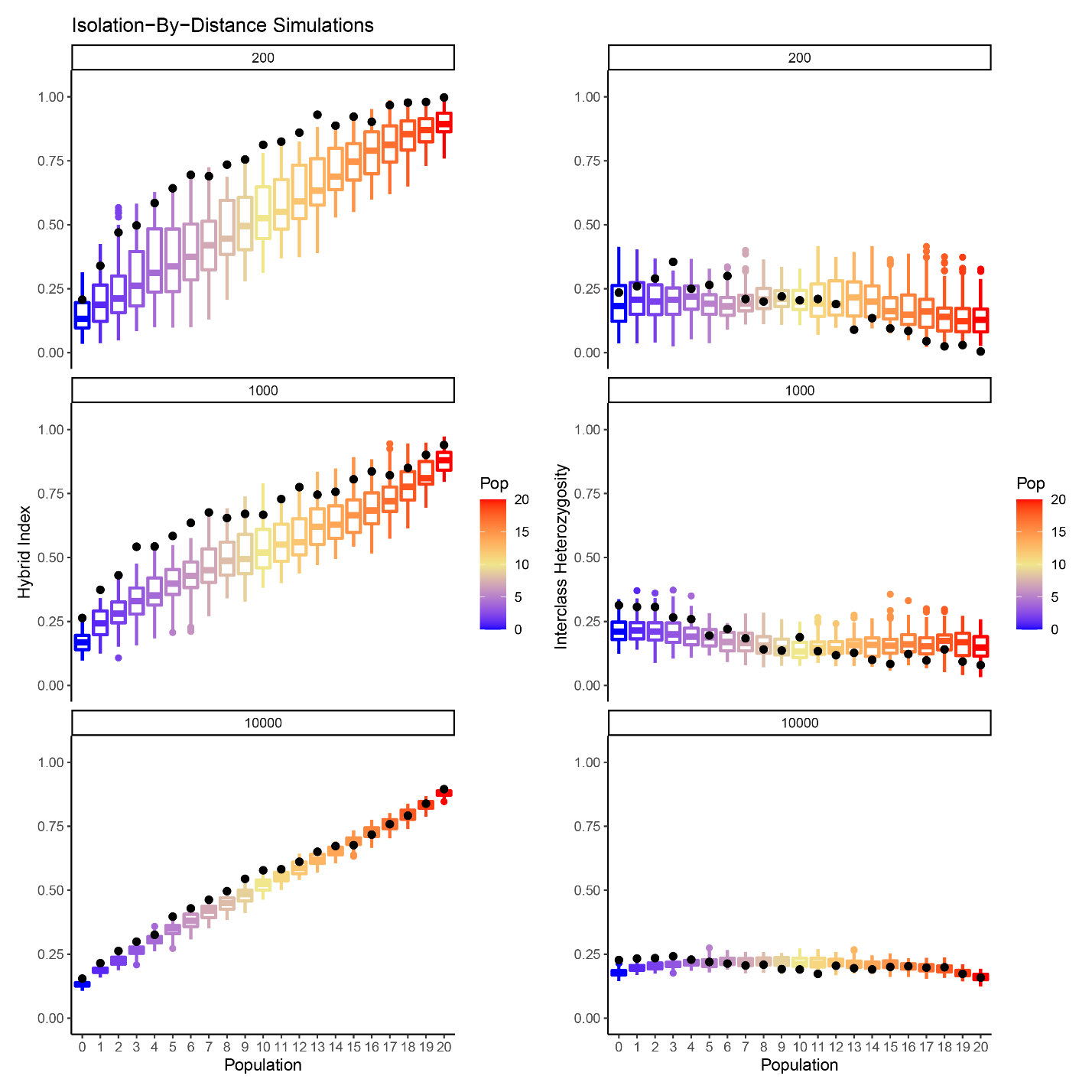

**Fig S1.** Distribution of hybrid index **(left)** and interclass heterozygosity **(right)** in each population at generations 200, 1,000, and 10,000 across ten replicates of the IBD simulation. Values for the full simulations are shown as black dots.

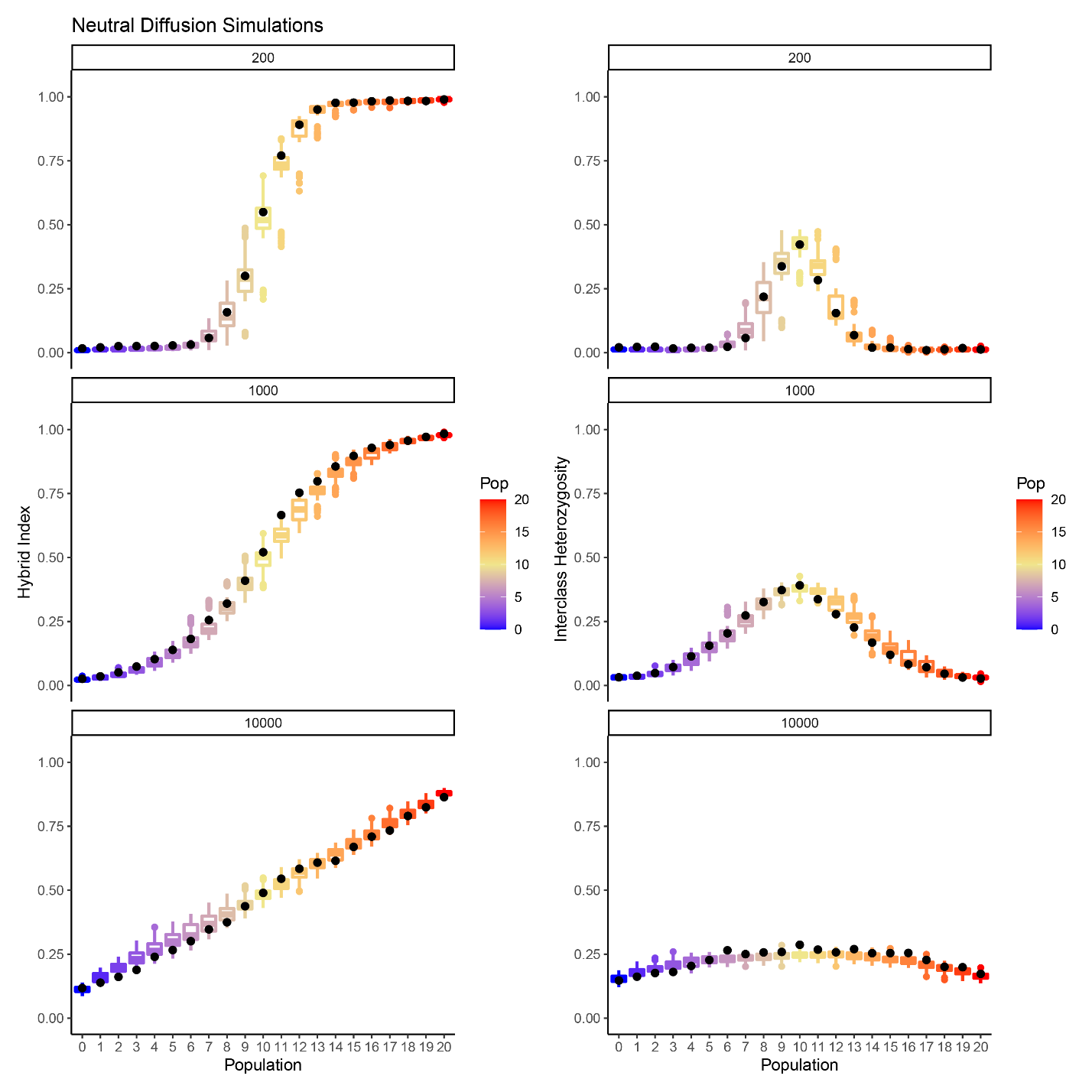

**Fig S2.** Distribution of hybrid index **(left)** and interclass heterozygosity **(right)** in each population at generations 200, 1,000, and 10,000 across ten replicates of the ND simulation. Values for the full simulations are shown as black dots.

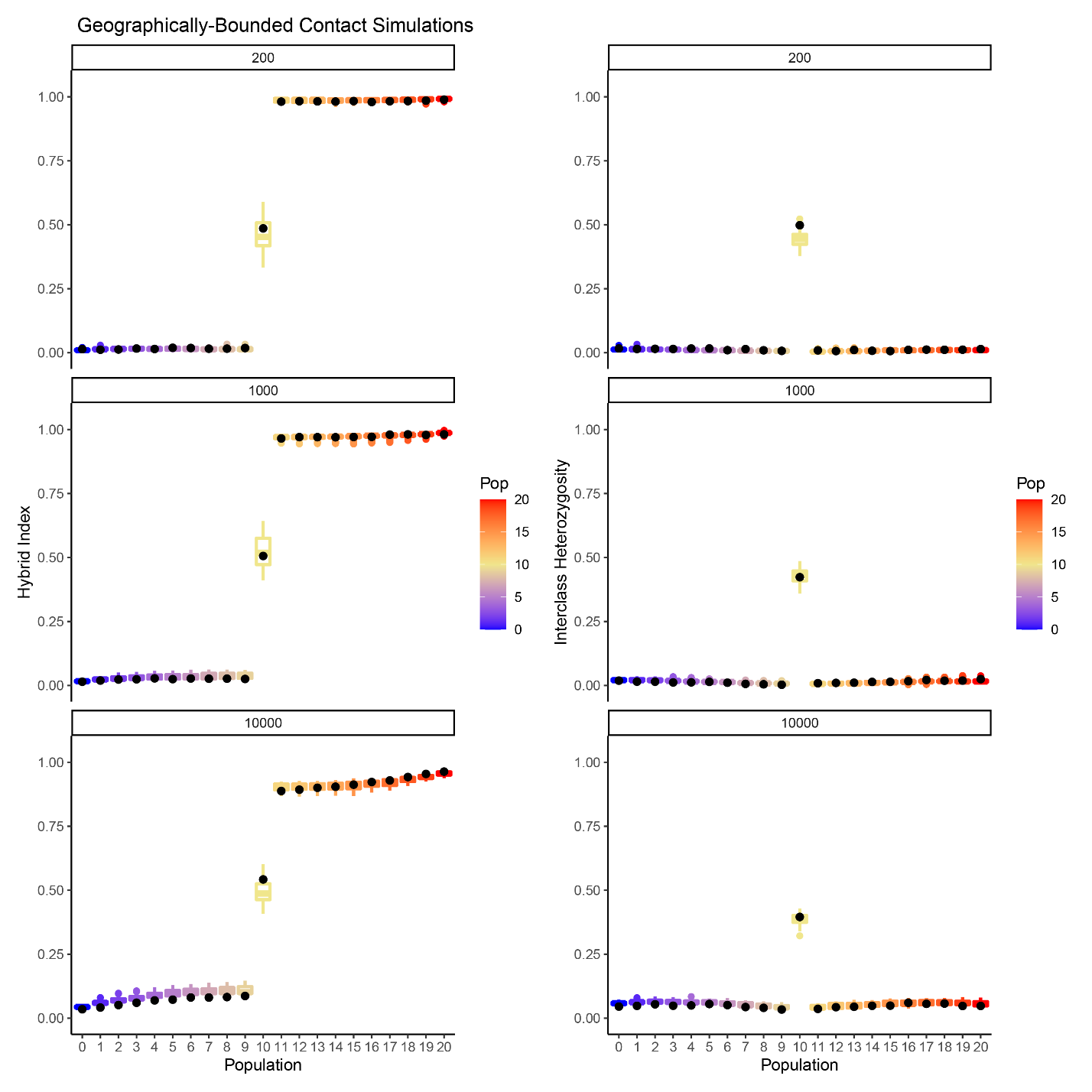

**Fig S3.** Distribution of hybrid index **(left)** and interclass heterozygosity **(right)** in each population at generations 200, 1,000, and 10,000 across ten replicates of the GBC simulation. Values for the full simulations are shown as black dots.

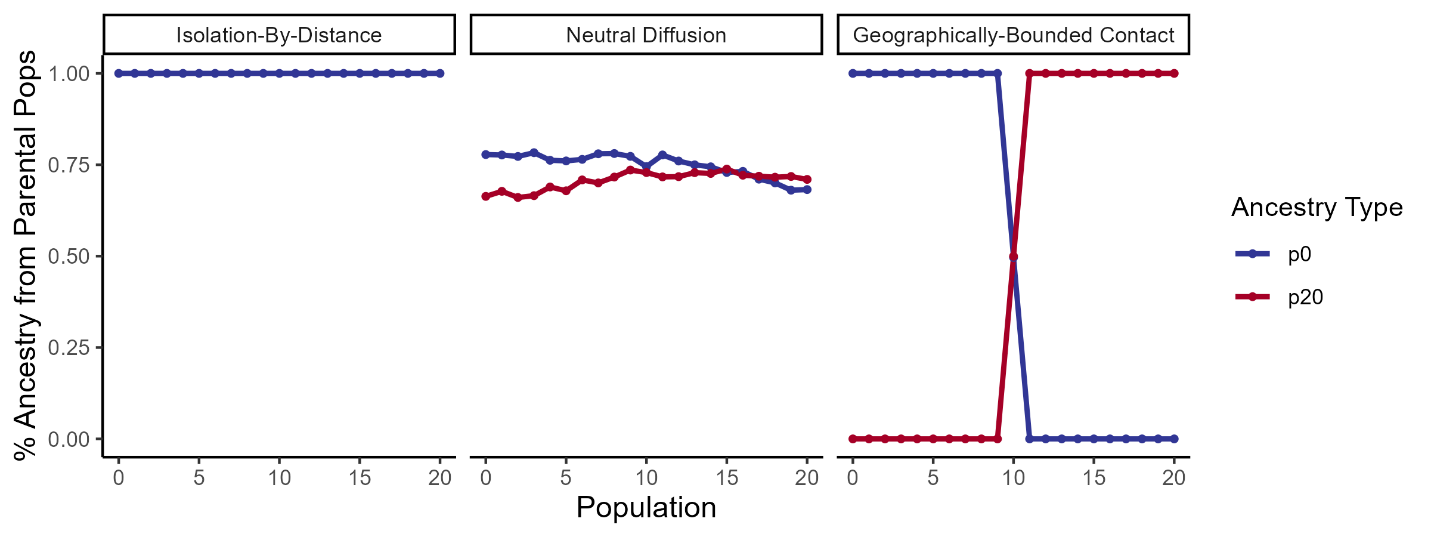

**Fig S4.** True ancestry proportions in each population of the full simulations at the end of Phase III (i.e. the end of the simulation). Calculated for each population based on the percentage of alleles which were fixed in p0 and p20 at the end of Phase II (i.e. at the end of the allopatric divergence phase) that are present in that population at the end of Phase III. Blue indicates alleles that are informative of p0 ancestry and red indicates alleles that are informative of p20 ancestry. Red not shown for IBD plot because p20 did not exist at the end of Phase II.

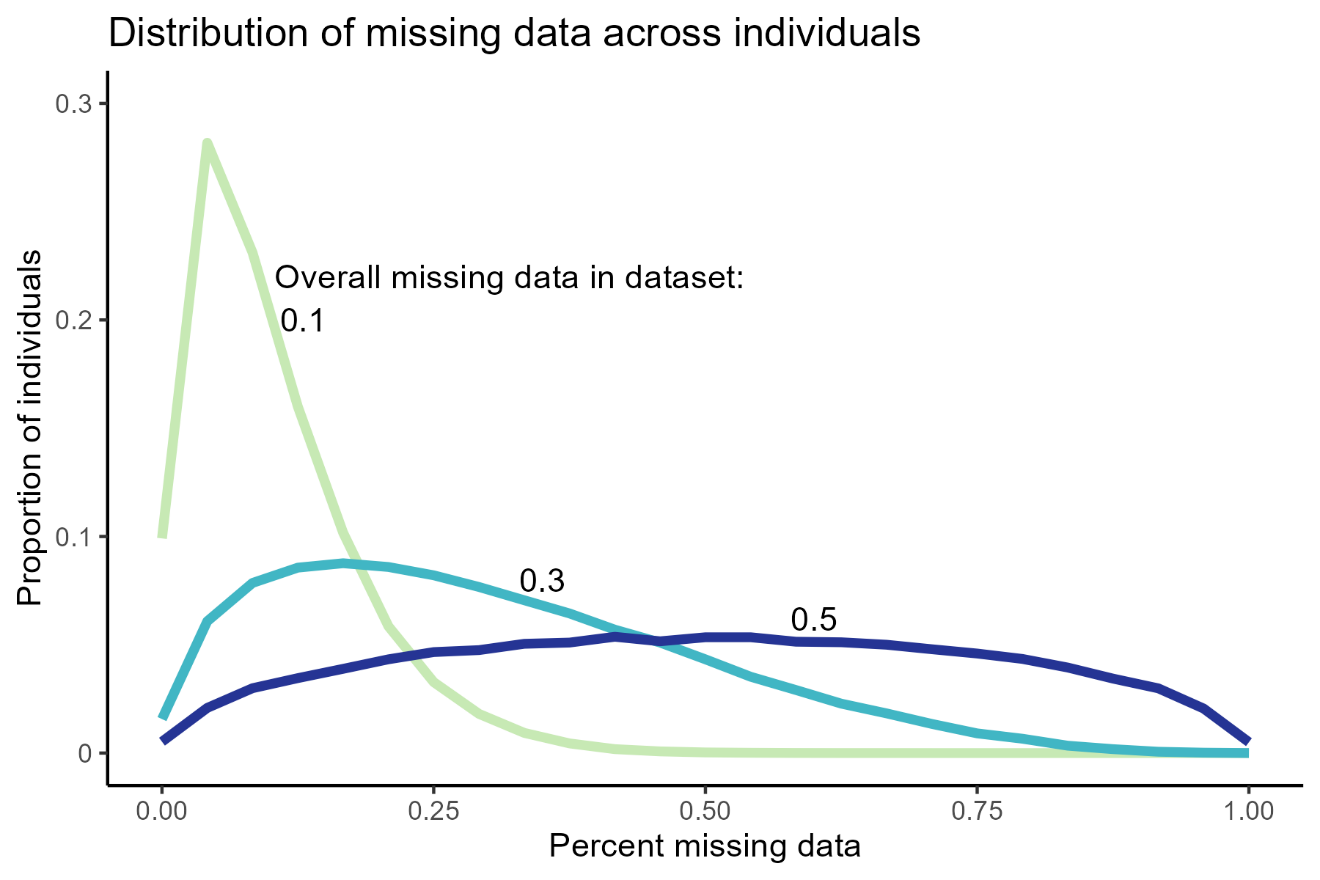

**Fig S5.** Distributions of amount of missing data per sample when overall missing data is 0.1, 0.3, and 0.5. The amount of missing data per sample follows a Beta distribution, such that most individuals have low missing data, as is often the case for empirical datasets.

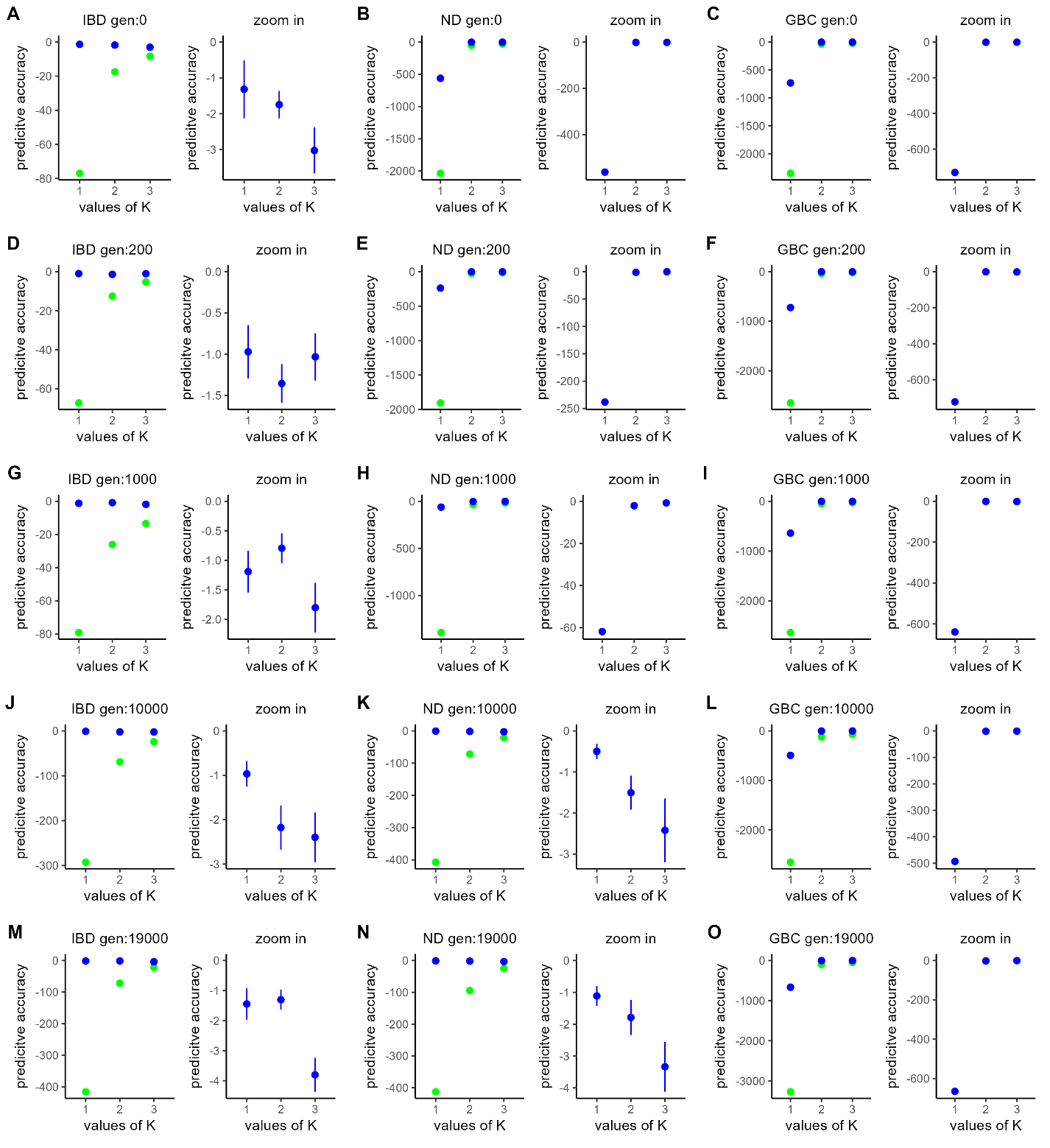

**Fig S6.** Cross validation of spatial (**green**) and nonspatial (**blue**) *conStruct* models for K=1-3 for each simulation at generations **(A-C)** 0, **(D-F)** 200, **(G-I)** 1000, **(J-L)** 10,000, and **(M-O)** 19,000. Simulations abbreviated as IBD (Isolation-By-Distance), ND (Neutral Diffusion), and GBC (Geographically-Bounded Contact). Each panel consists of the average predictive accuracy of each model based on eight replicates, with the best model standardized to 0 and lower scores indicating worse predictive accuracy. The left plot of each panel shows the spatial and non-spatial models, while the right plot zooms in on the spatial model.

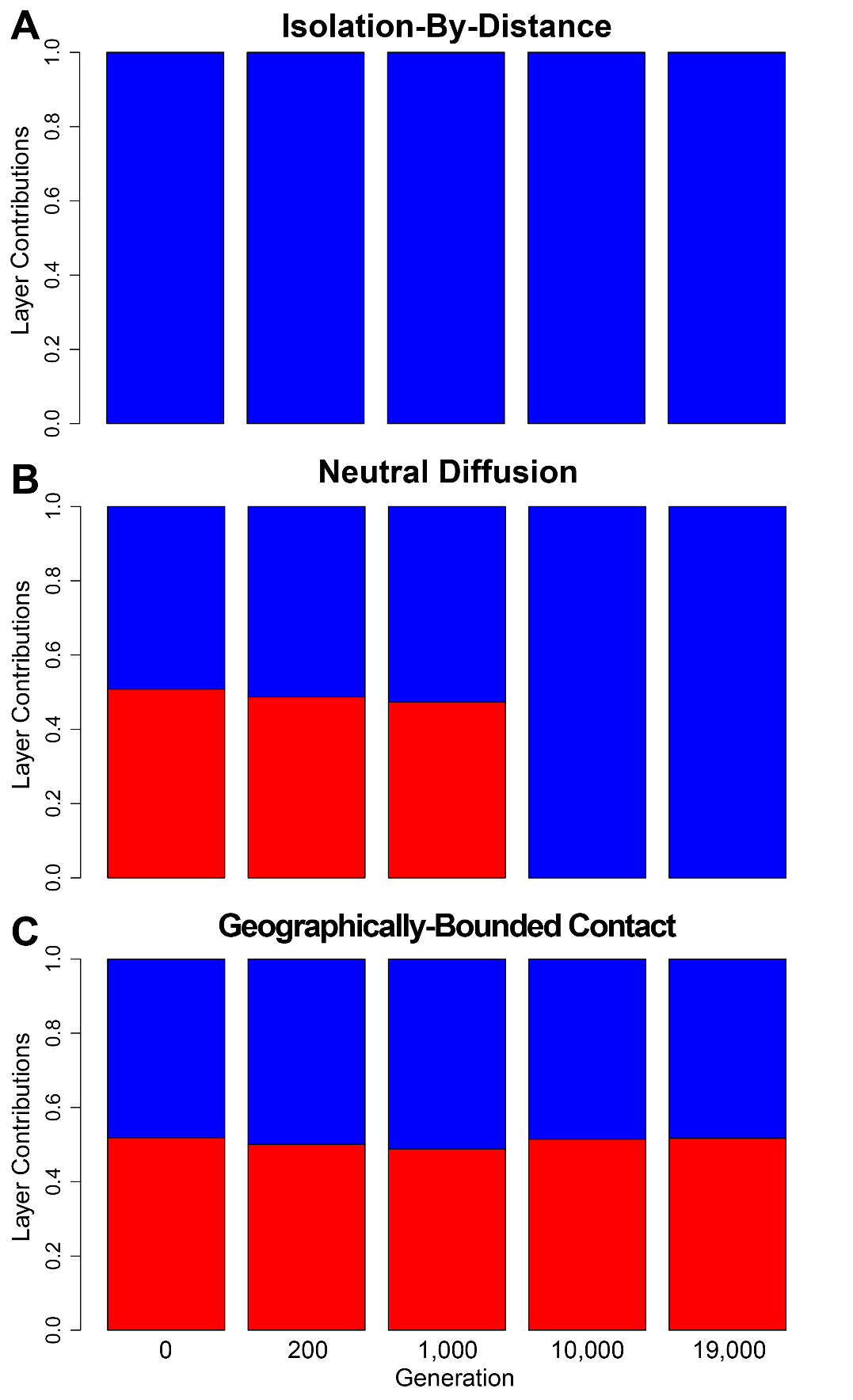

**Fig S7.** Covariance contributed by each layer (i.e. cluster) of the spatial *conStruct* model for K=2 for each simulation at generations 0, 200, 1000, 10000, and 19000. Each bar represents the total covariance across all populations at that generation. Blue indicates the proportion of covariance contributed by the first layer (i.e. the cluster that corresponds to p0 ancestry) and red indicates the proportion of covariance contributed by the second layer (i.e. the cluster that corresponds to p20 ancestry).

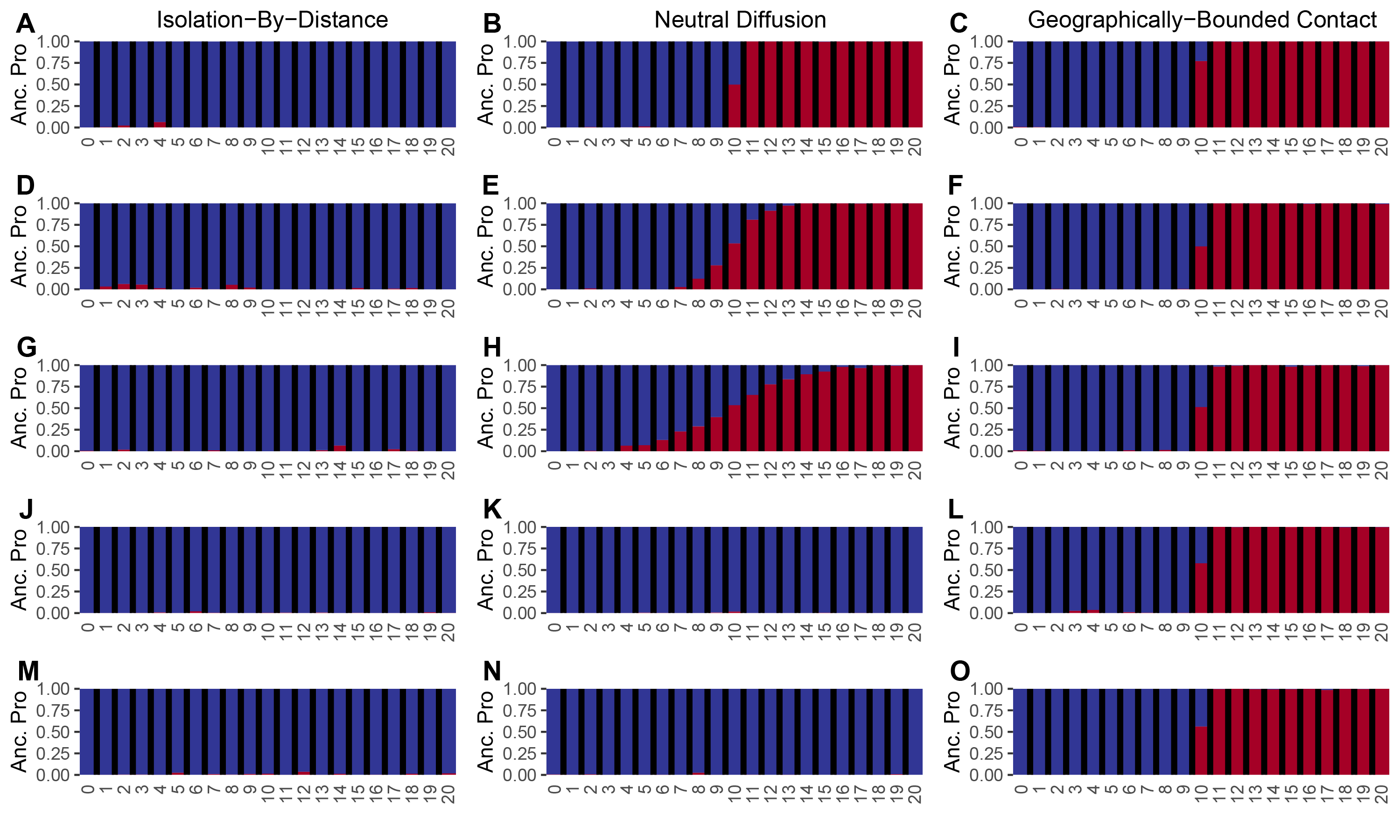

**Fig S8.** *conStruct* plots for each simulation at generations **(A-C)** 0, **(D-F)** 200, **(G-I)** 1000, **(J-L)** 10,000, and **(M-O)** 19,000 using the spatial model and K=2. Allele frequencies were pooled by population for clustering and bar plots show inferred ancestry proportions for each population.

**Table S1.** Linear models fit to the relationship between hybrid index and interclass heterozygosity. Significant linear relationships between hybrid index and interclass heterozygosity are bolded in the slope column (*p<0.05).

| Simulation | Generation | Slope | SE | t | p value |
| --- | --- | --- | --- | --- | --- |
| IBD | 0 | **-0.128** | 0.024 | -5.328 | 1.63E-07* |
|  | 200 | **-0.342** | 0.025 | -13.567 | 5.30E-35* |
|  | 1000 | **-0.408** | 0.020 | -20.030 | 4.72E-63* |
|  | 10000 | **-0.086** | 0.008 | -10.662 | 1.22E-23* |
|  | 19000 | 0.017 | 0.009 | 1.878 | 0.0611 |
| ND | 0 | 0.00 | 0.012 | -0.025 | 0.980 |
|  | 200 | -0.004 | 0.014 | -0.251 | 0.802 |
|  | 1000 | -0.016 | 0.017 | -0.970 | 0.332 |
|  | 10000 | **0.050** | 0.011 | 4.668 | 4.11E-06* |
|  | 19000 | **-0.065** | 0.009 | -7.573 | 2.37E-13* |
| GBC | 0 | 0.003 | 0.008 | 0.338 | 0.735 |
|  | 200 | -0.005 | 0.011 | -0.445 | 0.657 |
|  | 1000 | 0.005 | 0.009 | 0.565 | 0.573 |
|  | 10000 | 0.007 | 0.009 | 0.742 | 0.459 |
|  | 19000 | 0.002 | 0.009 | 0.251 | 0.802 |
