## Supplementary figures and images for "That’s not a Hybrid: How to Distinguish Patterns of Admixture and Isolation-by-Distance"

### IBD.200.str.png

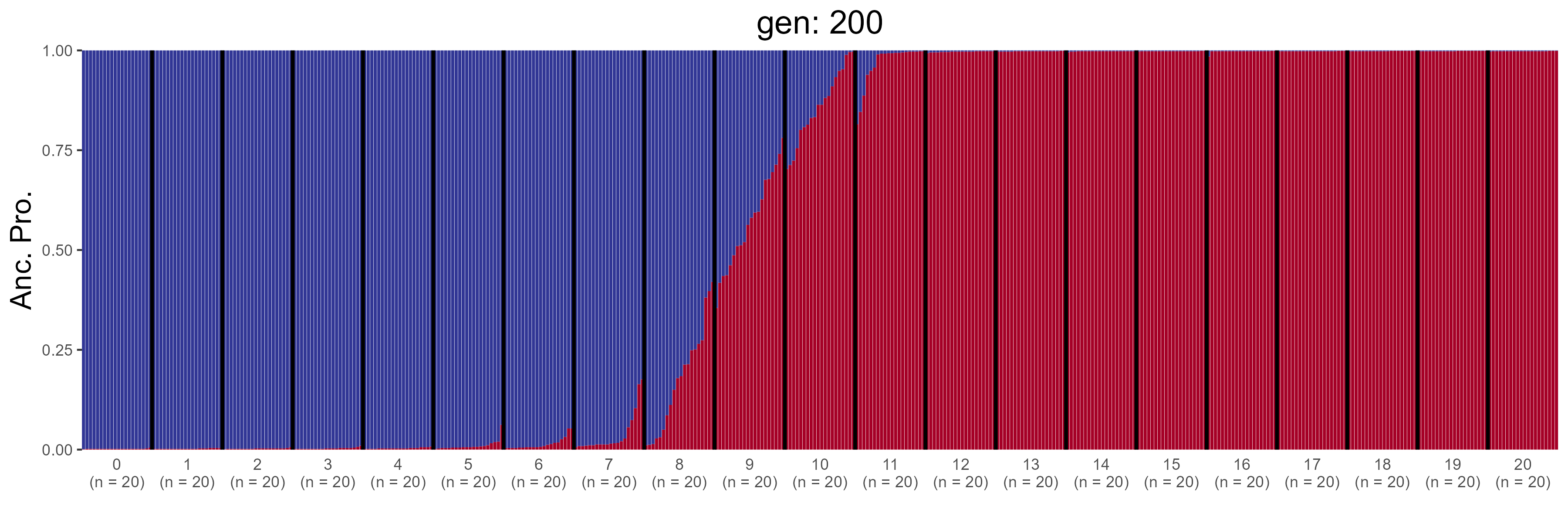

### IBD.400.str.png

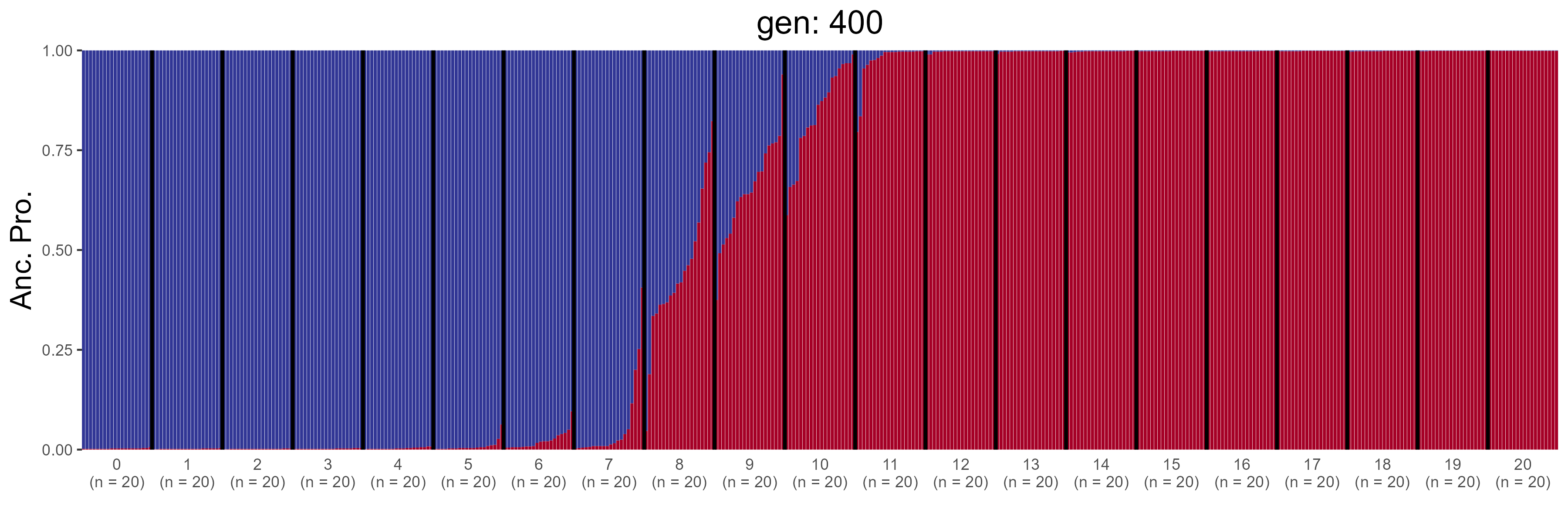

### IBD.600.str.png

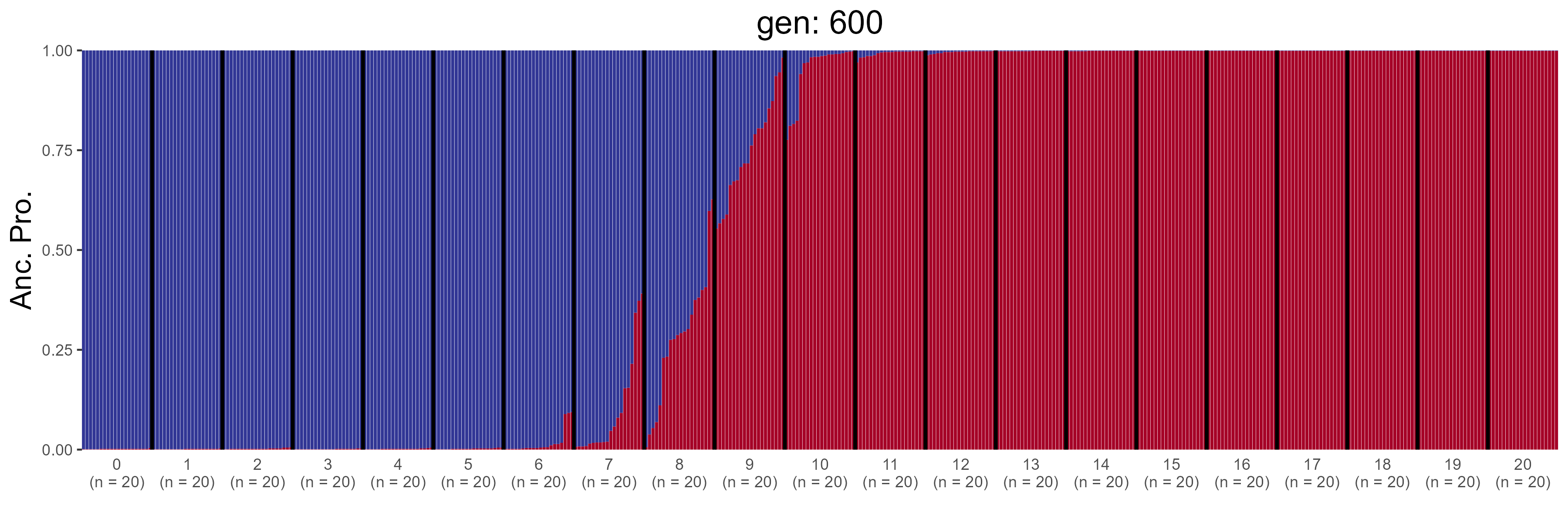

### IBD.800.str.png

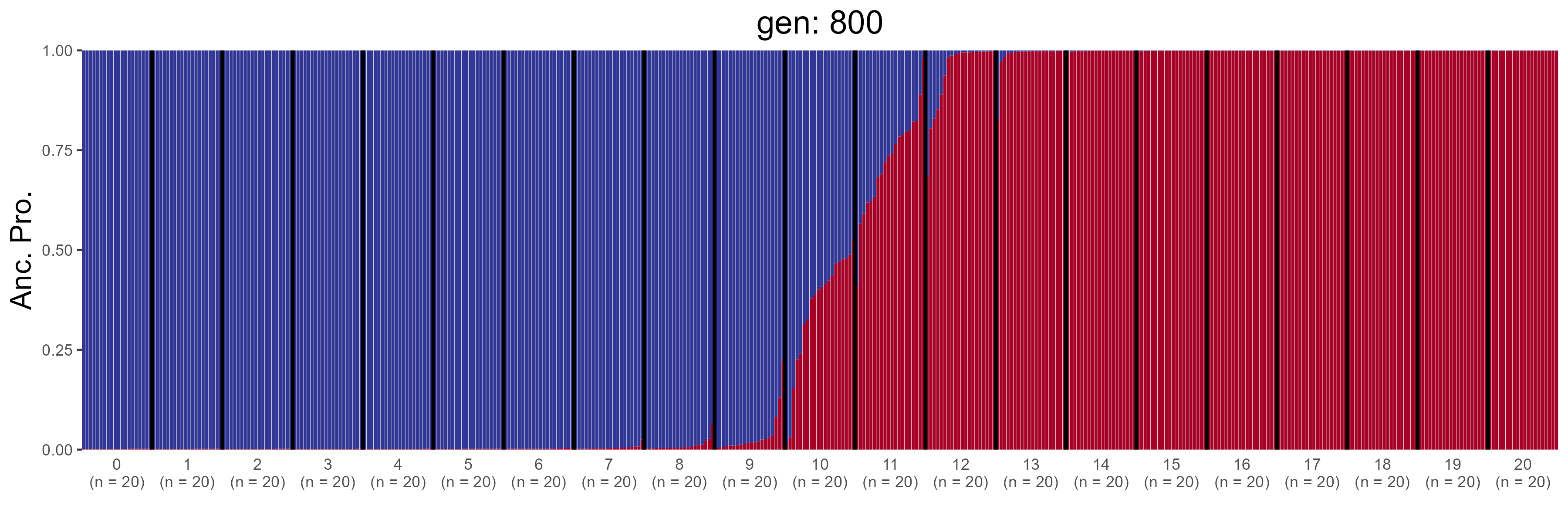

### IBD.1200.str.png

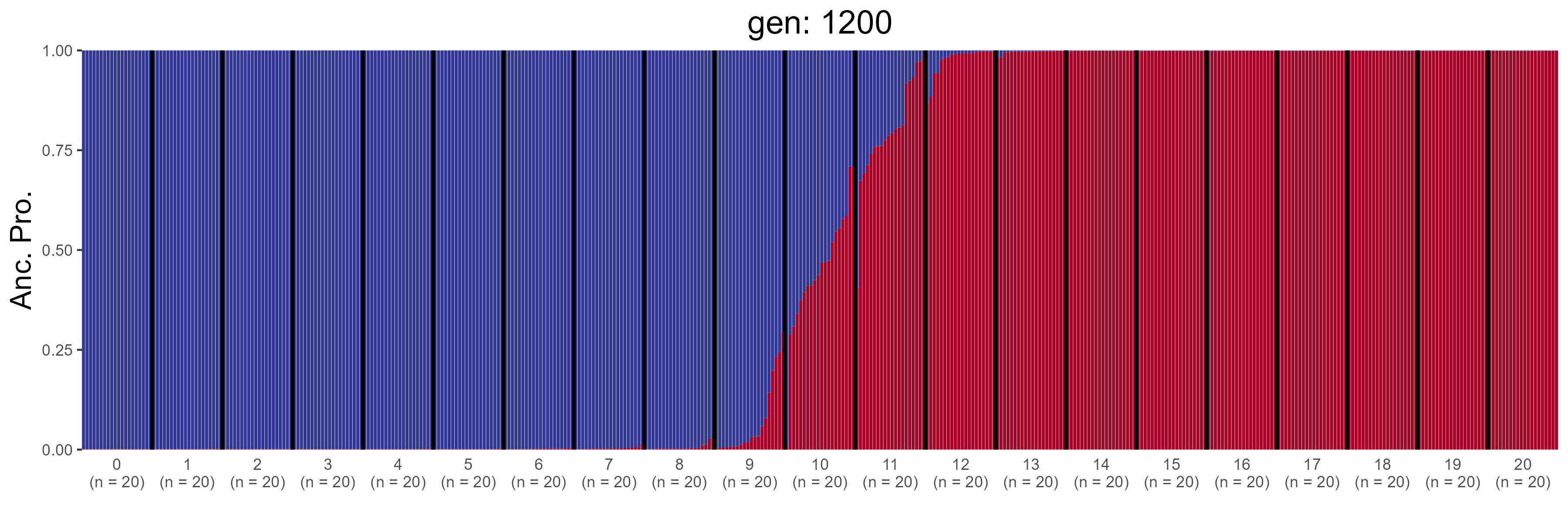

### IBD.1400.str.png

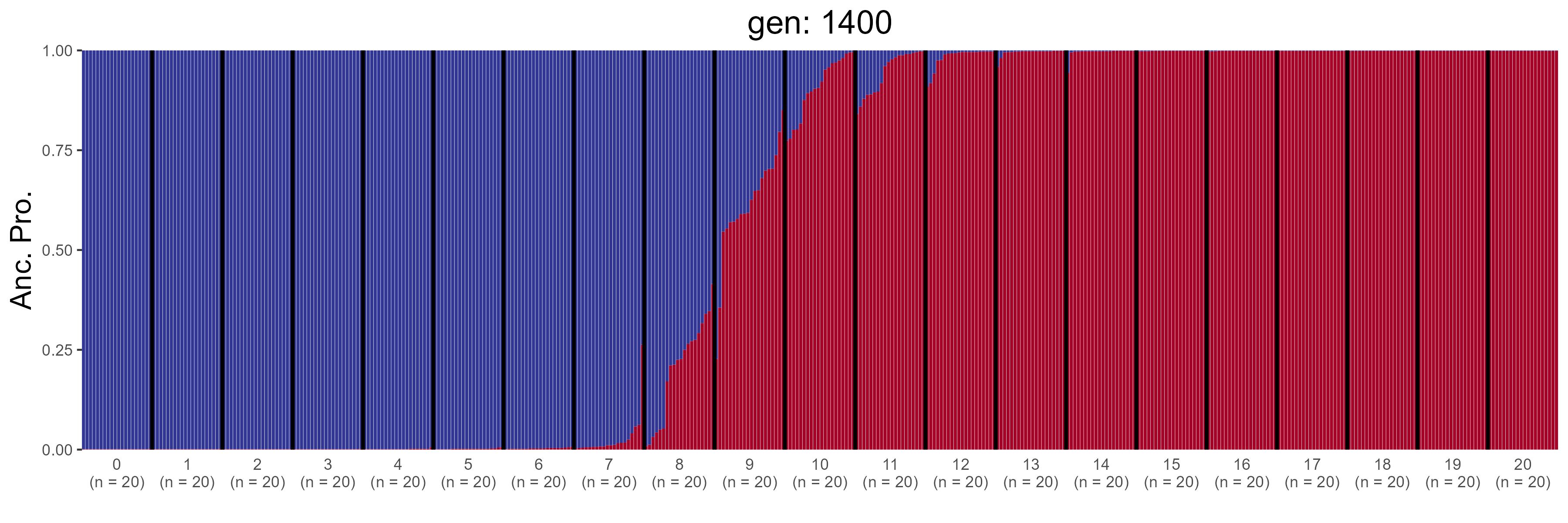

### IBD.1600.str.png

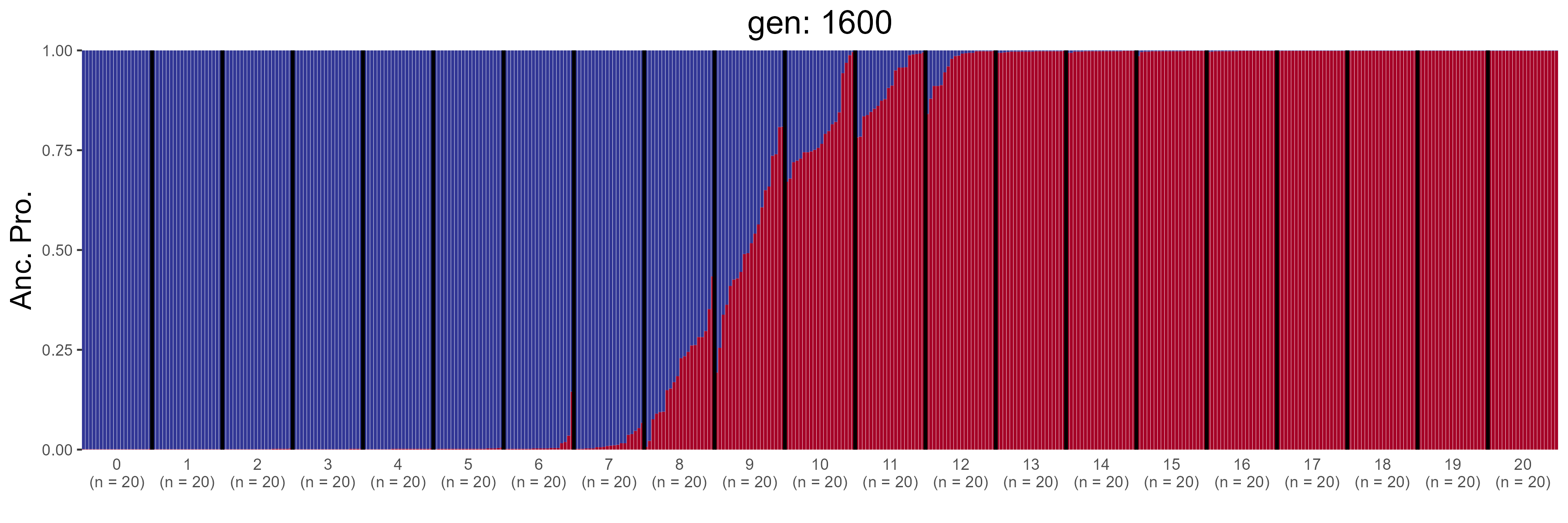

### IBD.1800.str.png

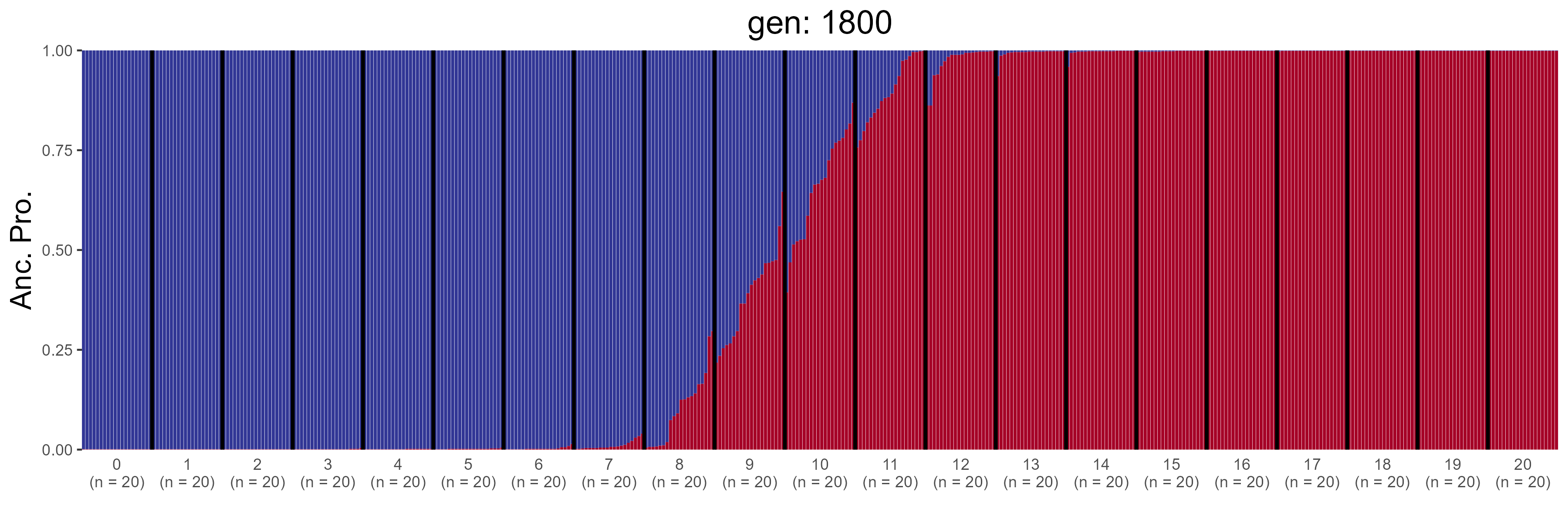

### IBD.2000.str.png

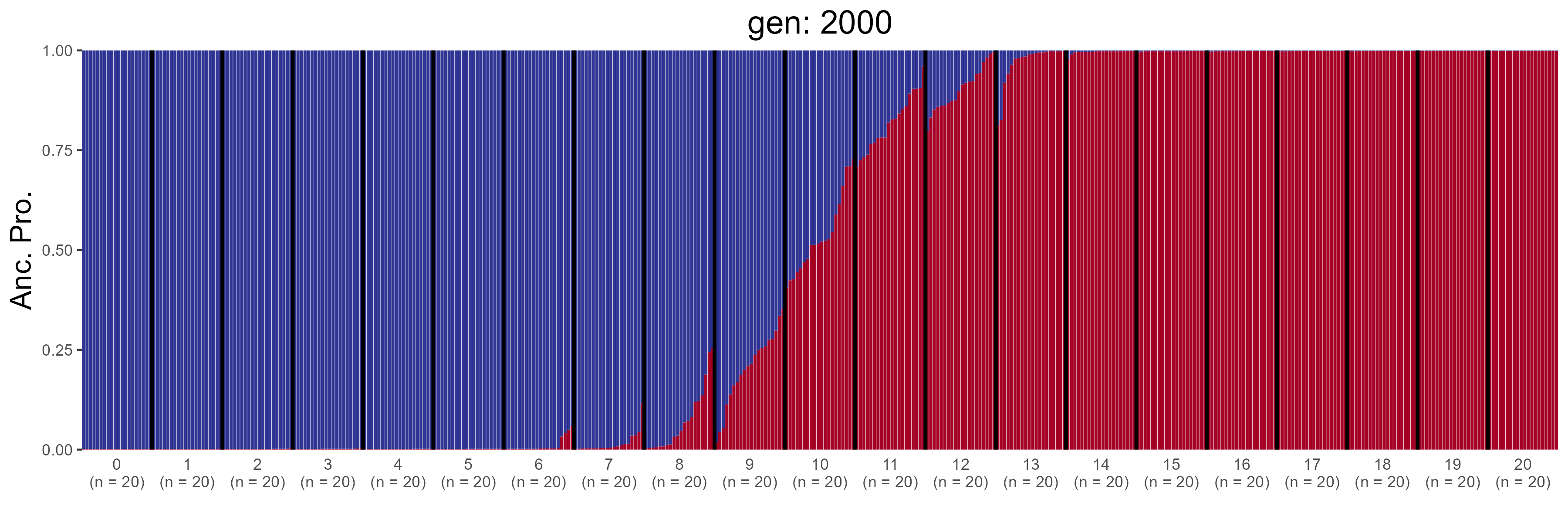

### IBD.2200.str.png

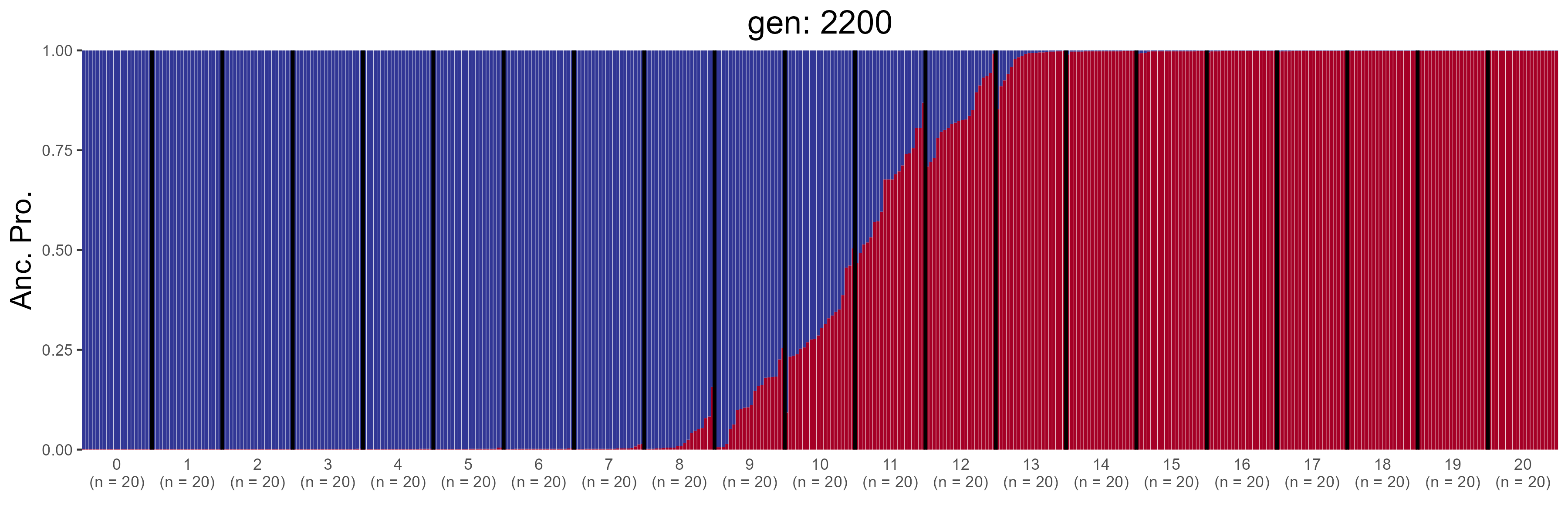

### IBD.2400.str.png

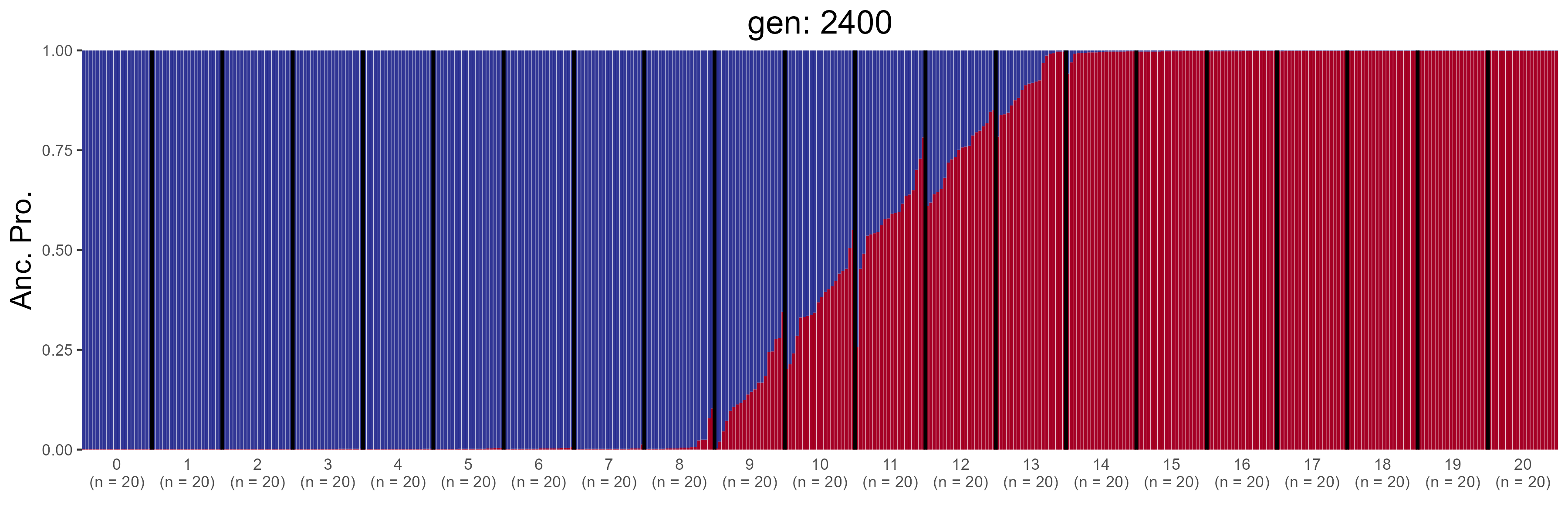

### IBD.2600.str.png

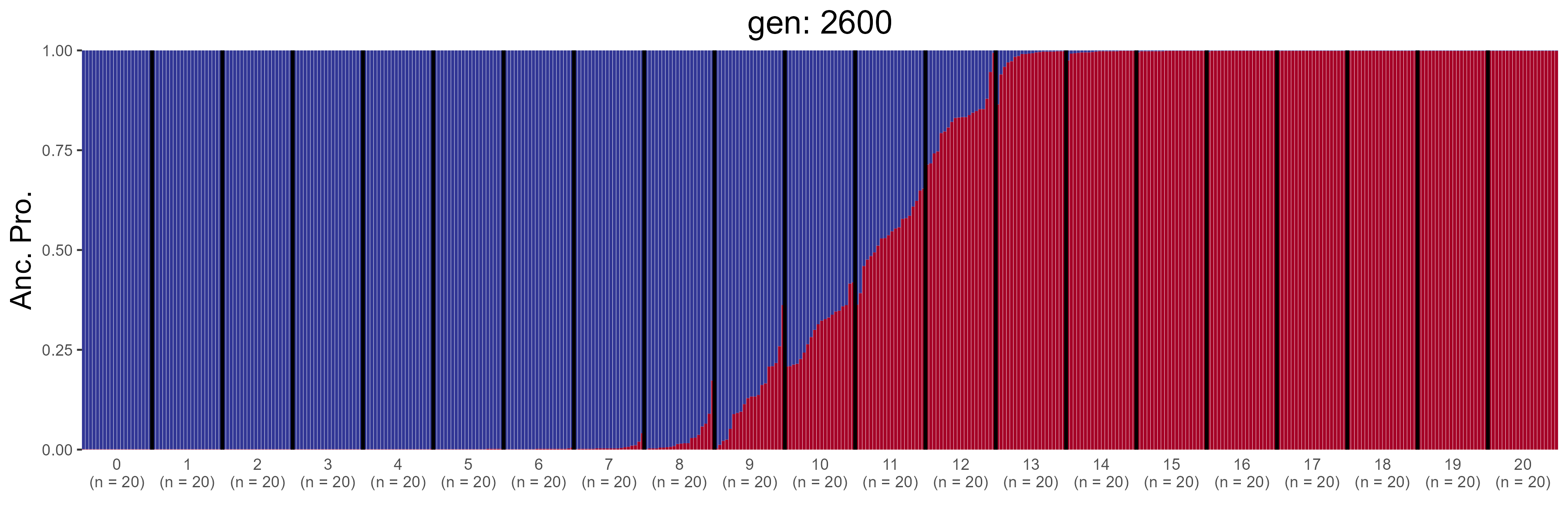

### IBD.2800.str.png

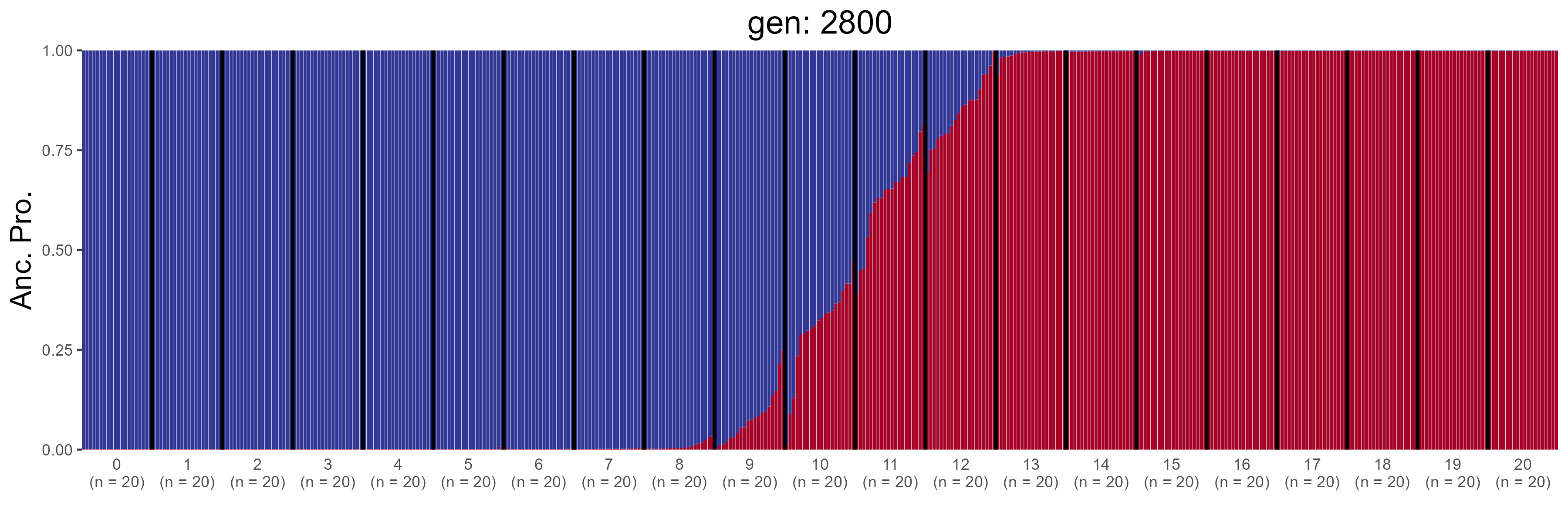

### IBD.3000.str.png

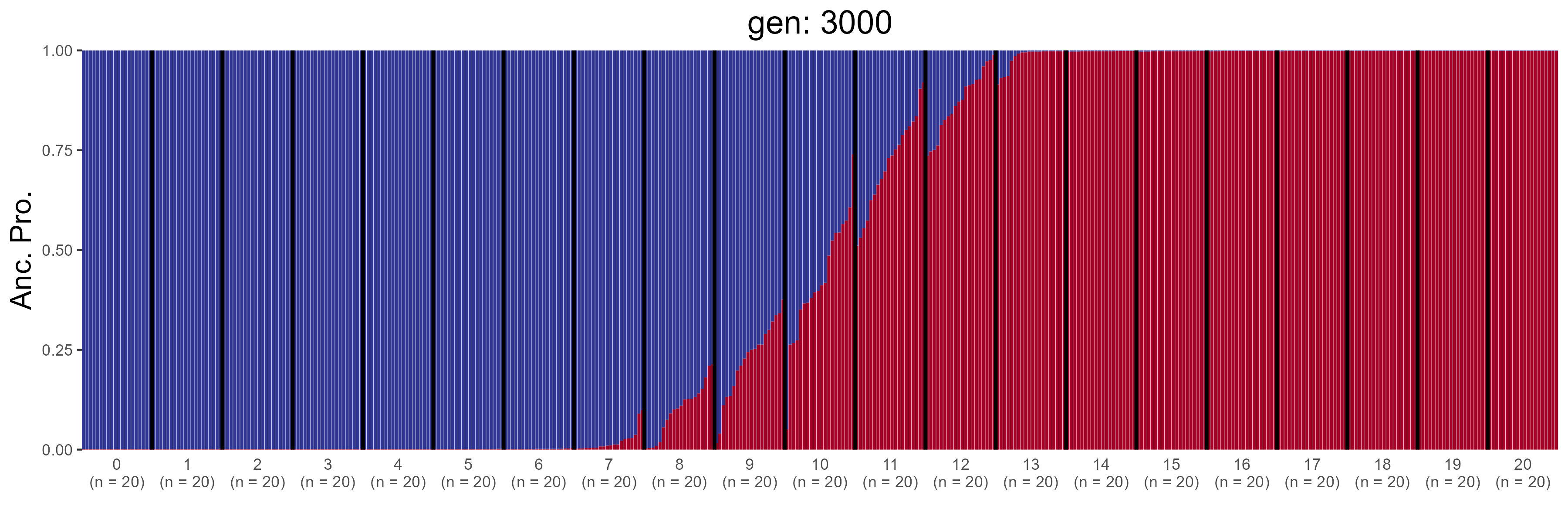

### IBD.3200.str.png

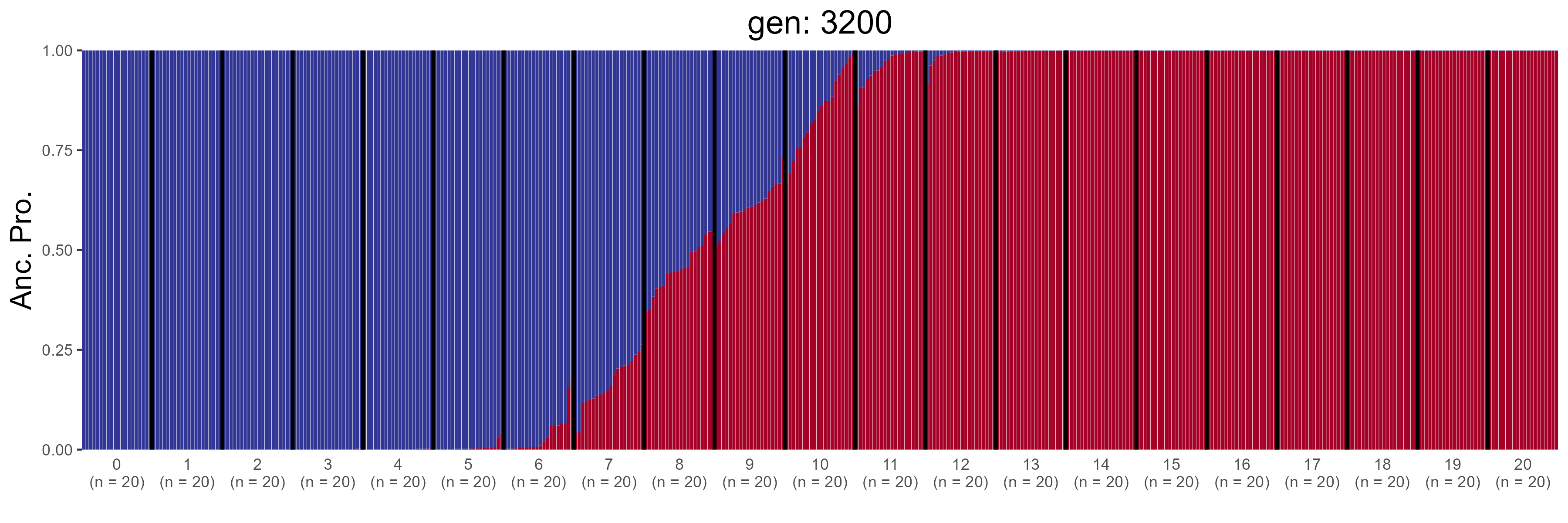

### IBD.3400.str.png

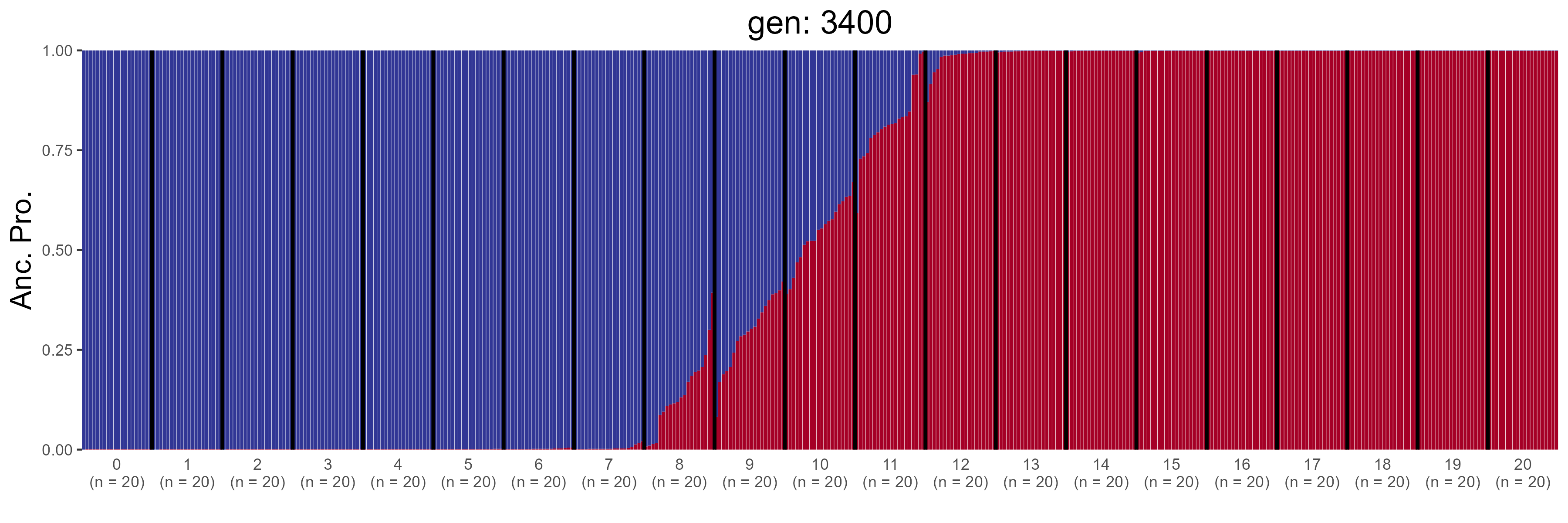

### IBD.3600.str.png

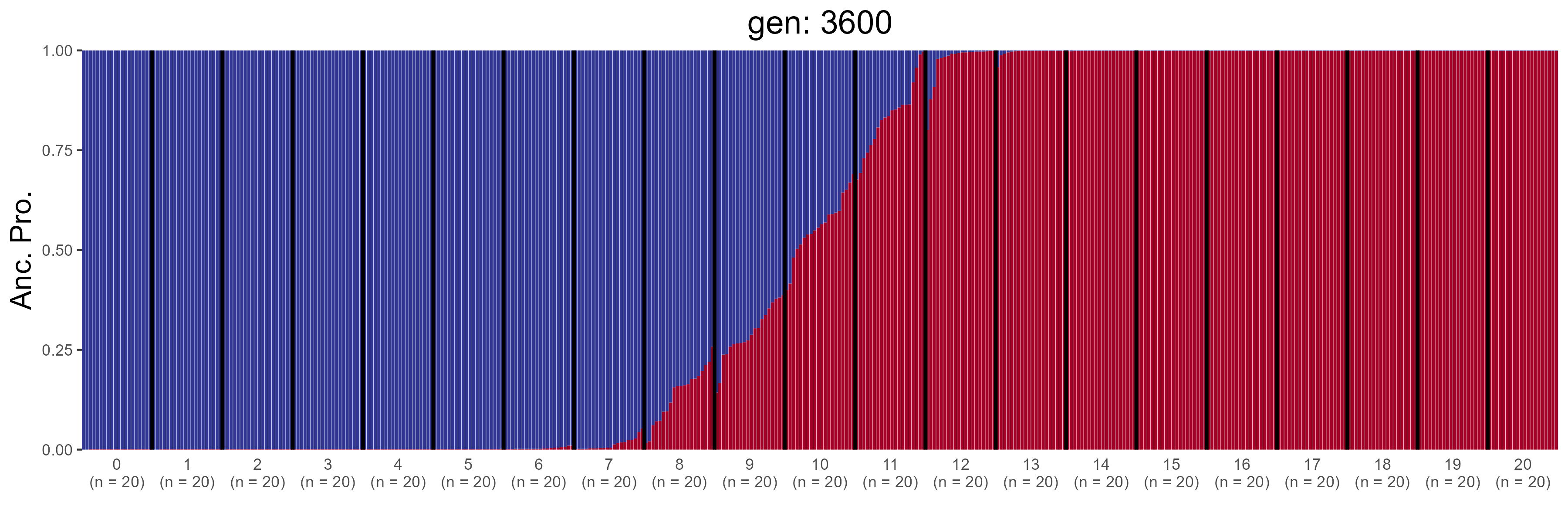

### IBD.3800.str.png

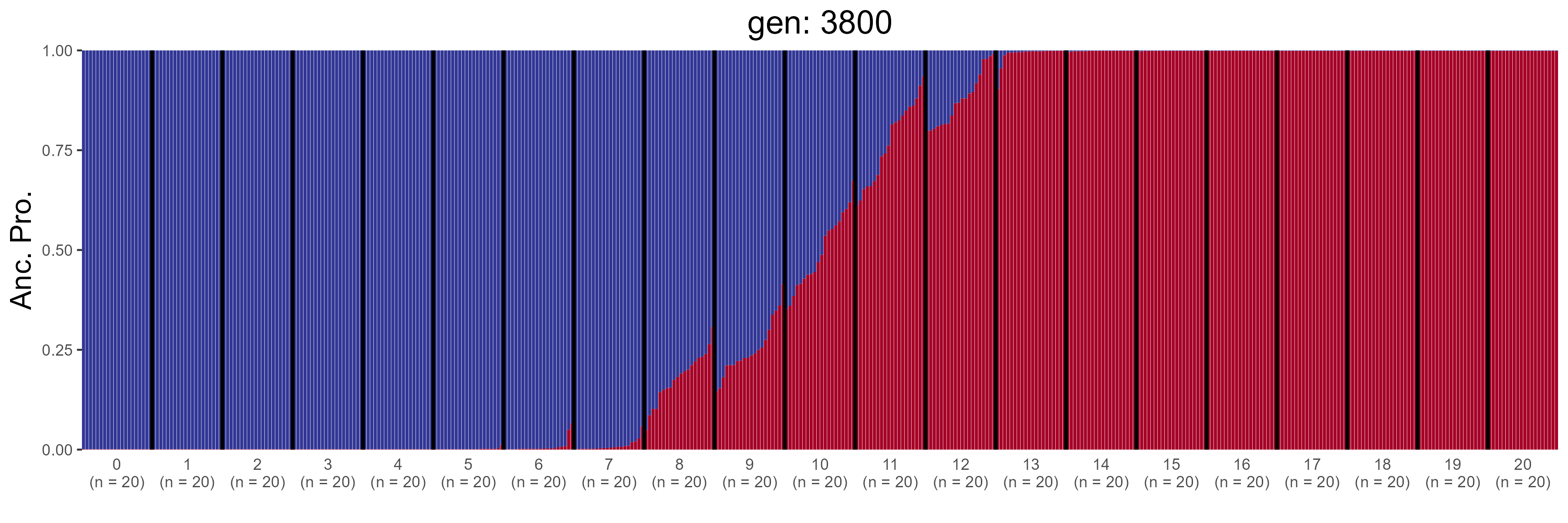

### IBD.4000.str.png

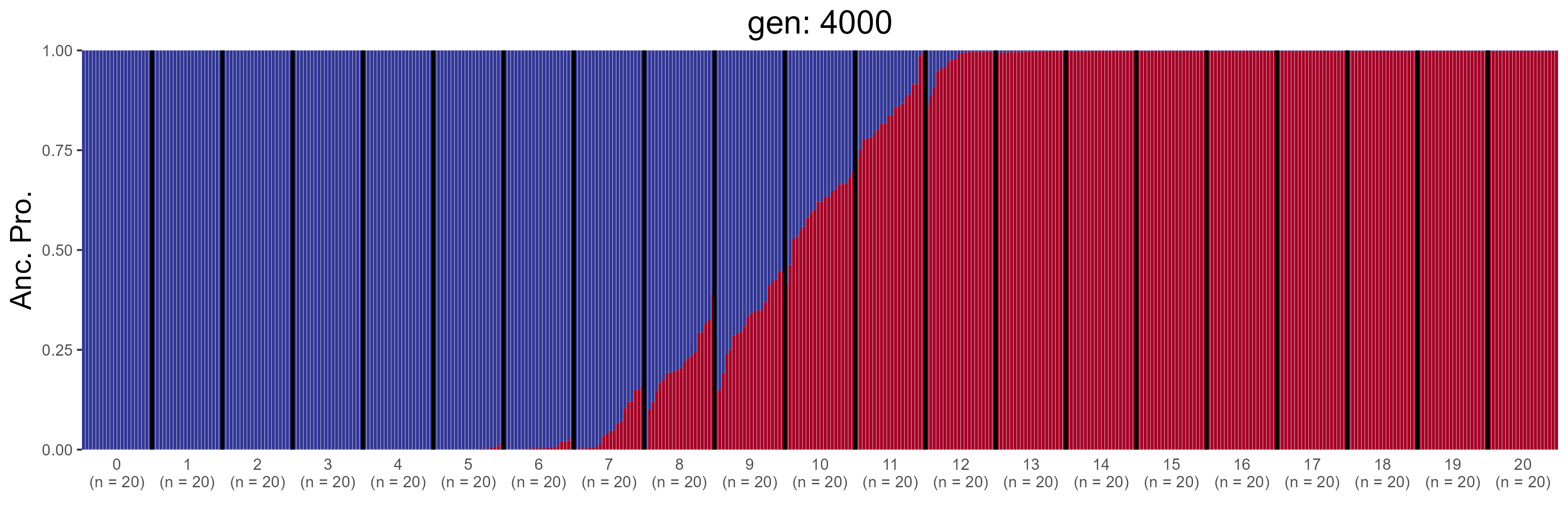

### IBD.4200.str.png

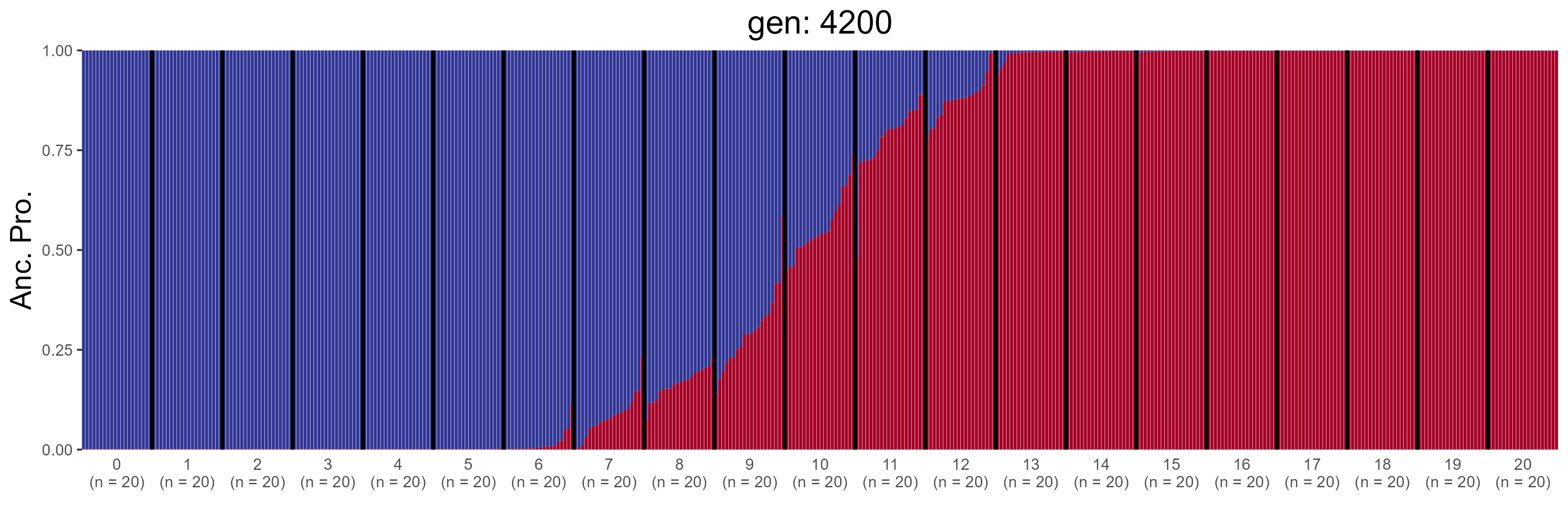

### IBD.4400.str.png

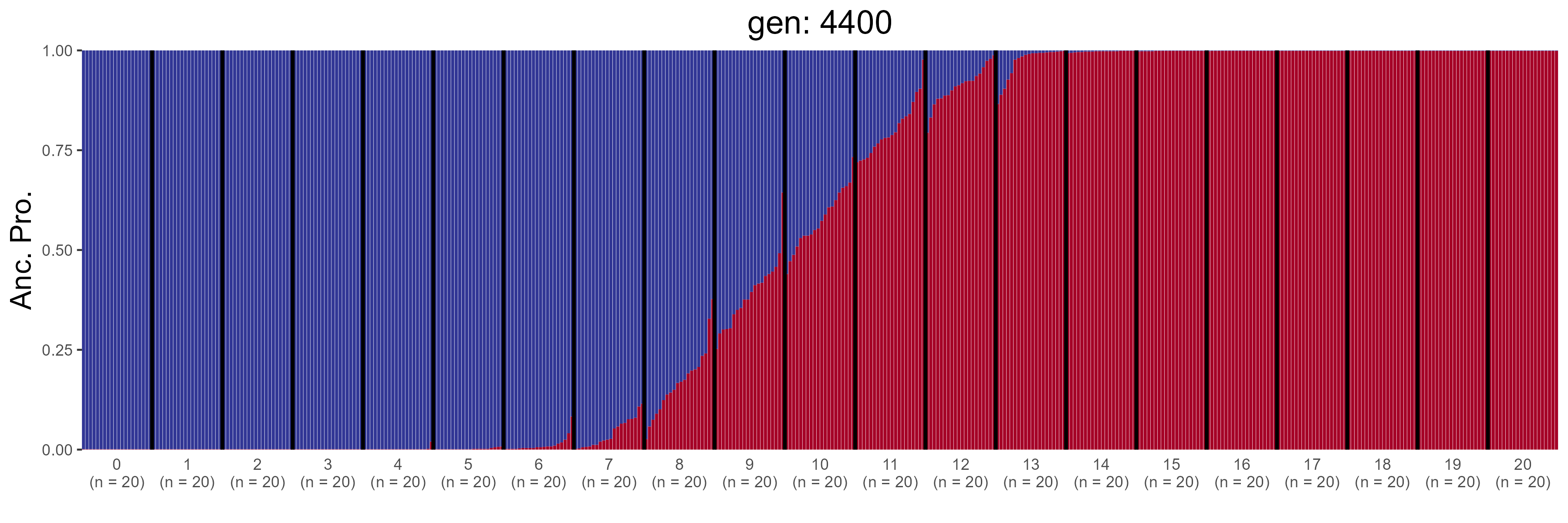

### IBD.4600.str.png

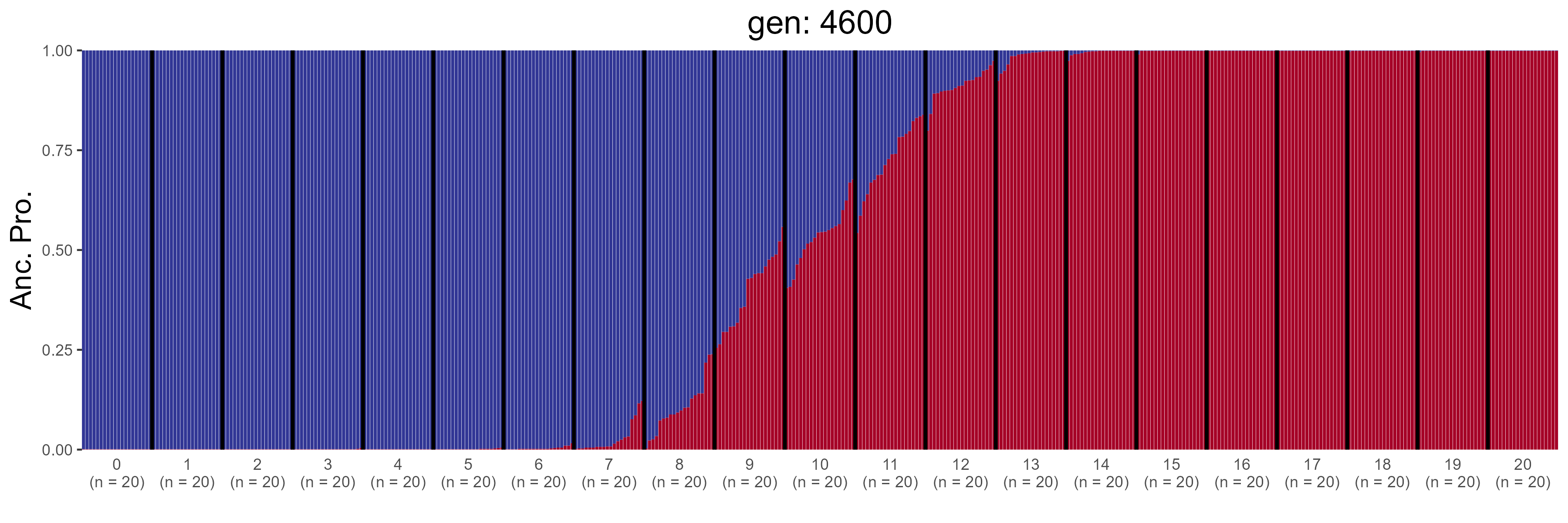

### IBD.4800.str.png

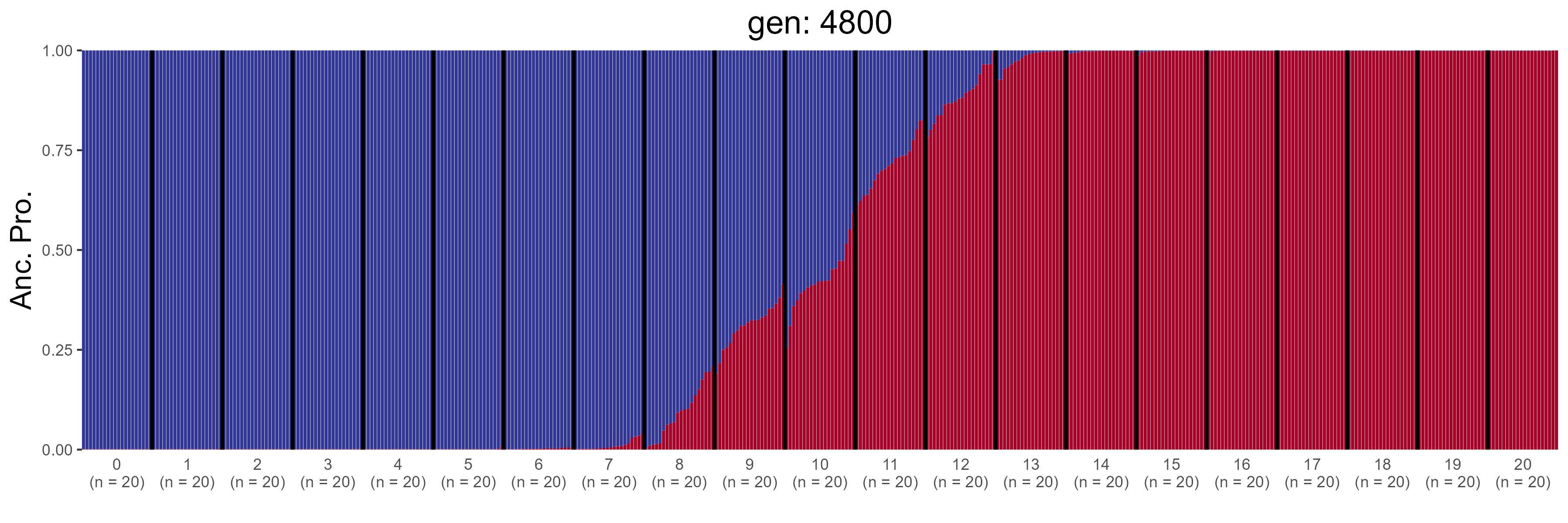

### IBD.5000.str.png

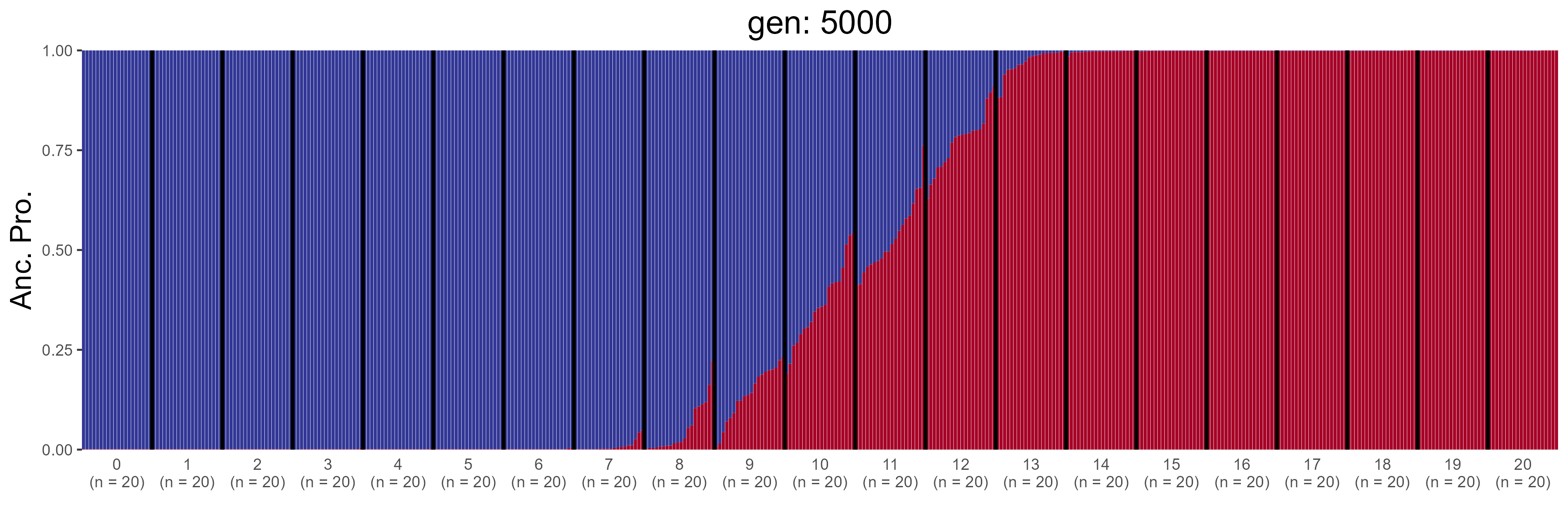

### IBD.5200.str.png

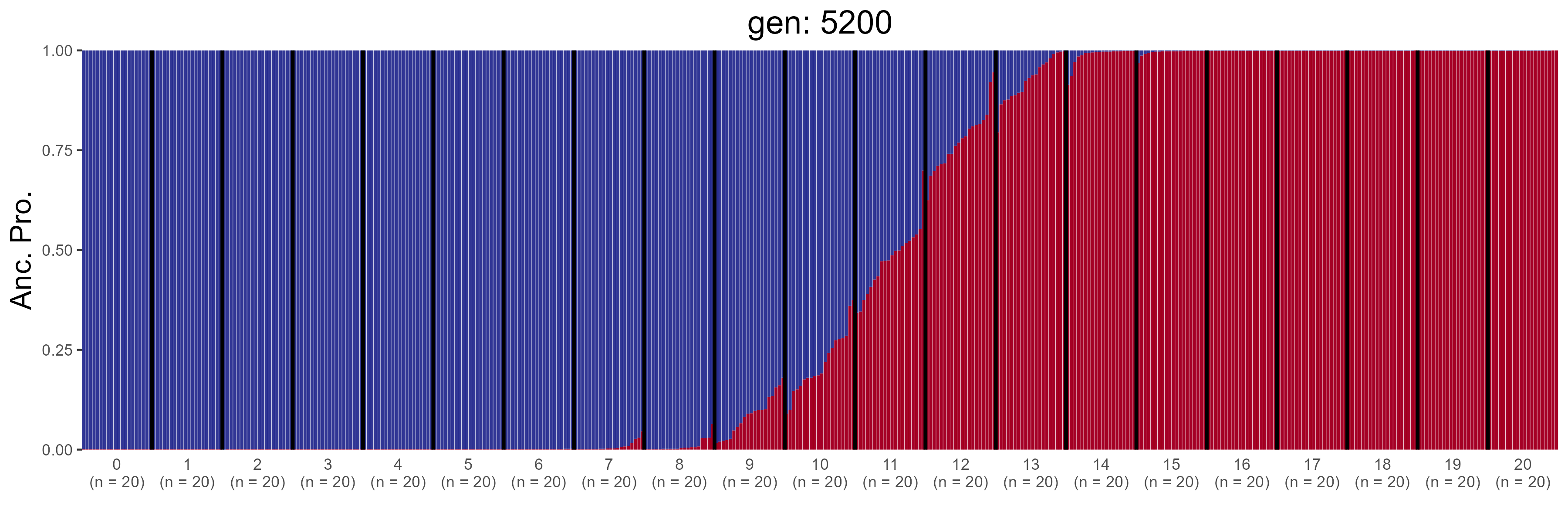

### IBD.5400.str.png

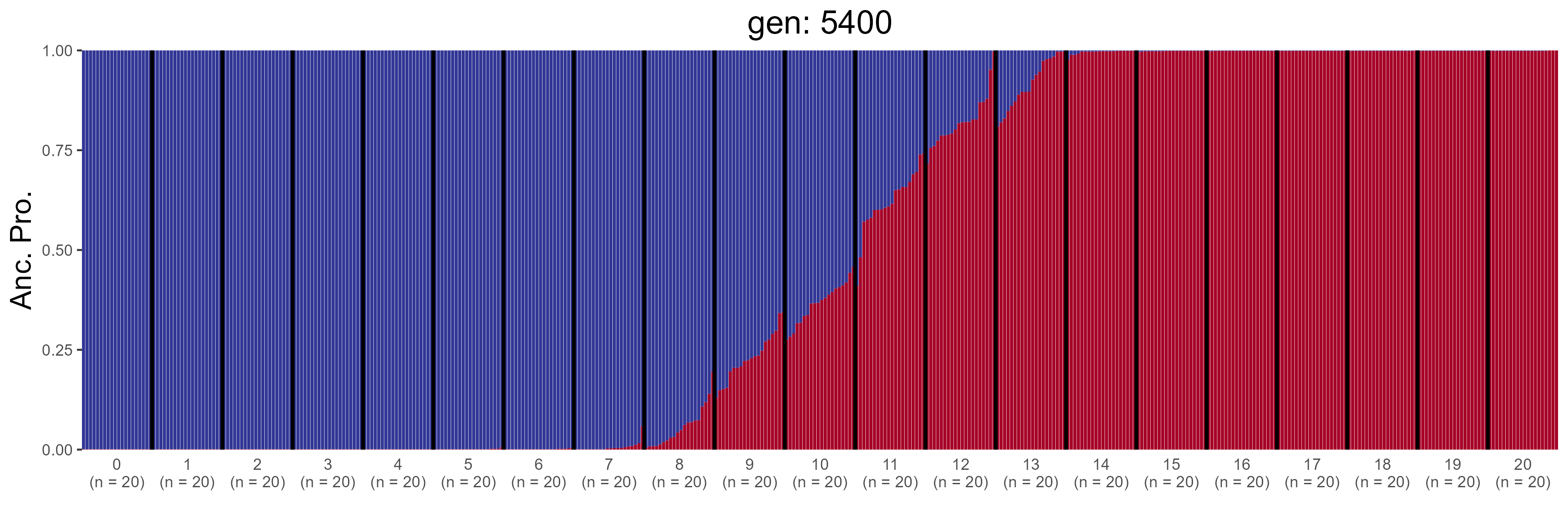

### IBD.5600.str.png

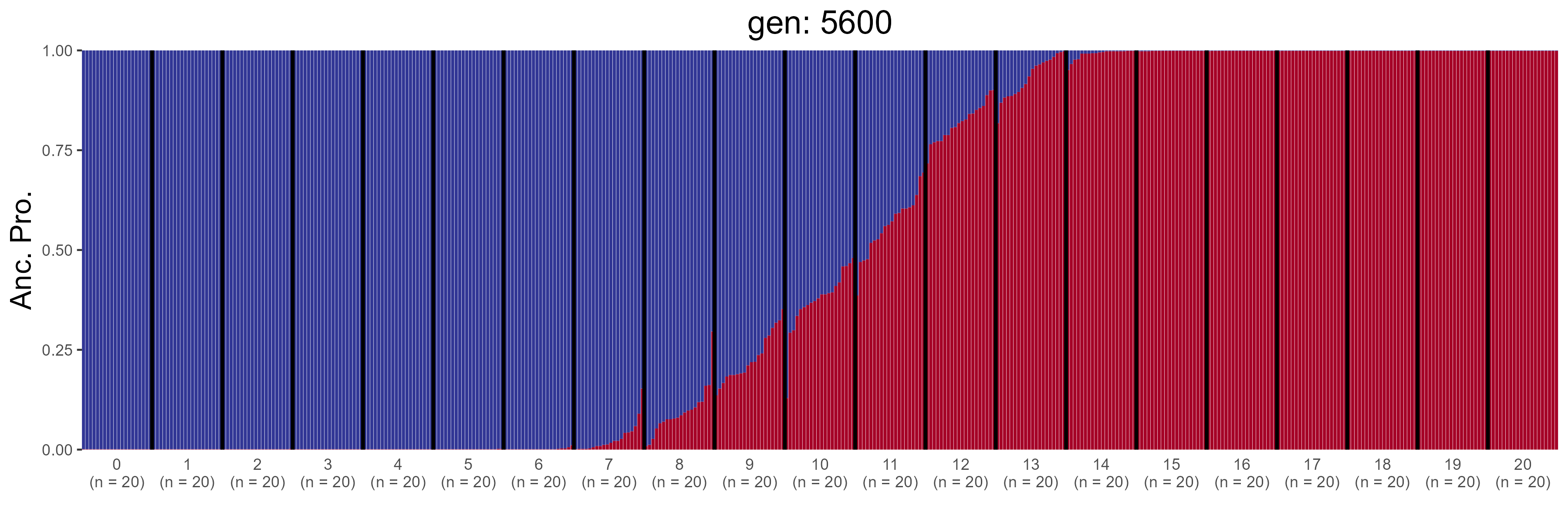

### IBD.5800.str.png

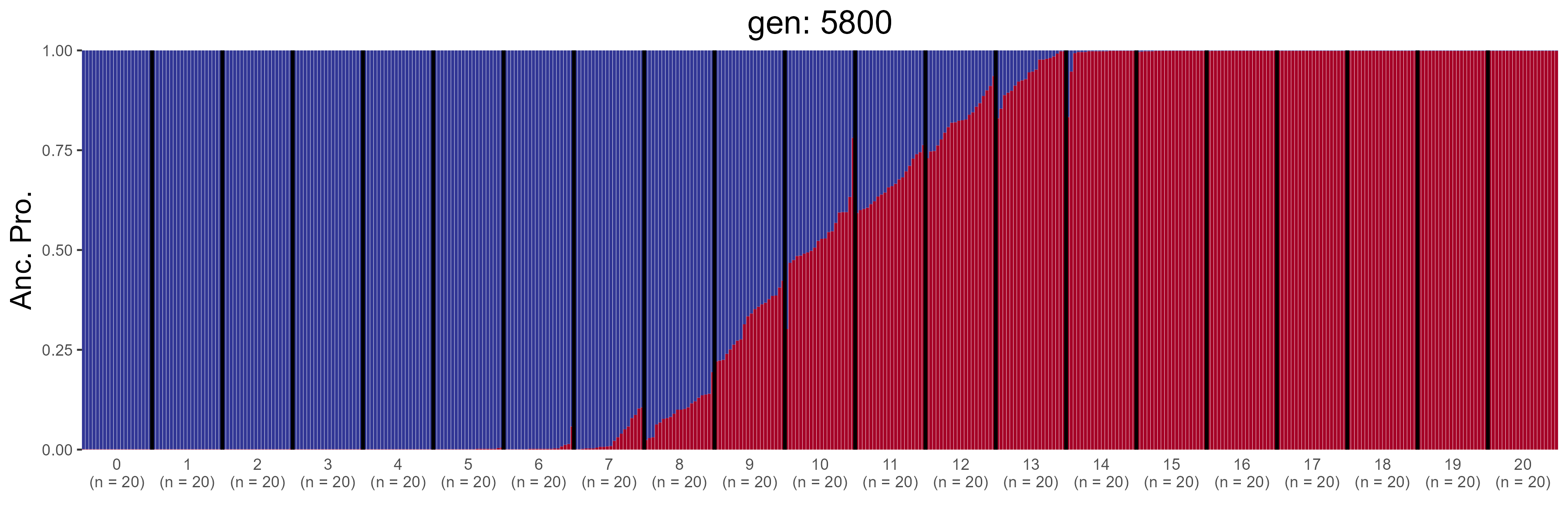

### IBD.6000.str.png

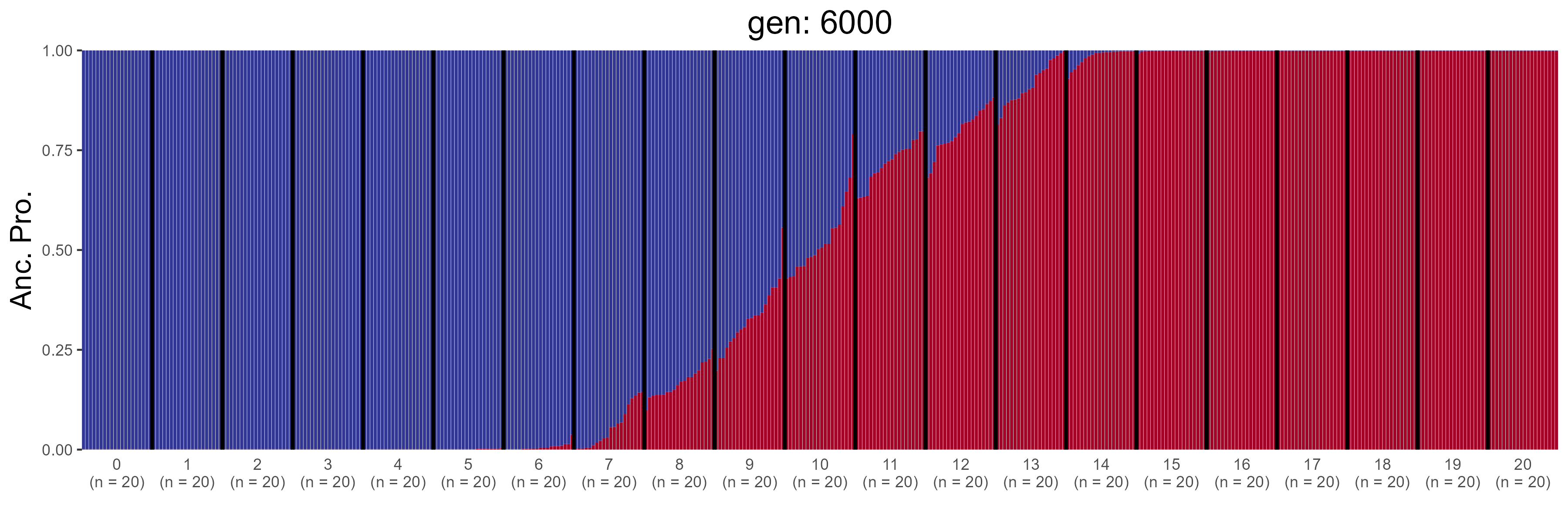

### IBD.6200.str.png

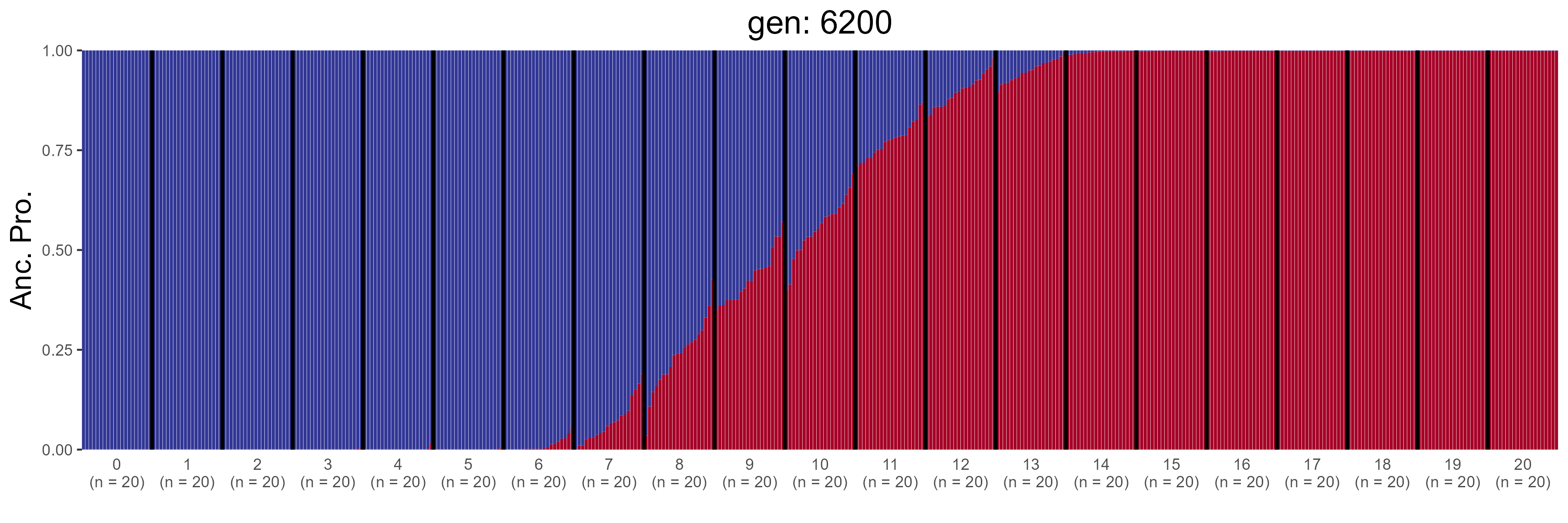

### IBD.6400.str.png

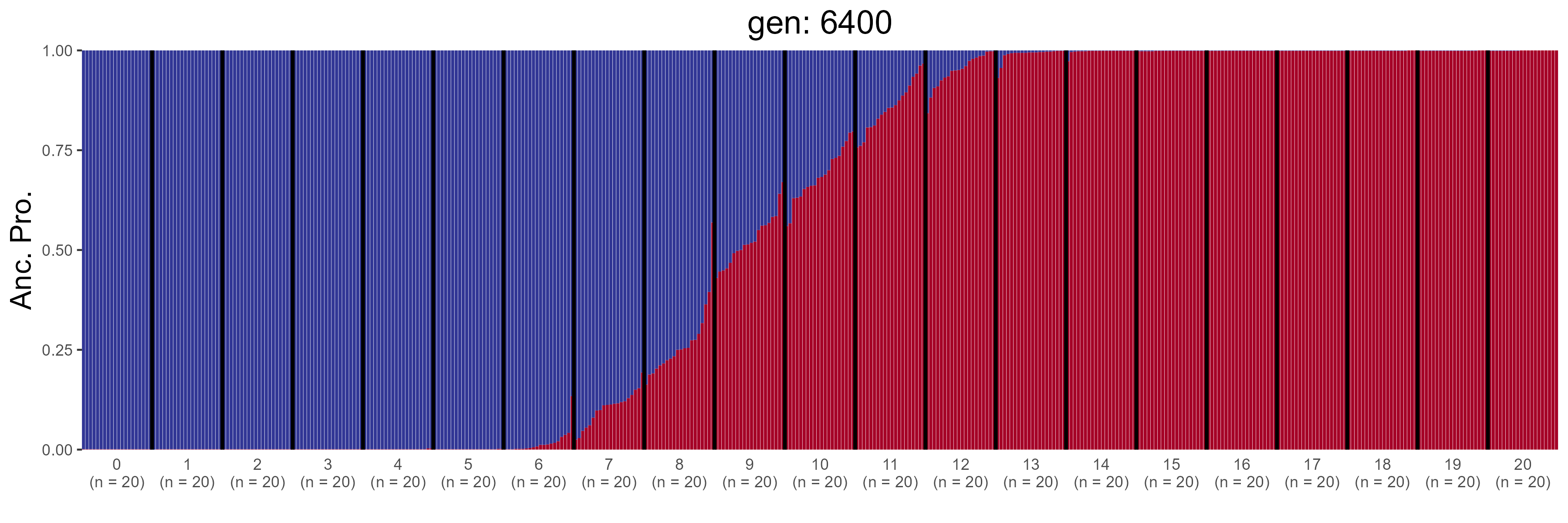

### IBD.6600.str.png

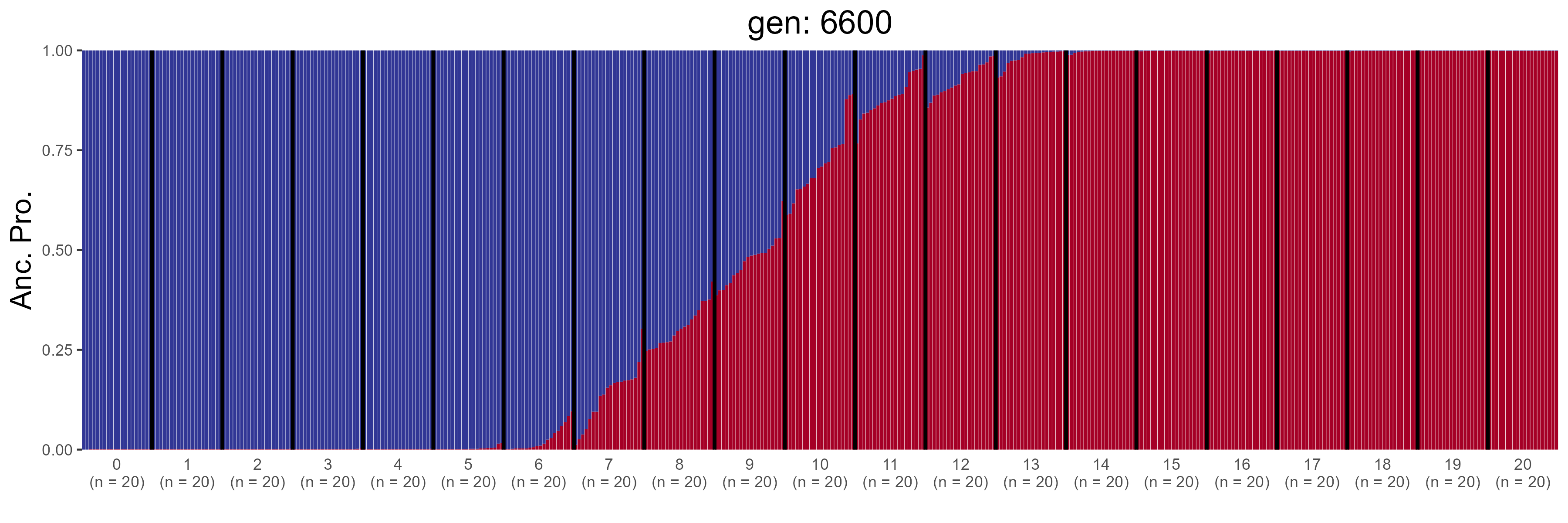

### IBD.6800.str.png

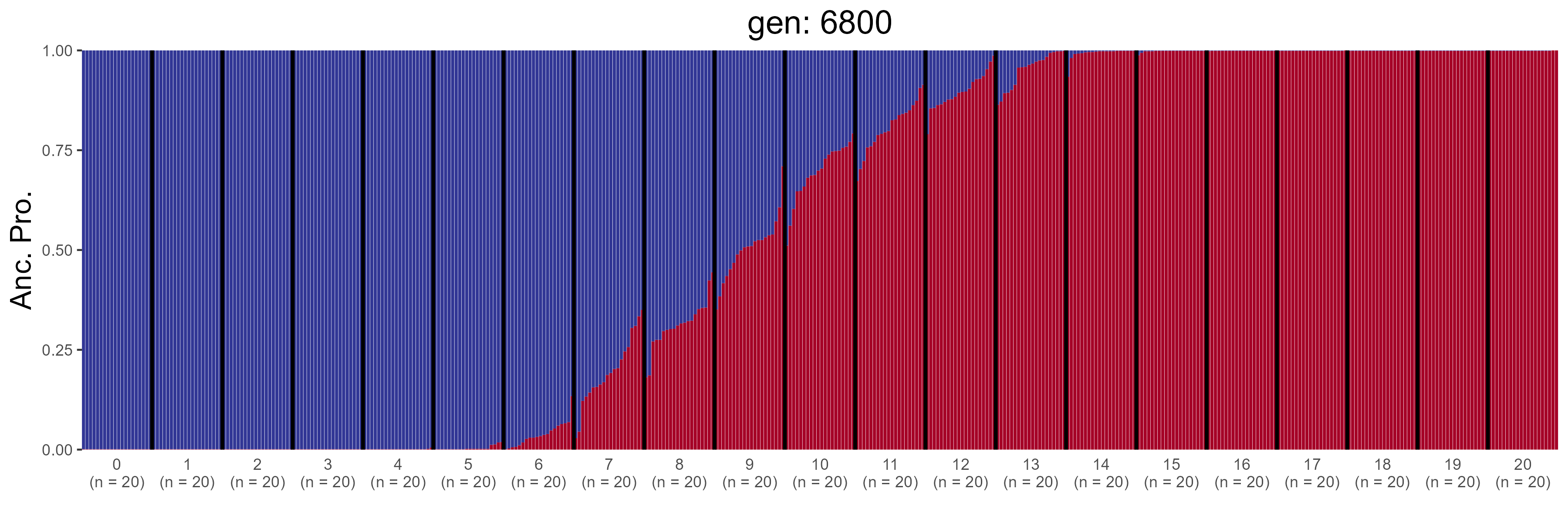

### IBD.7000.str.png

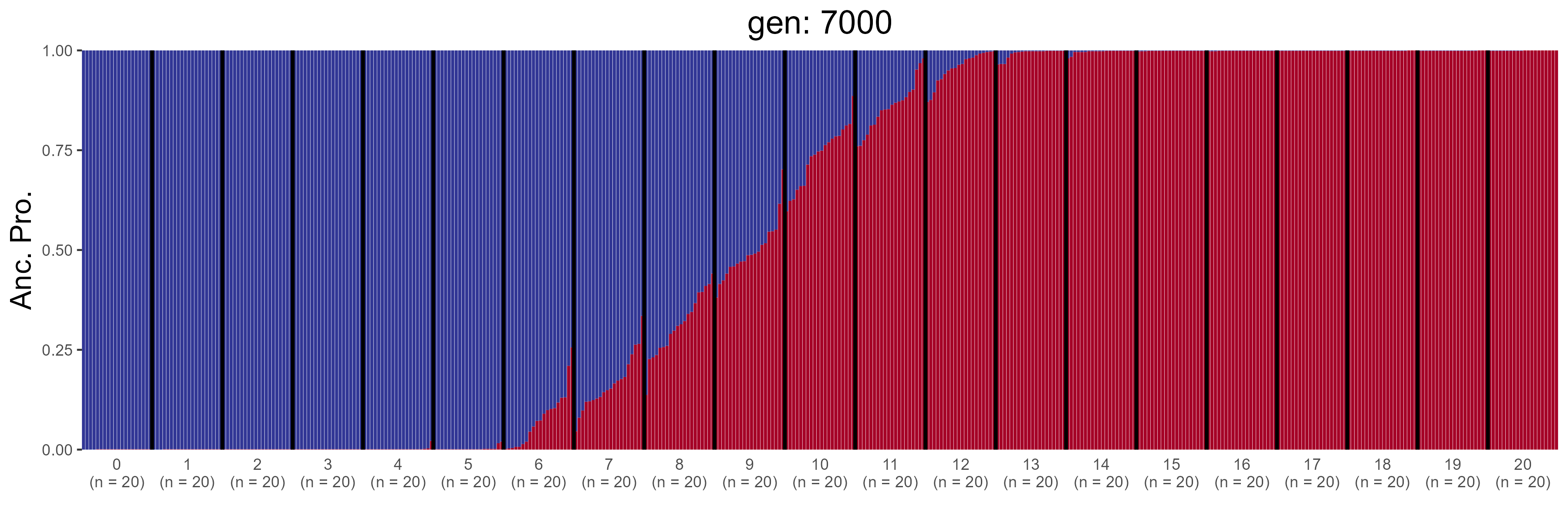

### IBD.7200.str.png

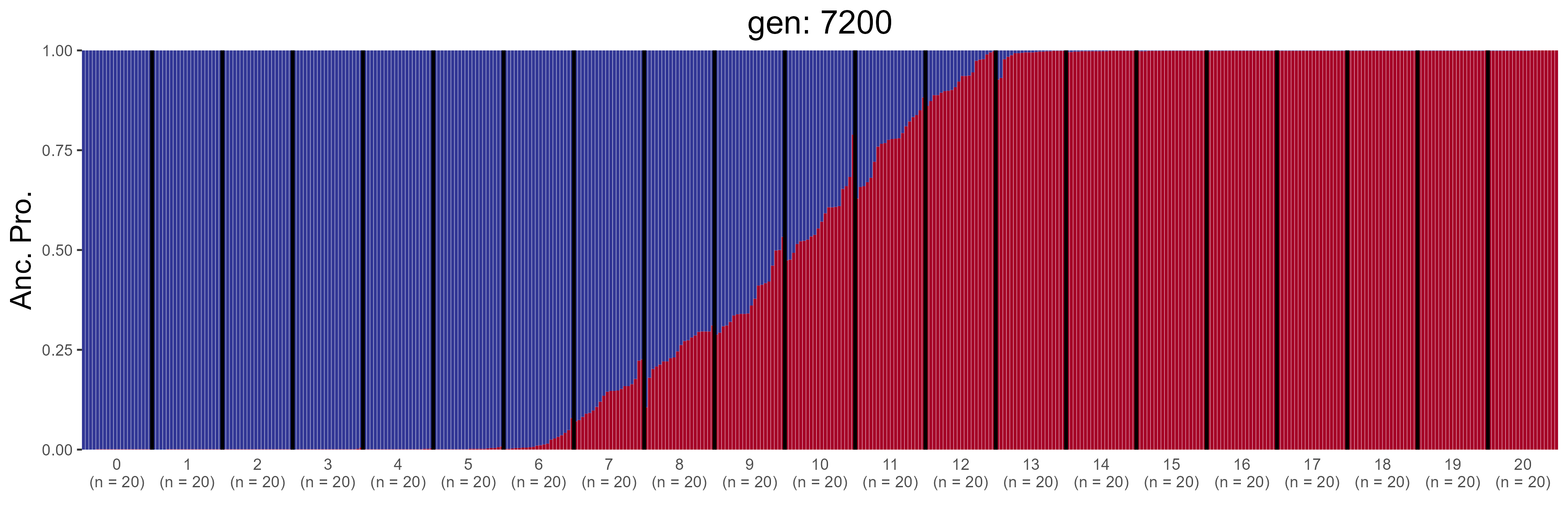

### IBD.7400.str.png

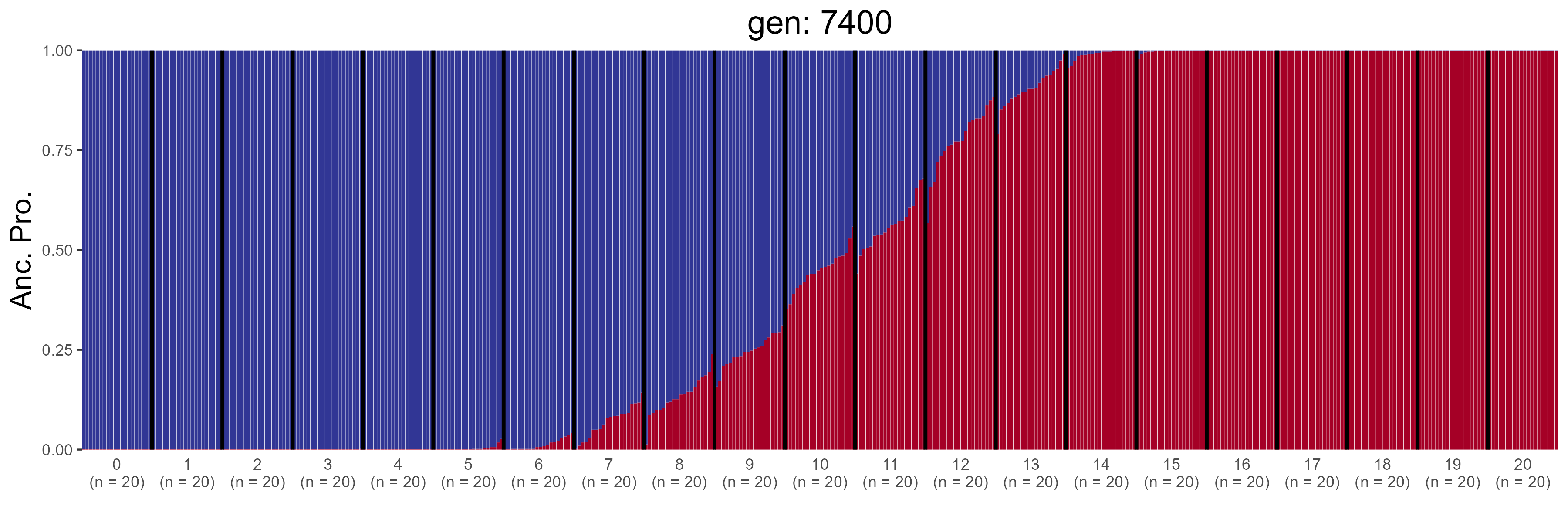

### IBD.7600.str.png

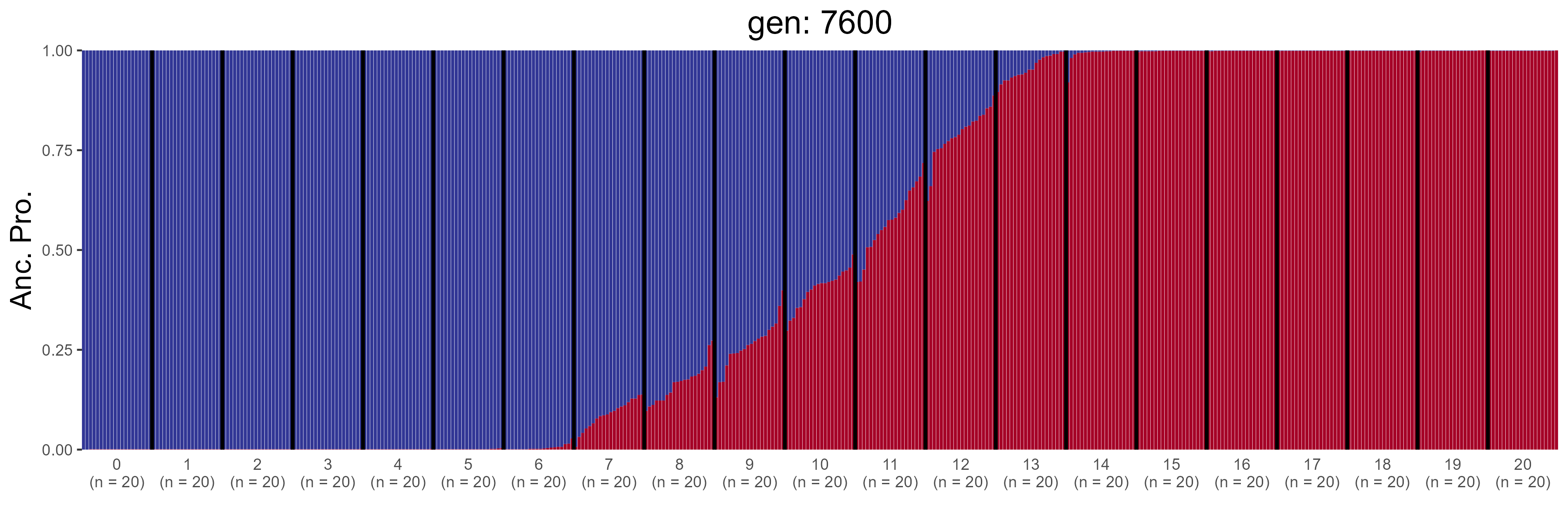

### IBD.7800.str.png

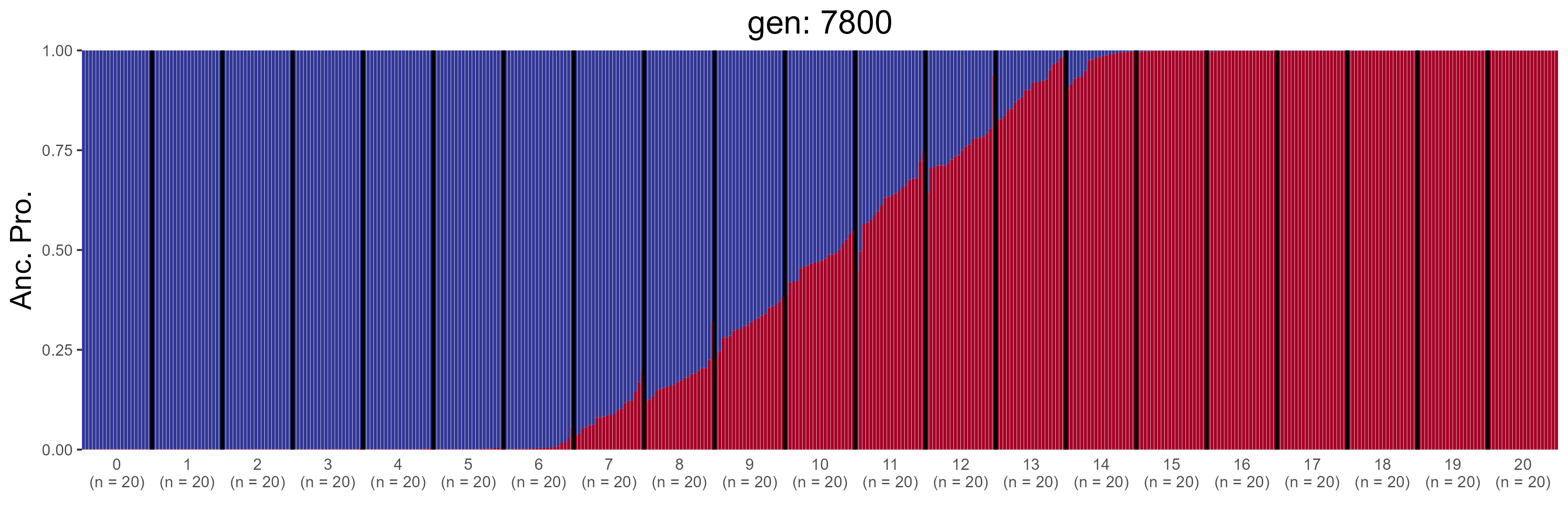

### IBD.8000.str.png

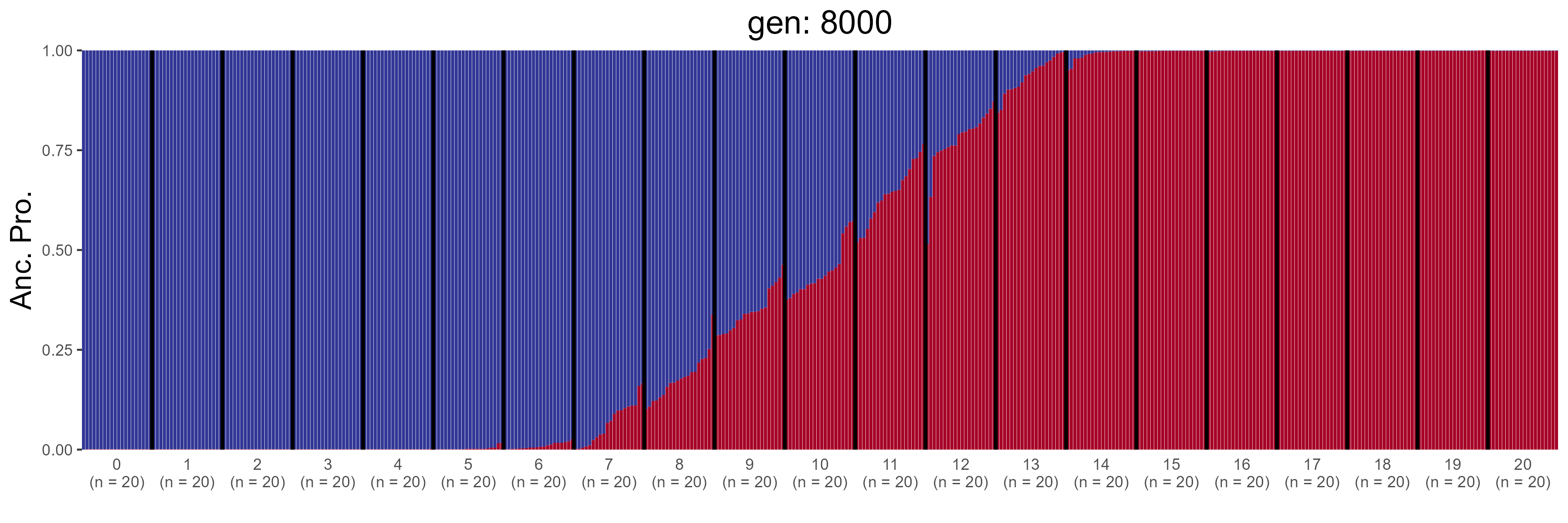

### IBD.8200.str.png

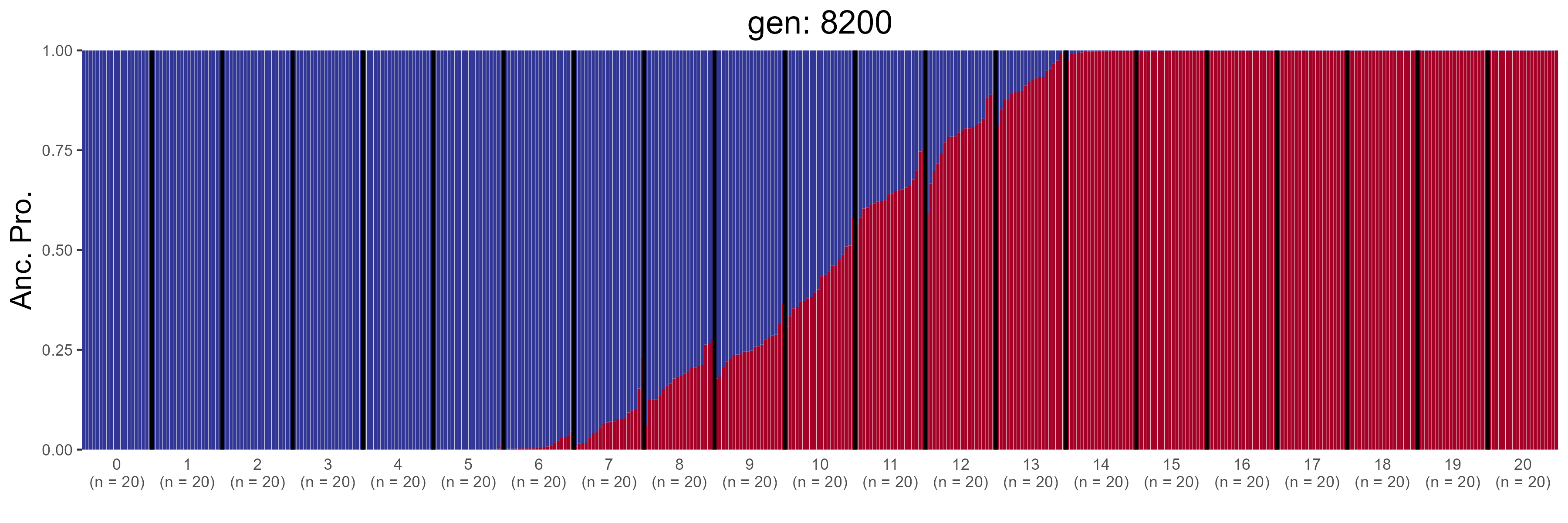

### IBD.8400.str.png

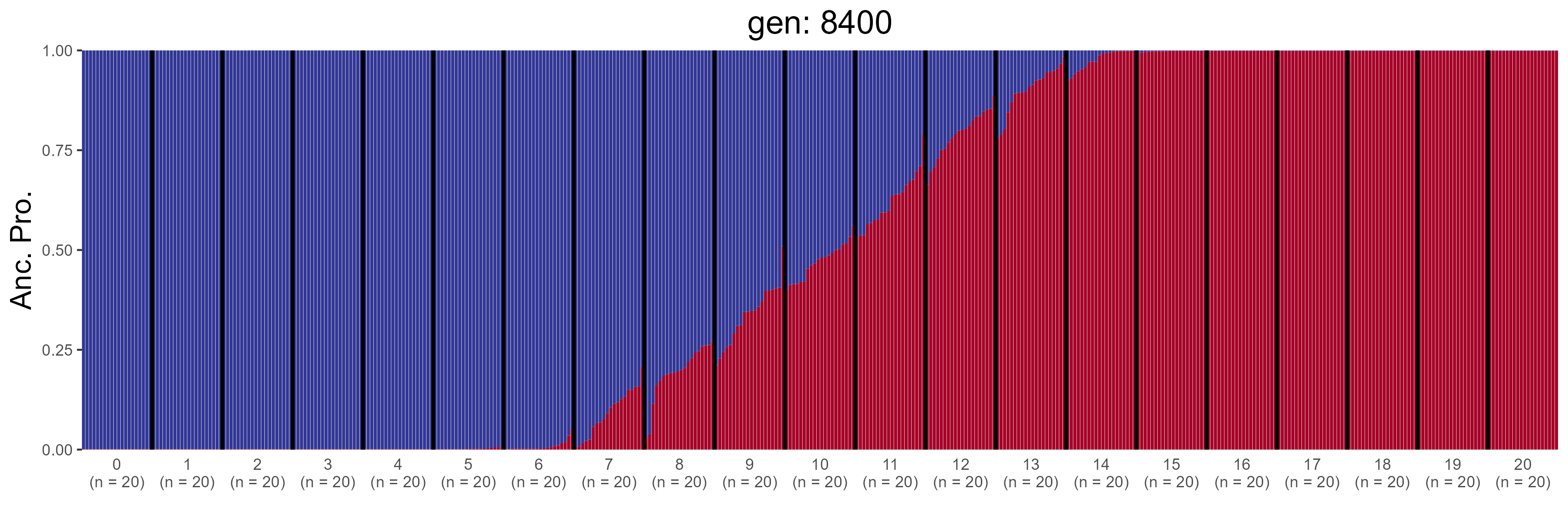

### IBD.8600.str.png

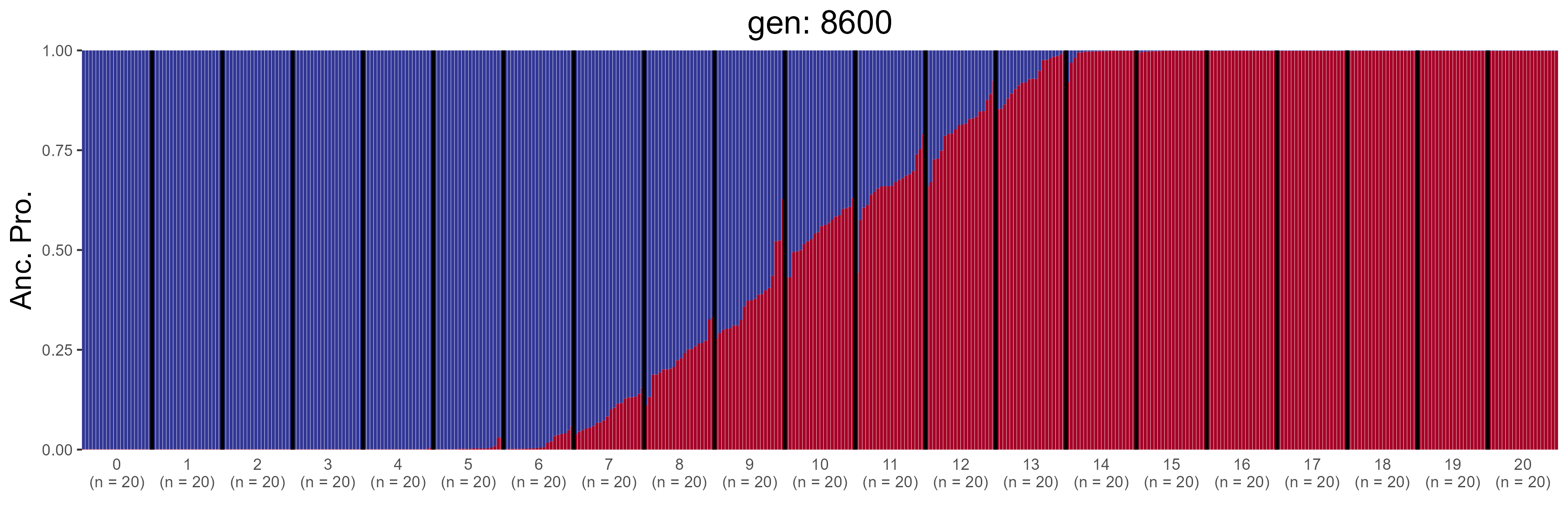

### IBD.8800.str.png

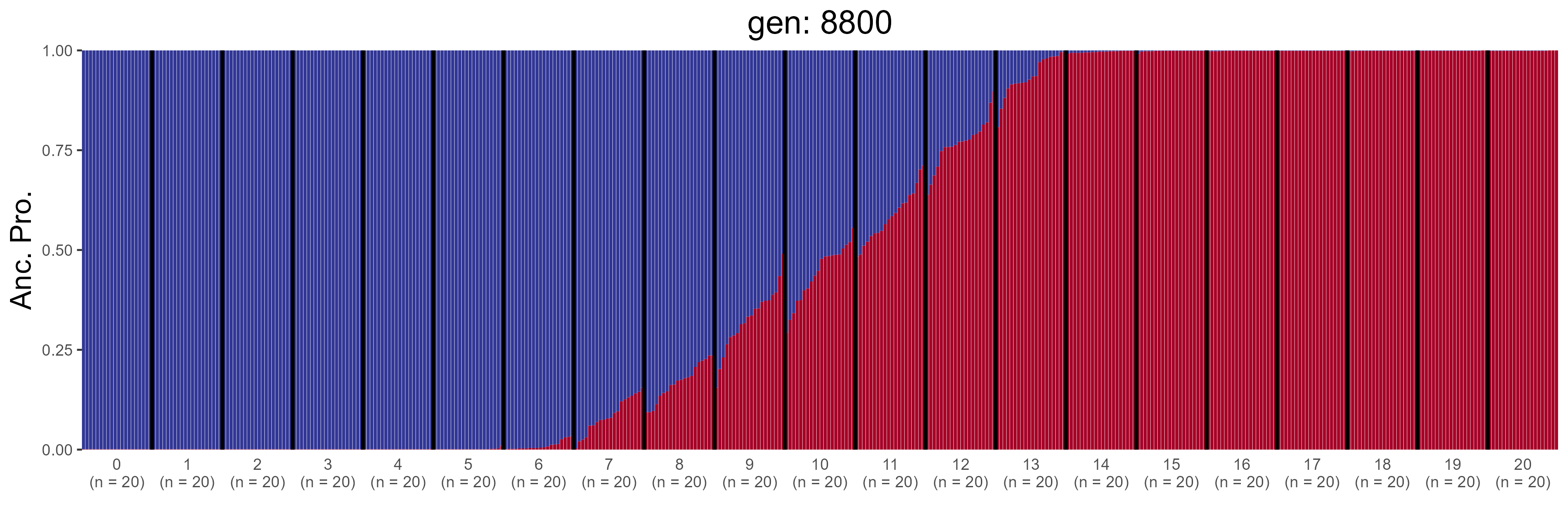

### IBD.9000.str.png

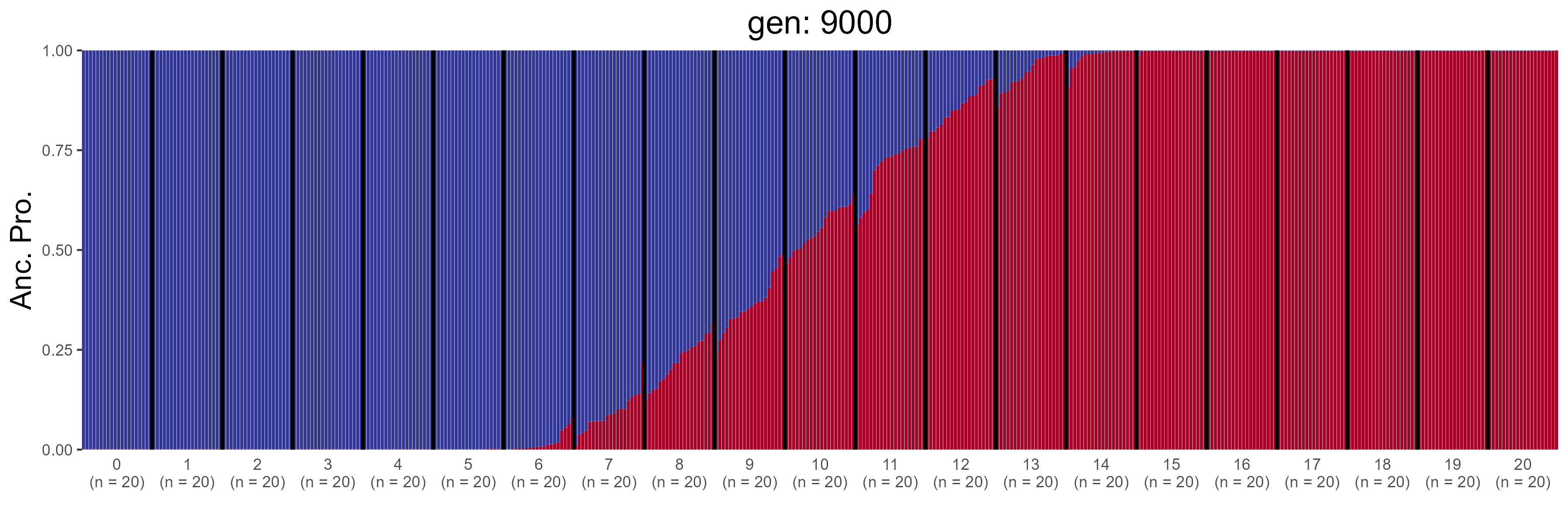

### IBD.9200.str.png

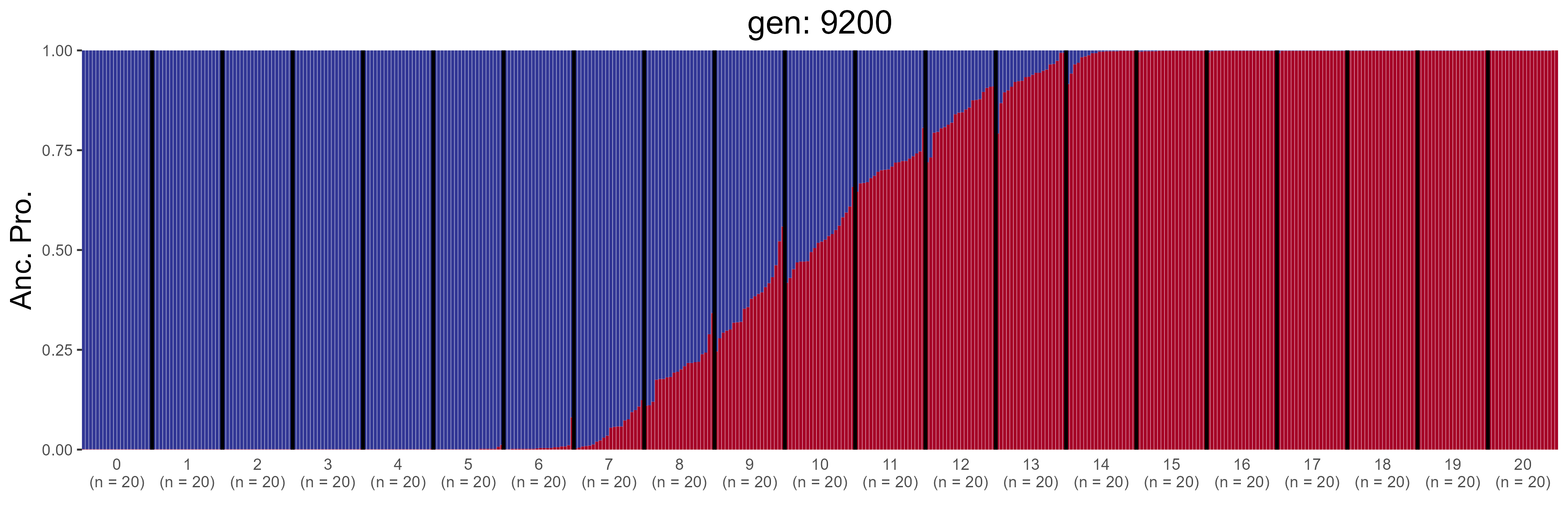

### IBD.9400.str.png

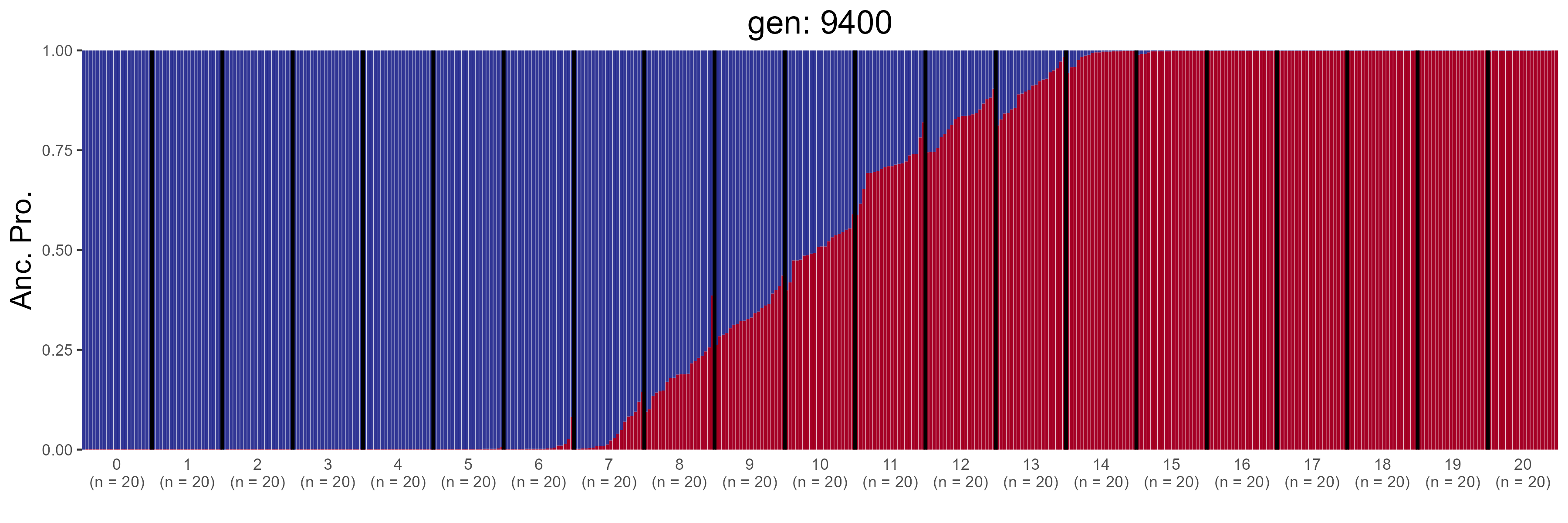

### IBD.9600.str.png

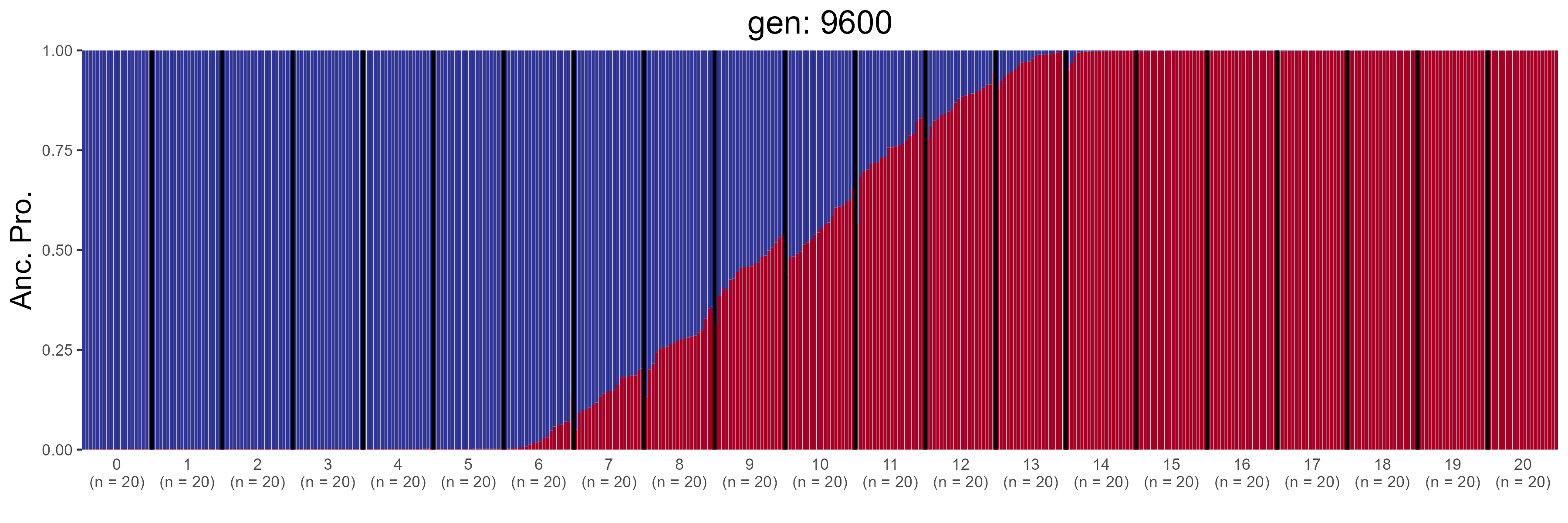

### IBD.9800.str.png

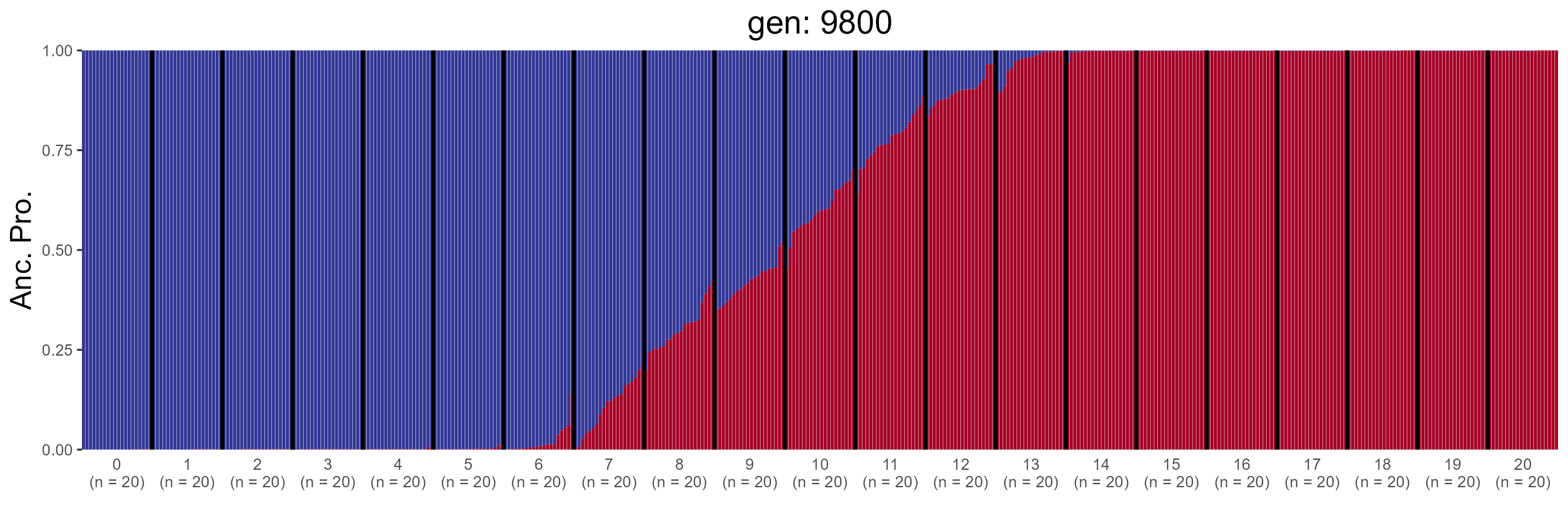

### IBD.10400.str.png

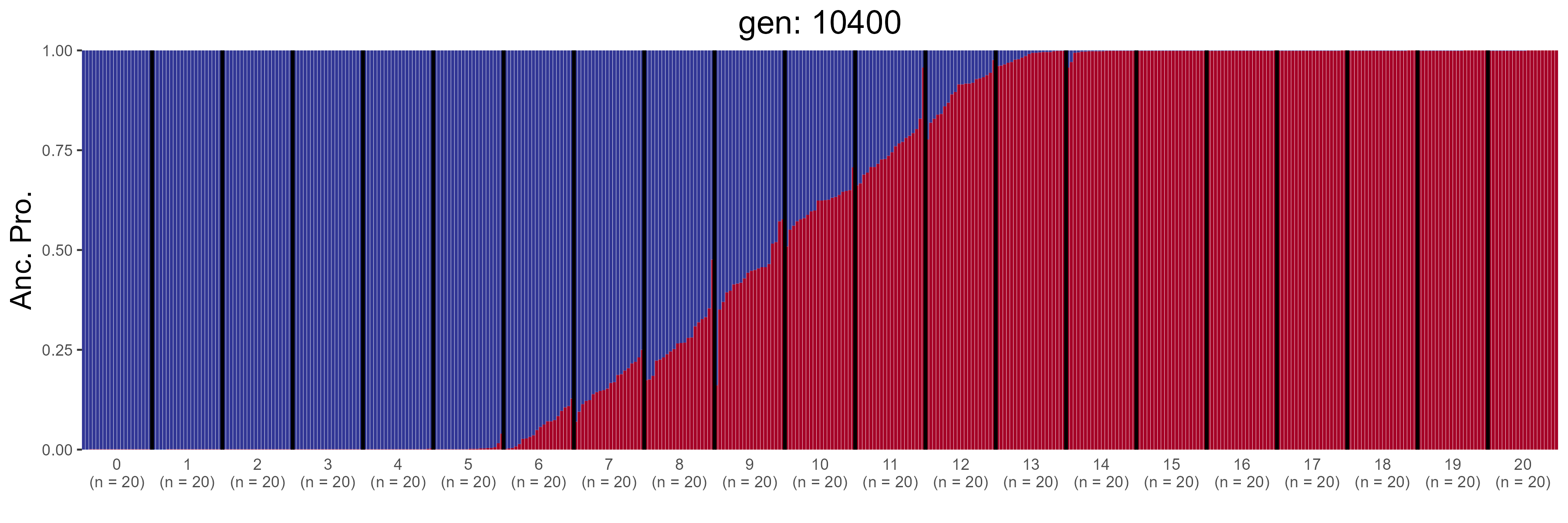

### IBD.10600.str.png

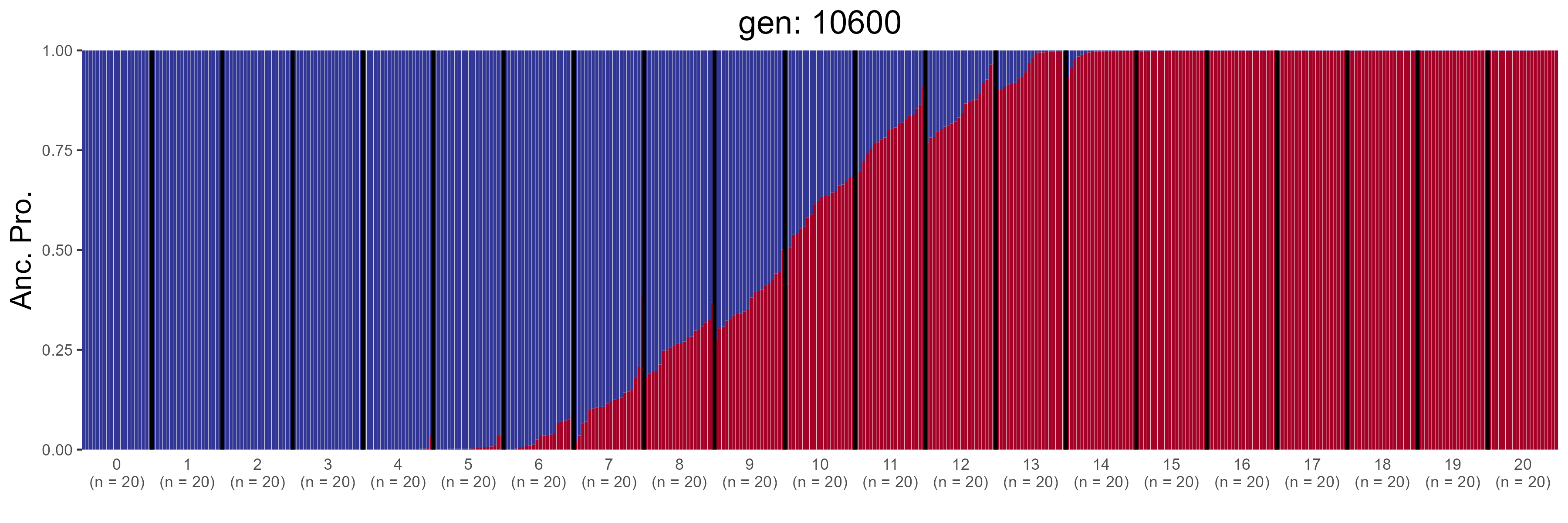

### IBD.10800.str.png

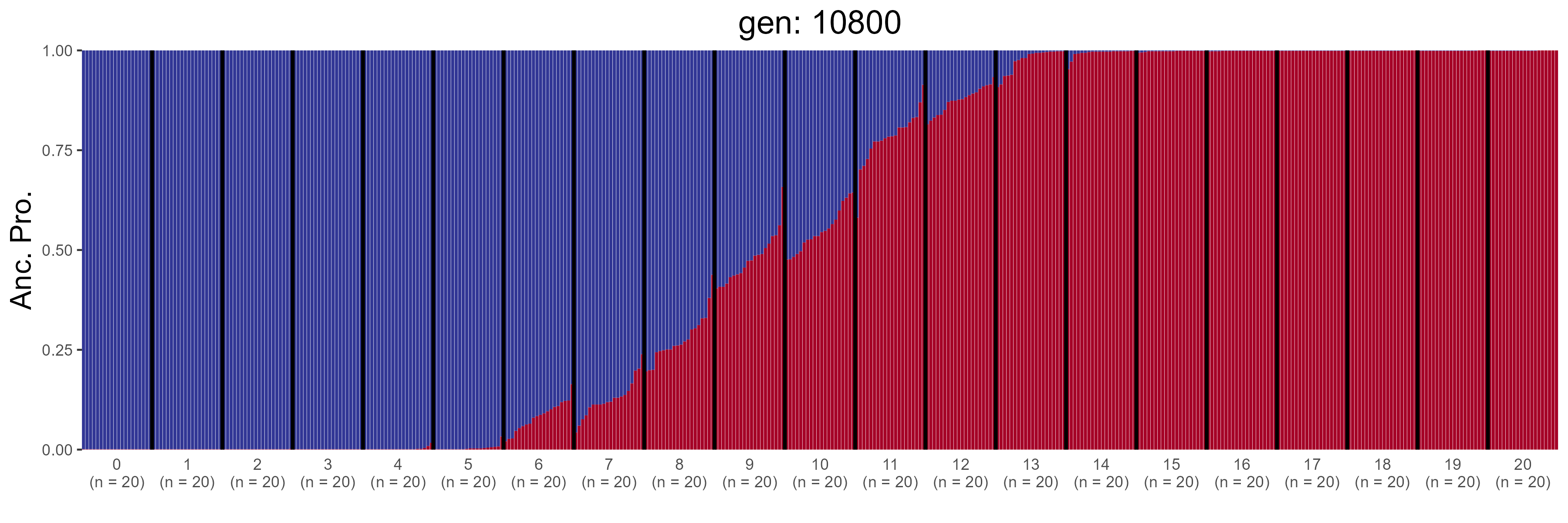

### IBD.11000.str.png

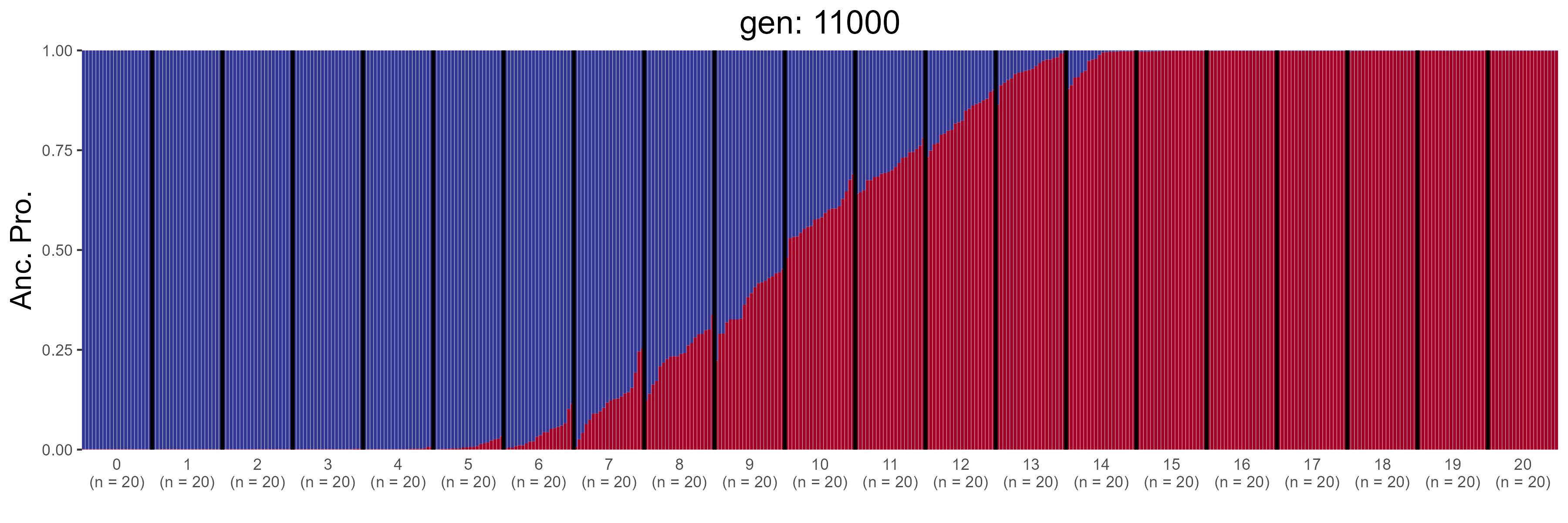

### IBD.11200.str.png

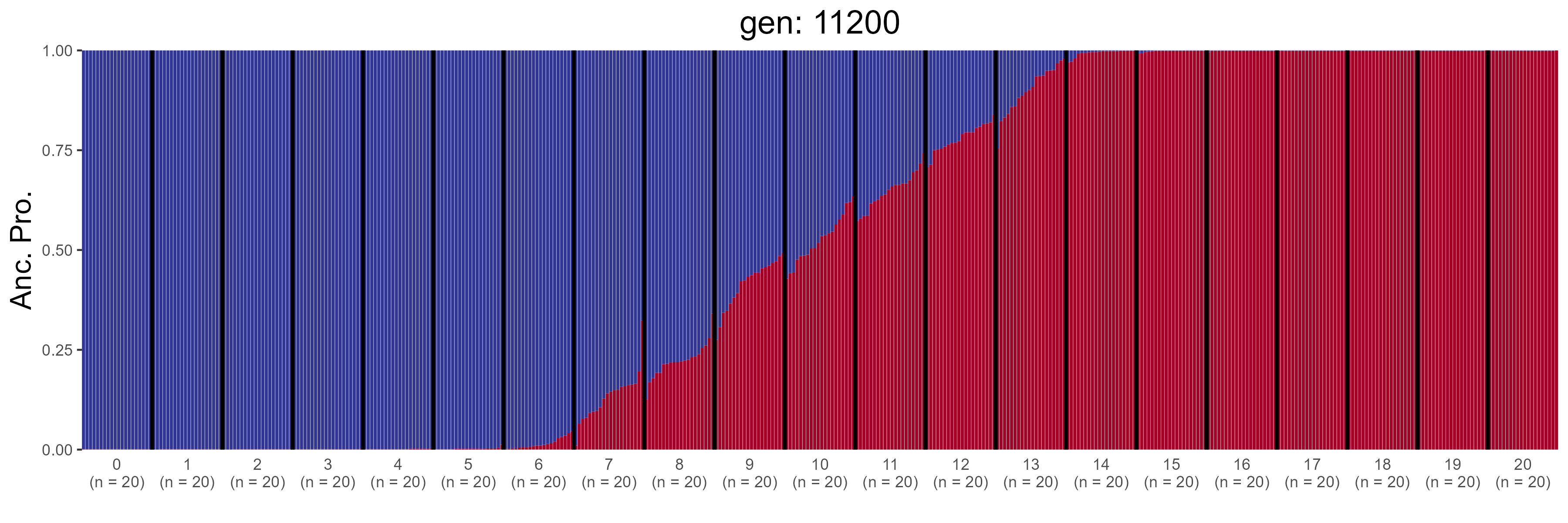

### IBD.11400.str.png

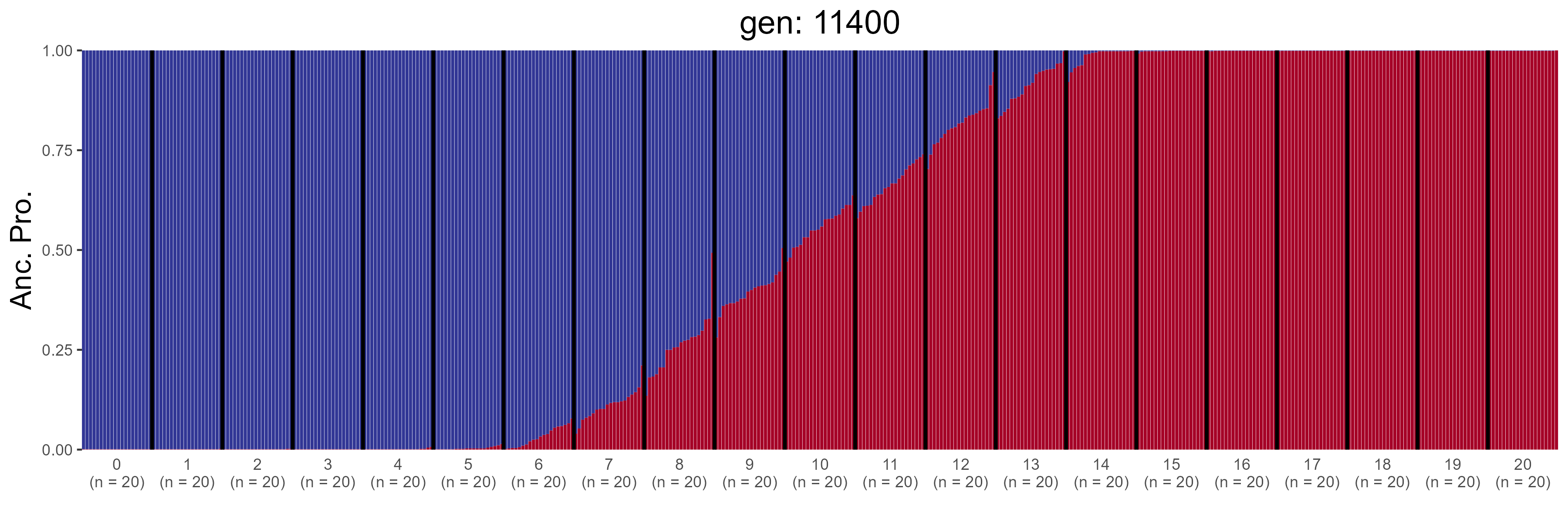

### IBD.11600.str.png

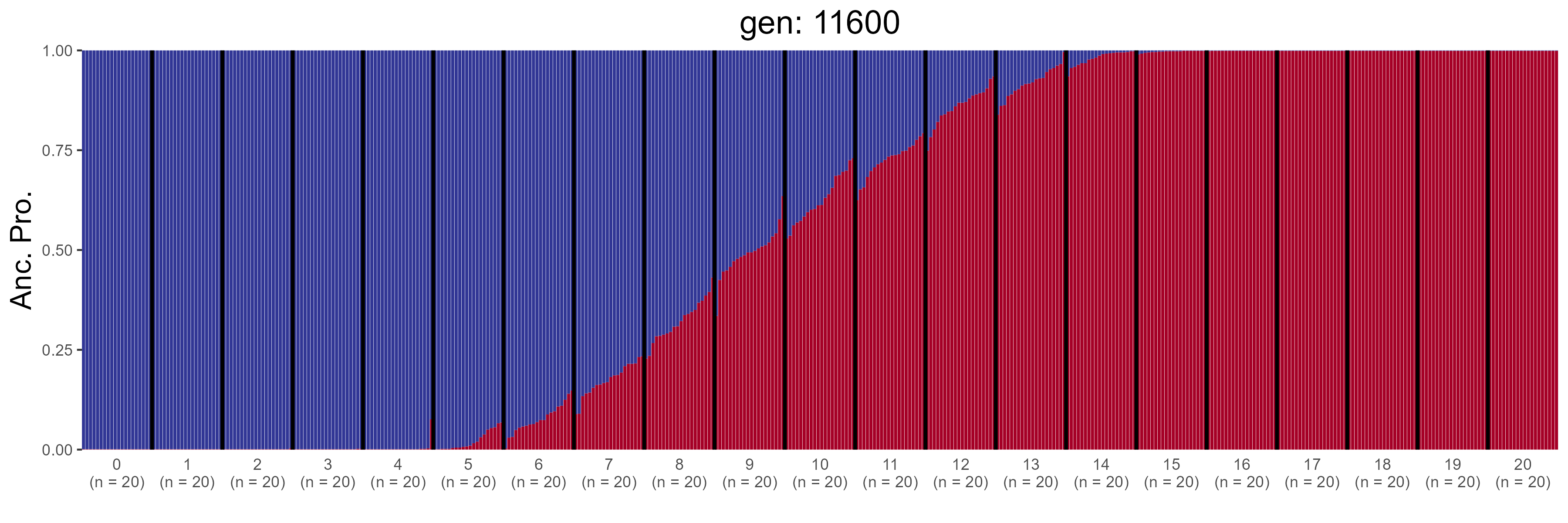

### IBD.11800.str.png

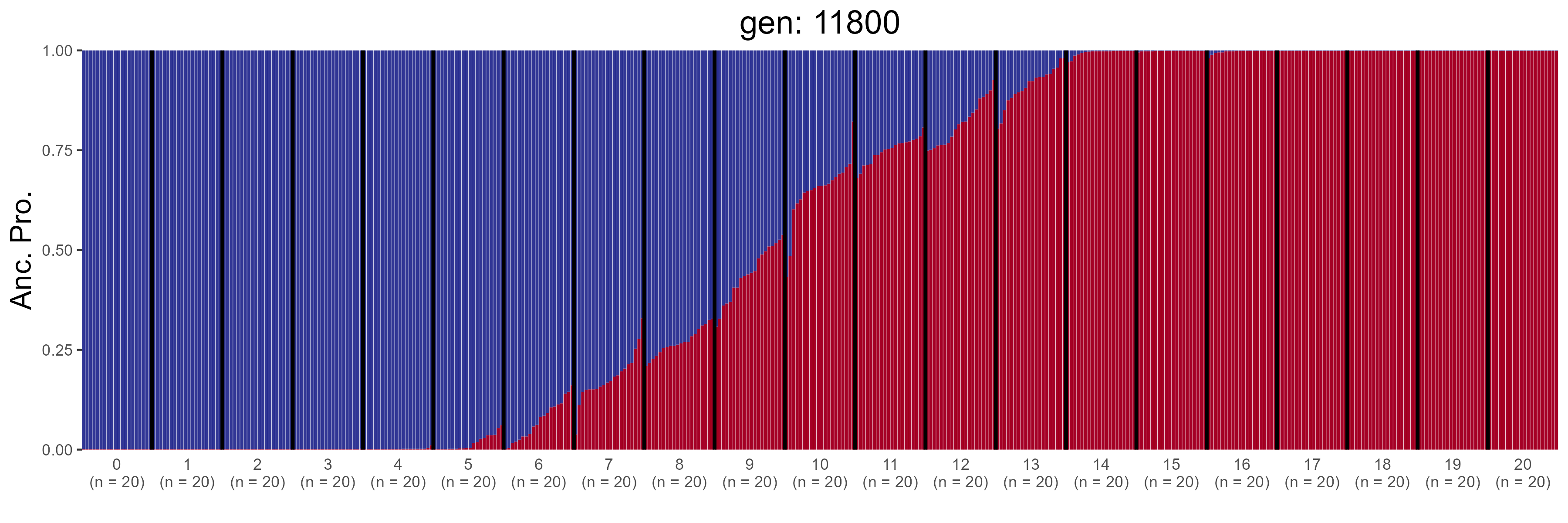

### IBD.12000.str.png

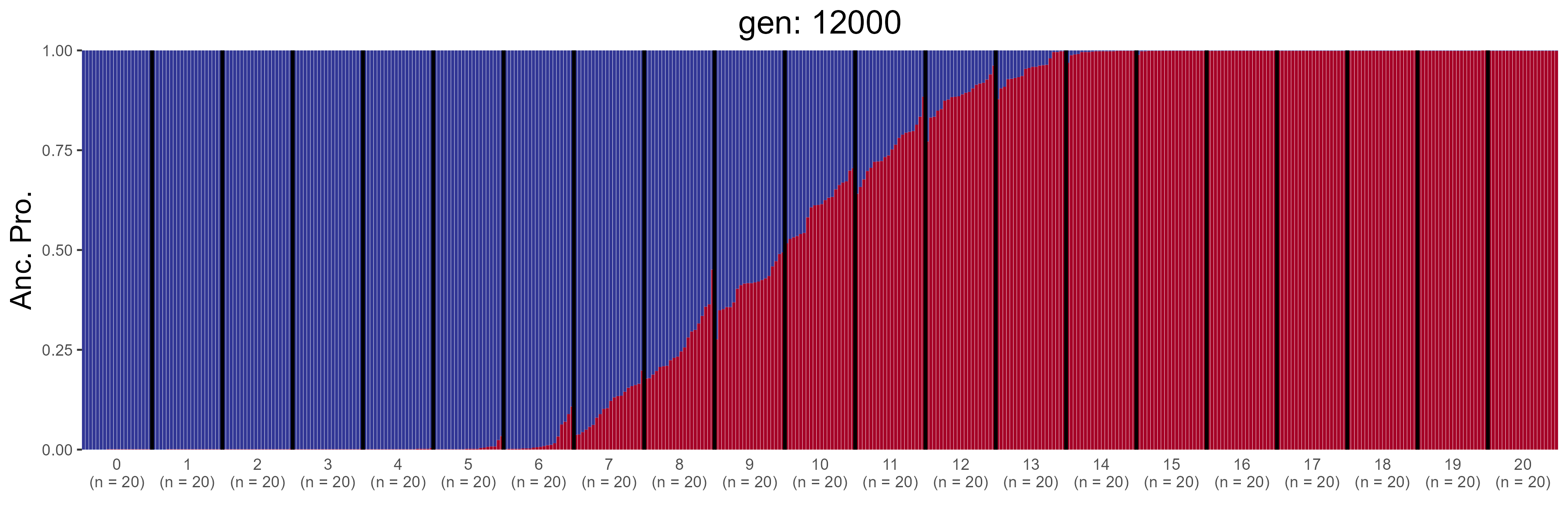

### IBD.12200.str.png

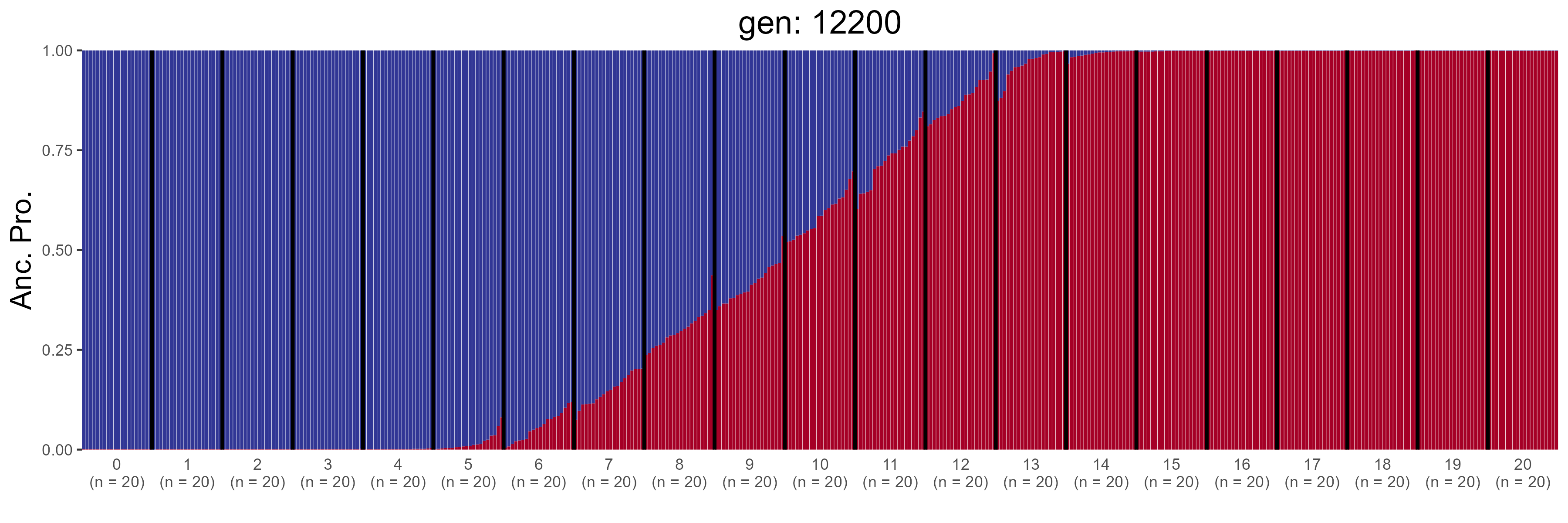

### IBD.12400.str.png

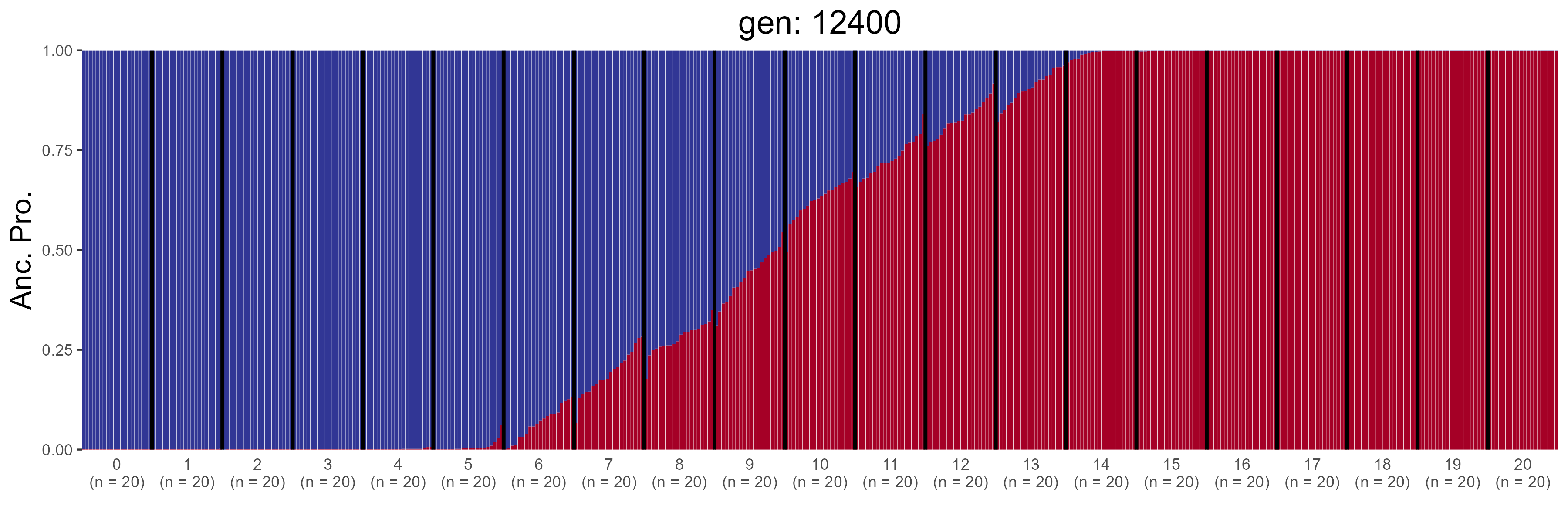

### IBD.12600.str.png

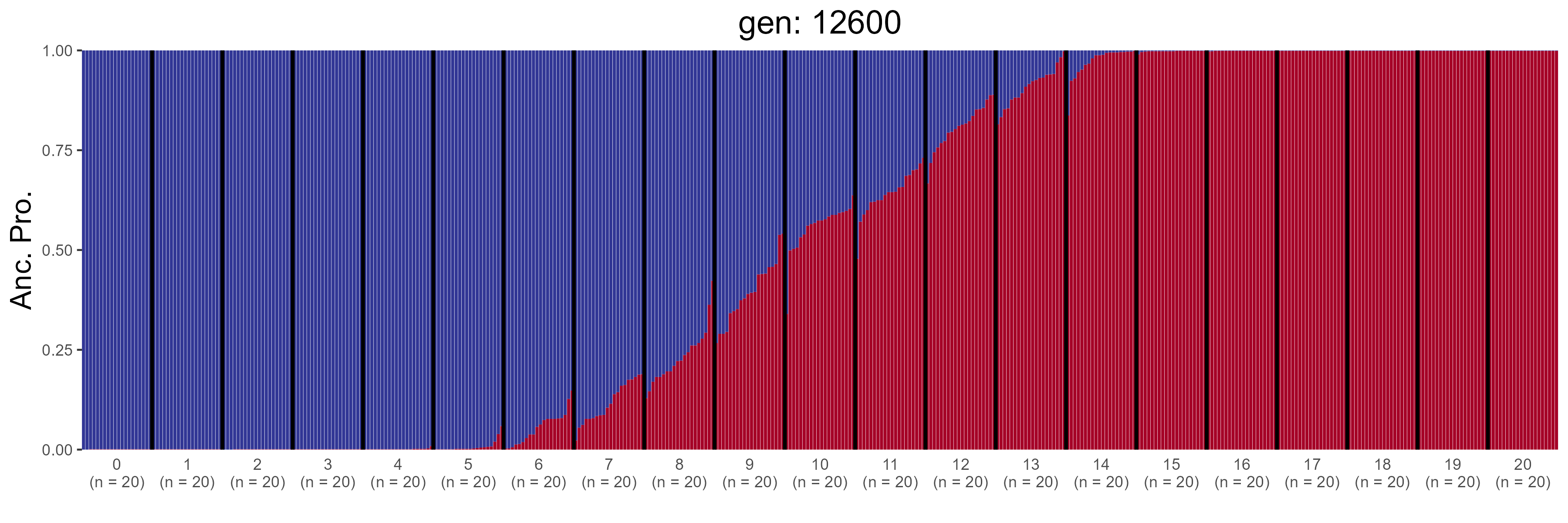

### IBD.12800.str.png

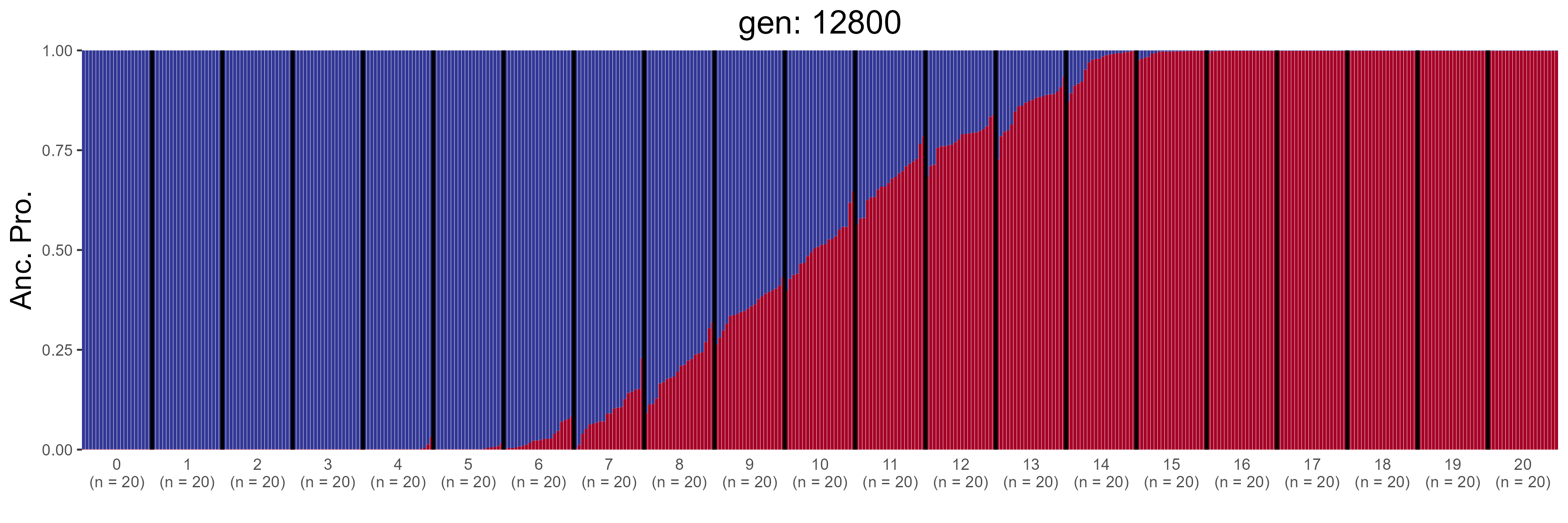

### IBD.13000.str.png

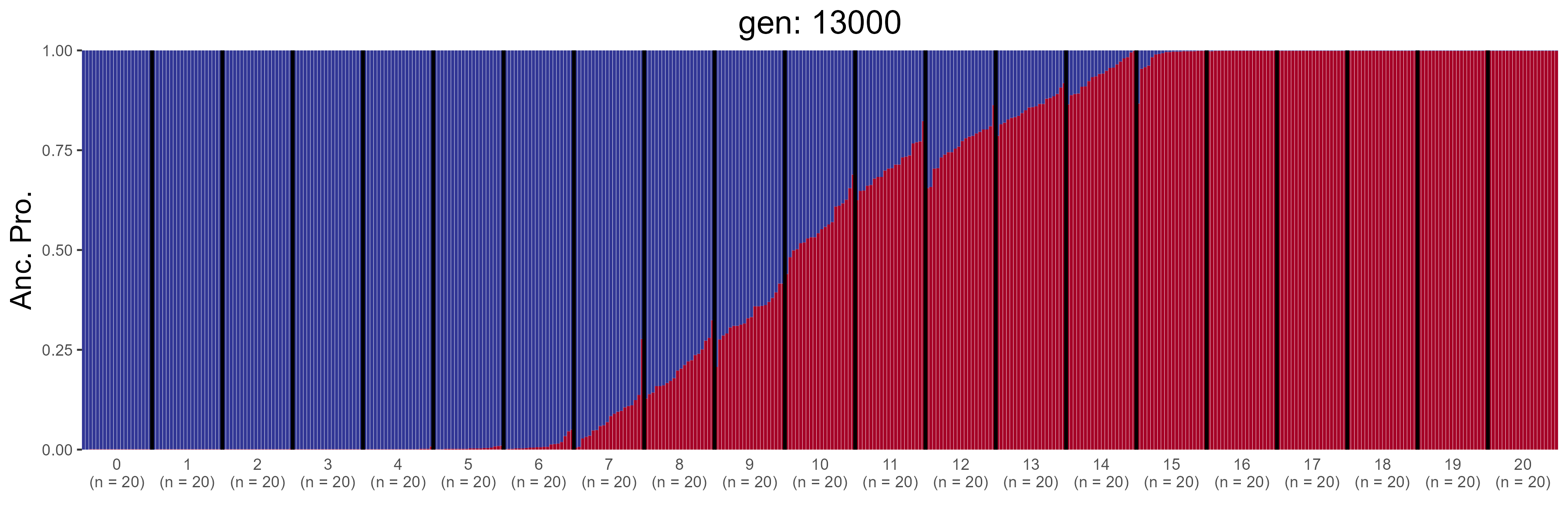

### IBD.13200.str.png

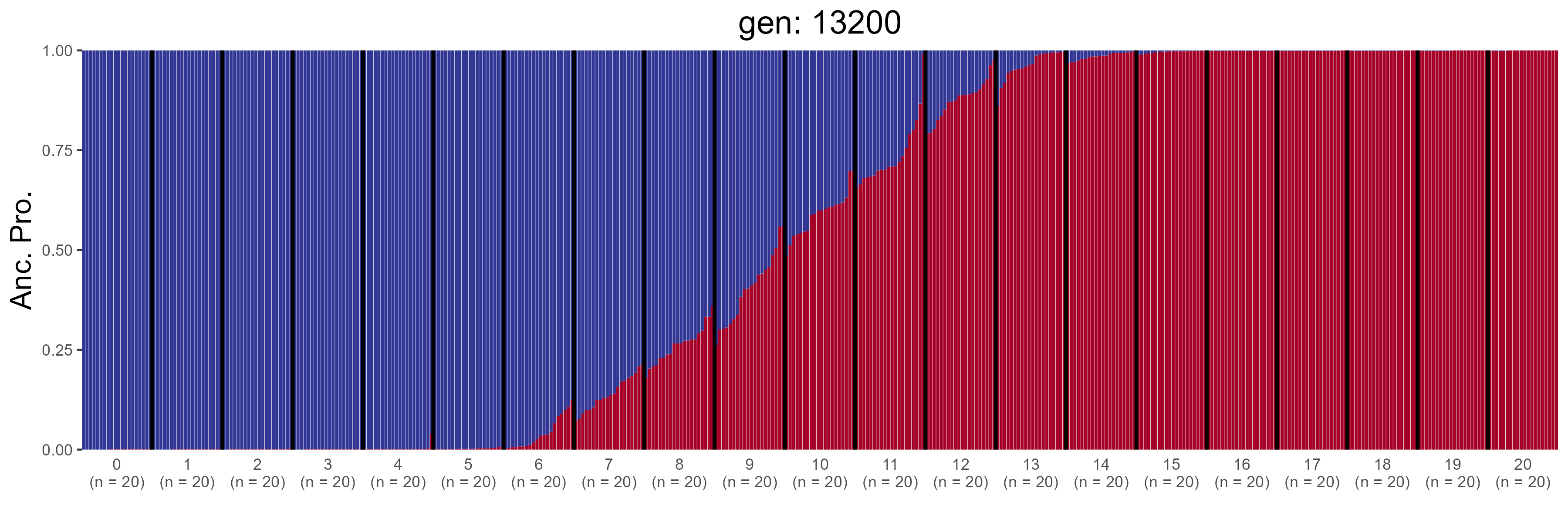

### IBD.13400.str.png

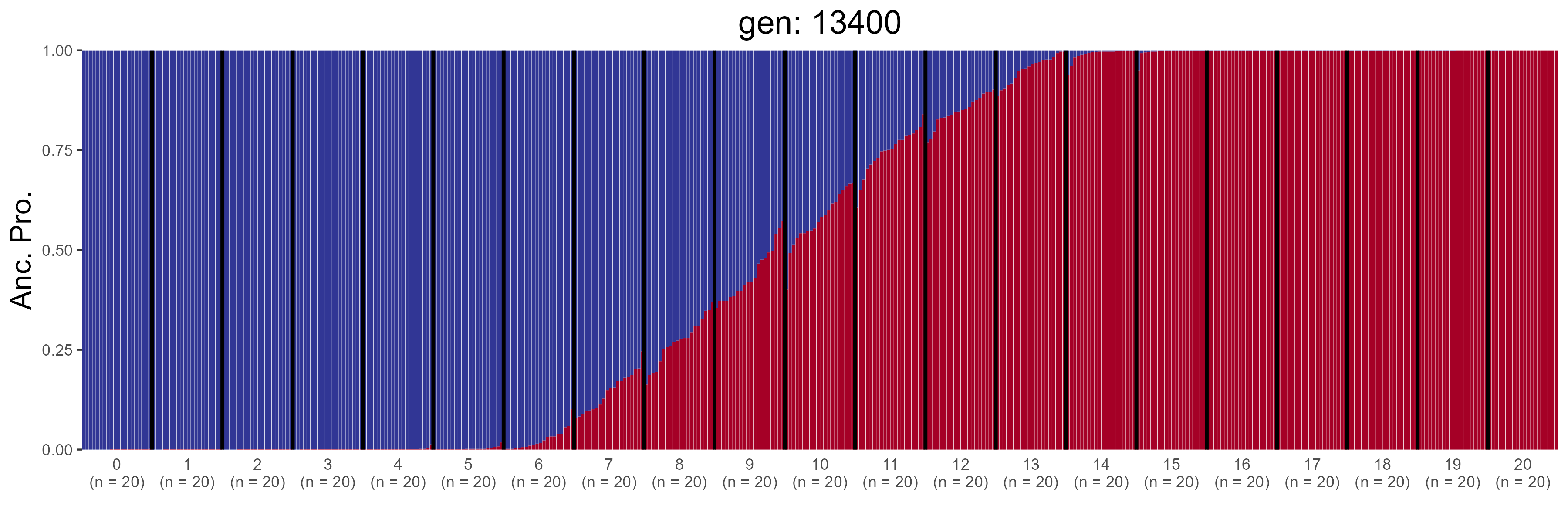

### IBD.13600.str.png

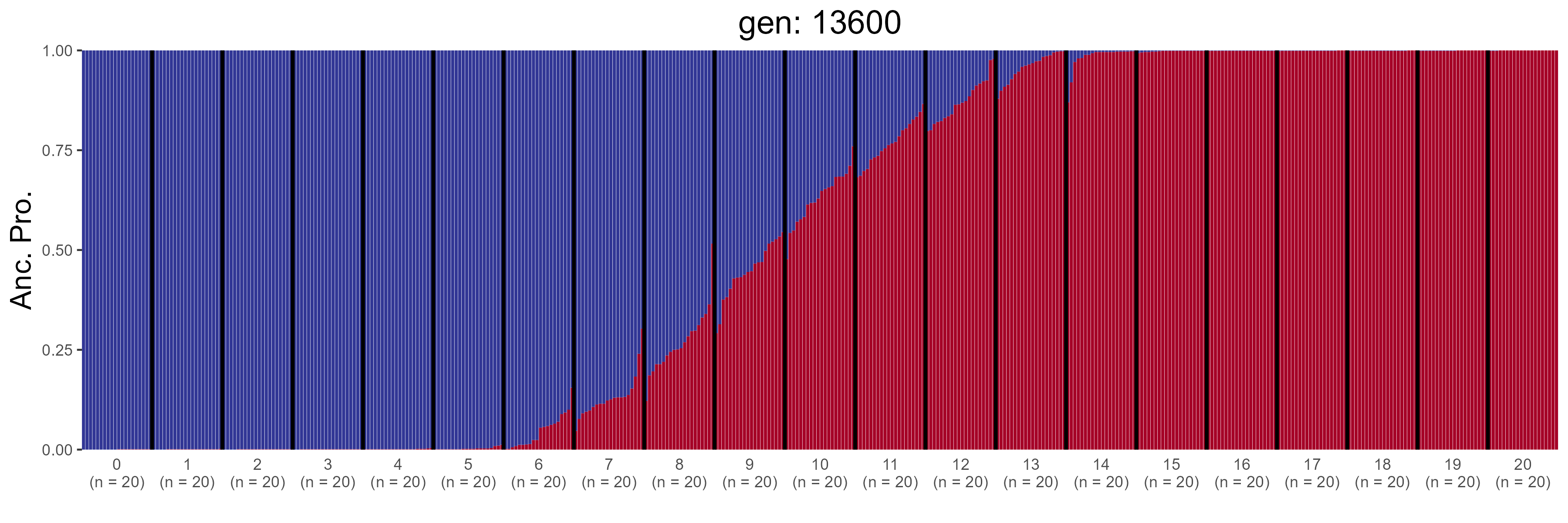

### IBD.13800.str.png

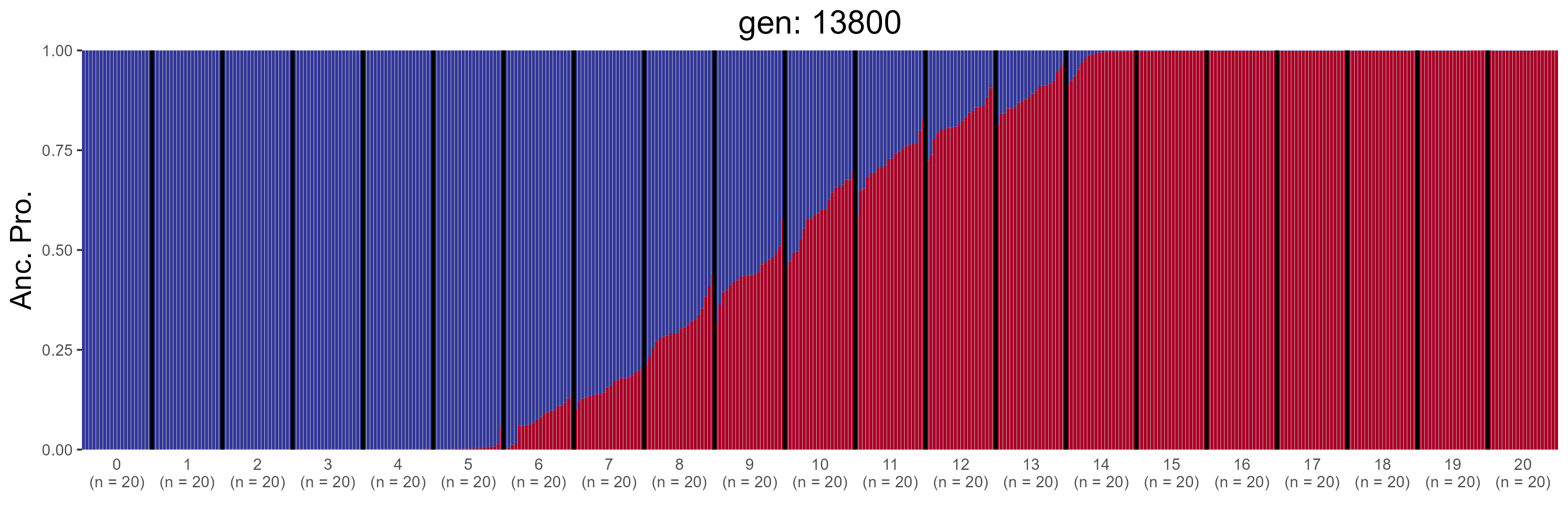

### IBD.14000.str.png

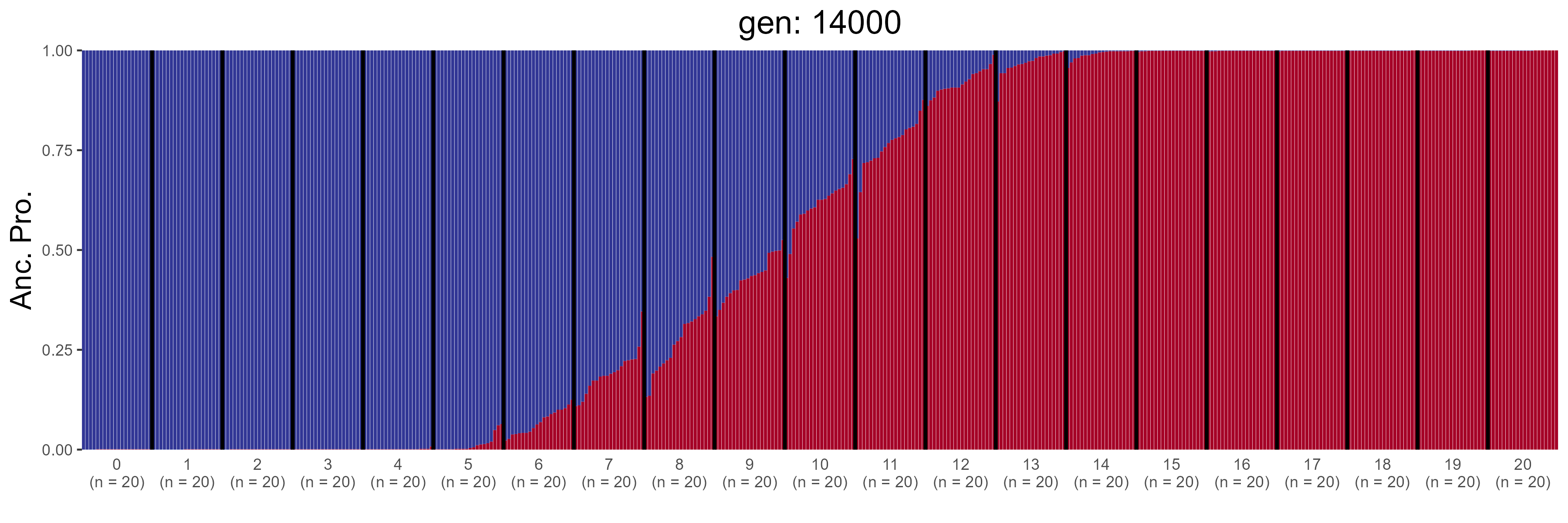

### IBD.14200.str.png

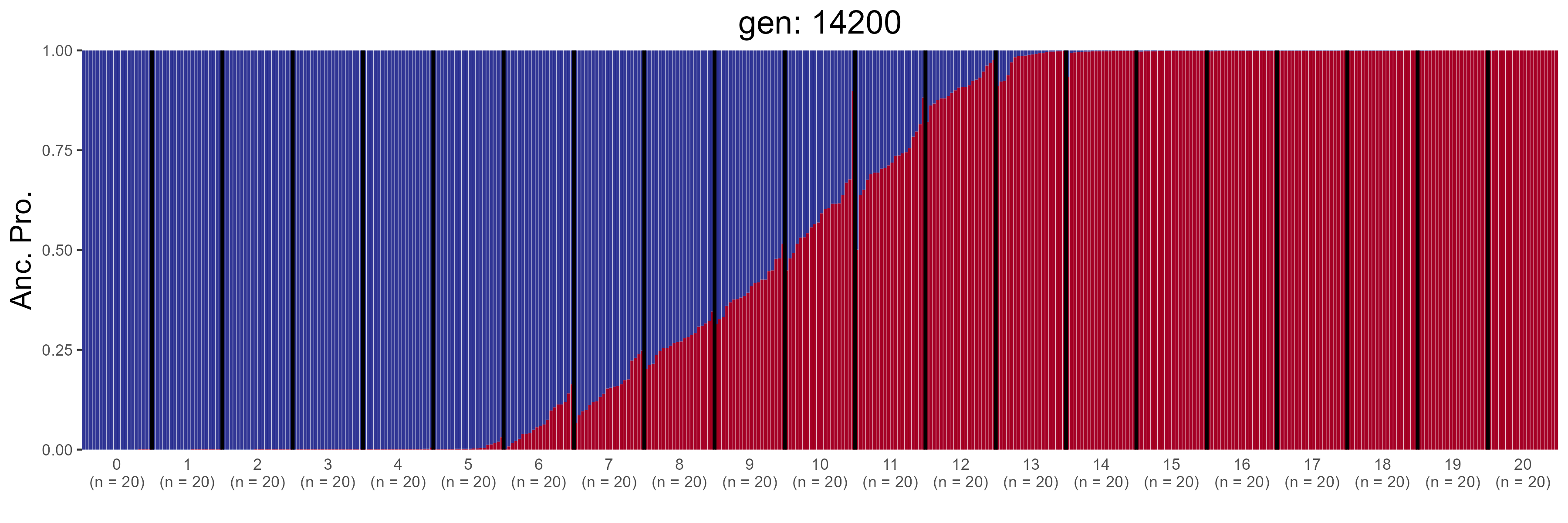

### IBD.14400.str.png

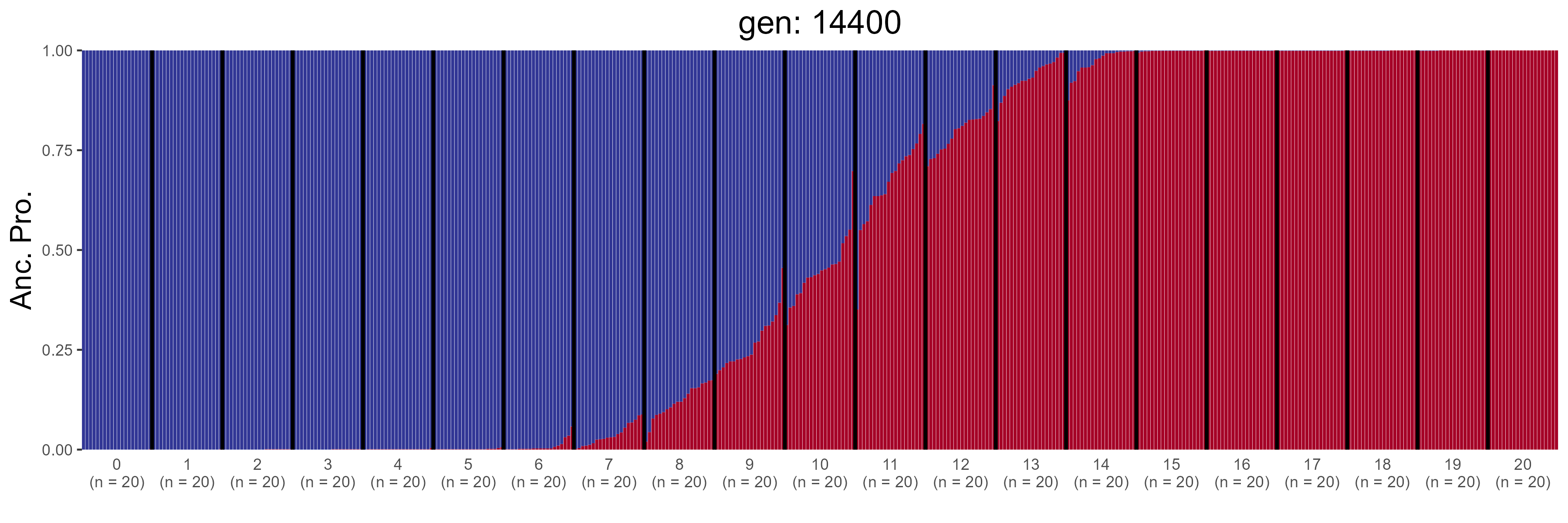

### IBD.14600.str.png

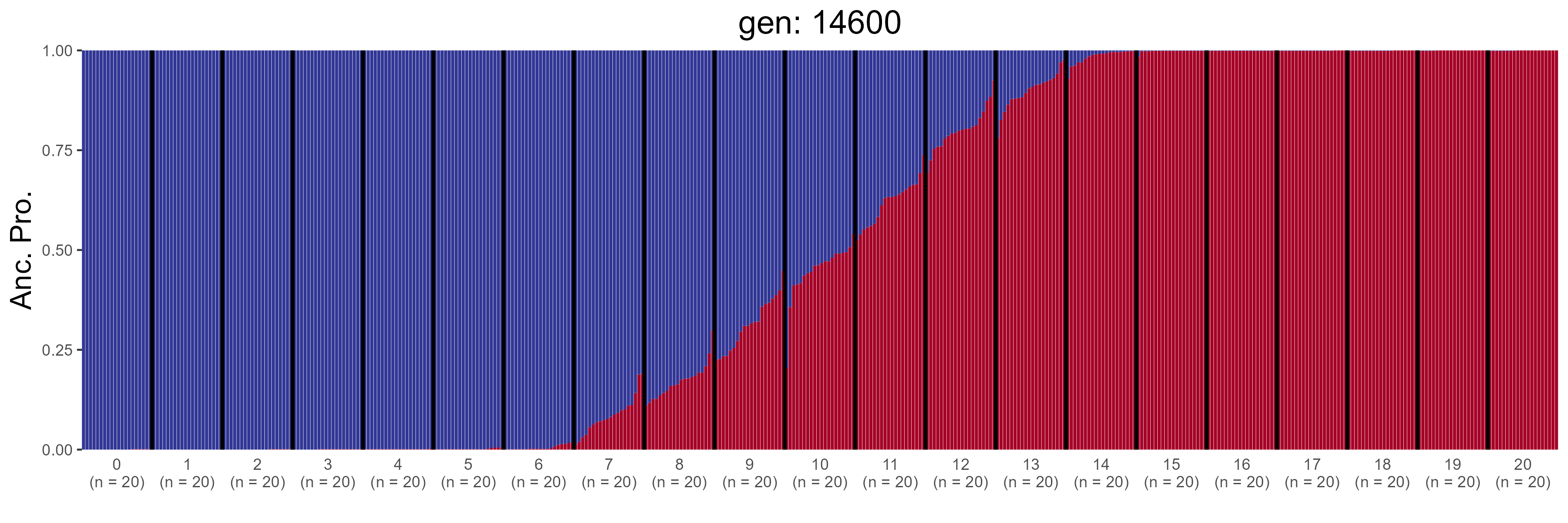

### IBD.14800.str.png

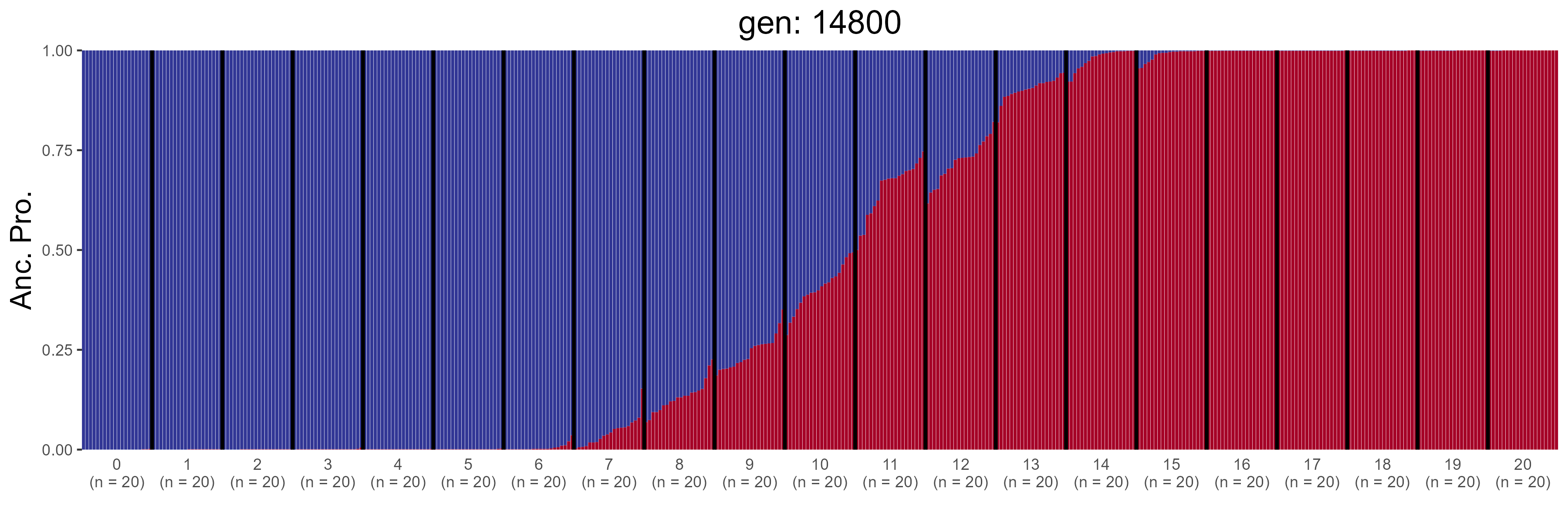

### IBD.15000.str.png

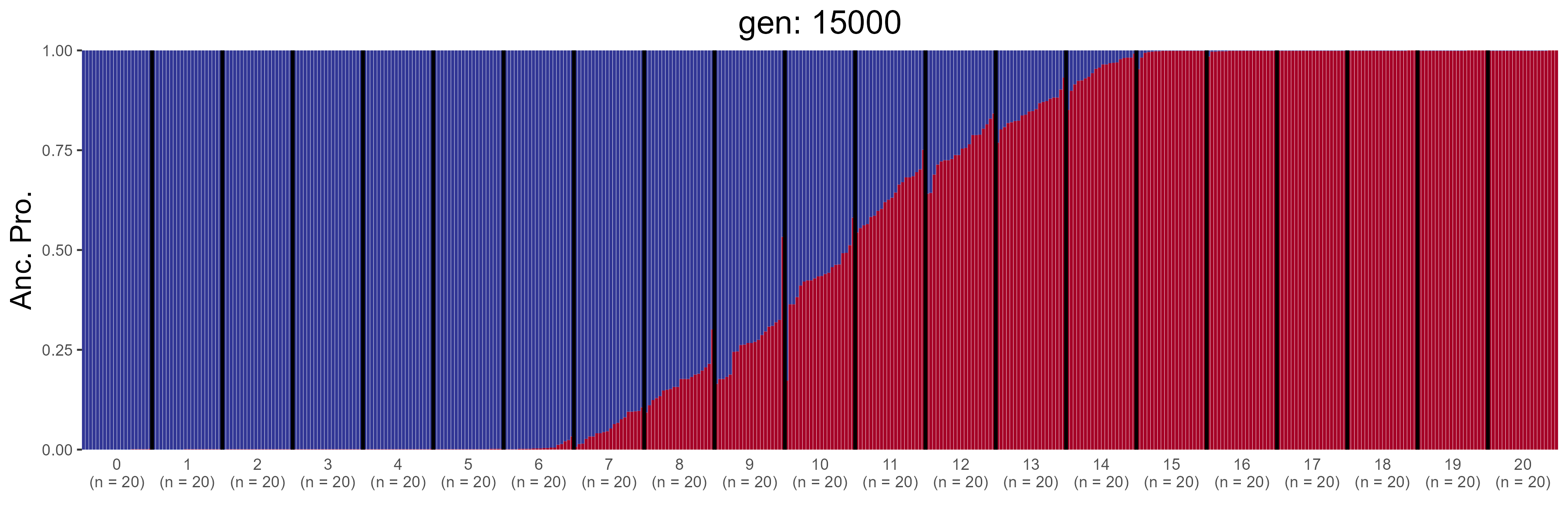

### IBD.15200.str.png

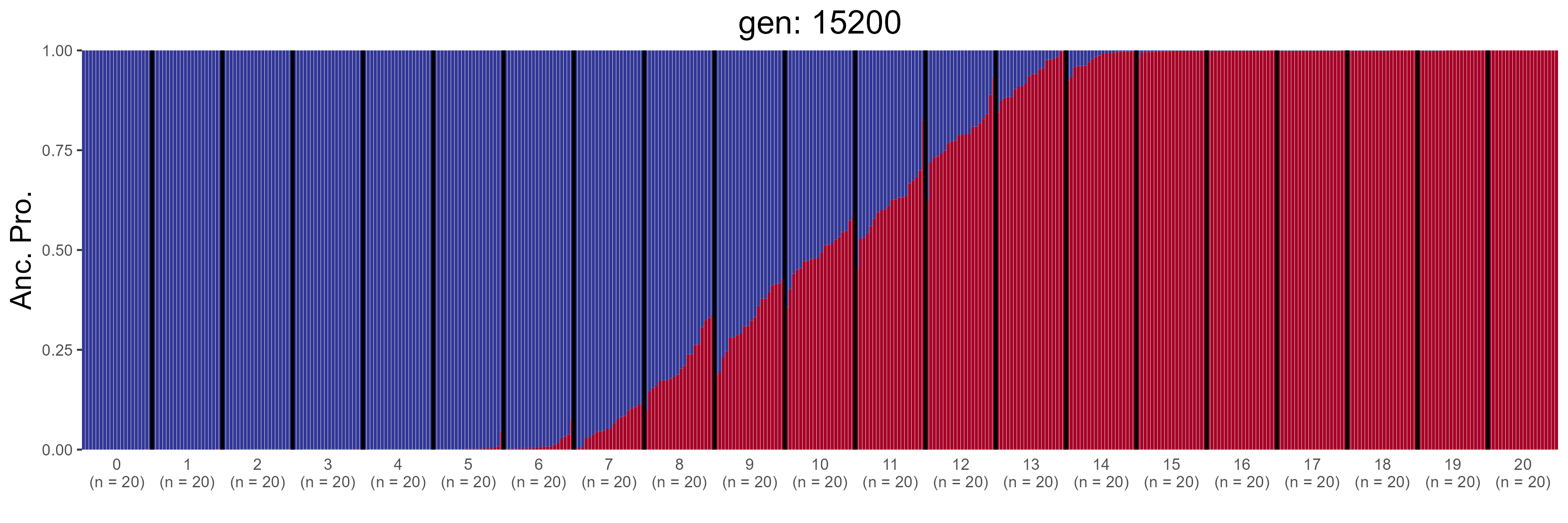

### IBD.15400.str.png

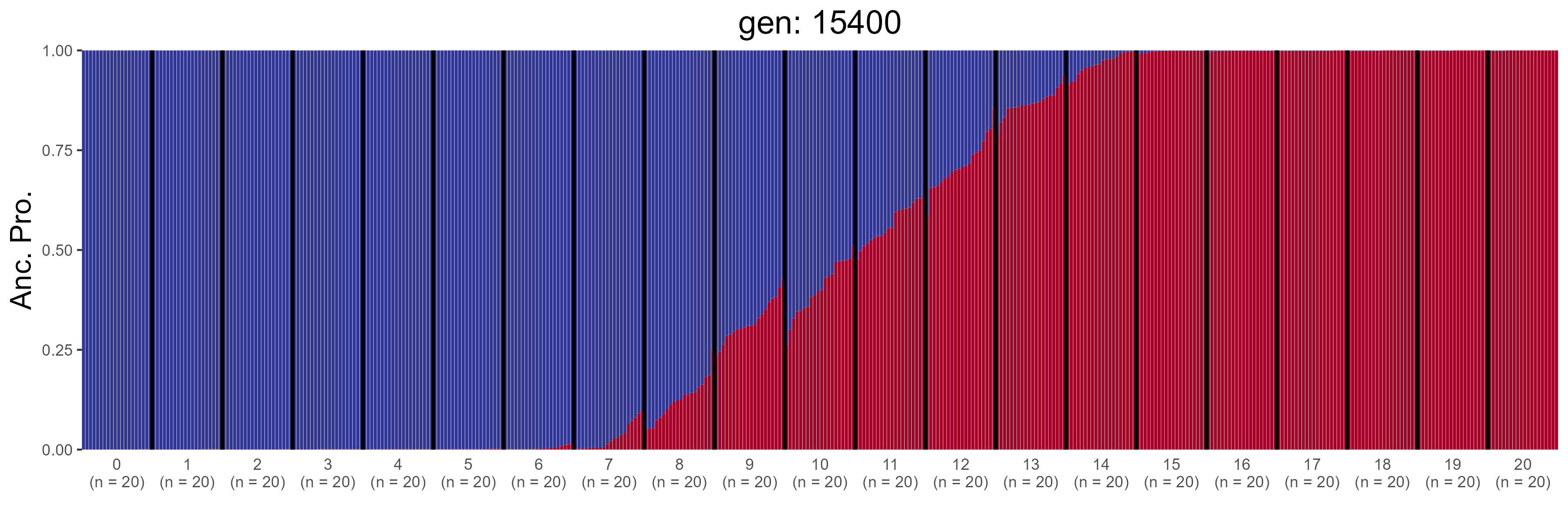

### IBD.15600.str.png

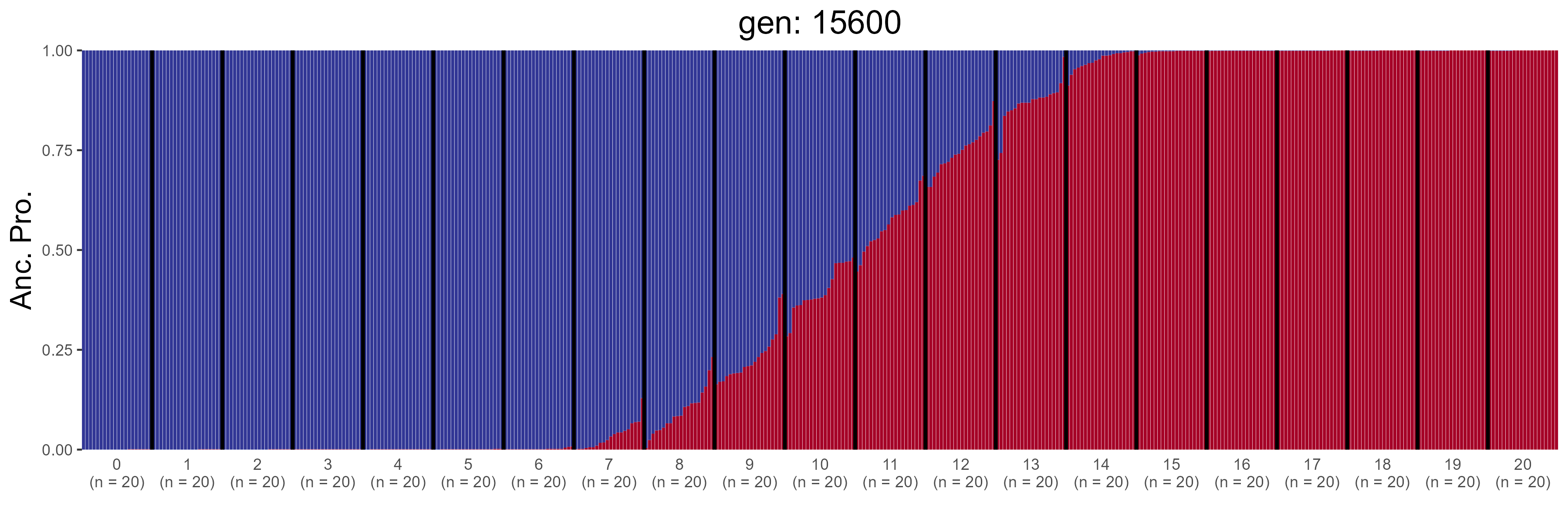

### IBD.15800.str.png

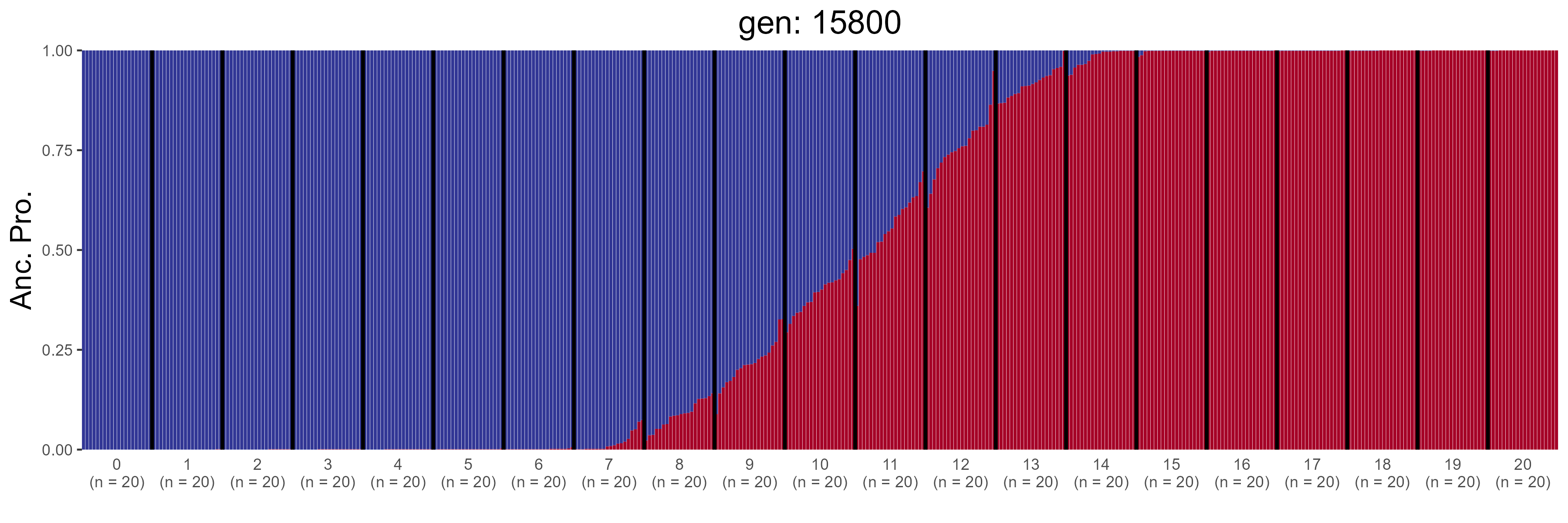

### IBD.16000.str.png

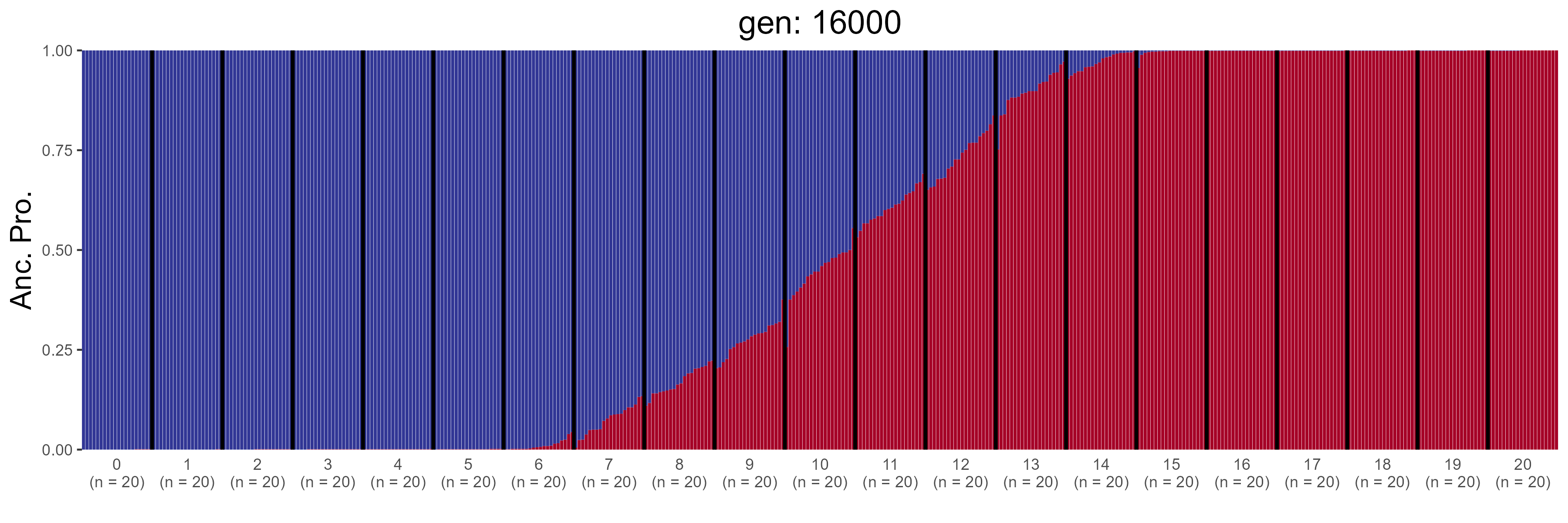

### IBD.16200.str.png

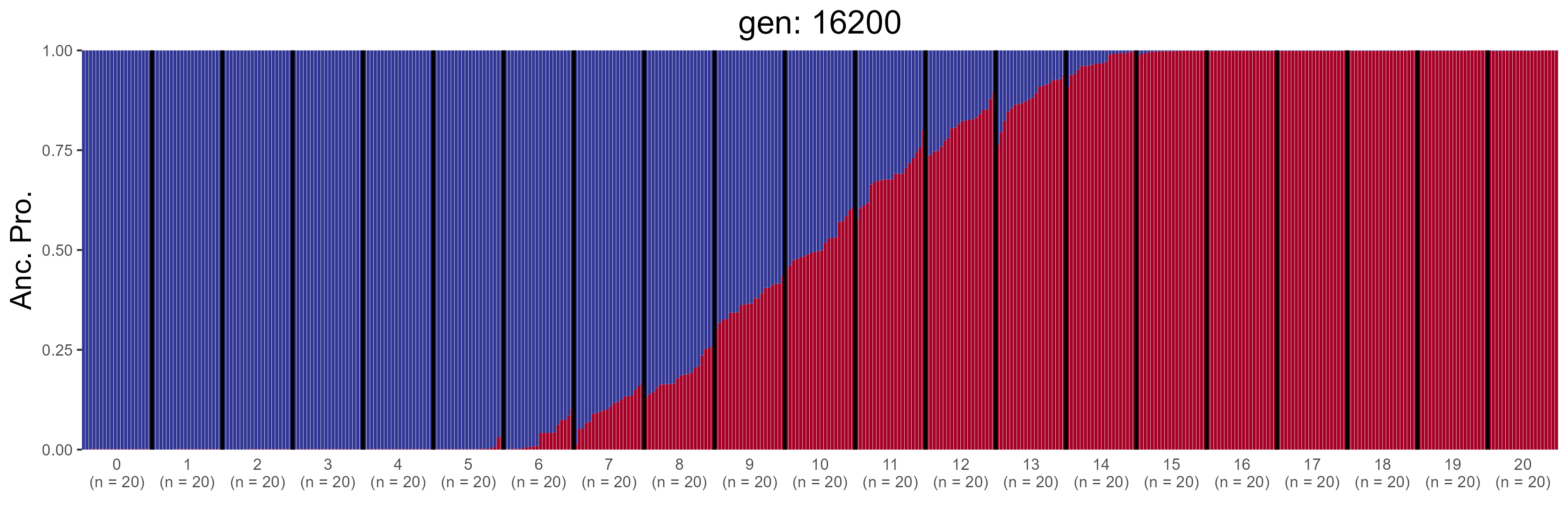

### IBD.16400.str.png

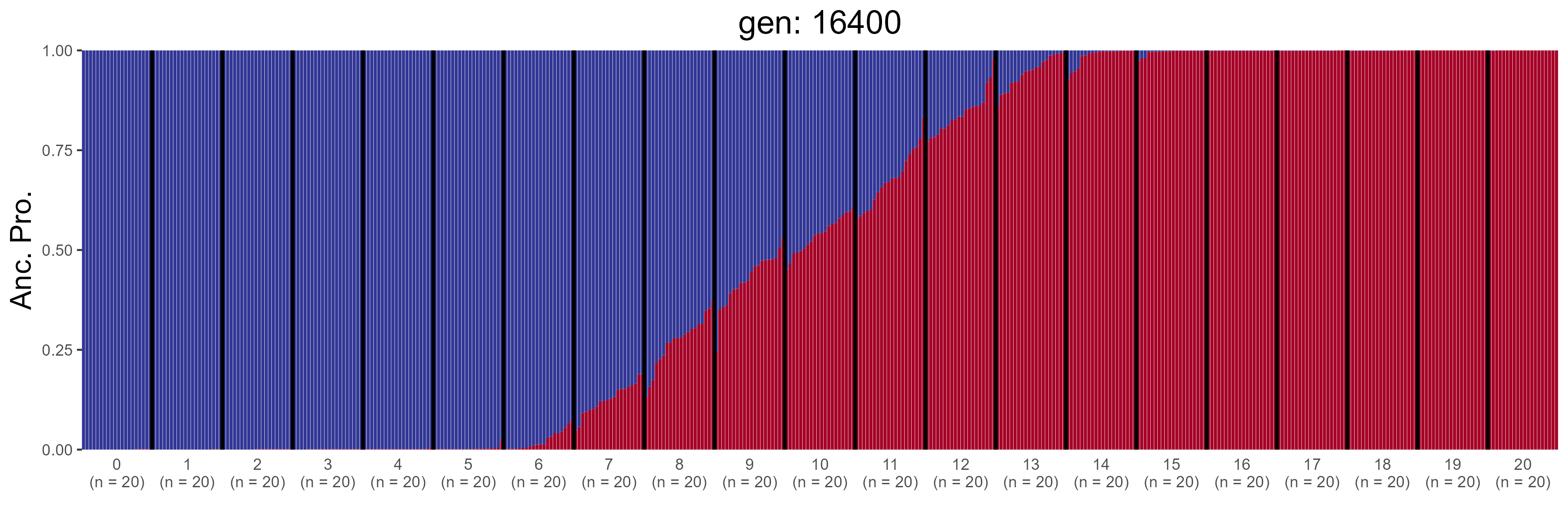

### IBD.16600.str.png

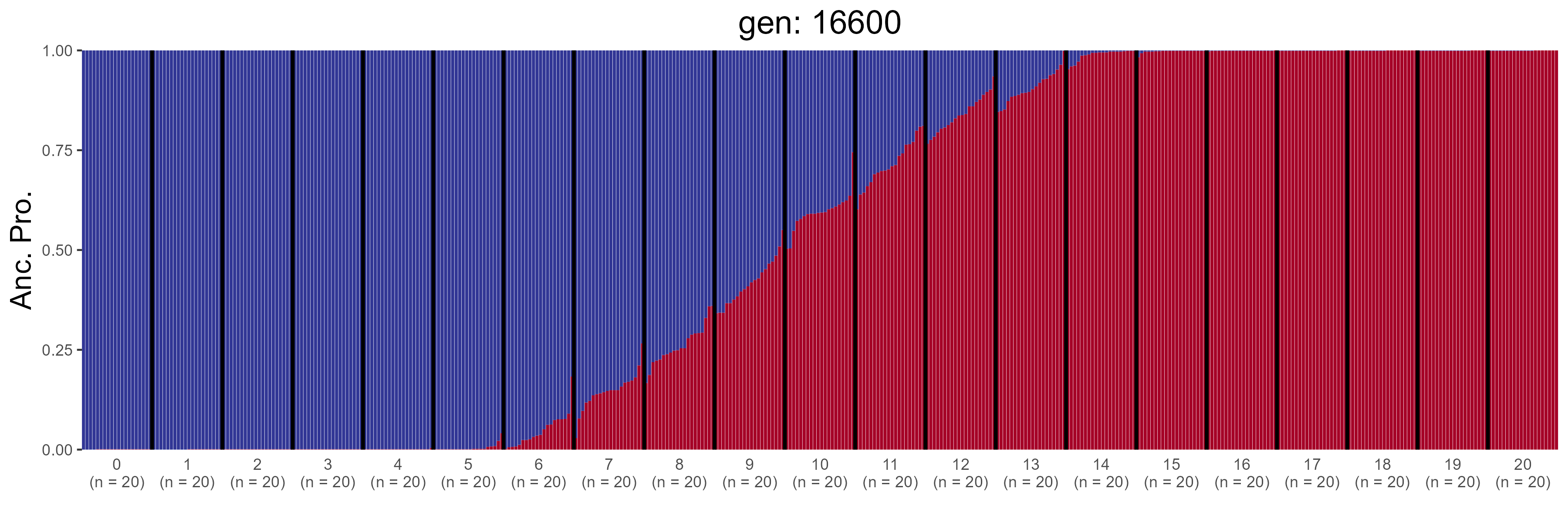

### IBD.16800.str.png

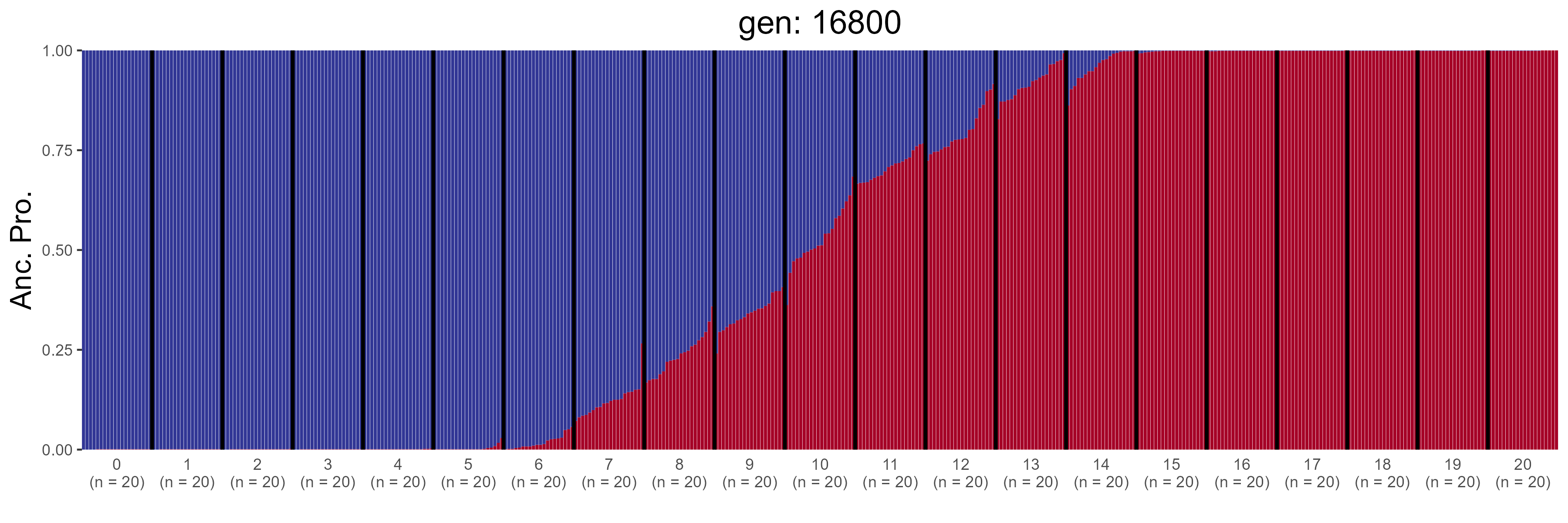

### IBD.17000.str.png

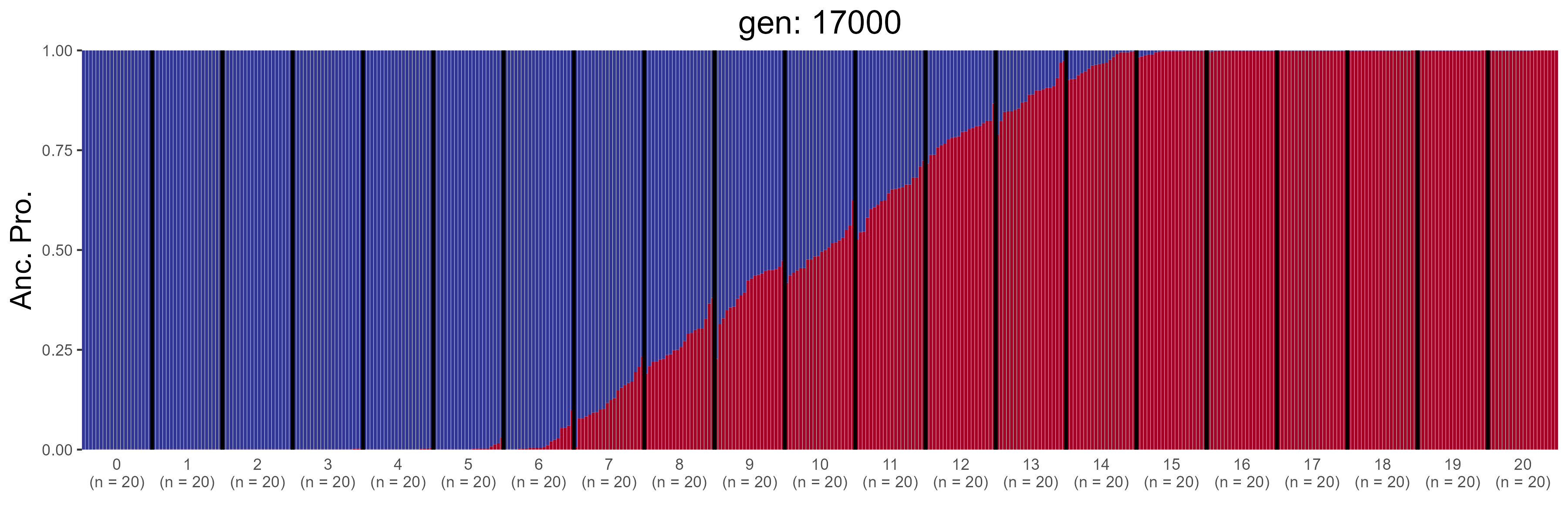

### IBD.17200.str.png

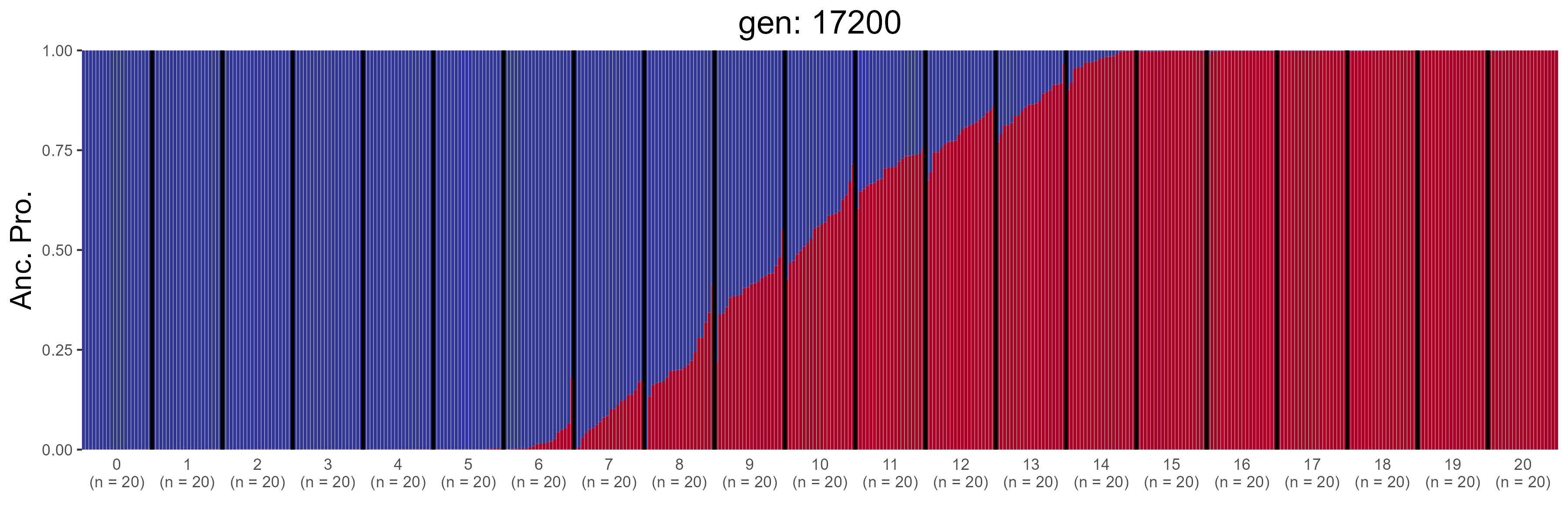

### IBD.17400.str.png

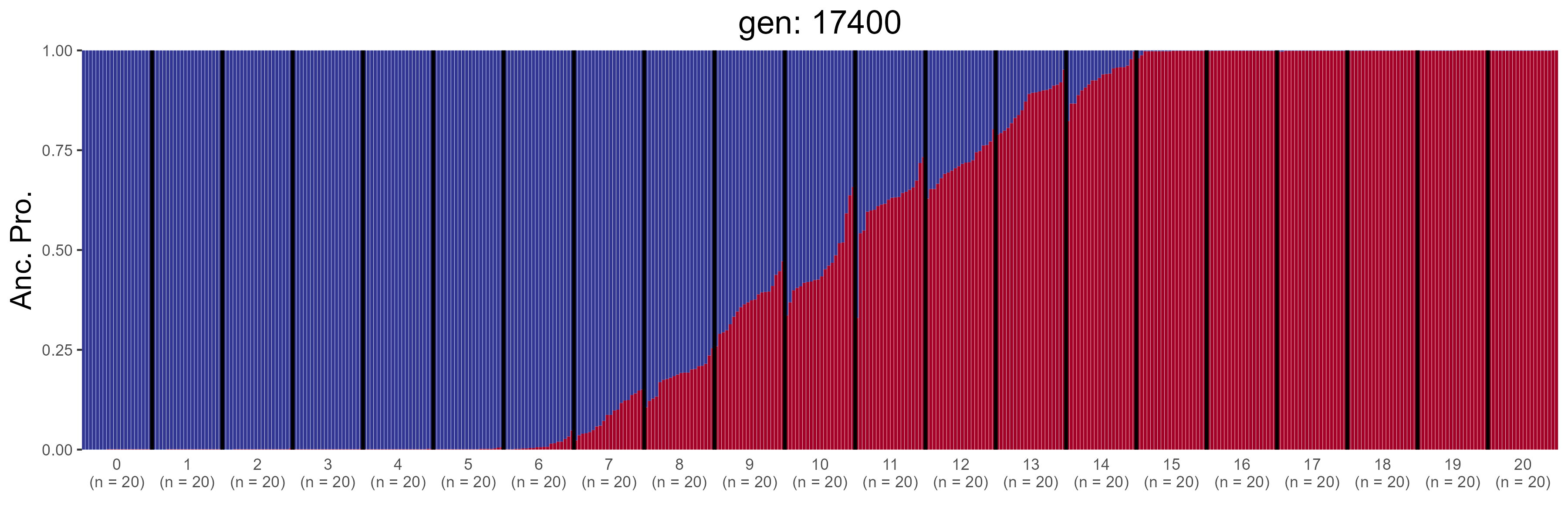

### IBD.17600.str.png

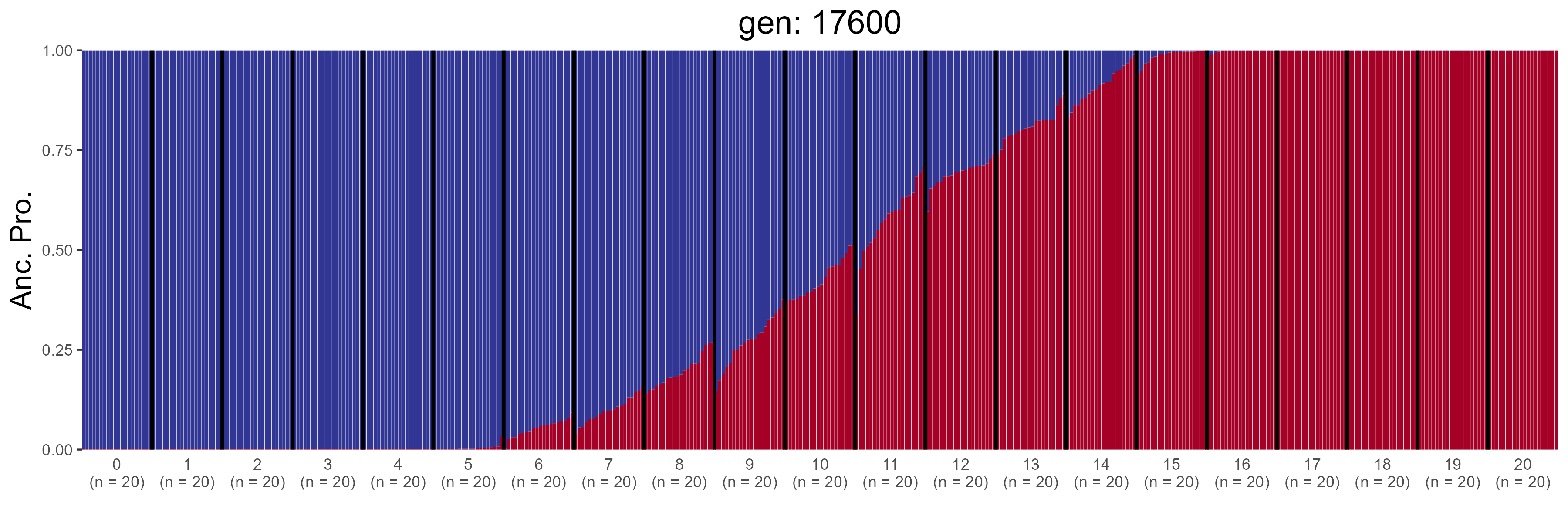

### IBD.17800.str.png

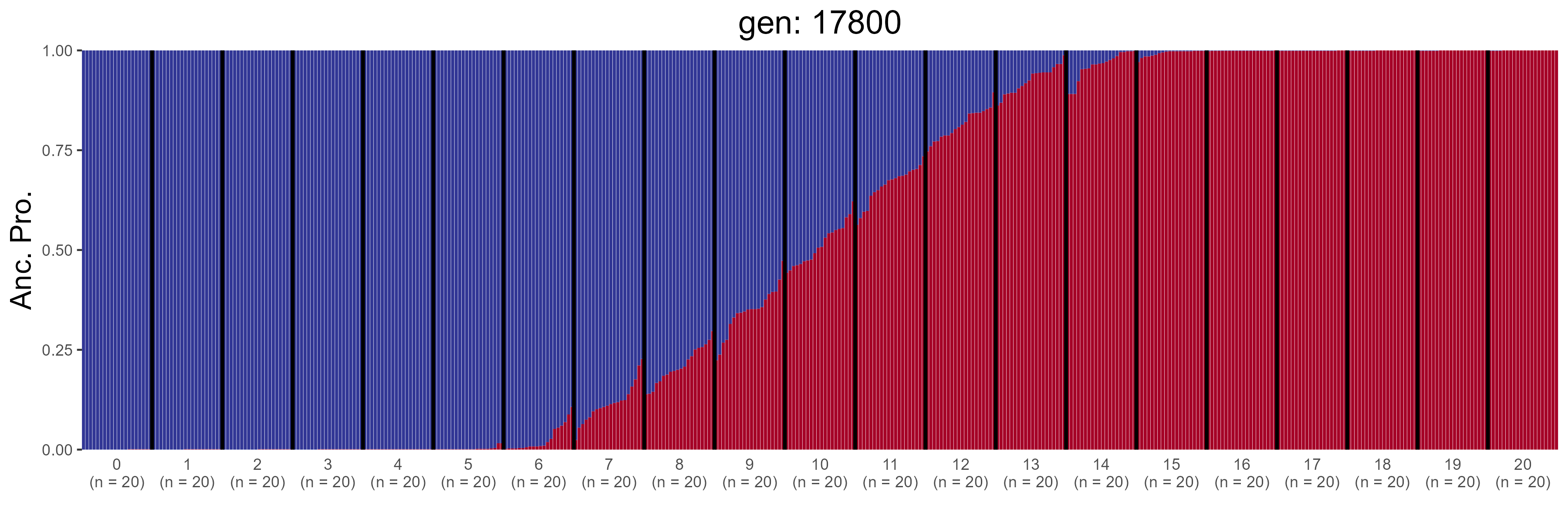

### IBD.18000.str.png

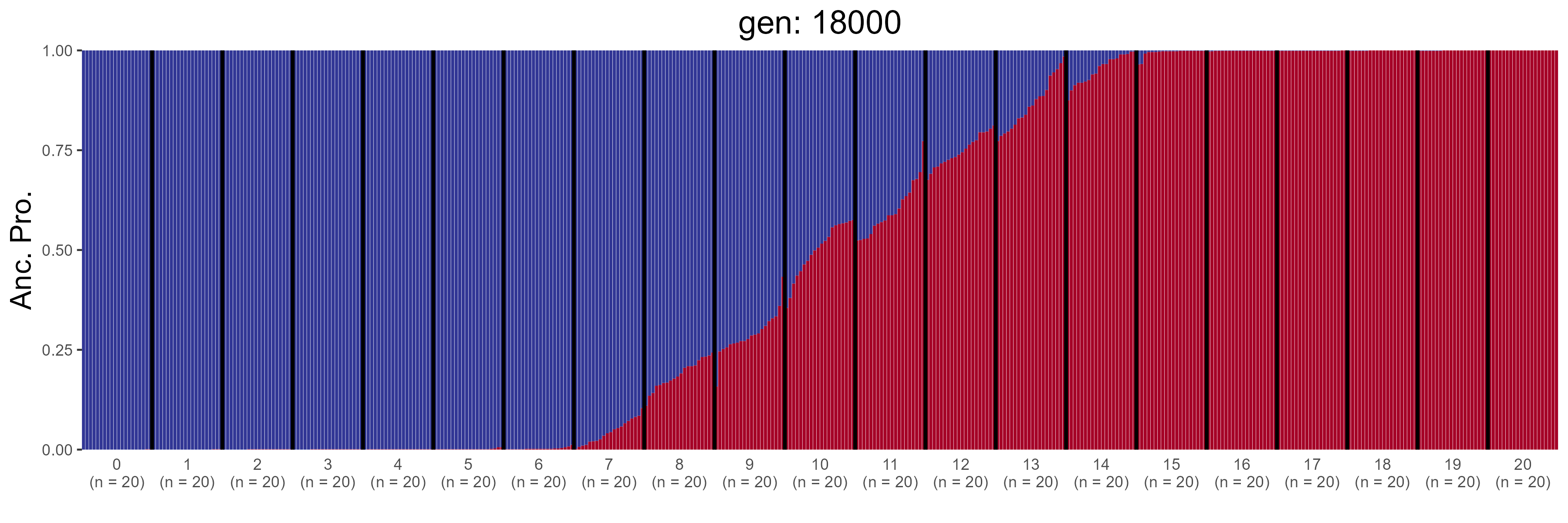

### IBD.18200.str.png

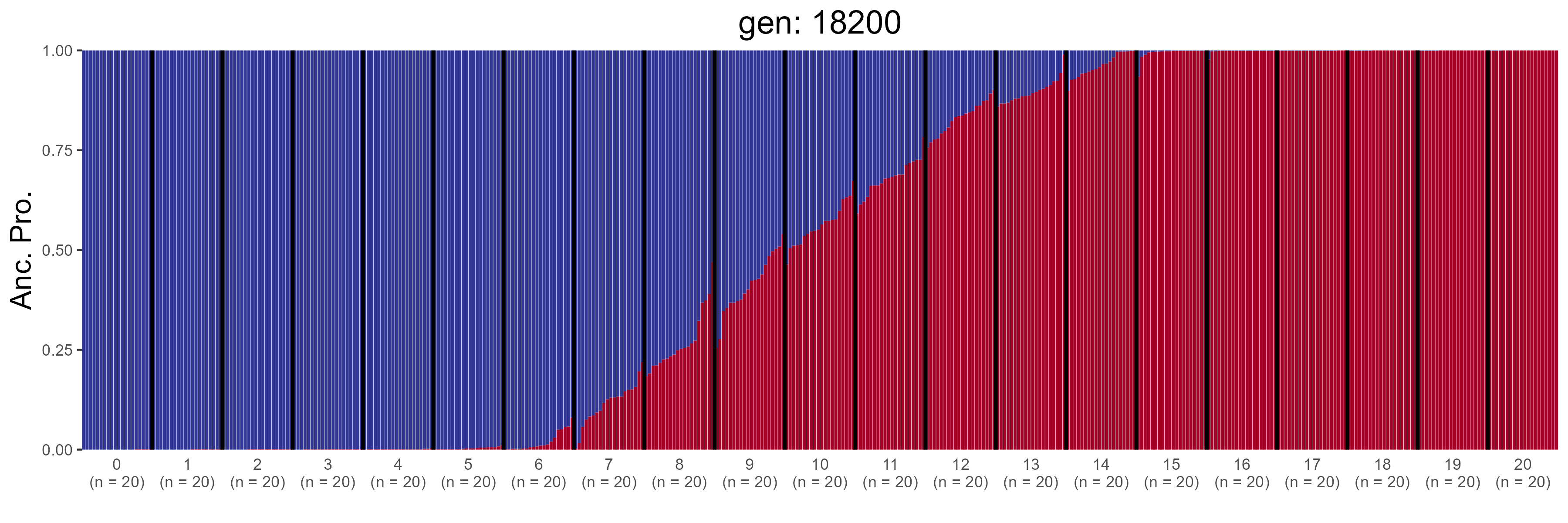

### IBD.18400.str.png

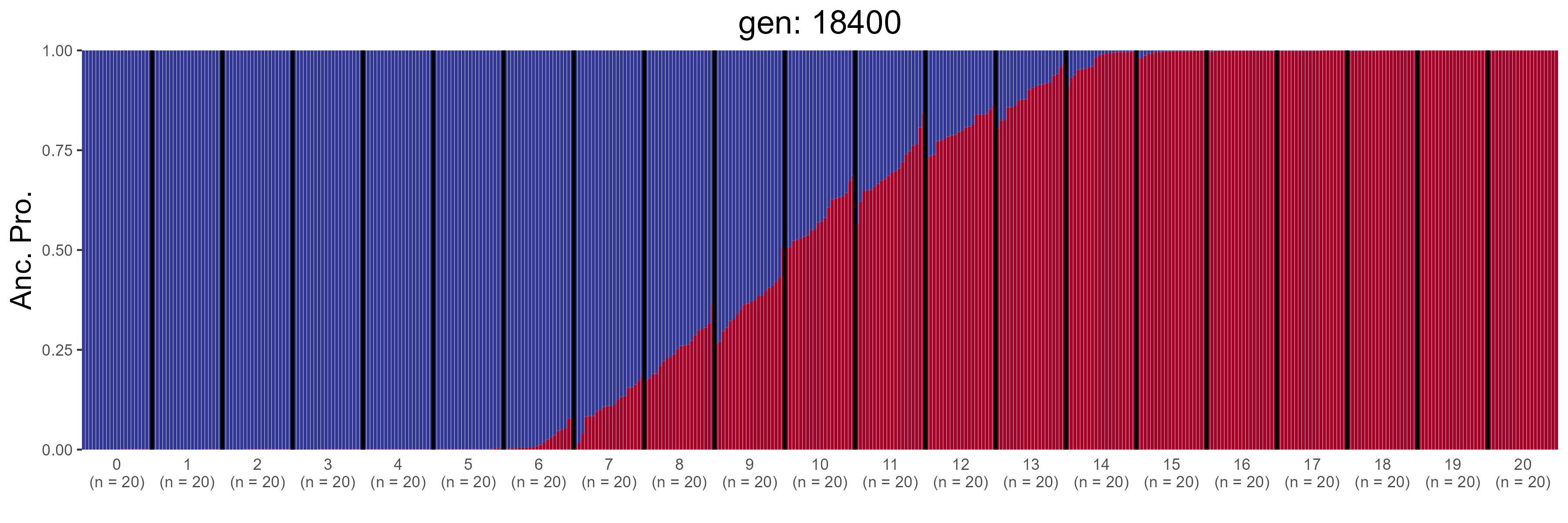

### IBD.18600.str.png

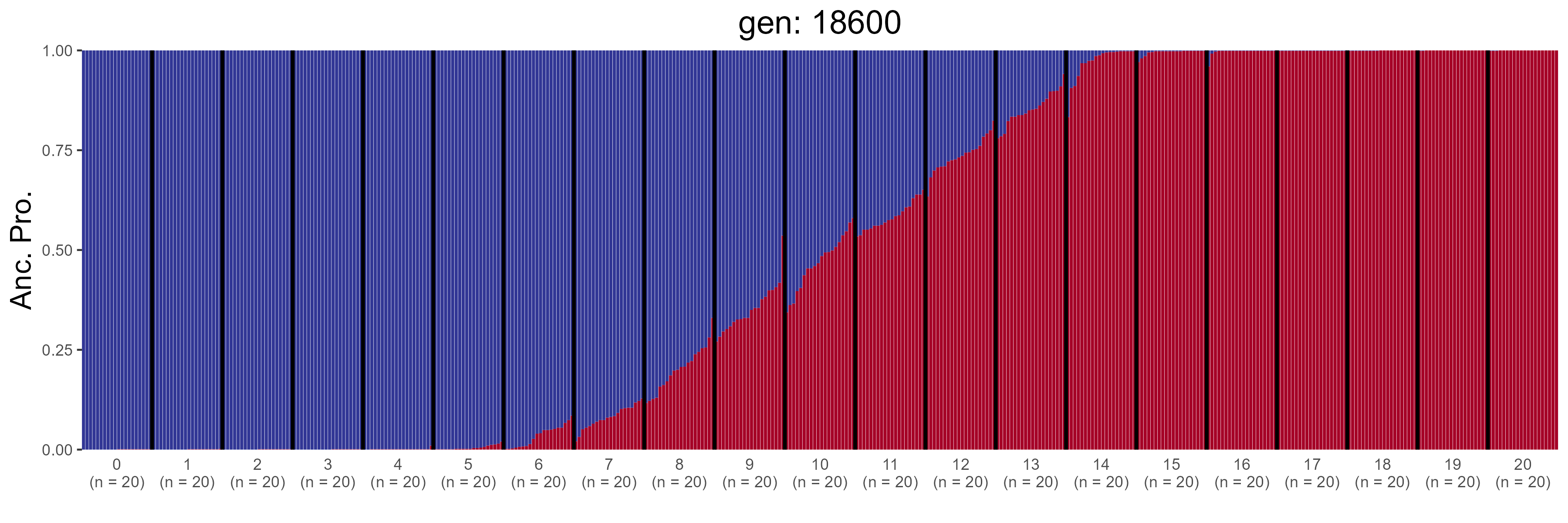

### IBD.18800.str.png

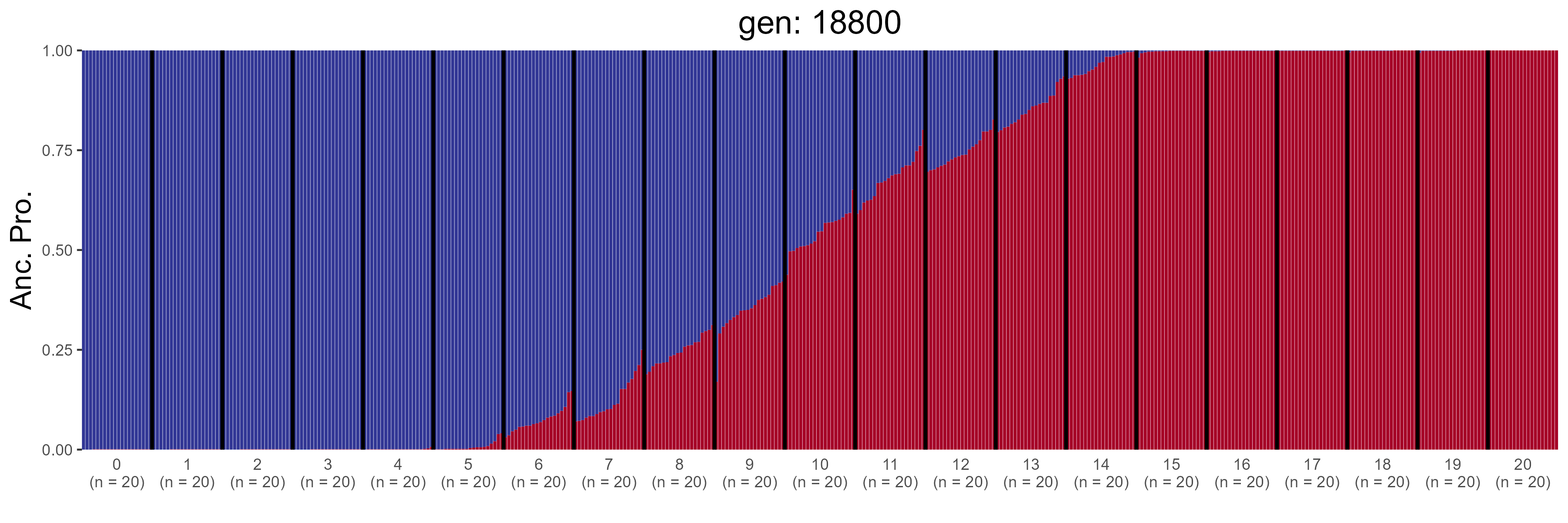

### IBD.19000.str.png

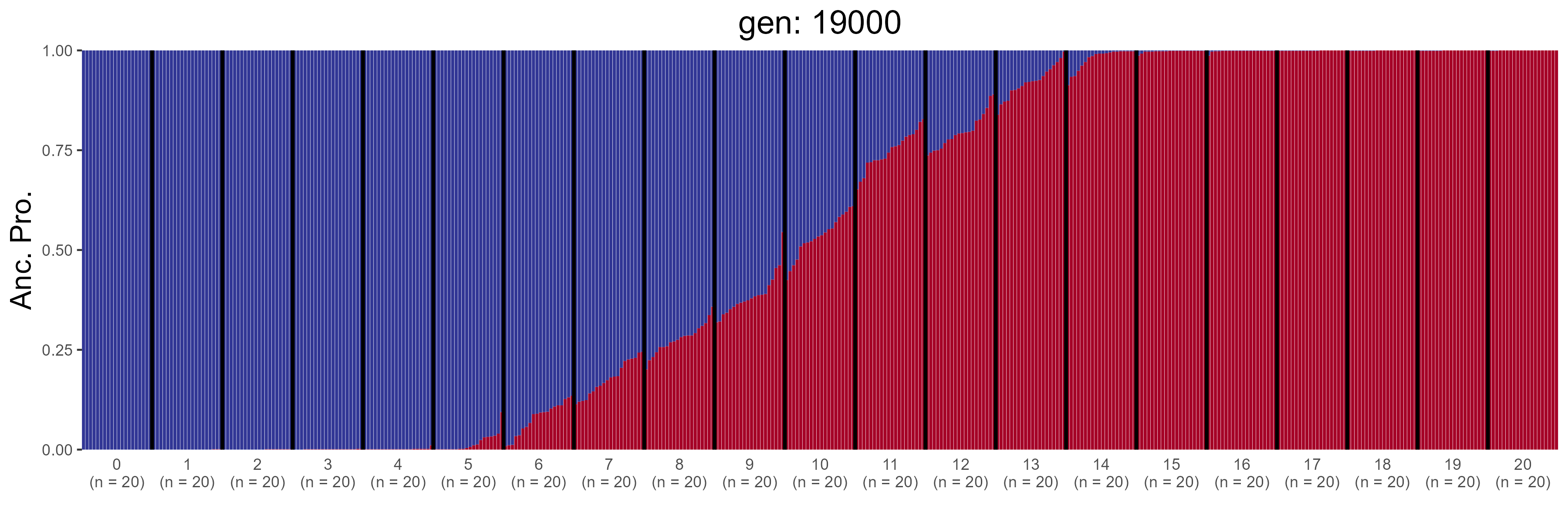

### ND.0.str.png

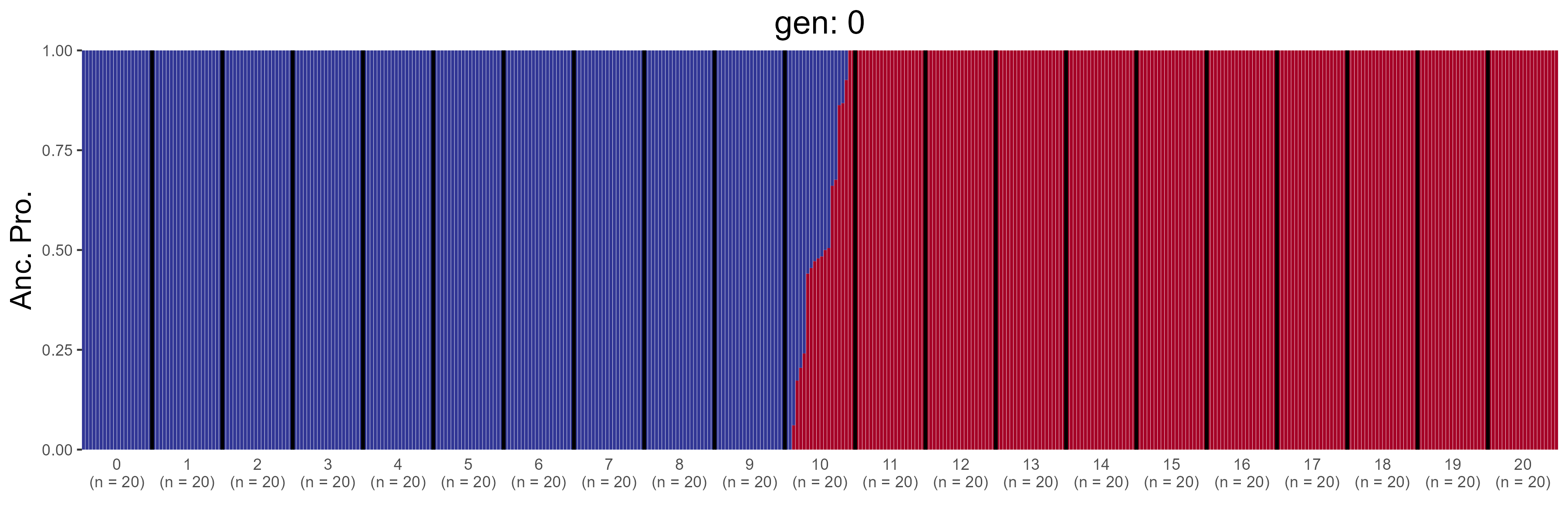

### ND.1000.str.png

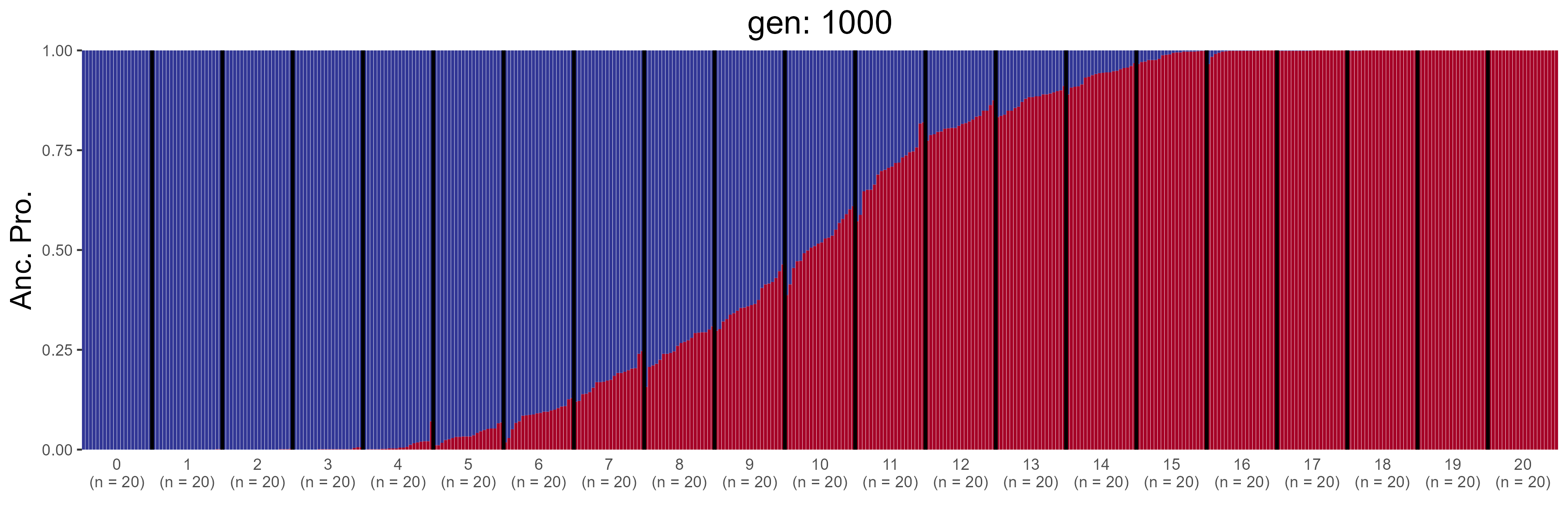

### ND.10000.str.png

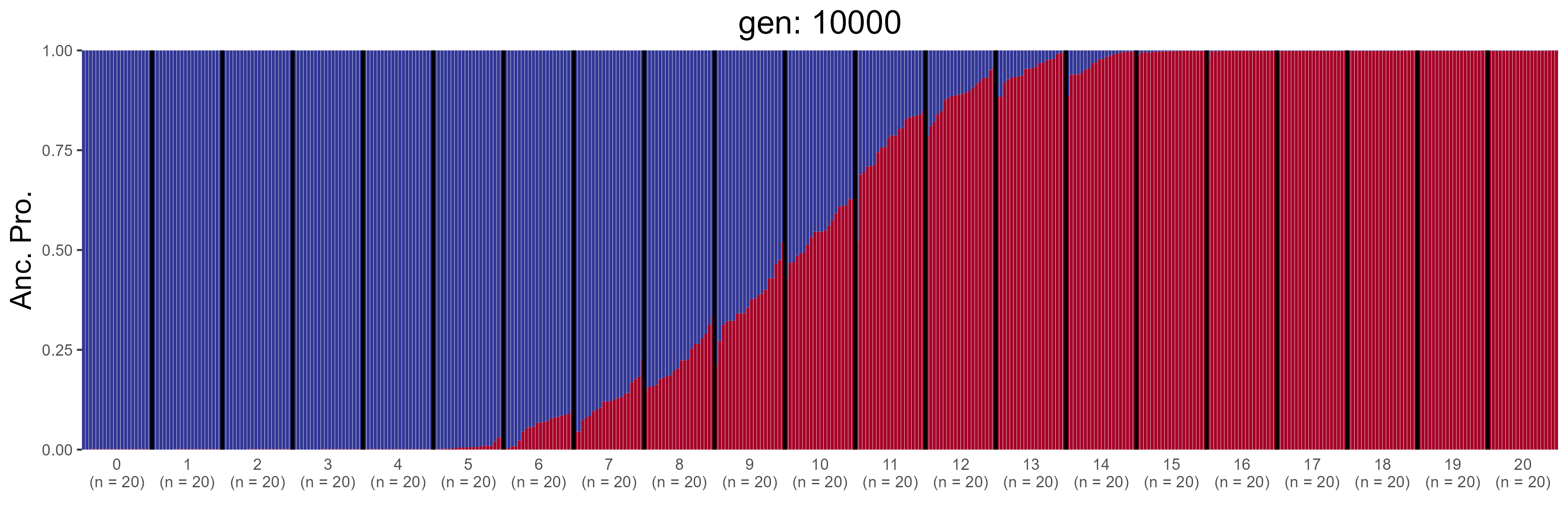

### ND.10200.str.png

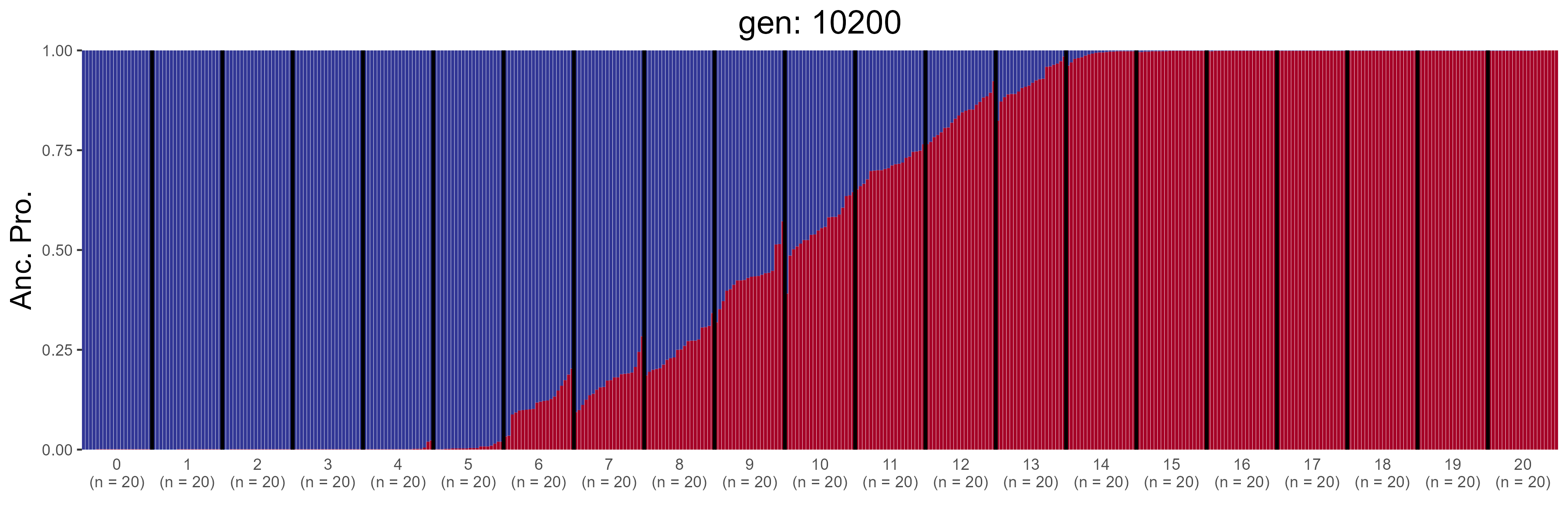

### ND.10400.str.png

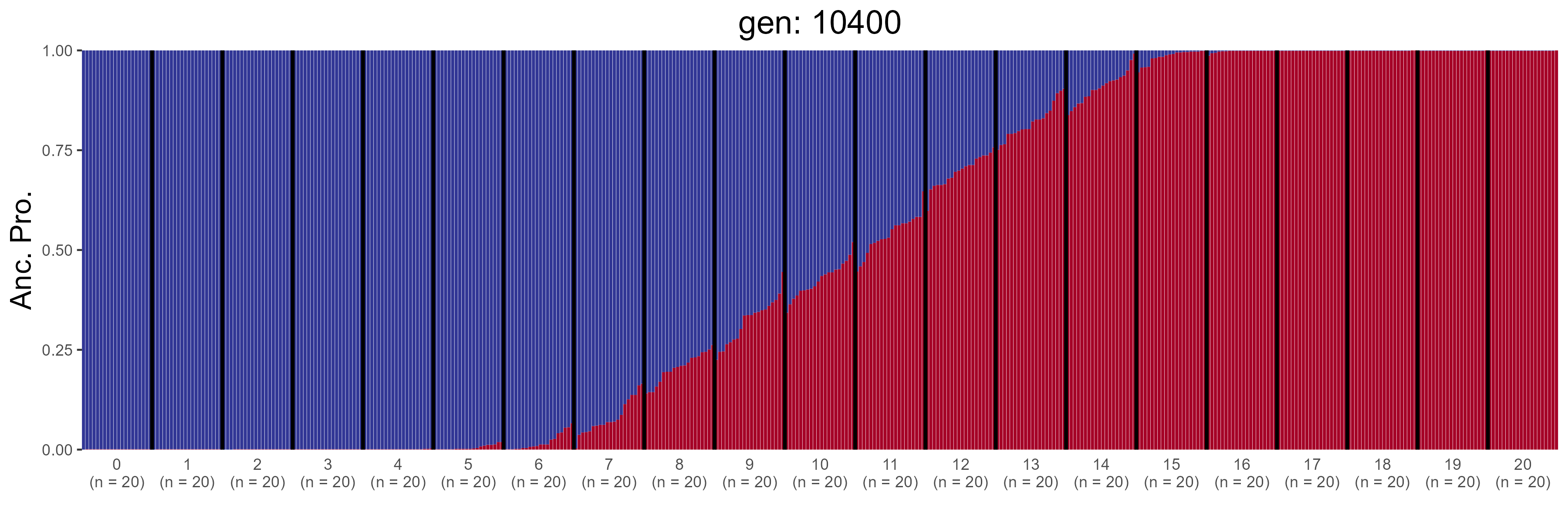

### ND.10600.str.png

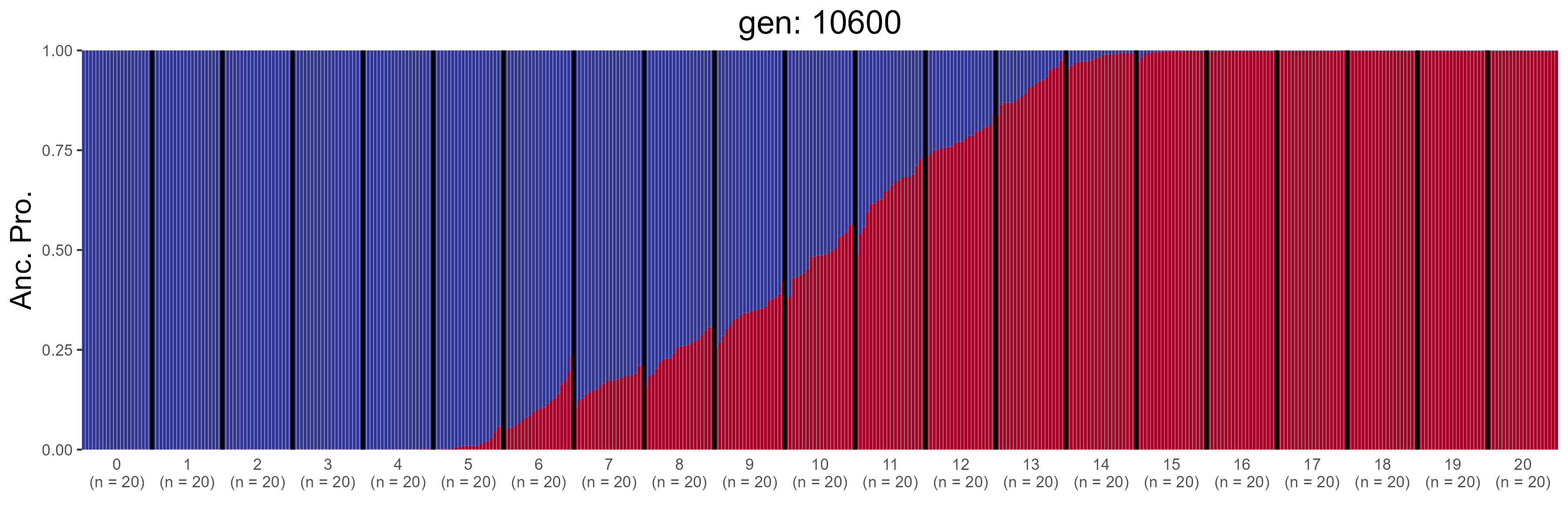

### ND.10800.str.png

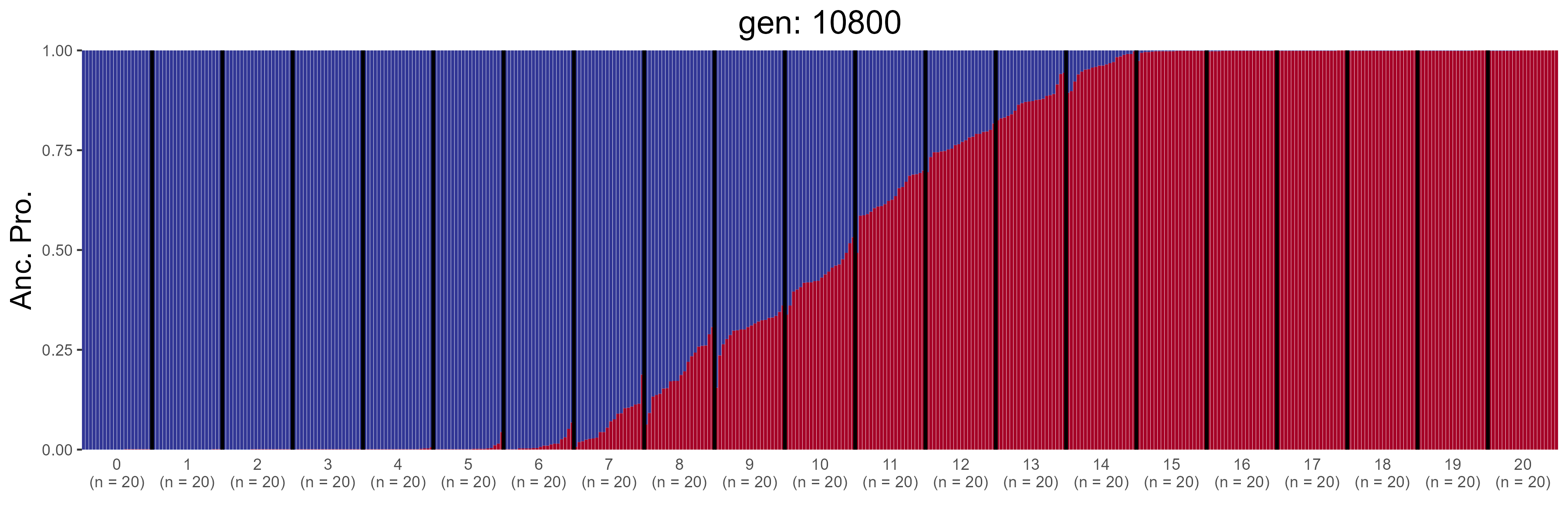

### ND.11000.str.png

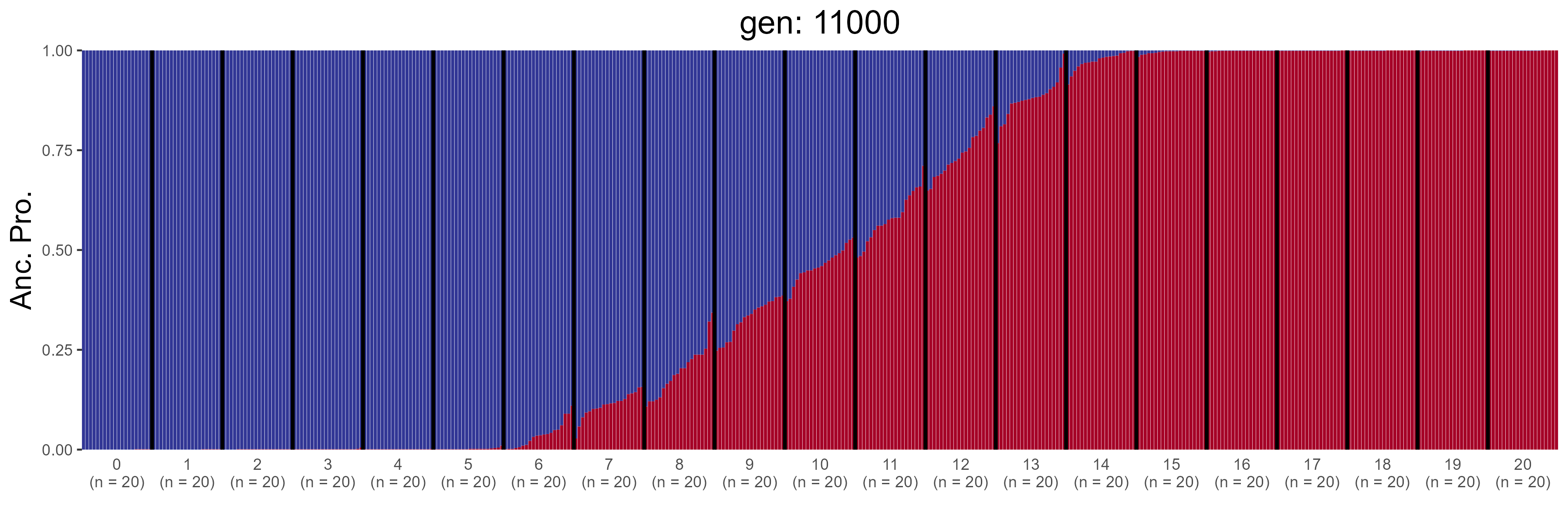
