## Supplementary figures and images for "That’s not a Hybrid: How to Distinguish Patterns of Admixture and Isolation-by-Distance"

### GBC.0.str.png

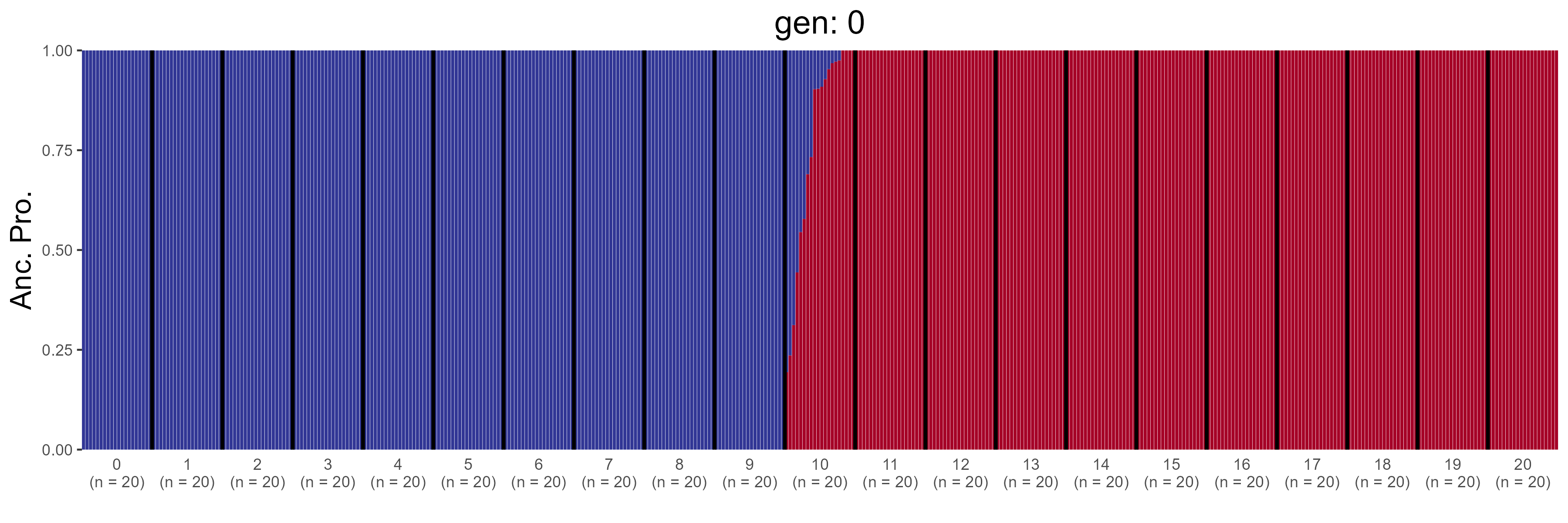

### GBC.200.str.png

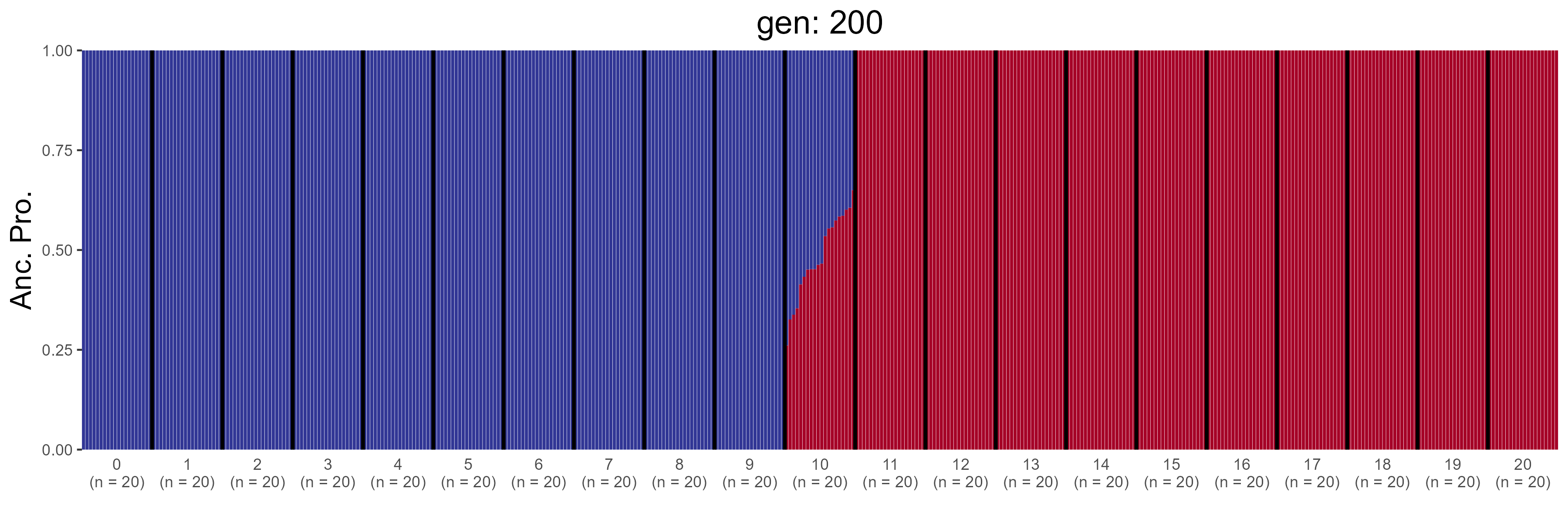

### GBC.400.str.png

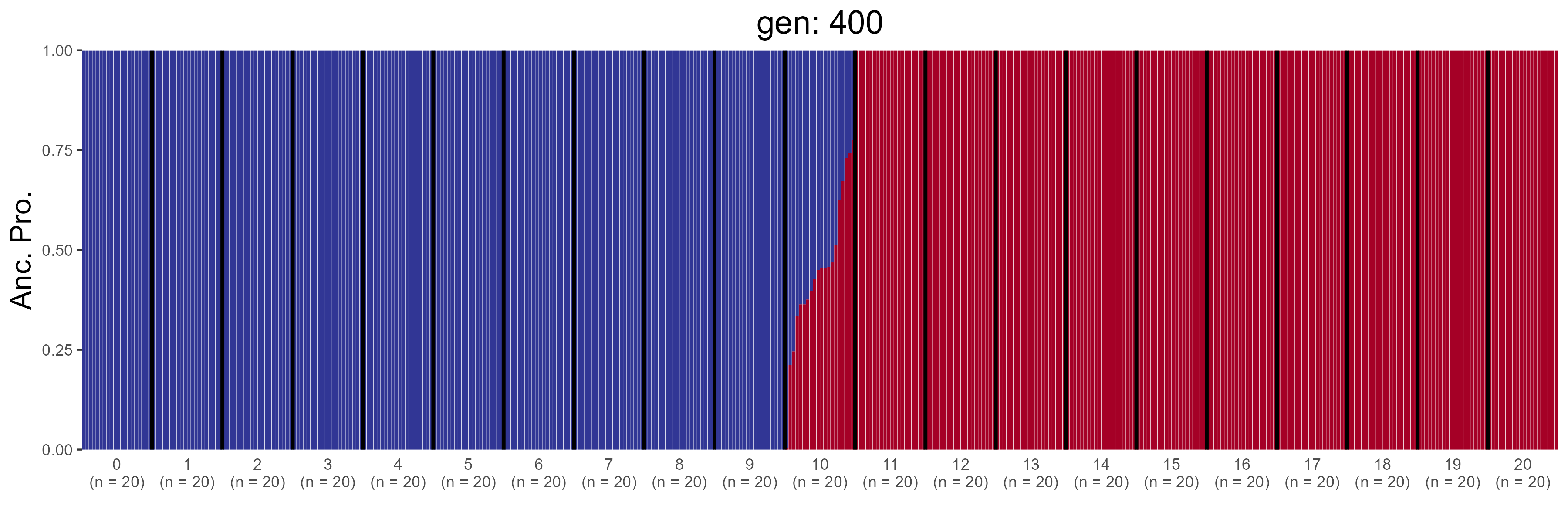

### GBC.600.str.png

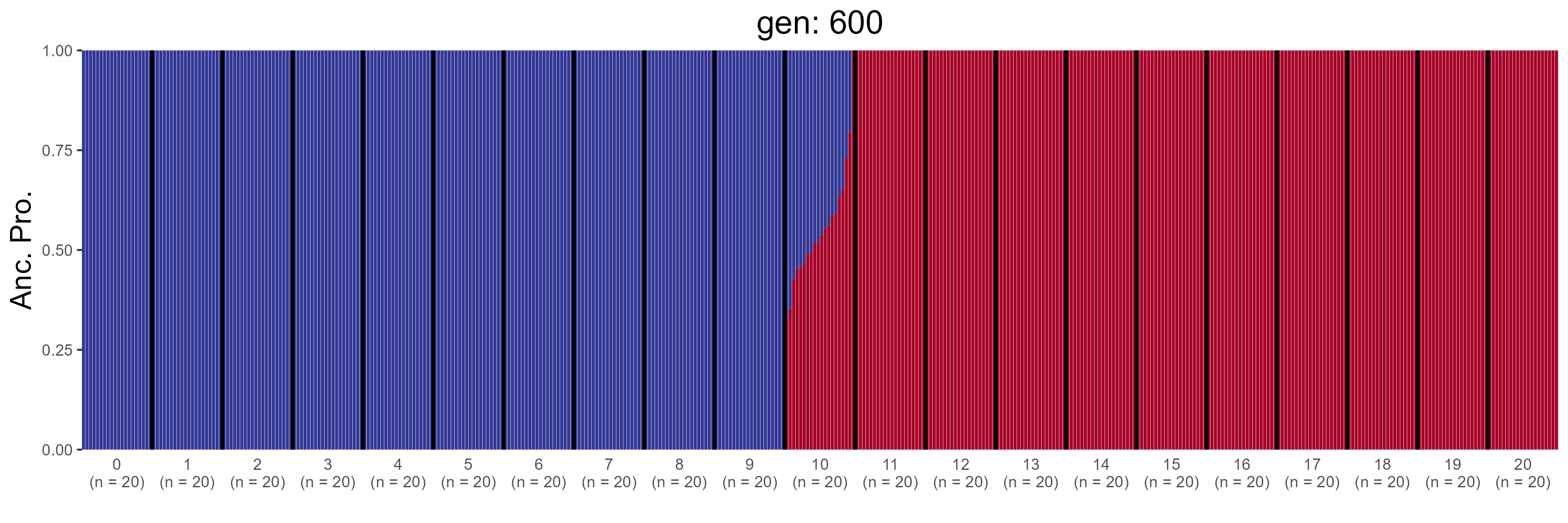

### GBC.800.str.png

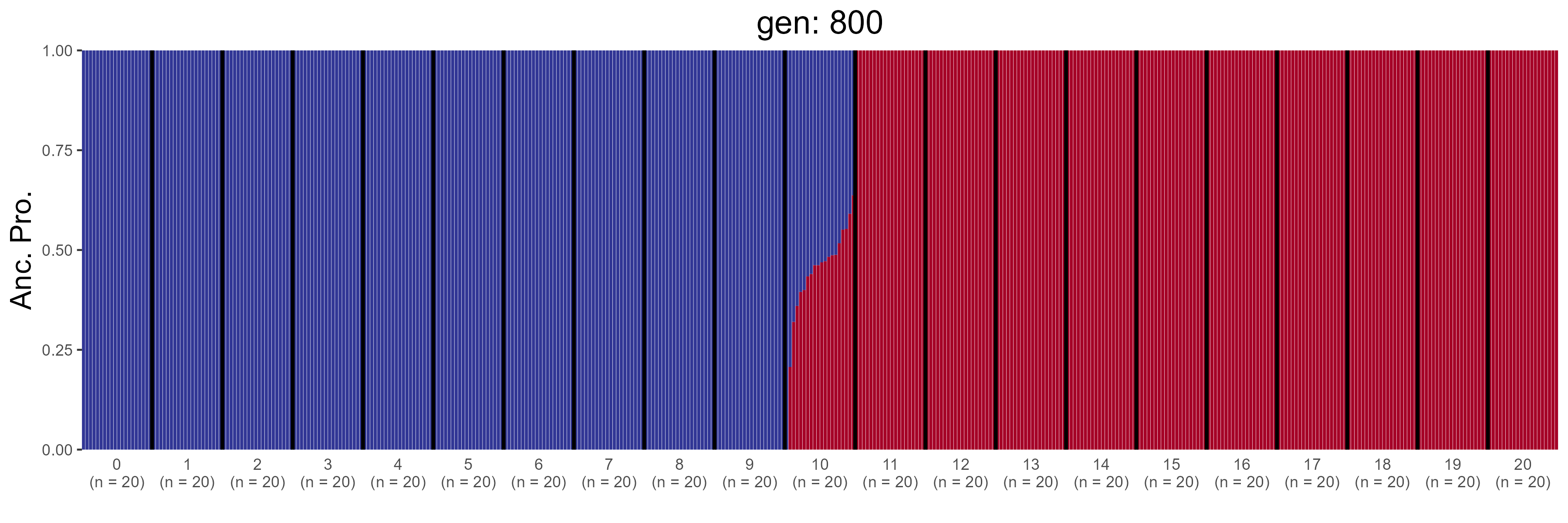

### GBC.1000.str.png

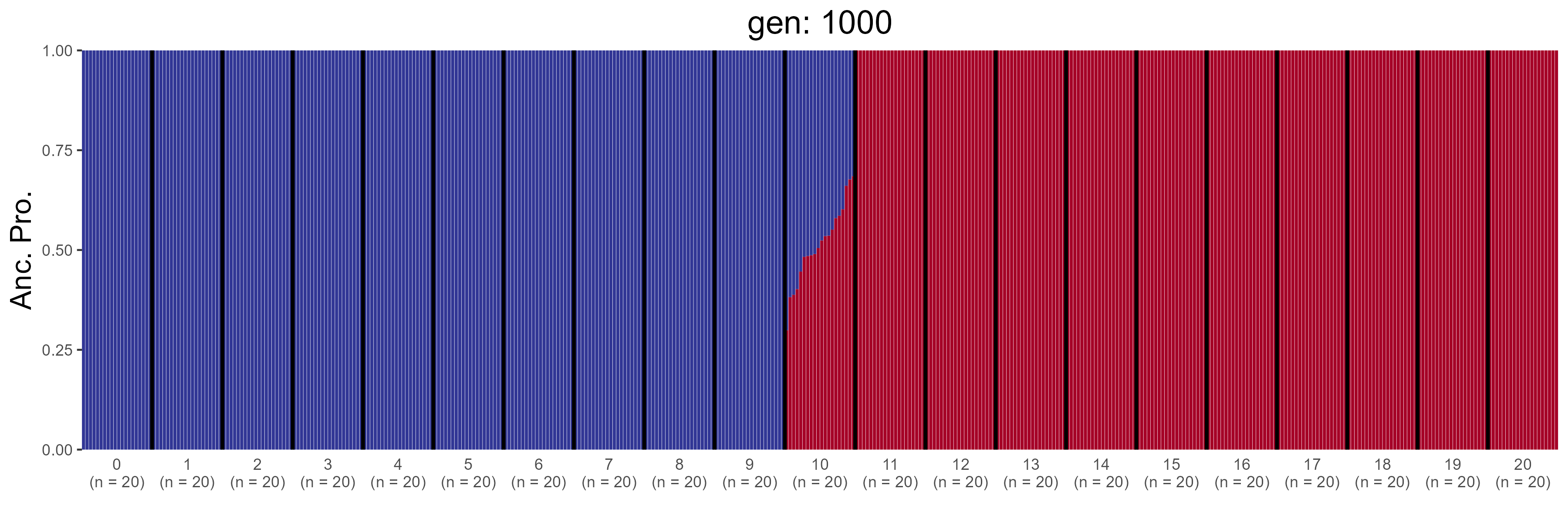

### GBC.1200.str.png

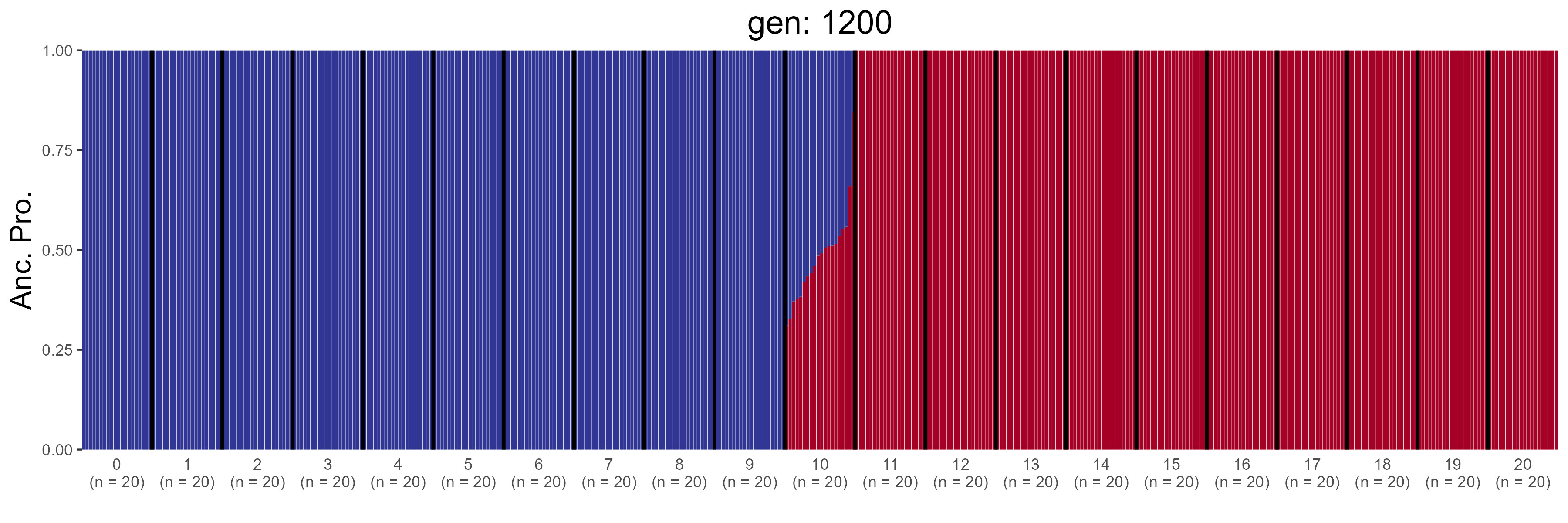

### GBC.1400.str.png

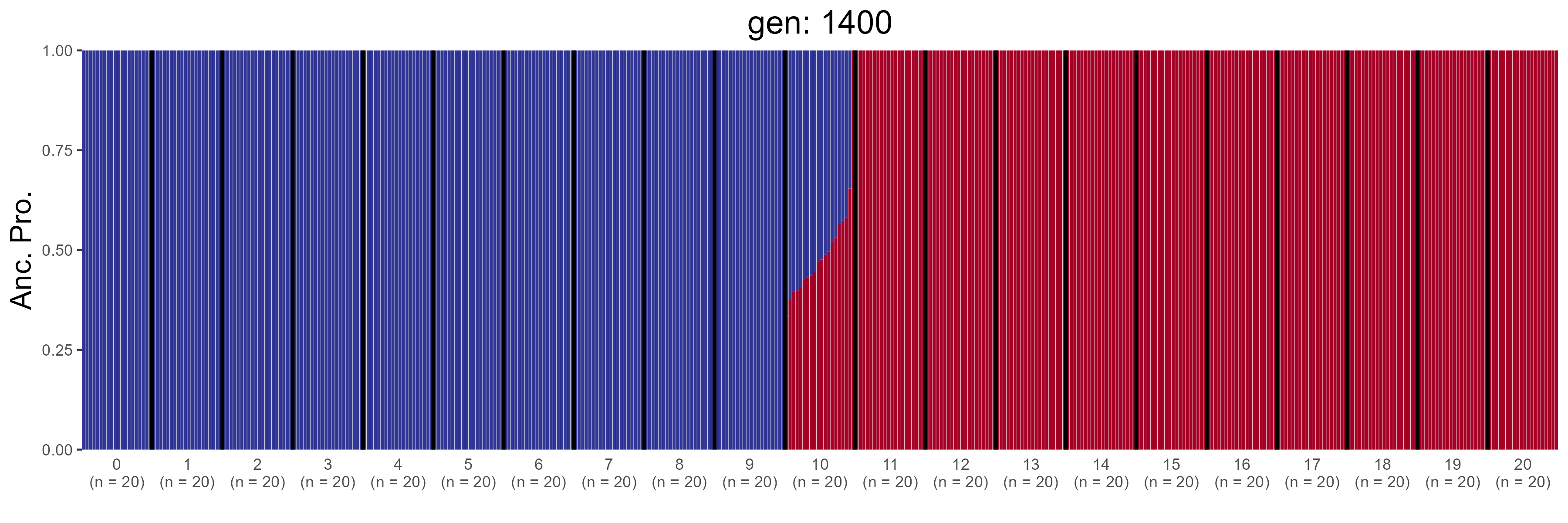

### GBC.1600.str.png

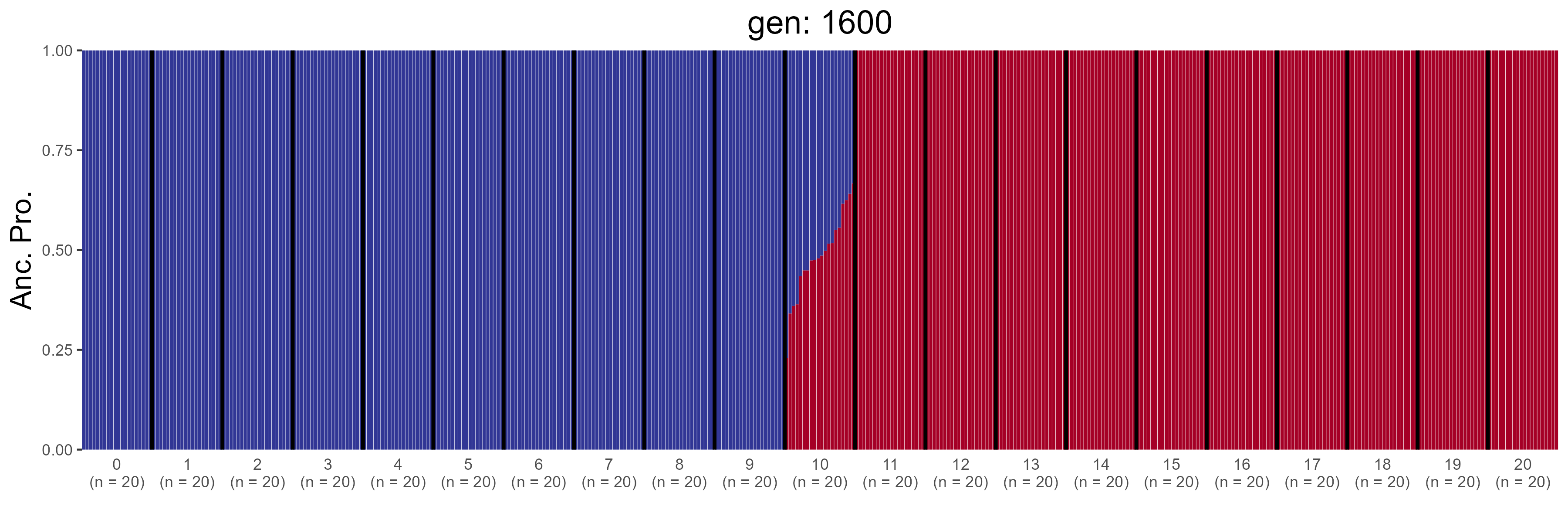

### GBC.1800.str.png

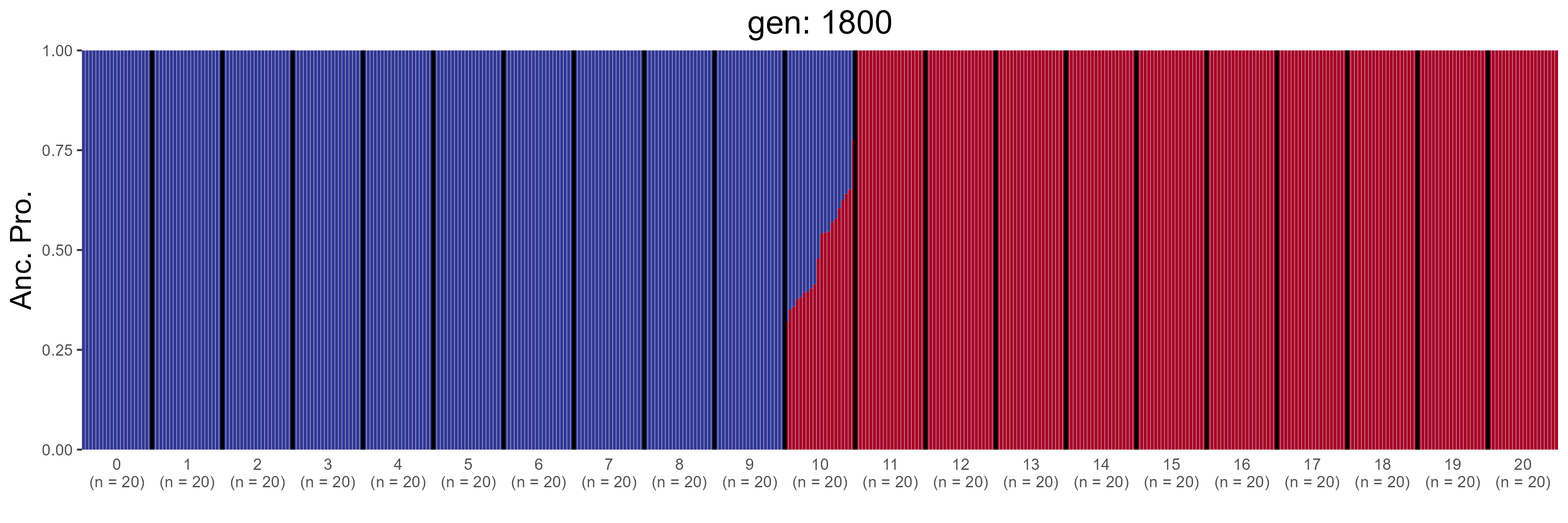

### GBC.2000.str.png

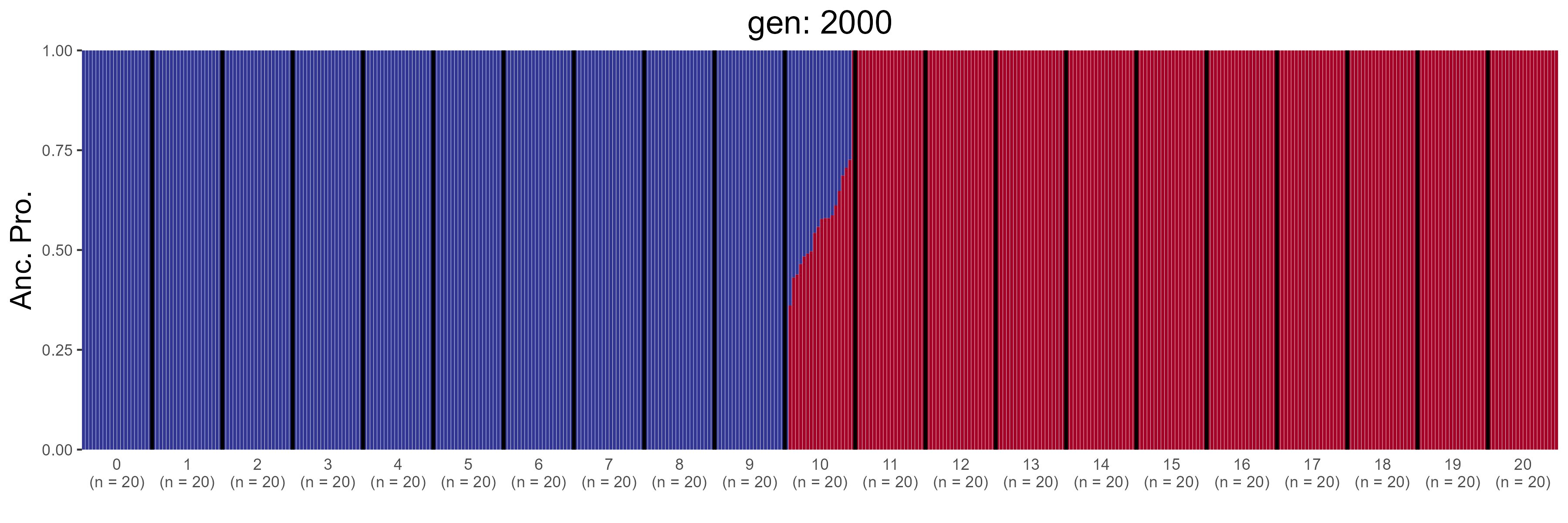

### GBC.2200.str.png

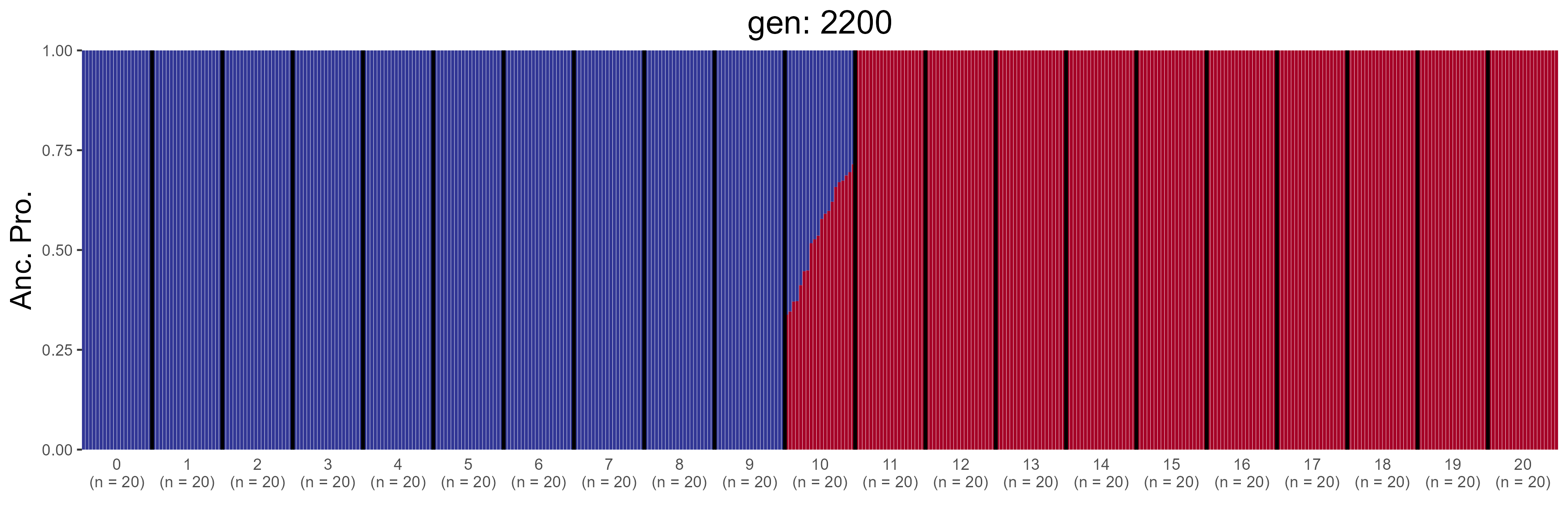

### GBC.2400.str.png

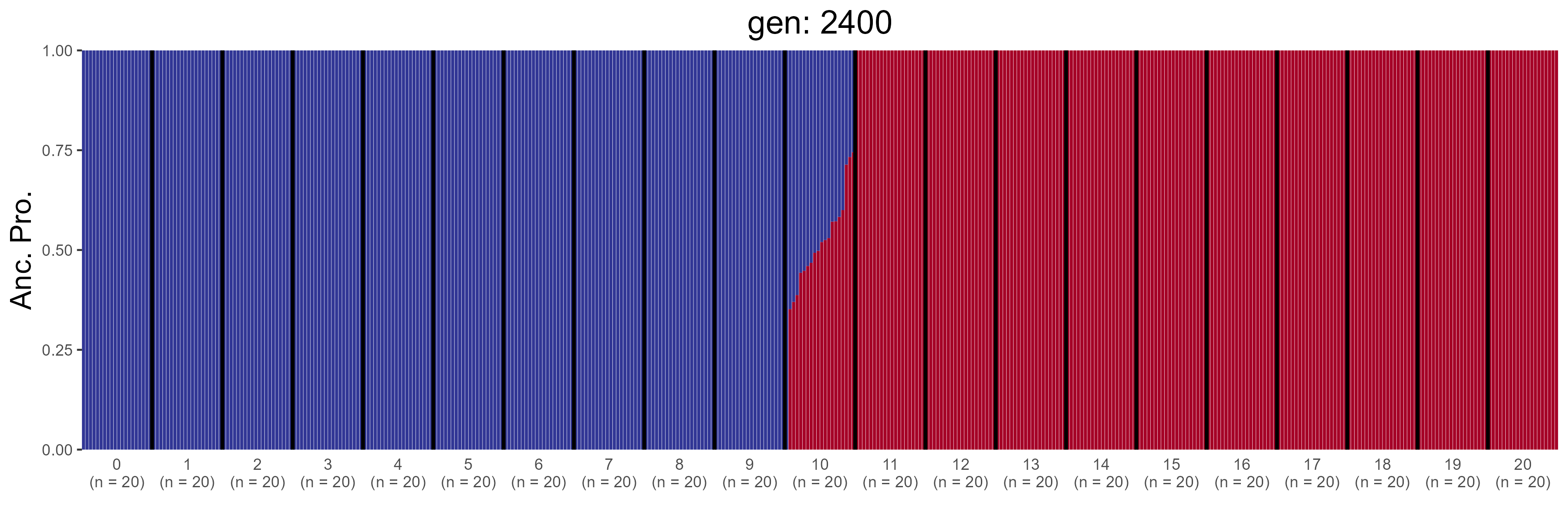

### GBC.2600.str.png

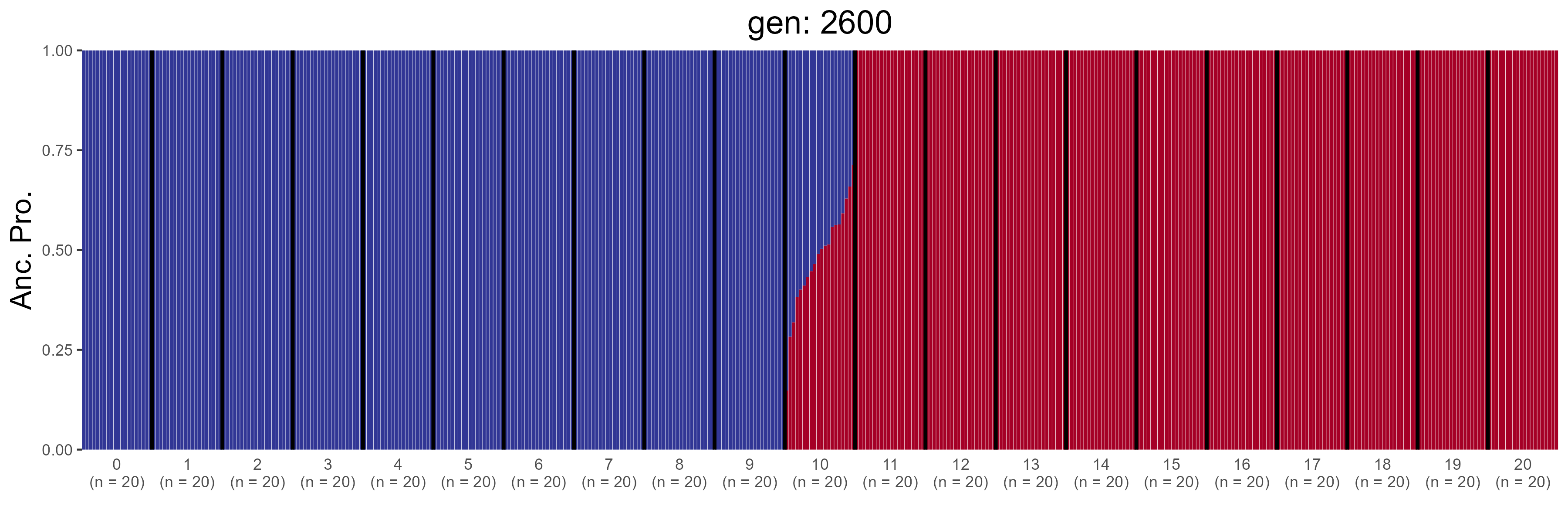

### GBC.2800.str.png

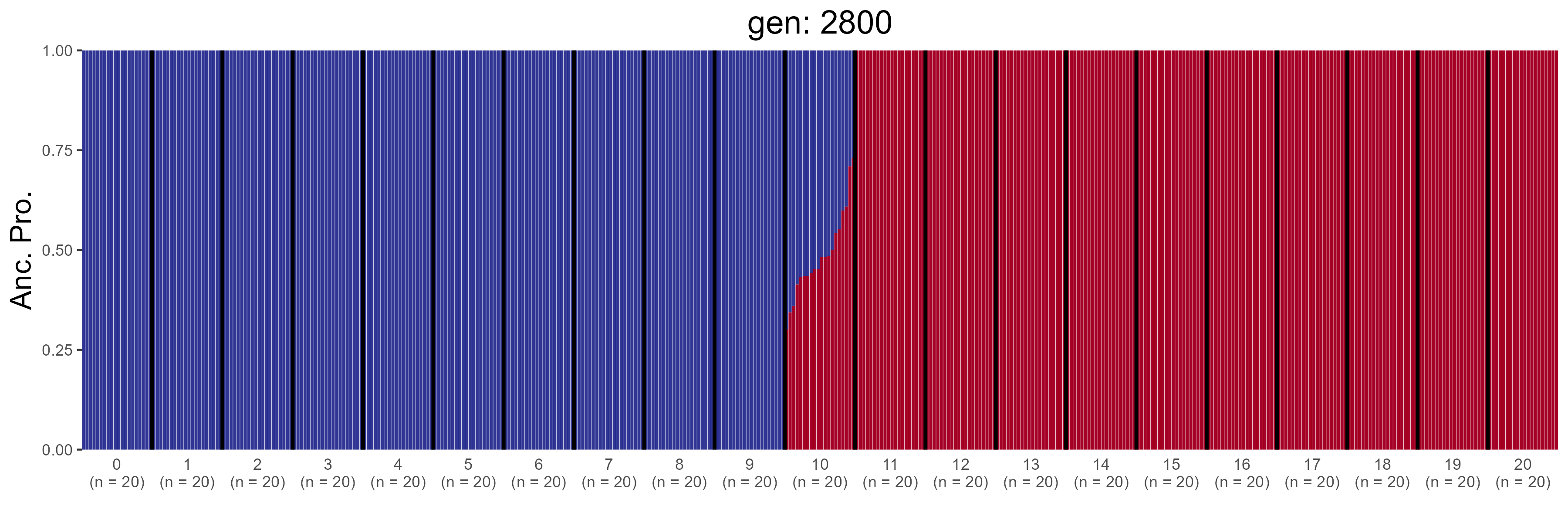

### GBC.3000.str.png

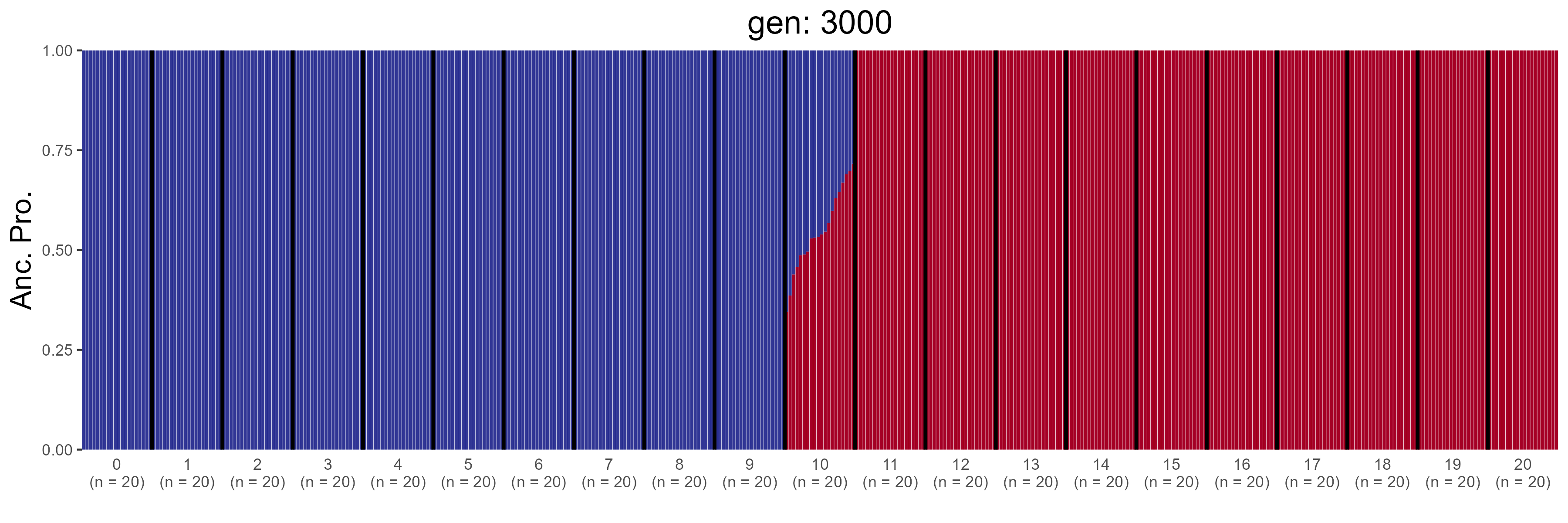

### GBC.3200.str.png

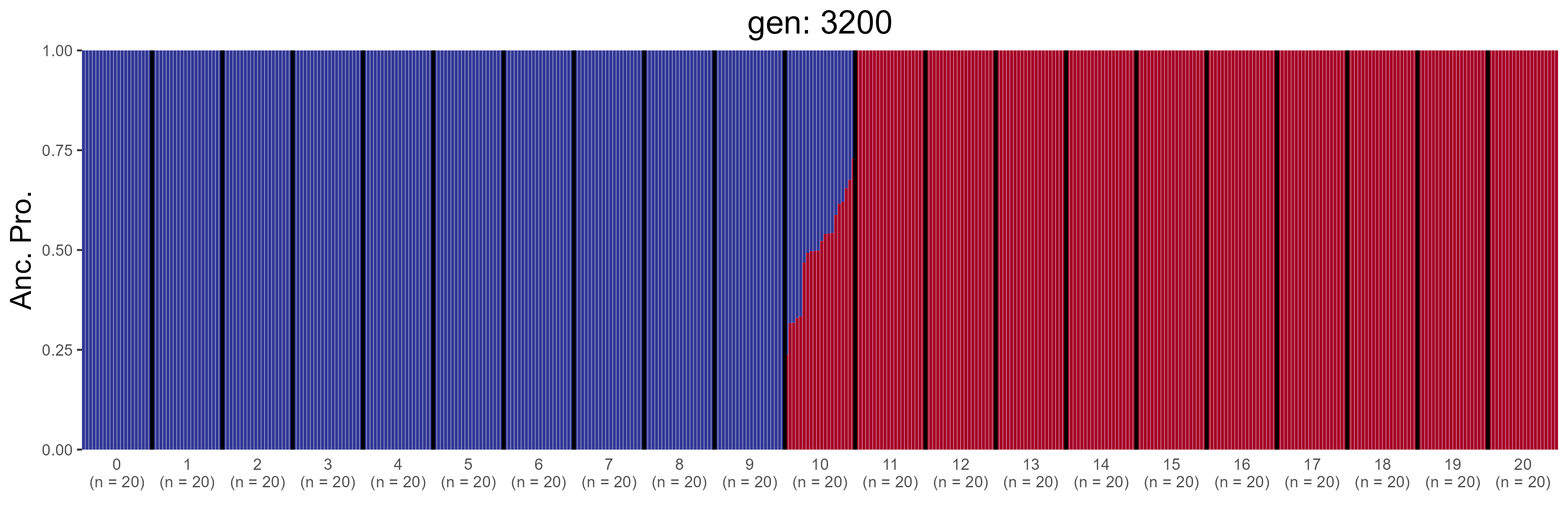

### GBC.3400.str.png

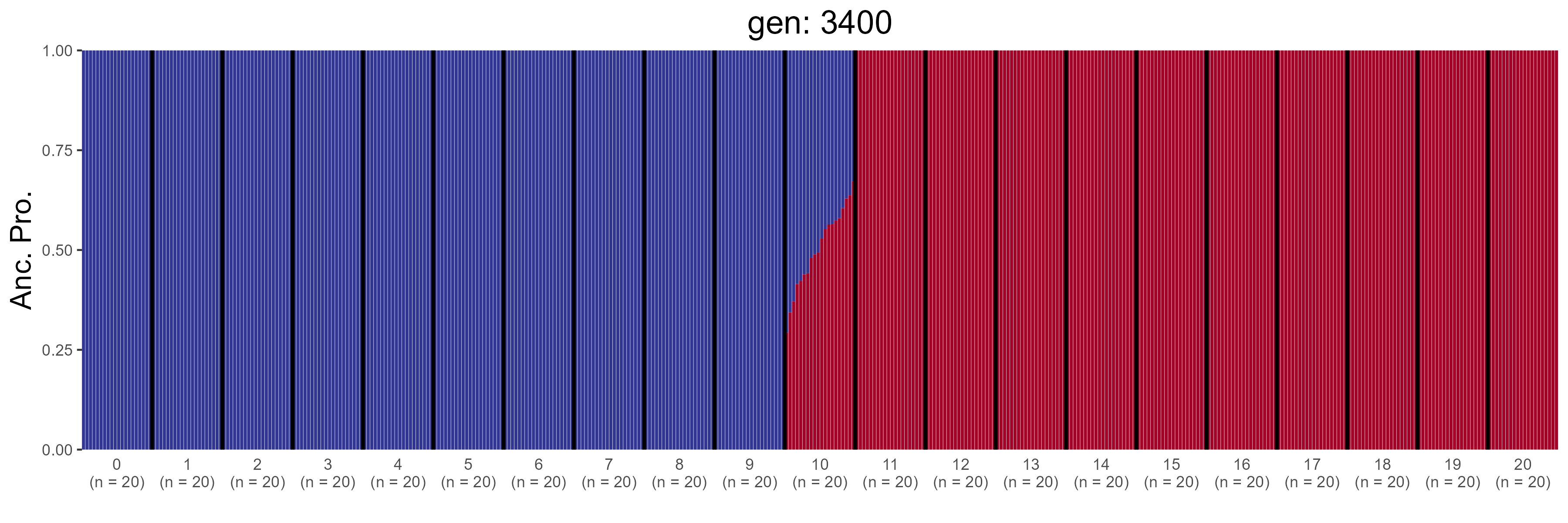

### GBC.3600.str.png

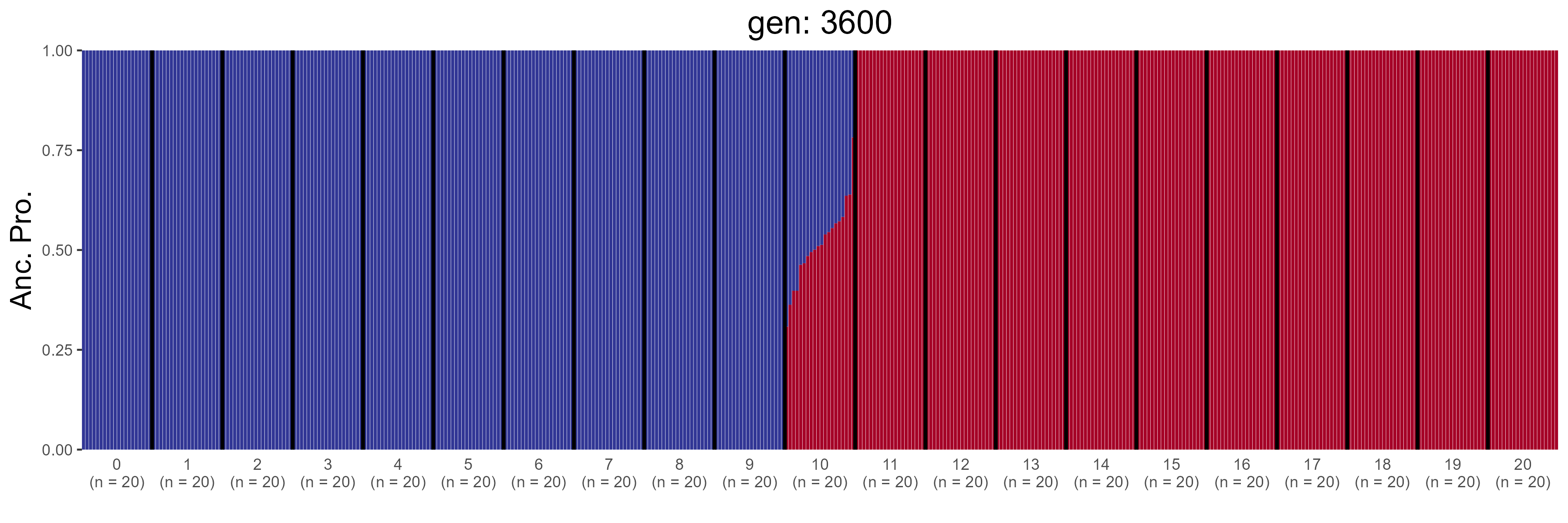

### GBC.3800.str.png

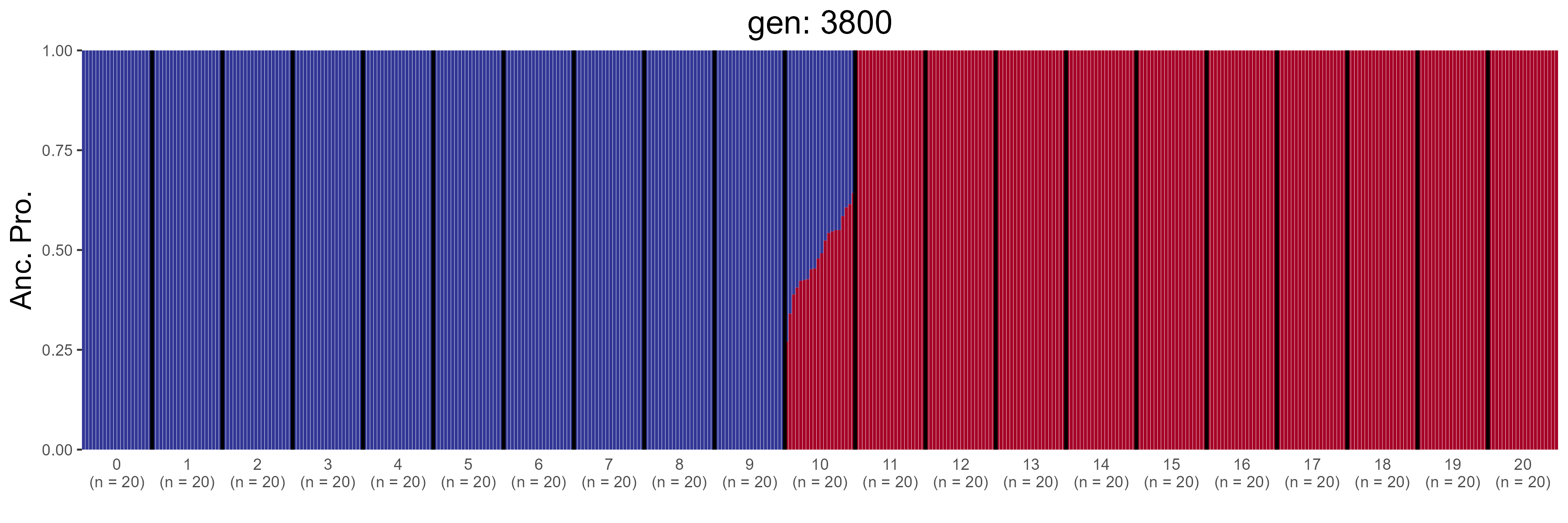

### GBC.4000.str.png

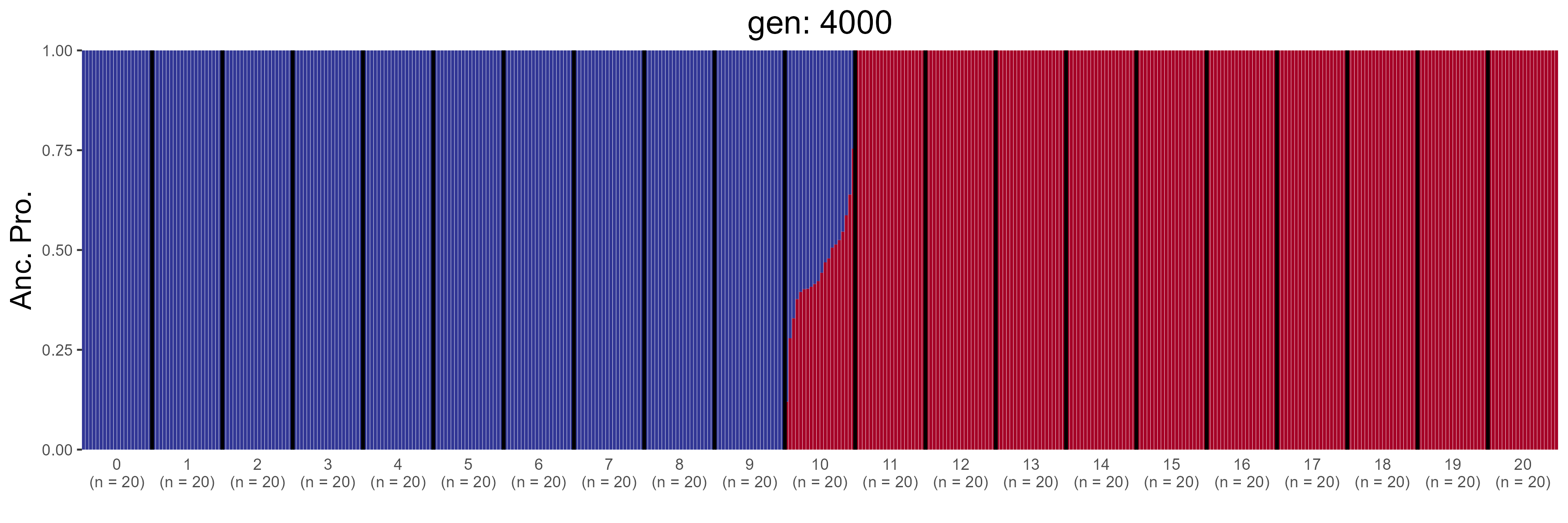

### GBC.4200.str.png

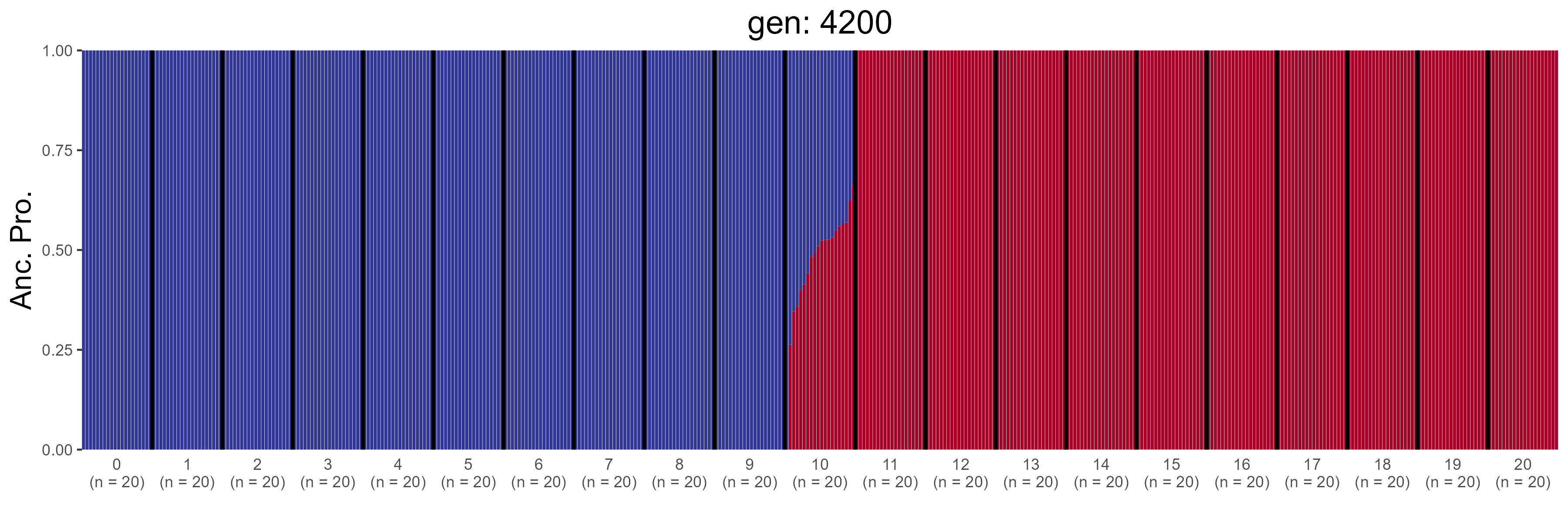

### GBC.4400.str.png

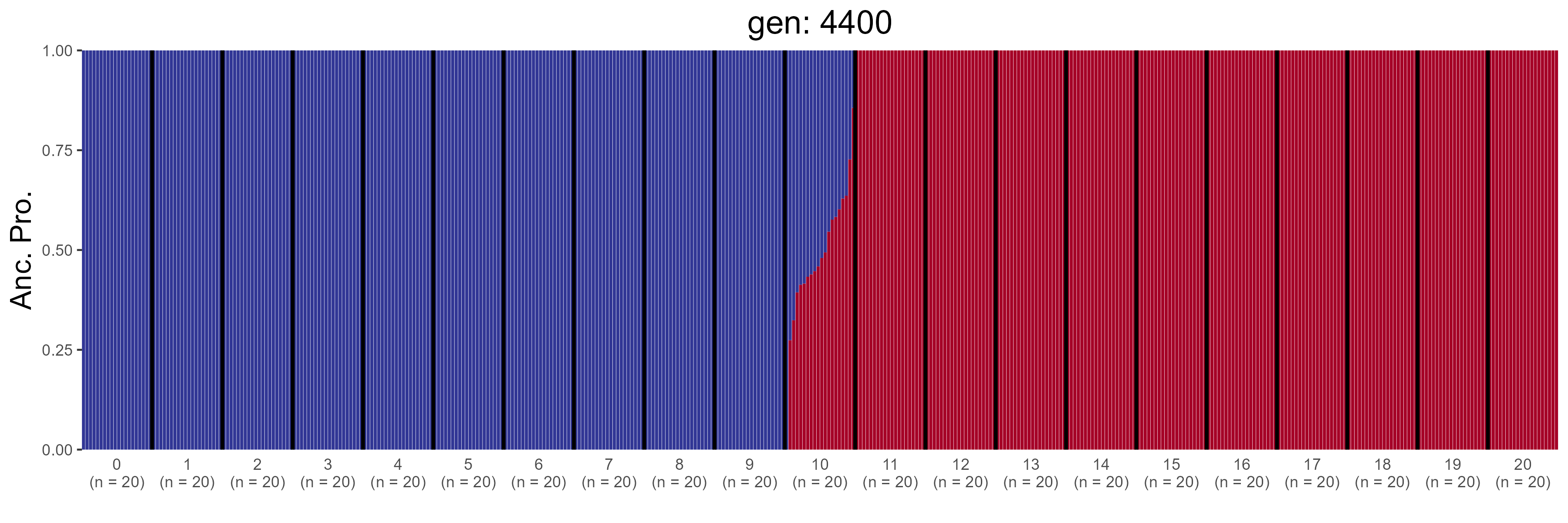

### GBC.4600.str.png

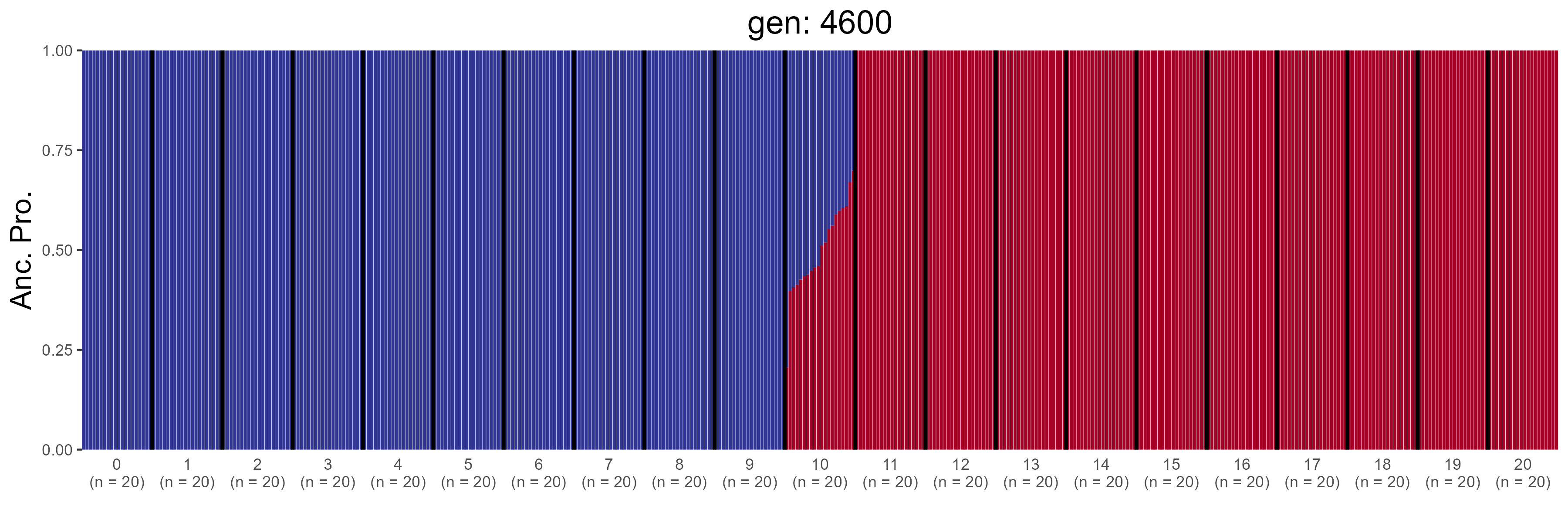

### GBC.4800.str.png

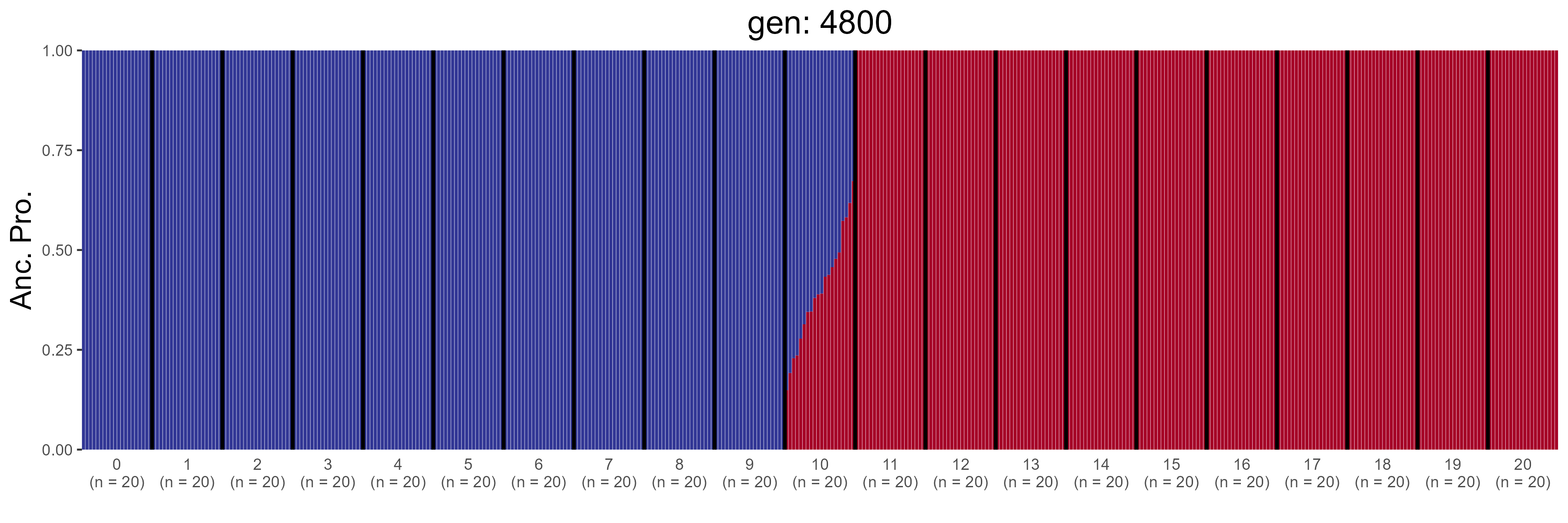

### GBC.5000.str.png

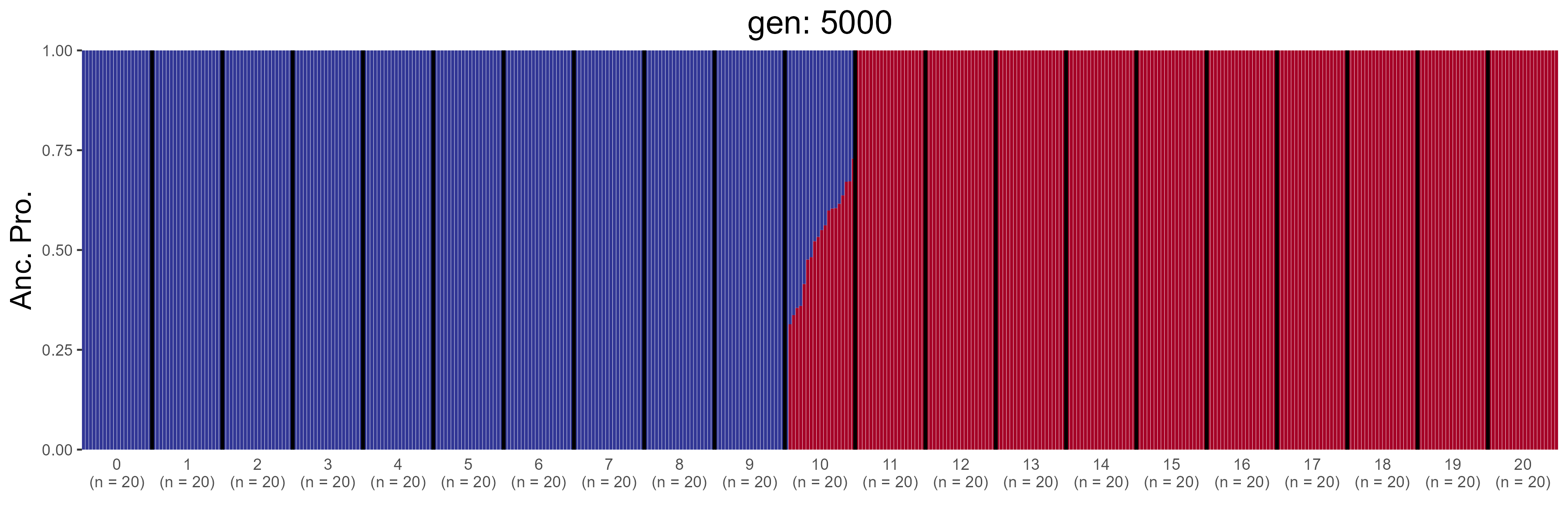

### GBC.5200.str.png

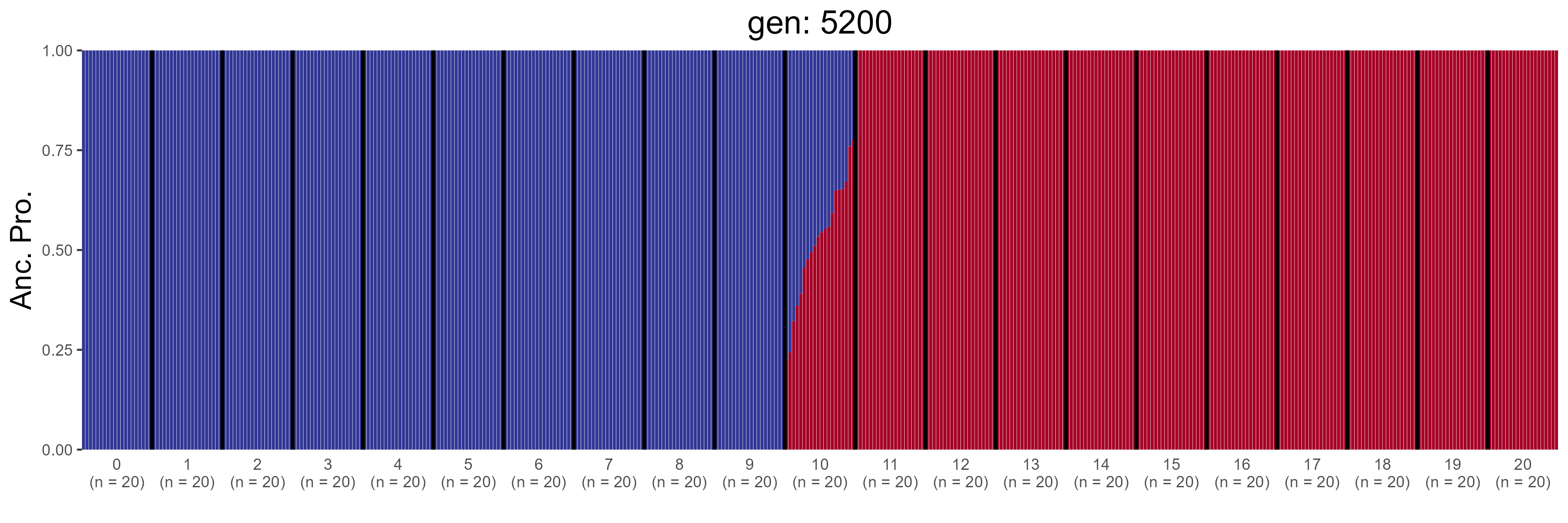

### GBC.5400.str.png

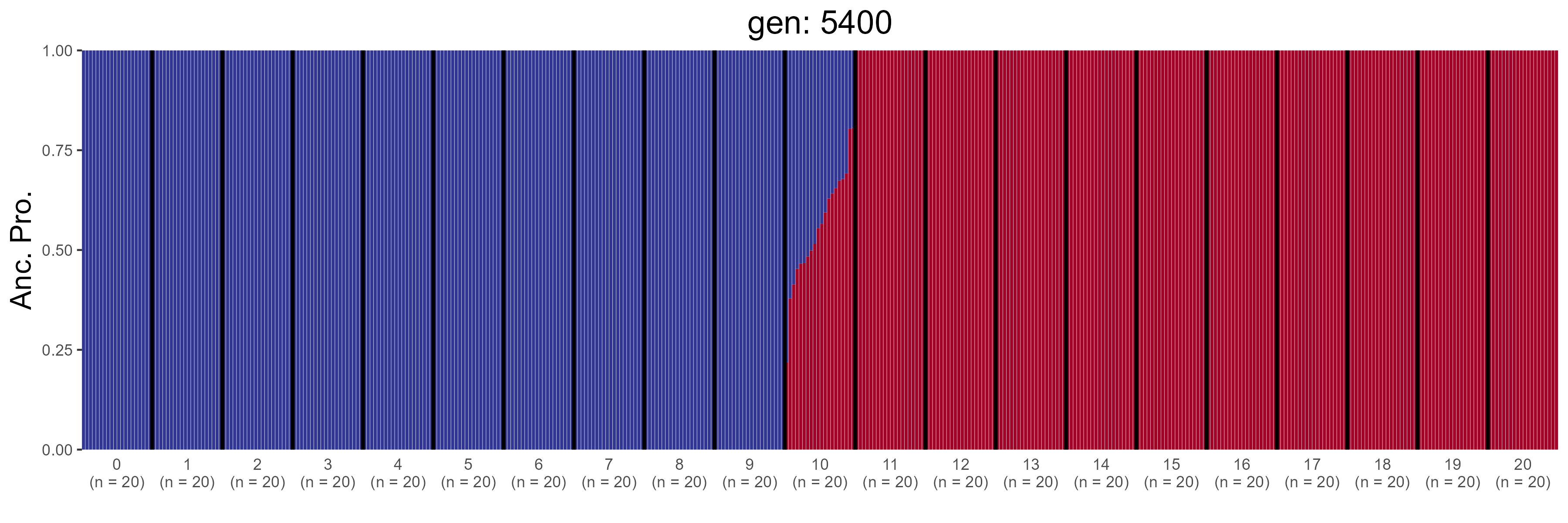

### GBC.5600.str.png

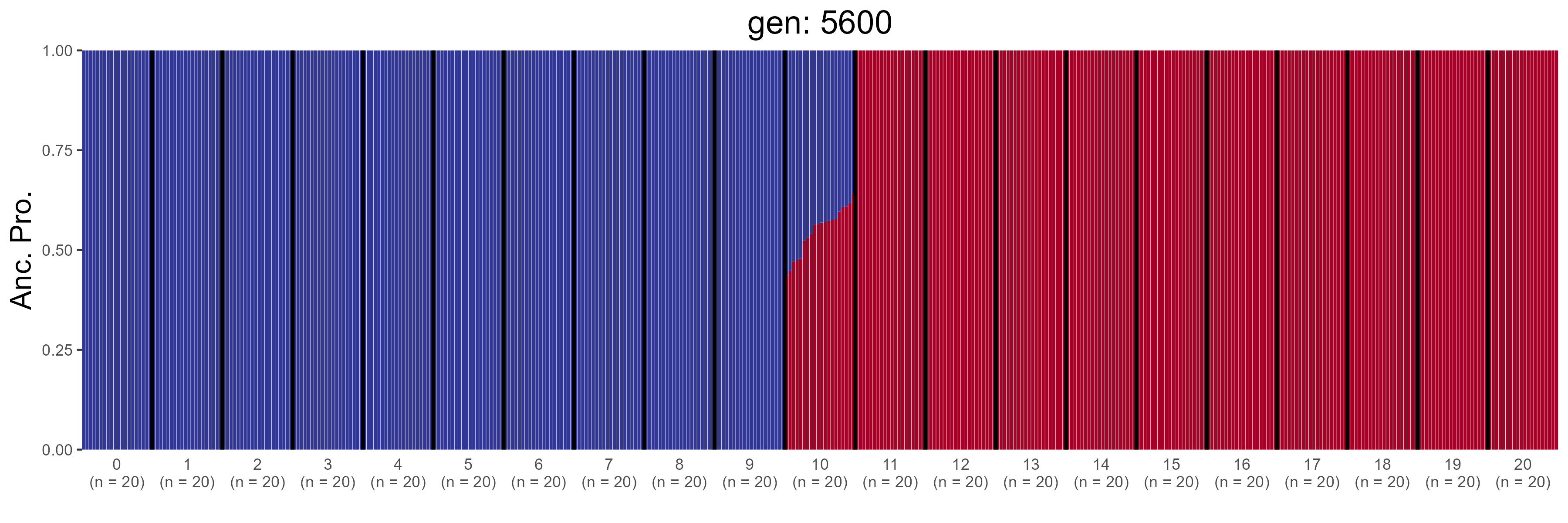

### GBC.5800.str.png

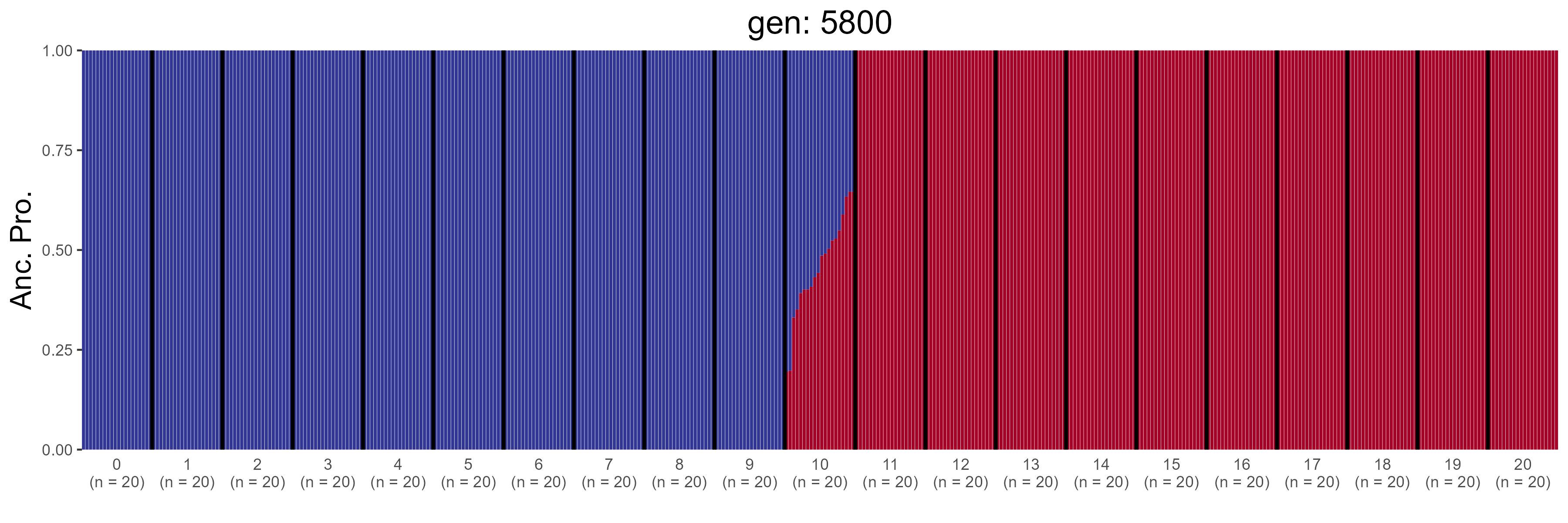

### GBC.6000.str.png

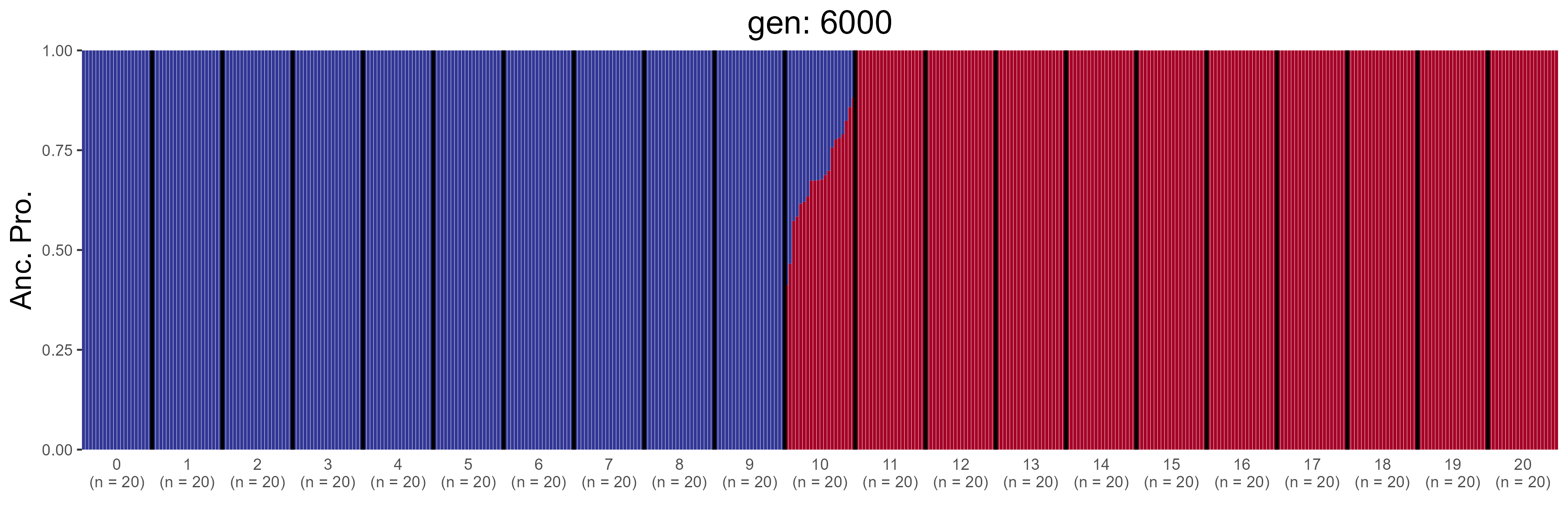

### GBC.6200.str.png

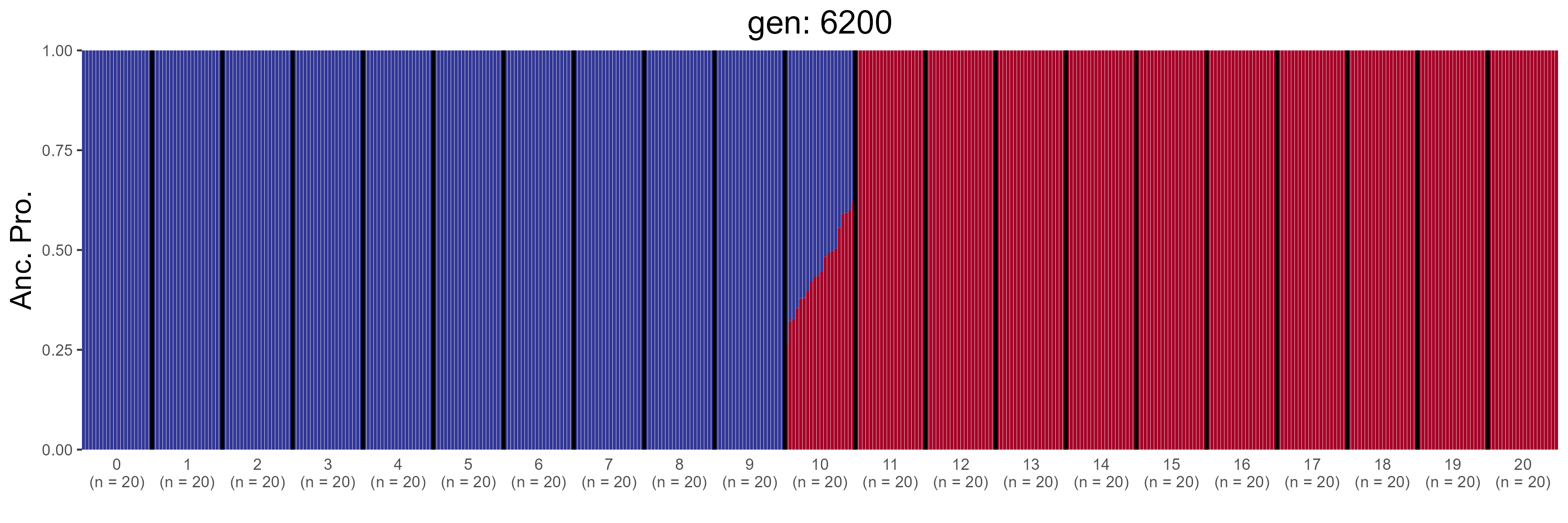

### GBC.6400.str.png

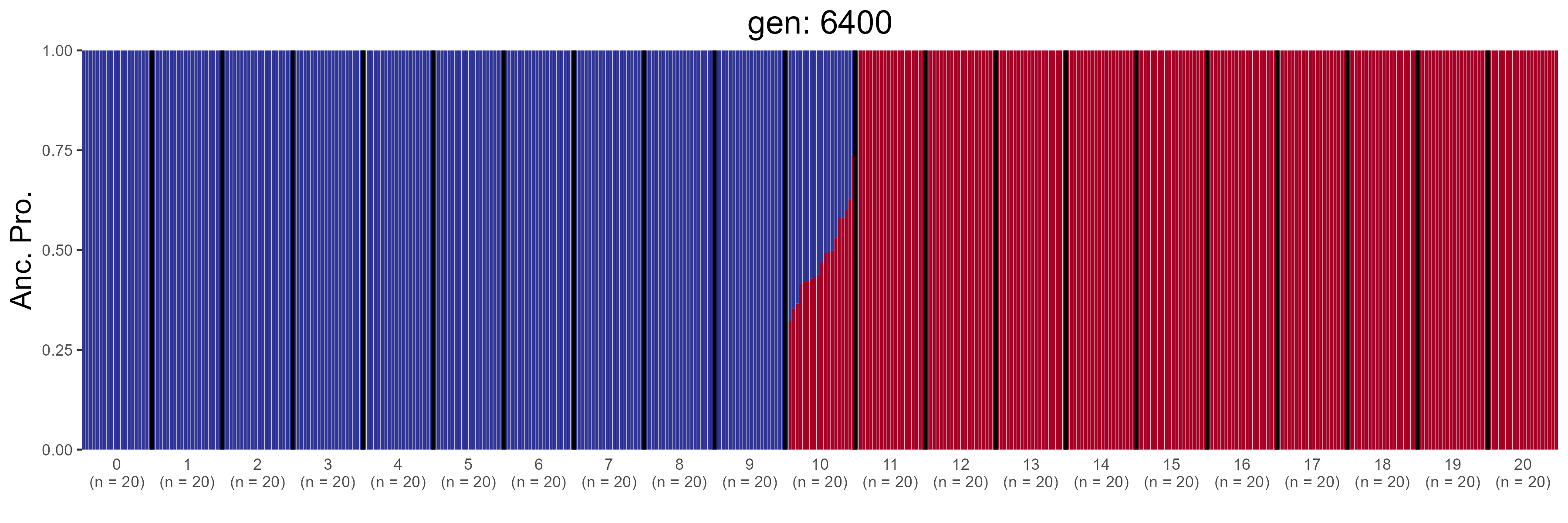

### GBC.6600.str.png

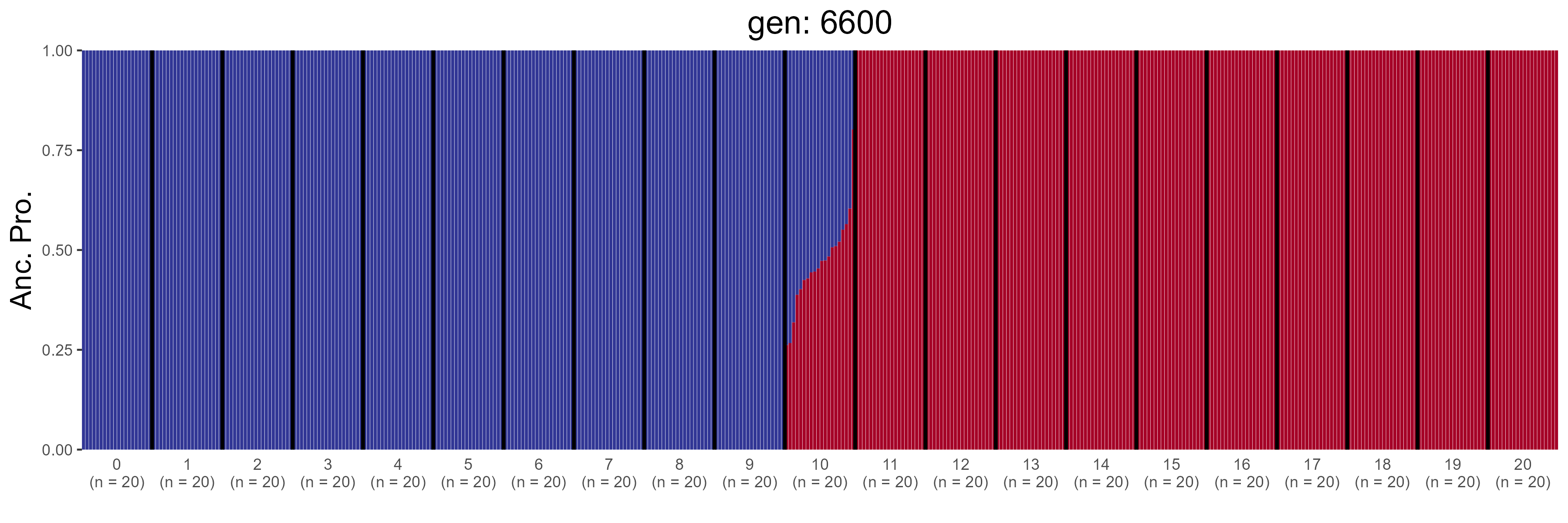

### GBC.6800.str.png

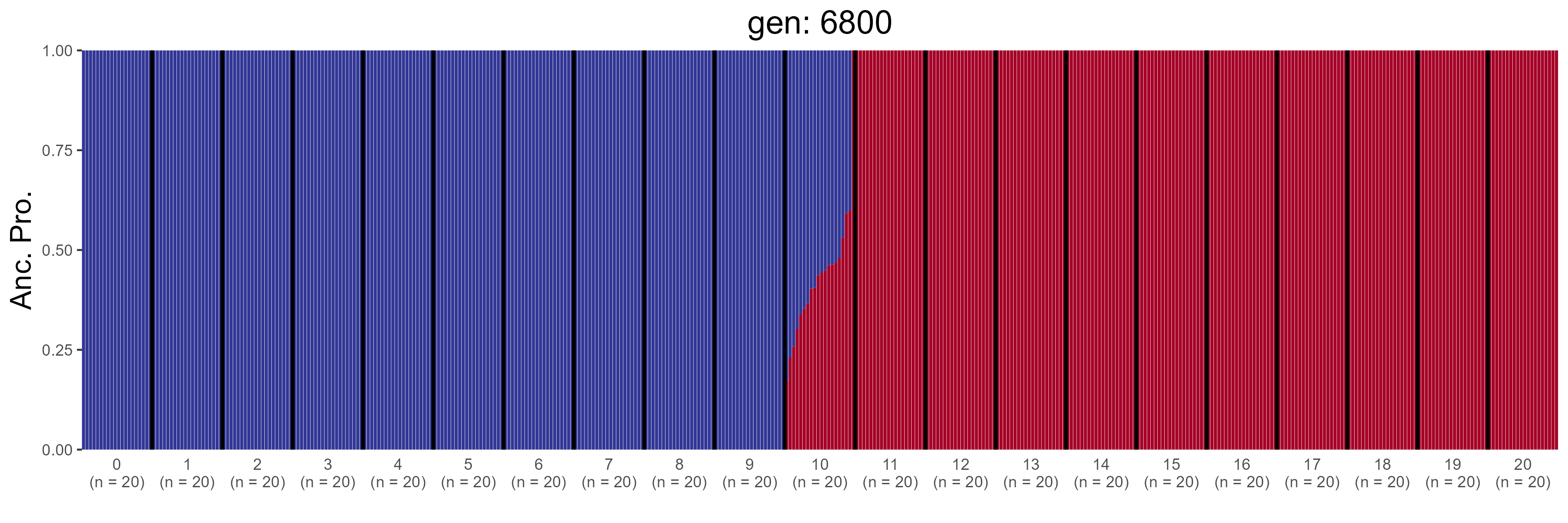

### GBC.7000.str.png

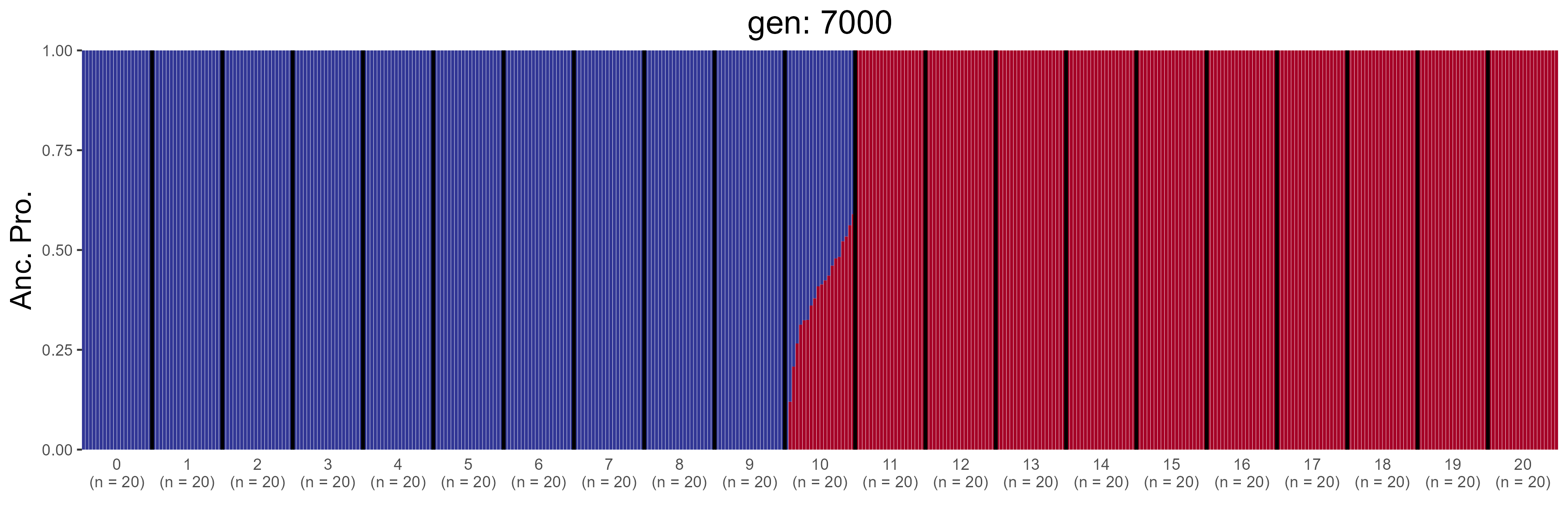

### GBC.7200.str.png

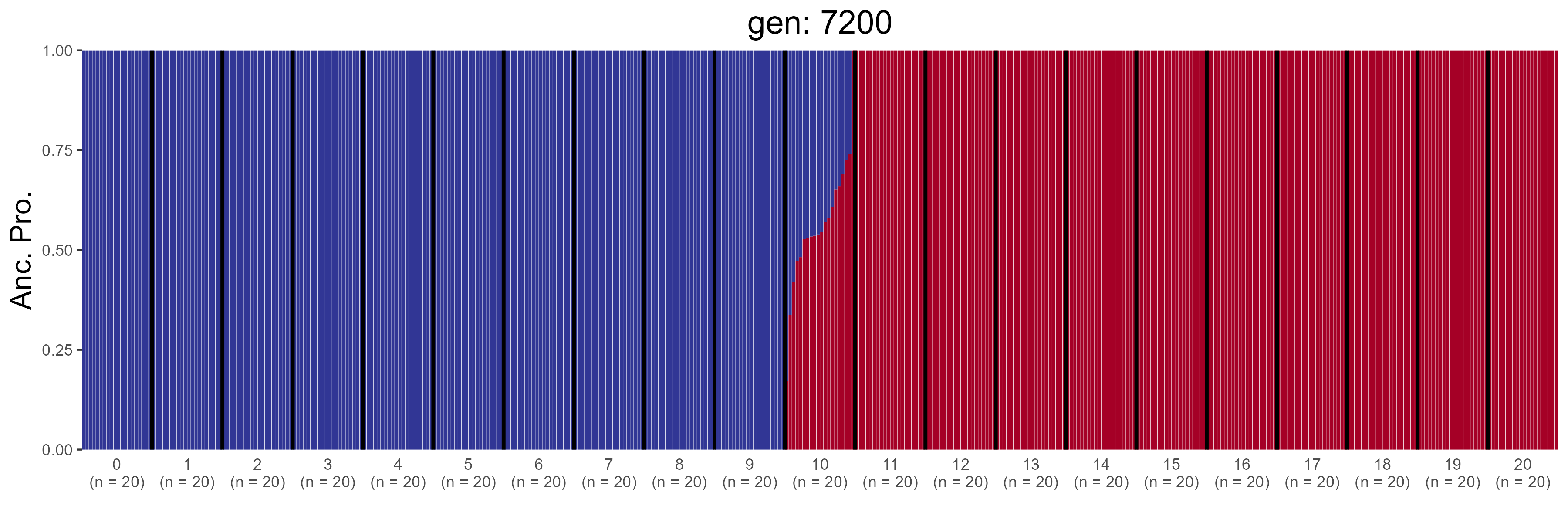

### GBC.7400.str.png

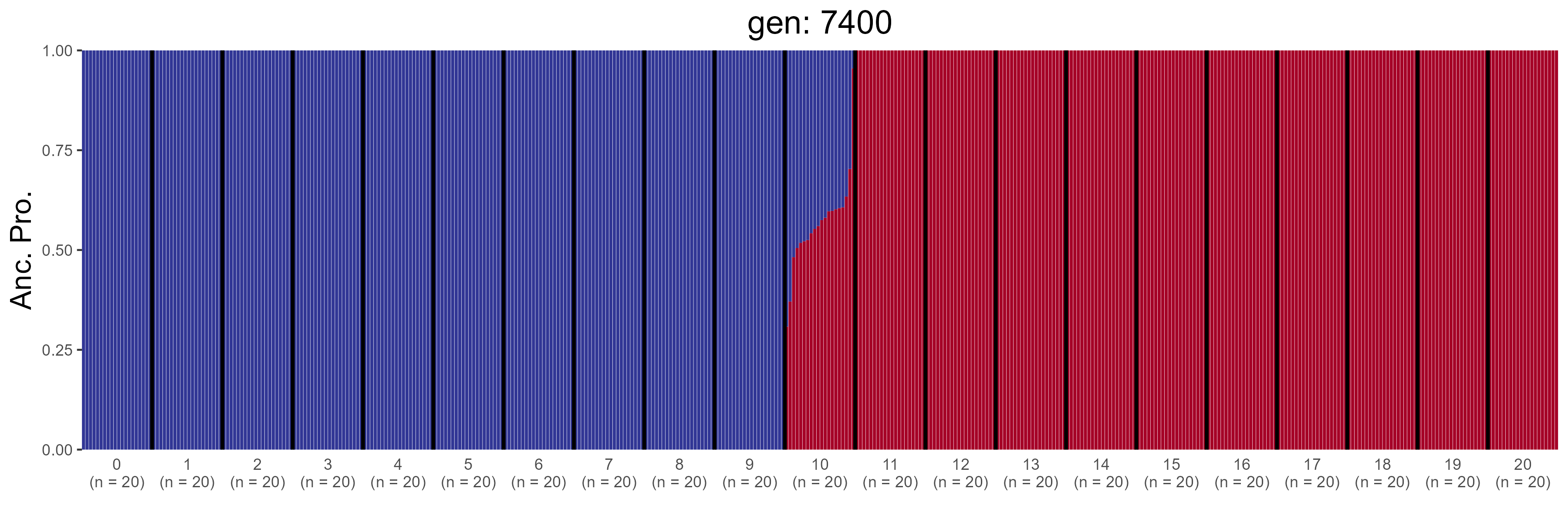

### GBC.7600.str.png

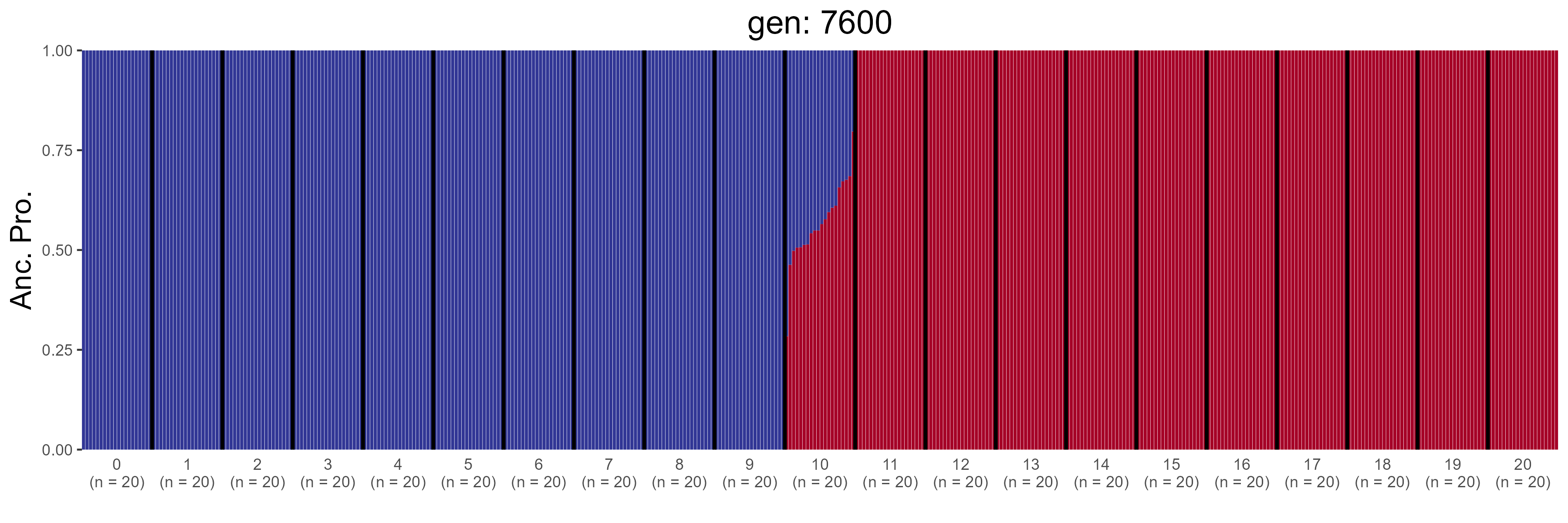

### GBC.7800.str.png

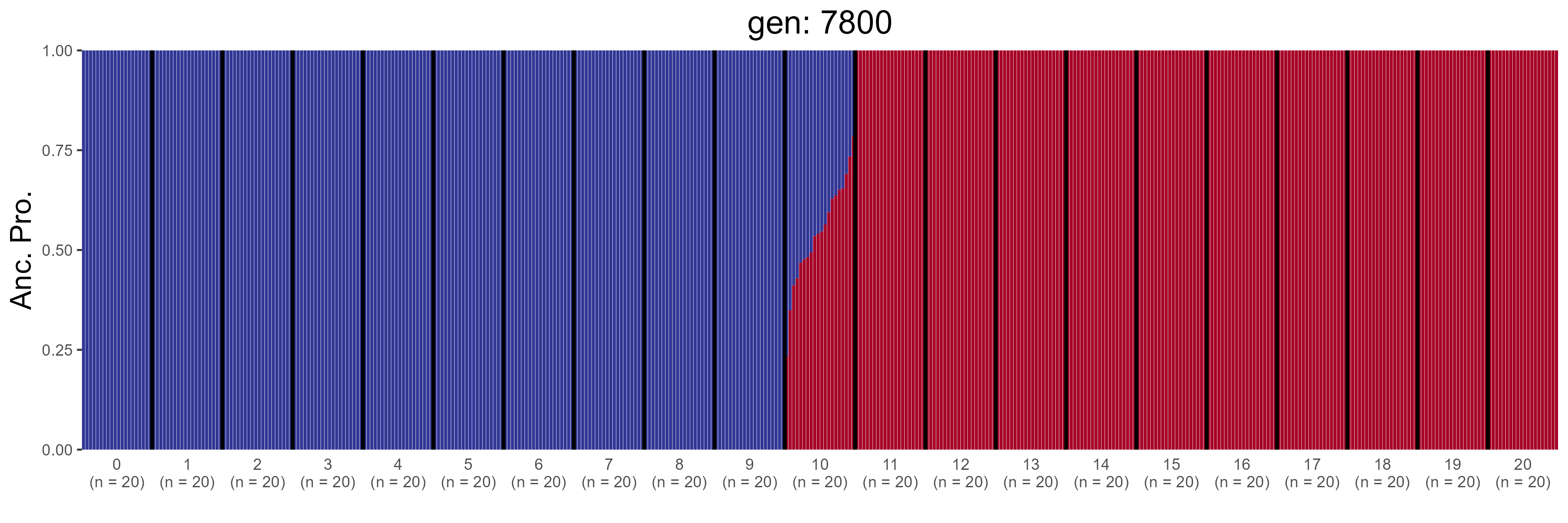

### GBC.8000.str.png

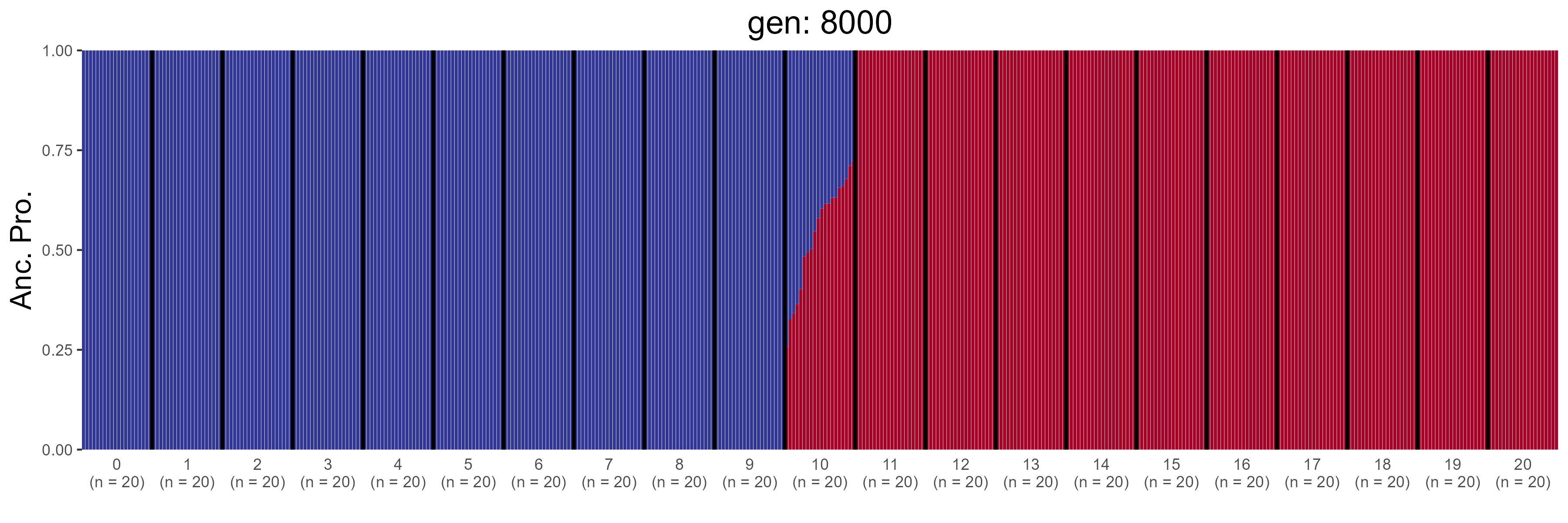

### GBC.8200.str.png

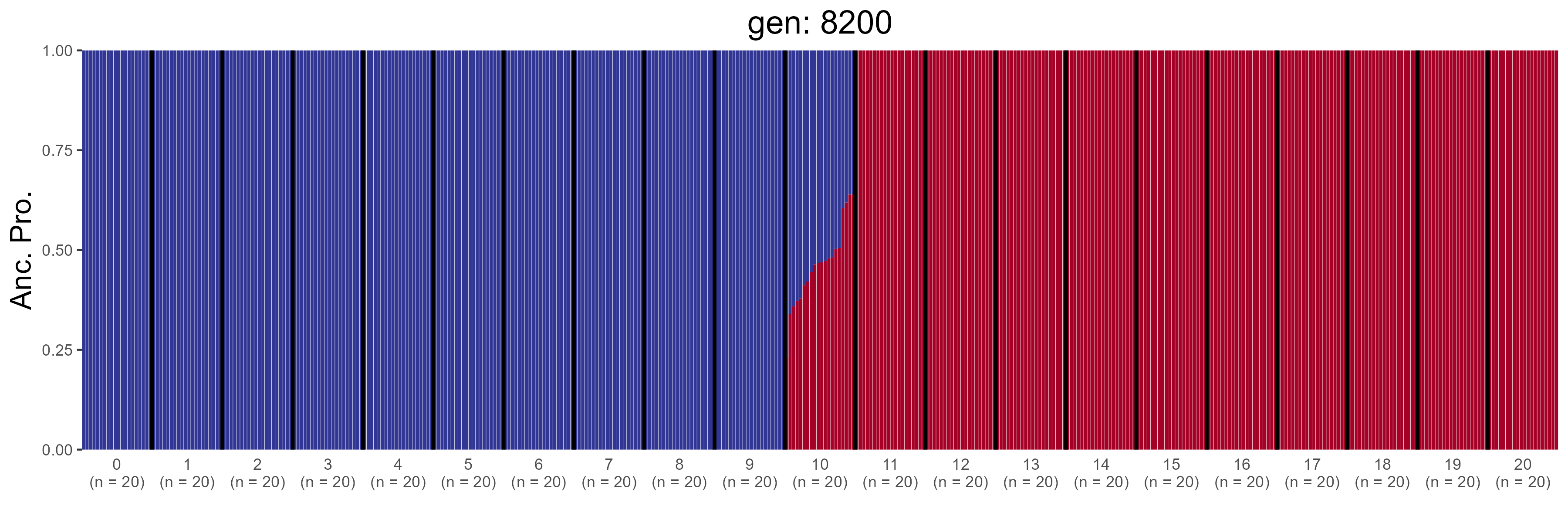

### GBC.8400.str.png

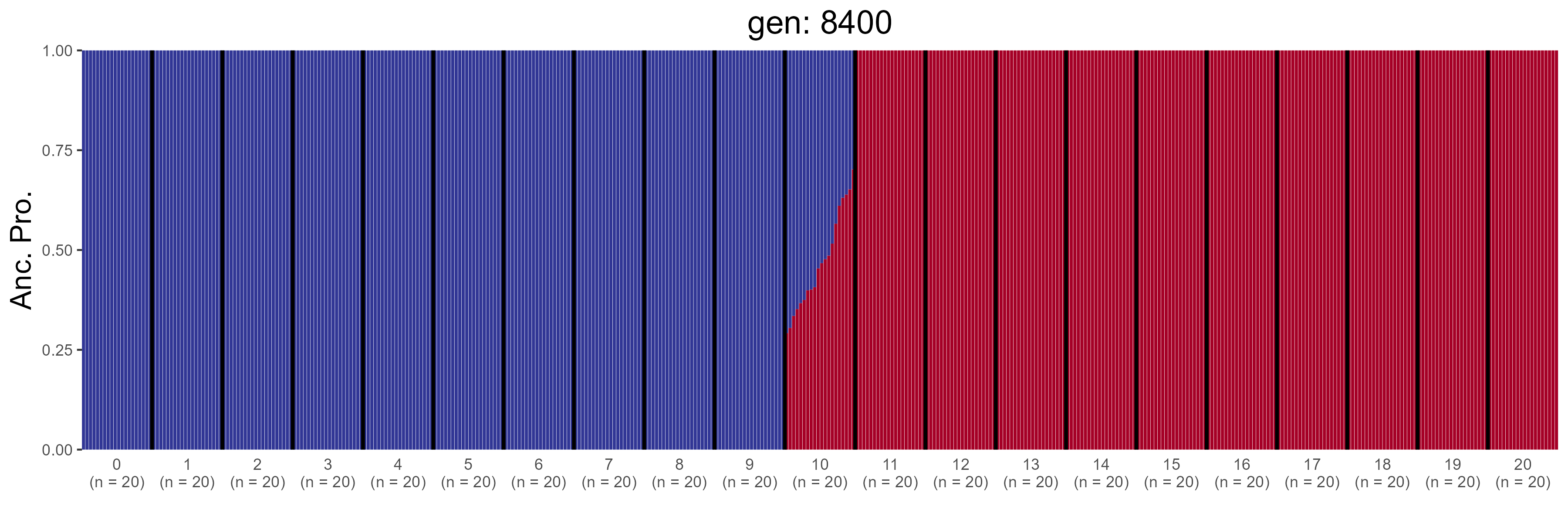

### GBC.8600.str.png

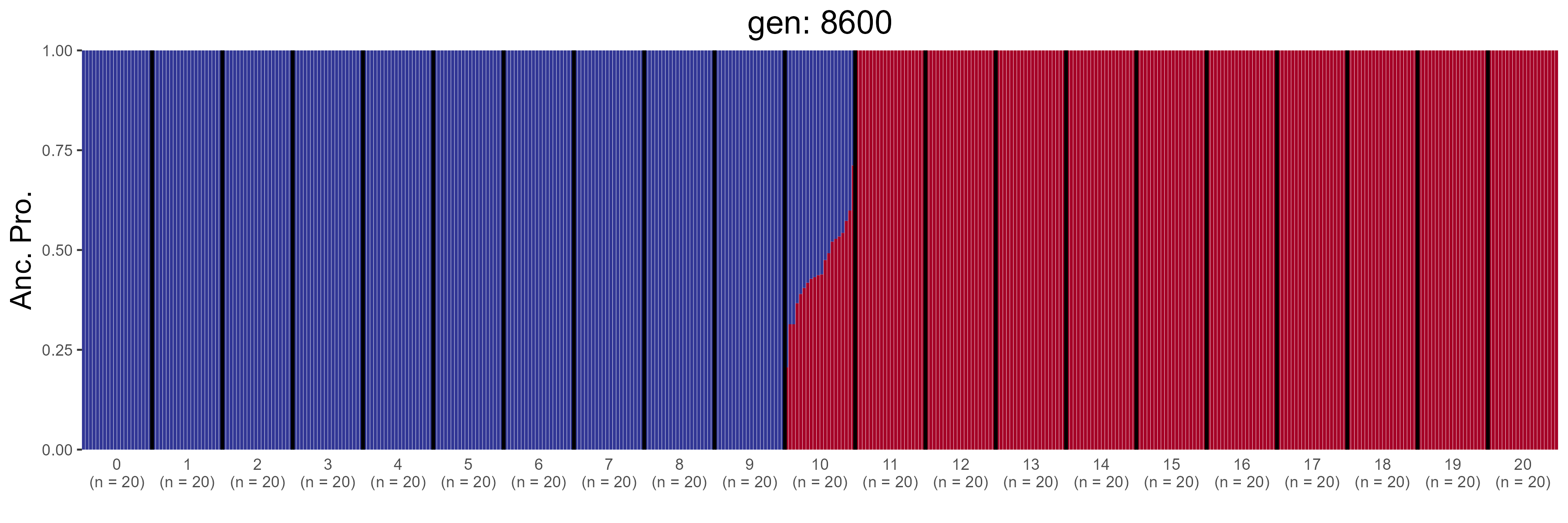

### GBC.8800.str.png

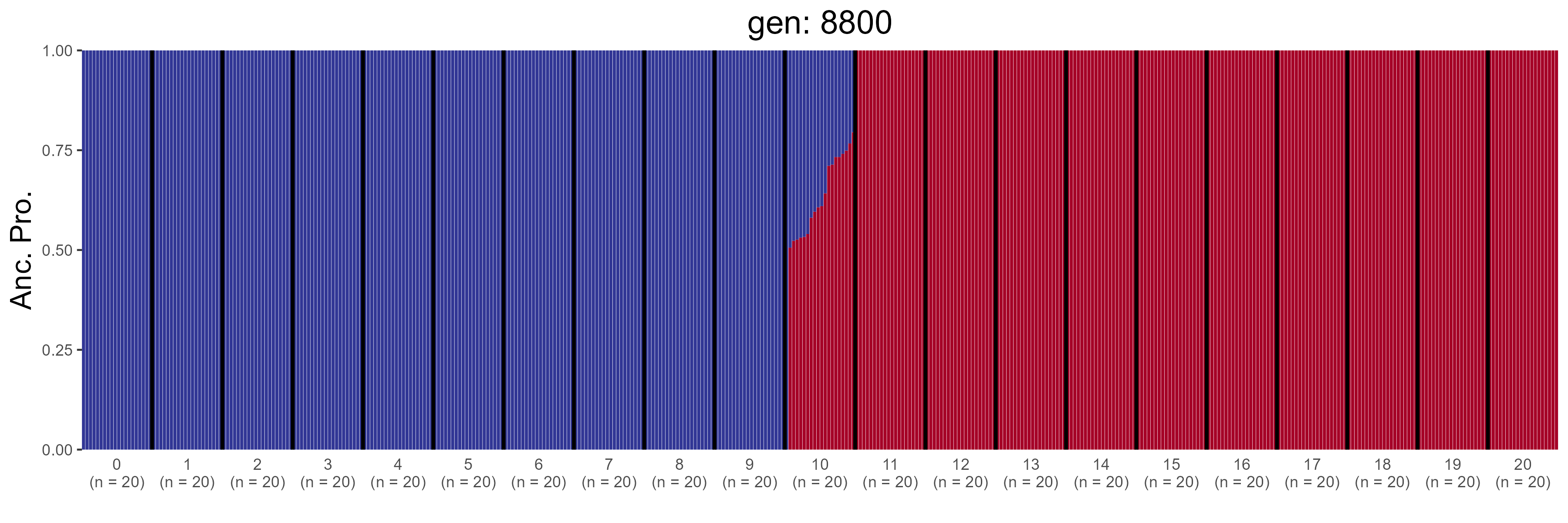

### GBC.9000.str.png

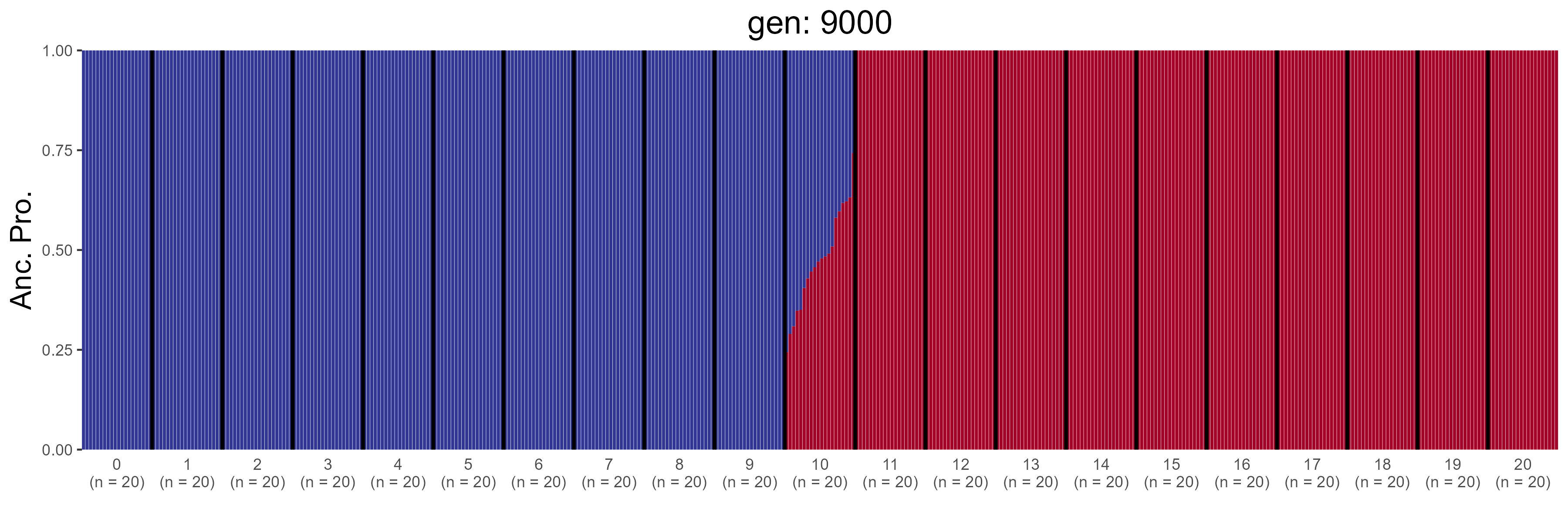

### GBC.9200.str.png

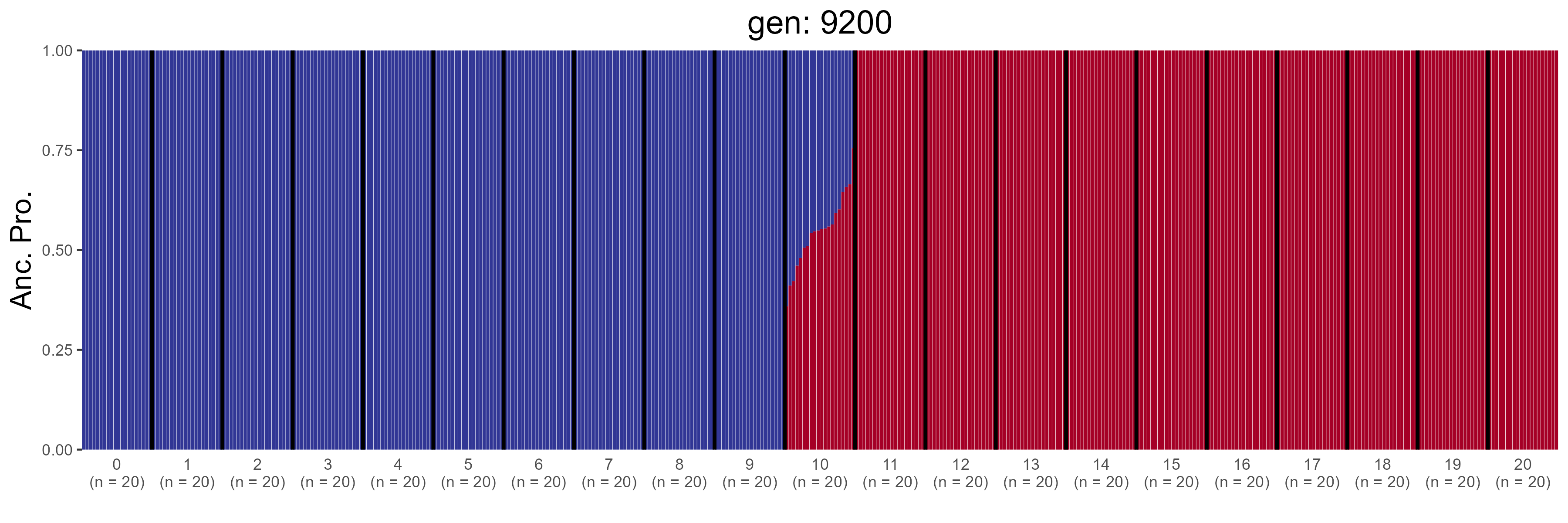

### GBC.9400.str.png

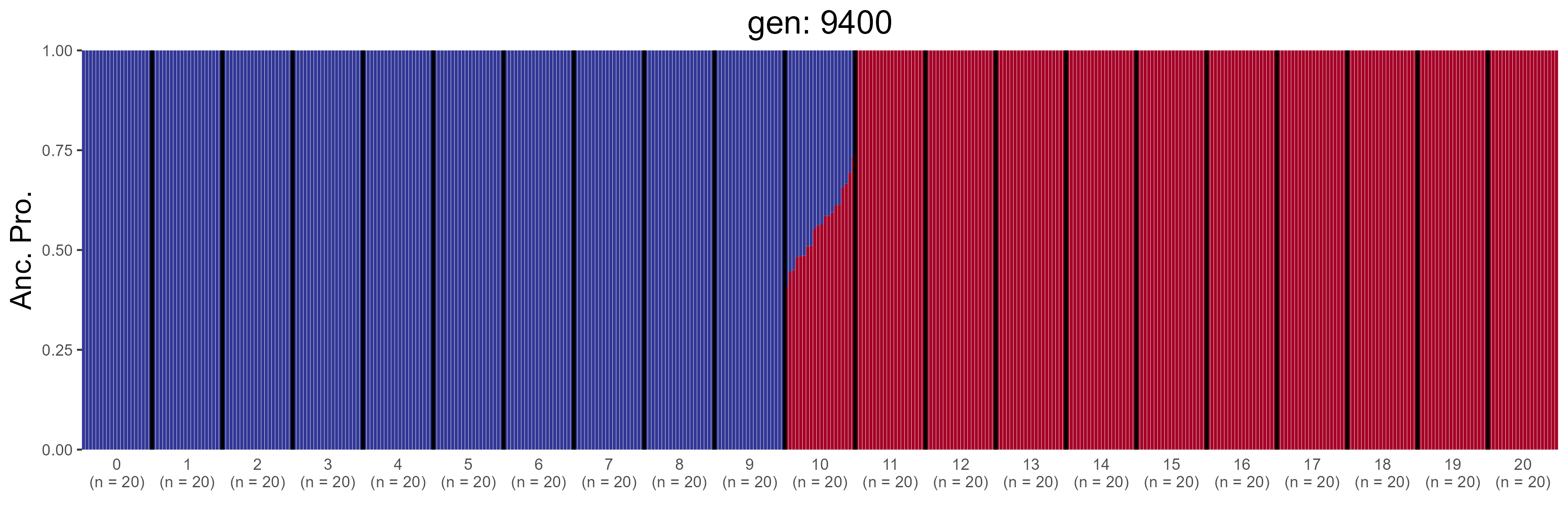

### GBC.9600.str.png

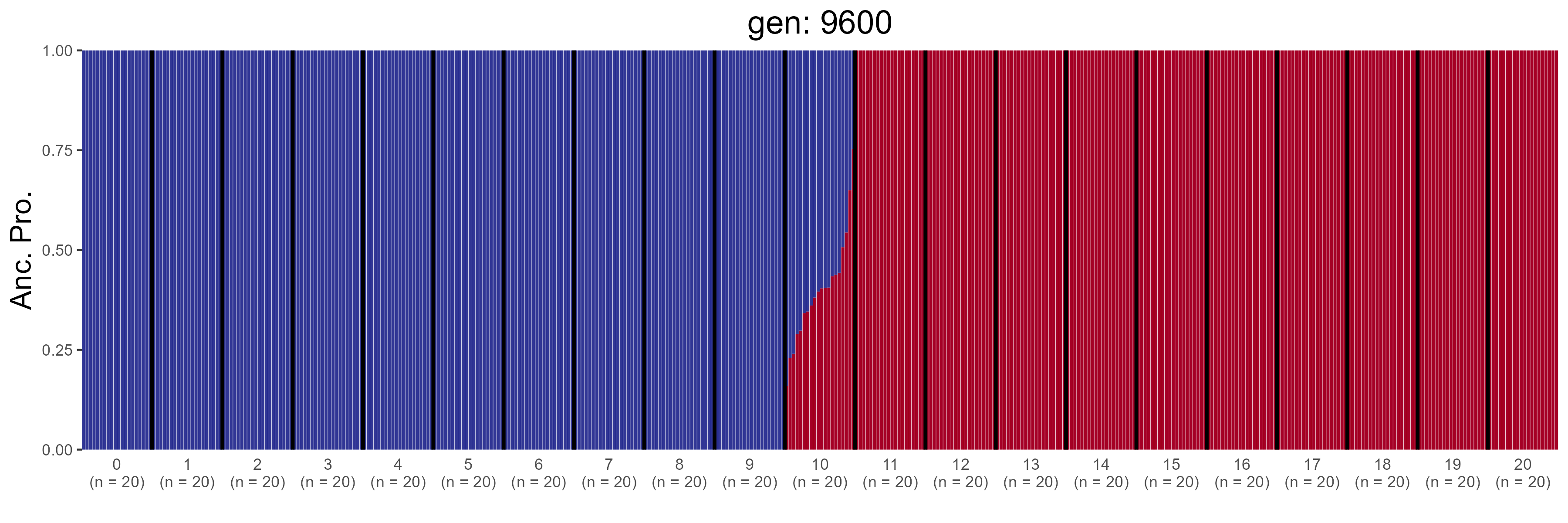

### GBC.9800.str.png

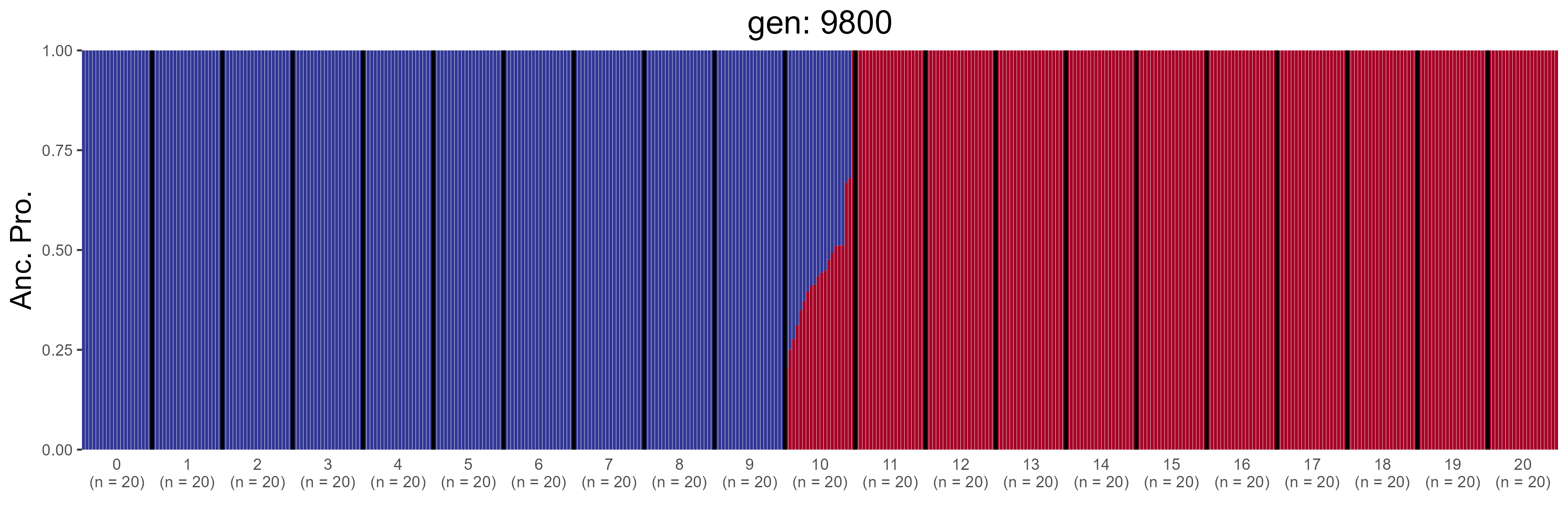

### GBC.10000.str.png

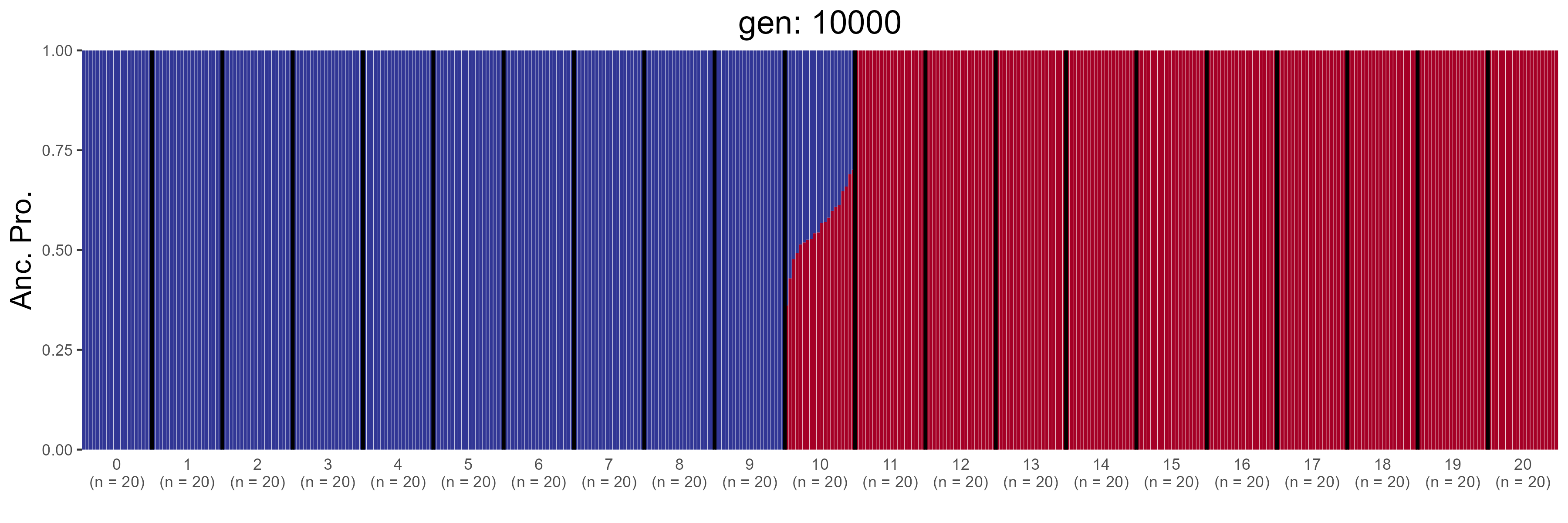

### GBC.10200.str.png

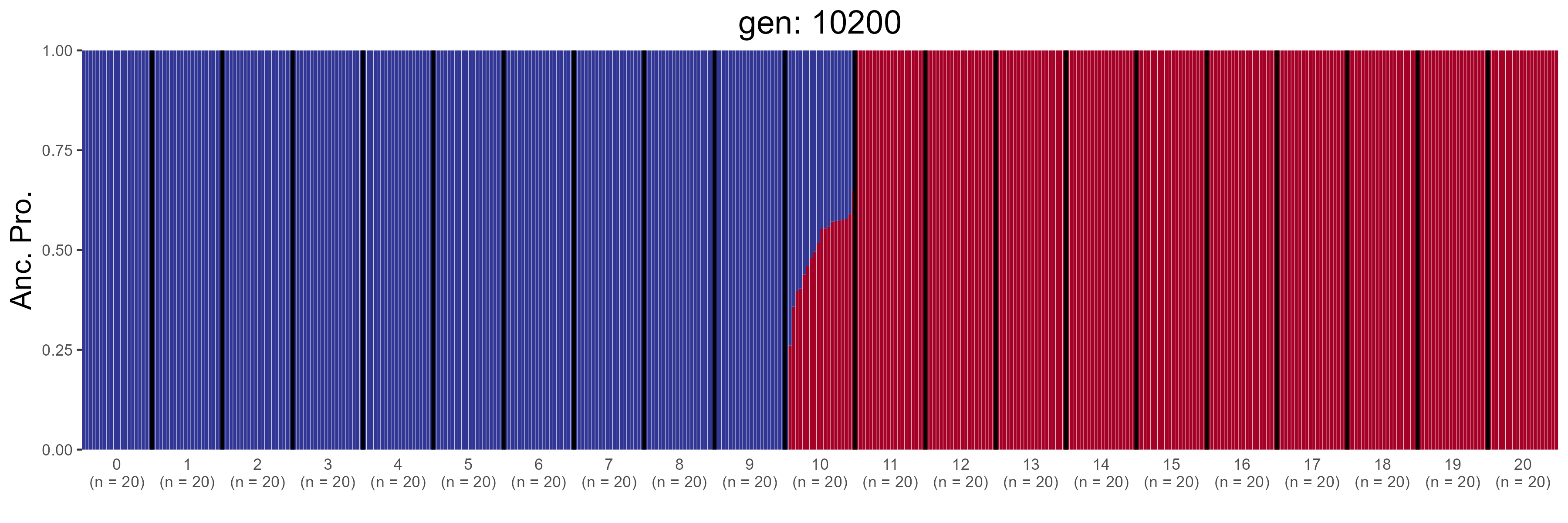

### GBC.10400.str.png

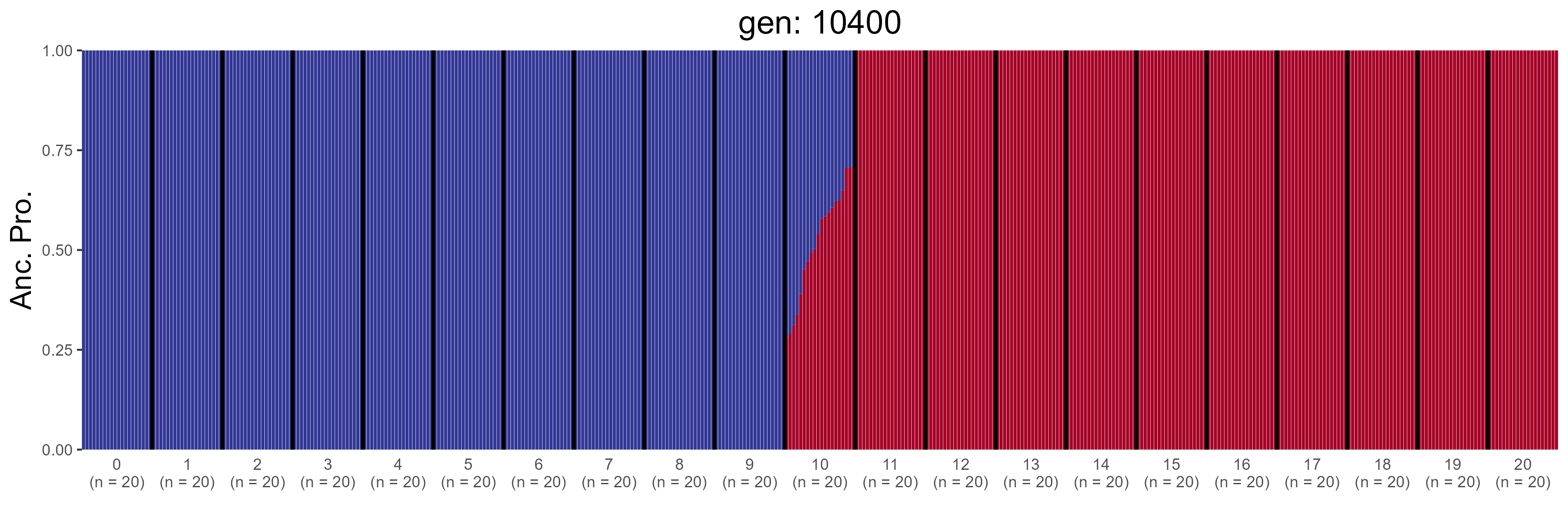

### GBC.10600.str.png

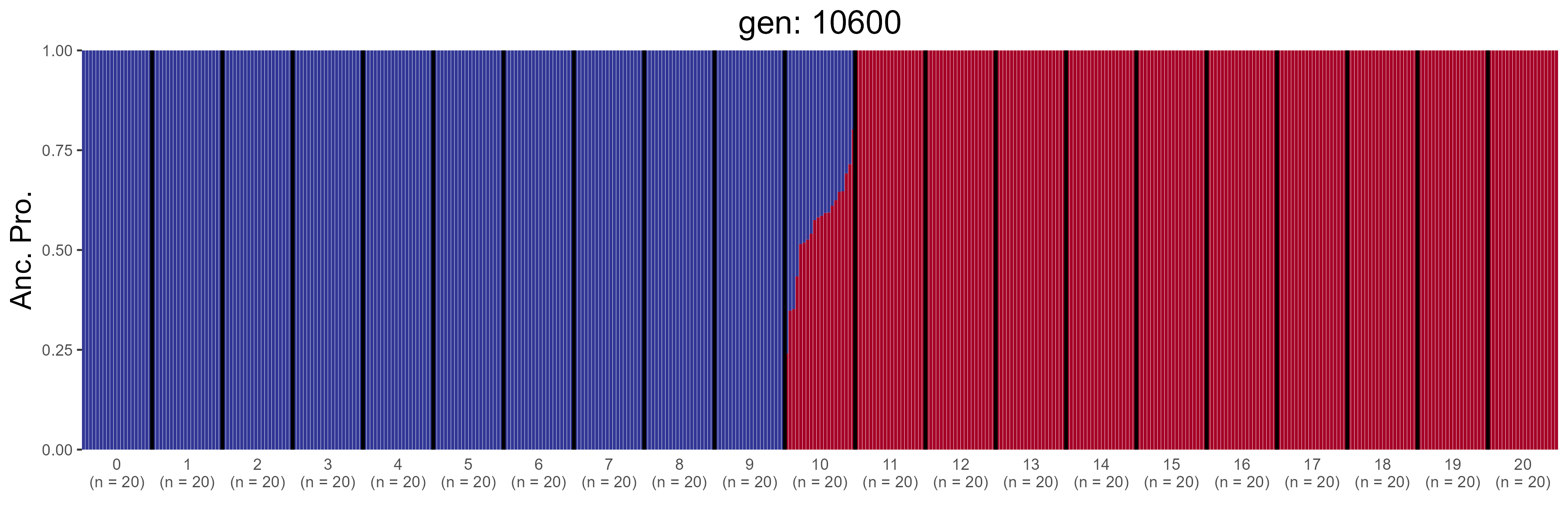

### GBC.10800.str.png

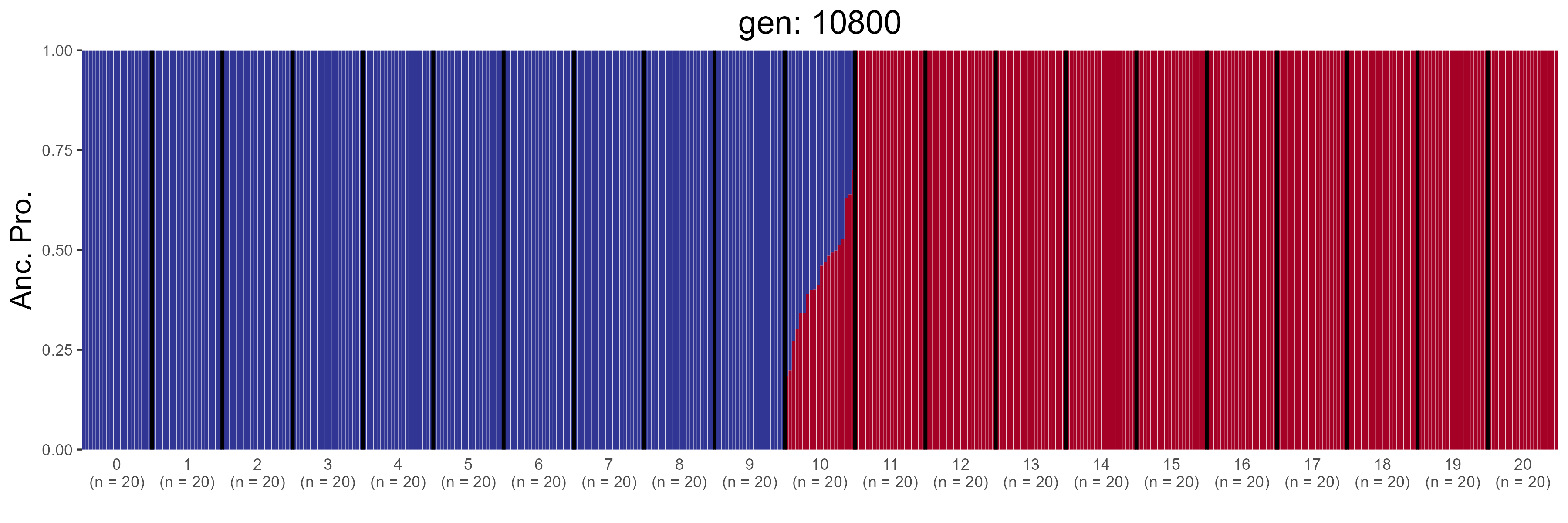

### GBC.11000.str.png

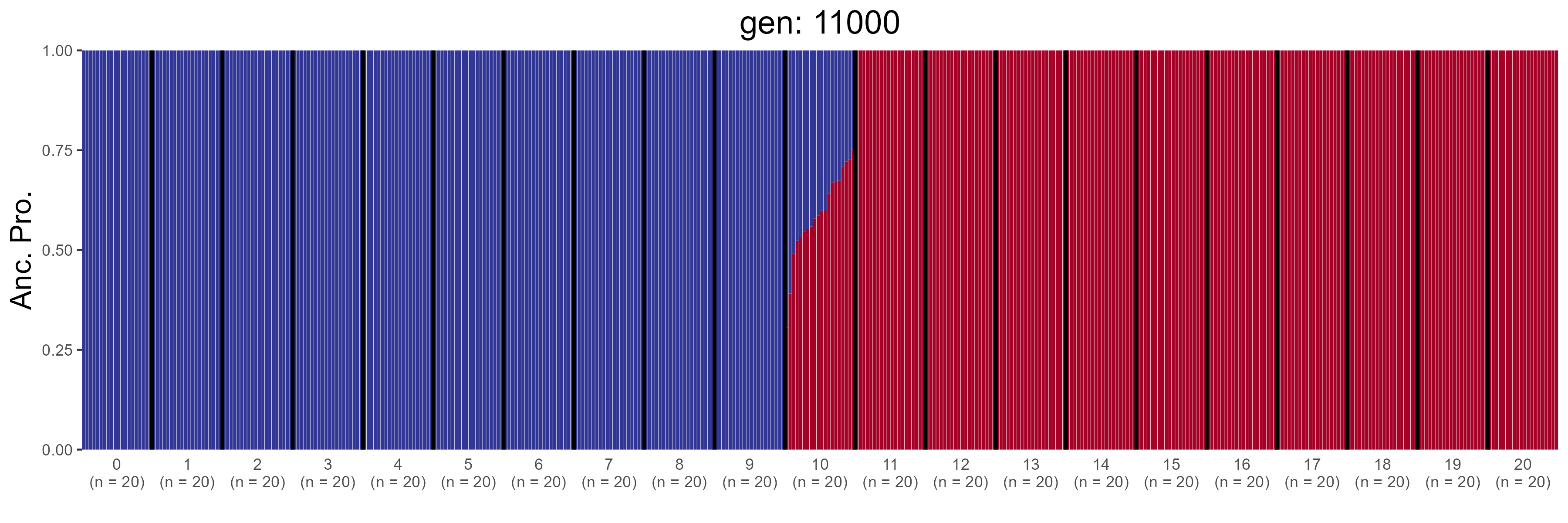

### GBC.11200.str.png

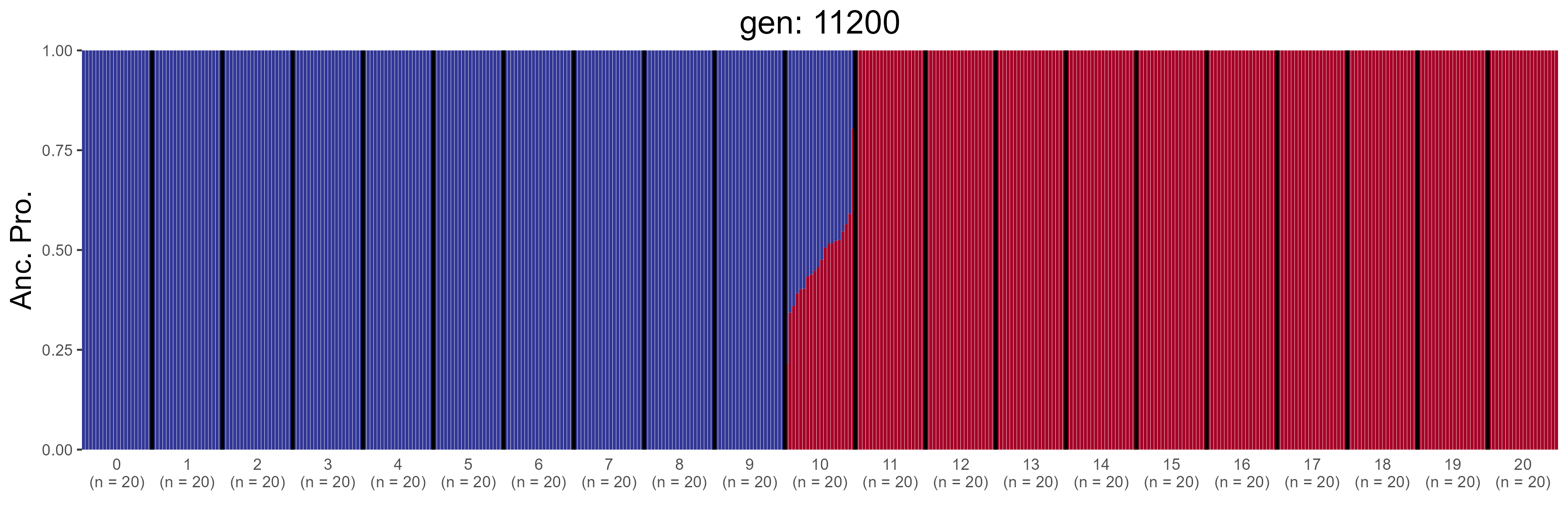

### GBC.11400.str.png

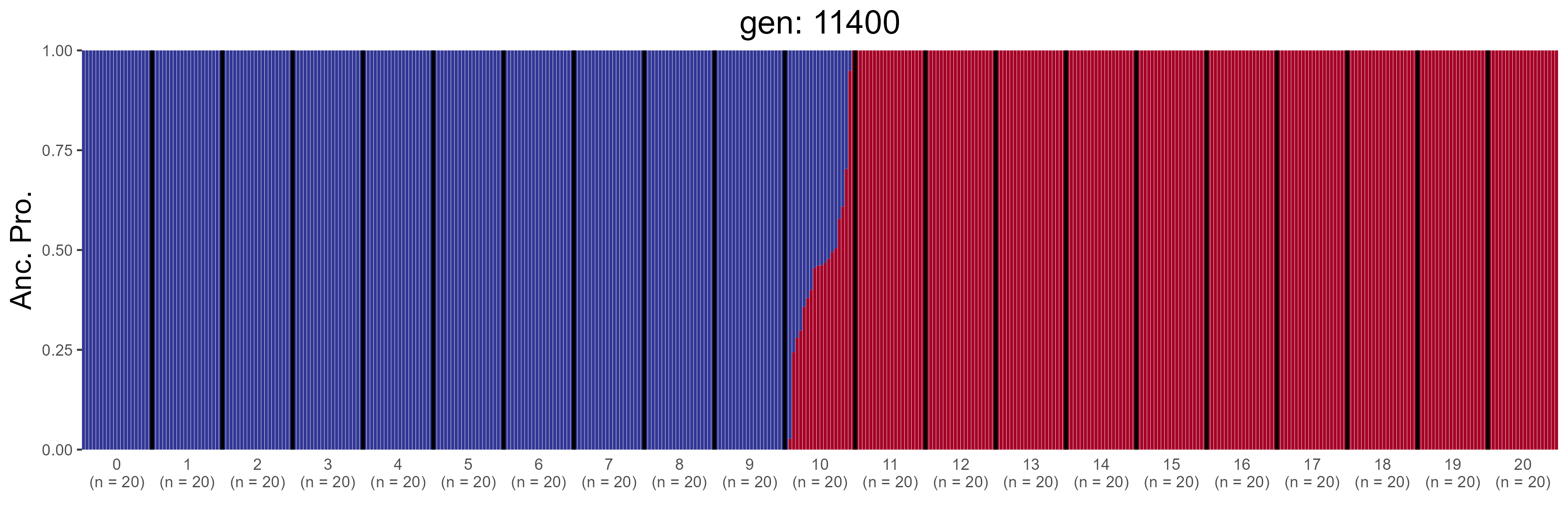

### GBC.11600.str.png

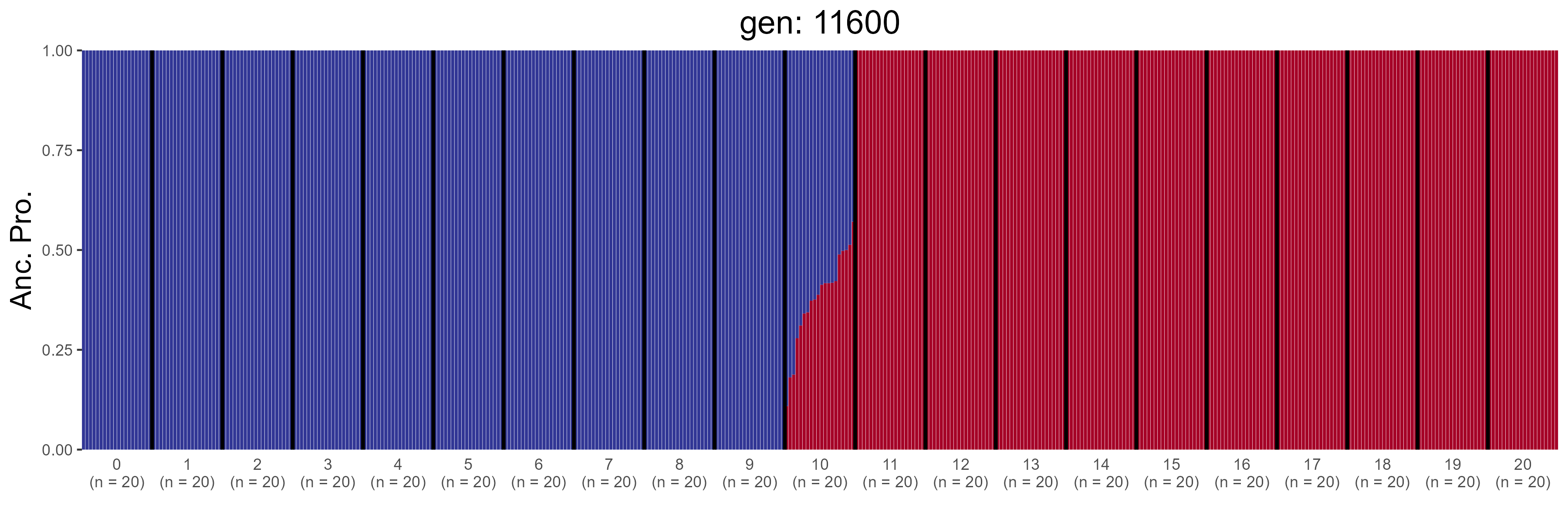

### GBC.11800.str.png

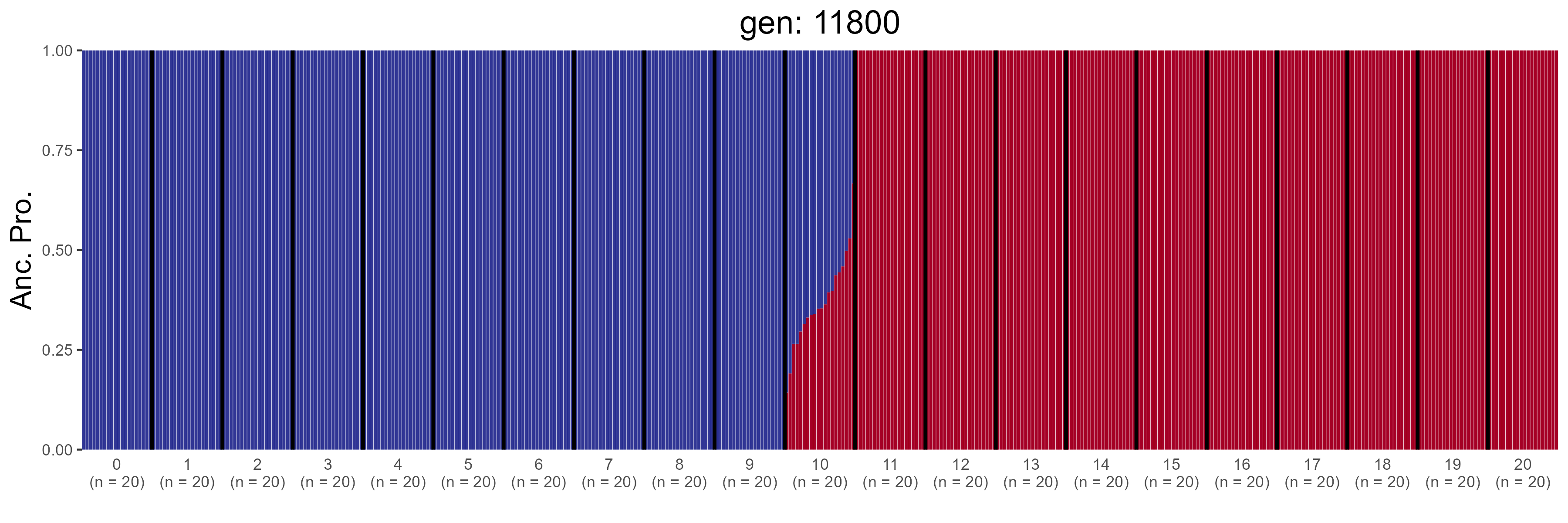

### GBC.12000.str.png

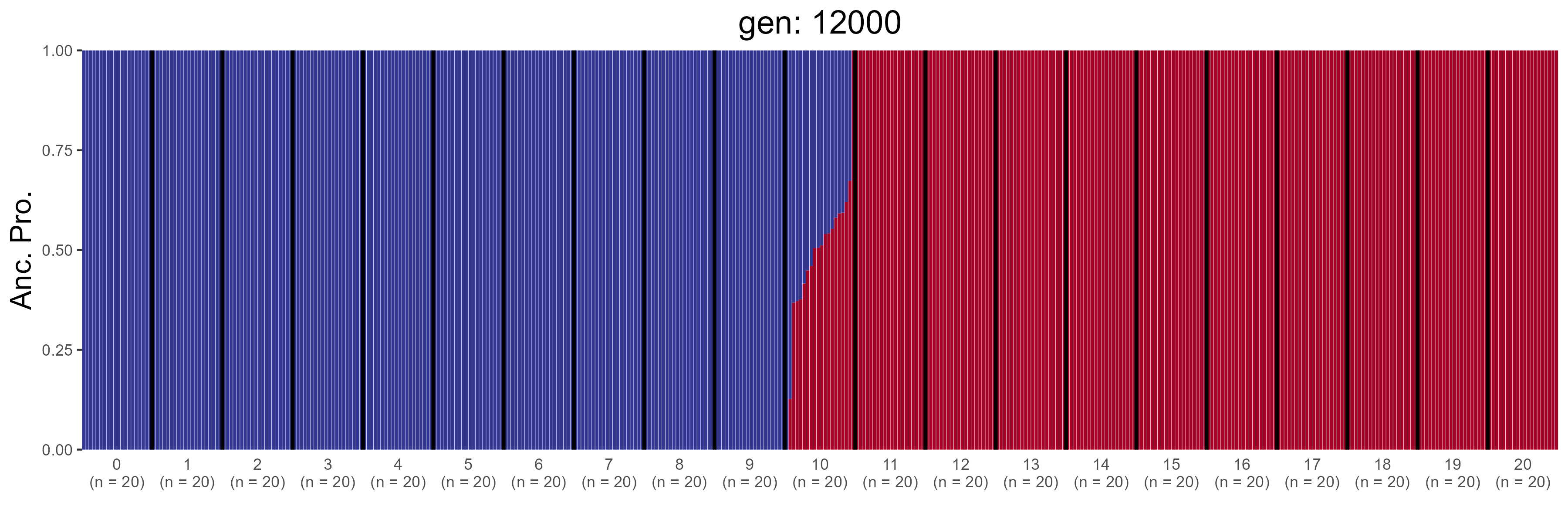

### GBC.12200.str.png

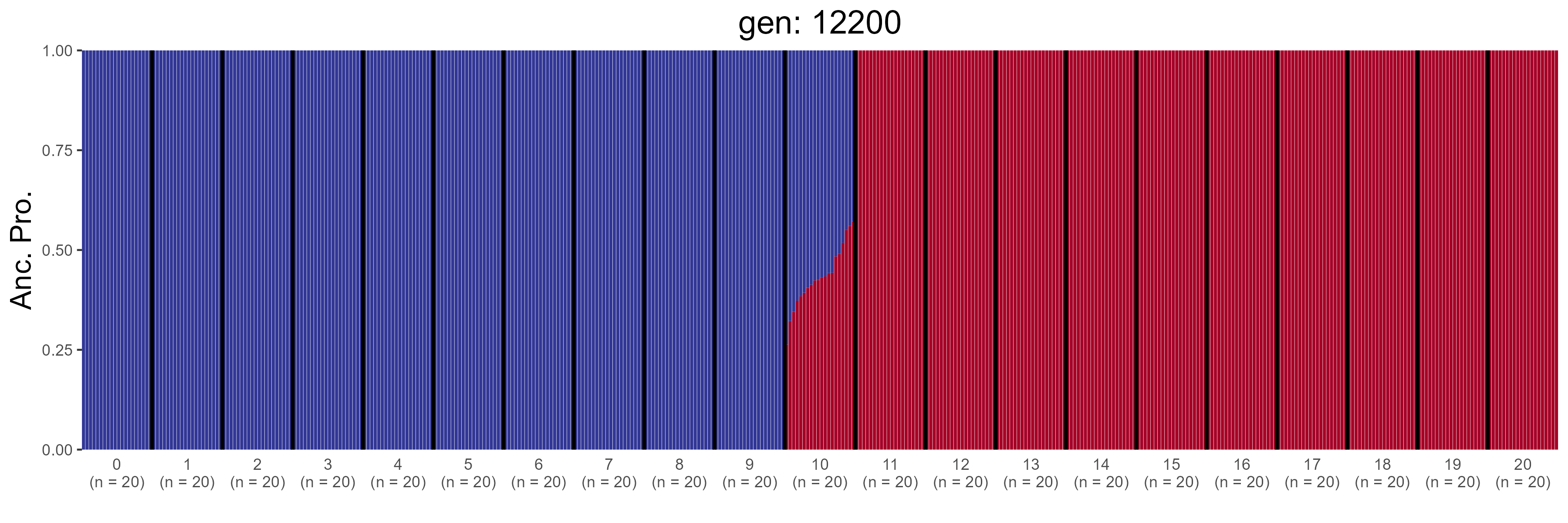

### GBC.12400.str.png

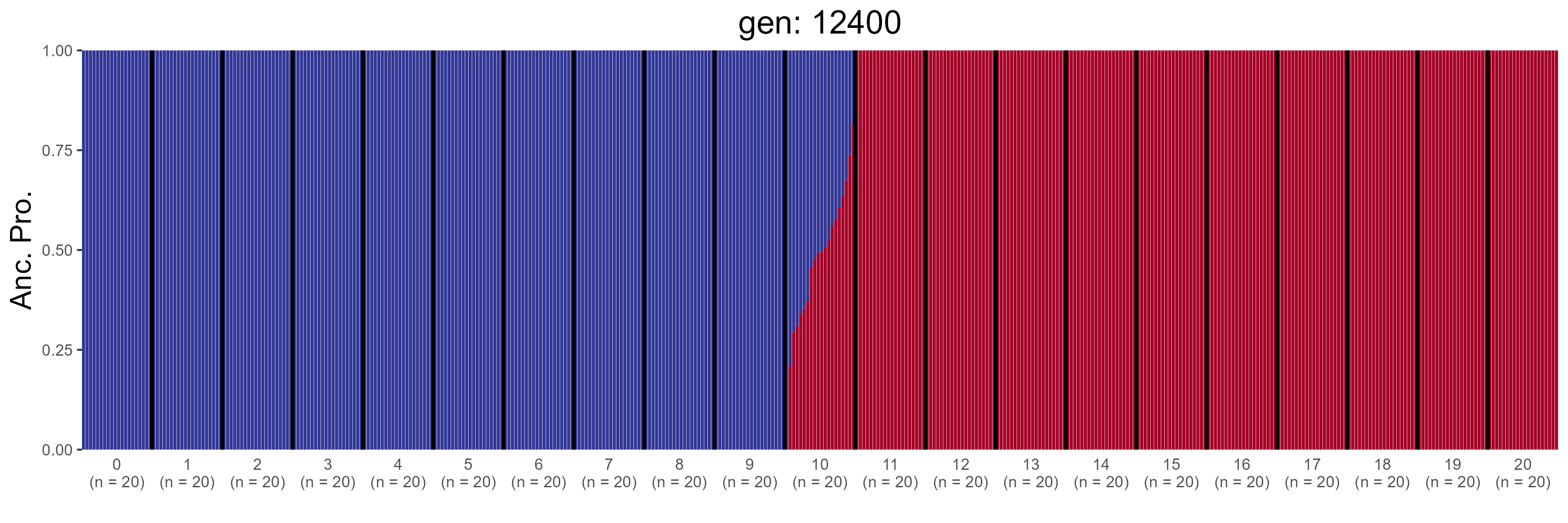

### GBC.12600.str.png

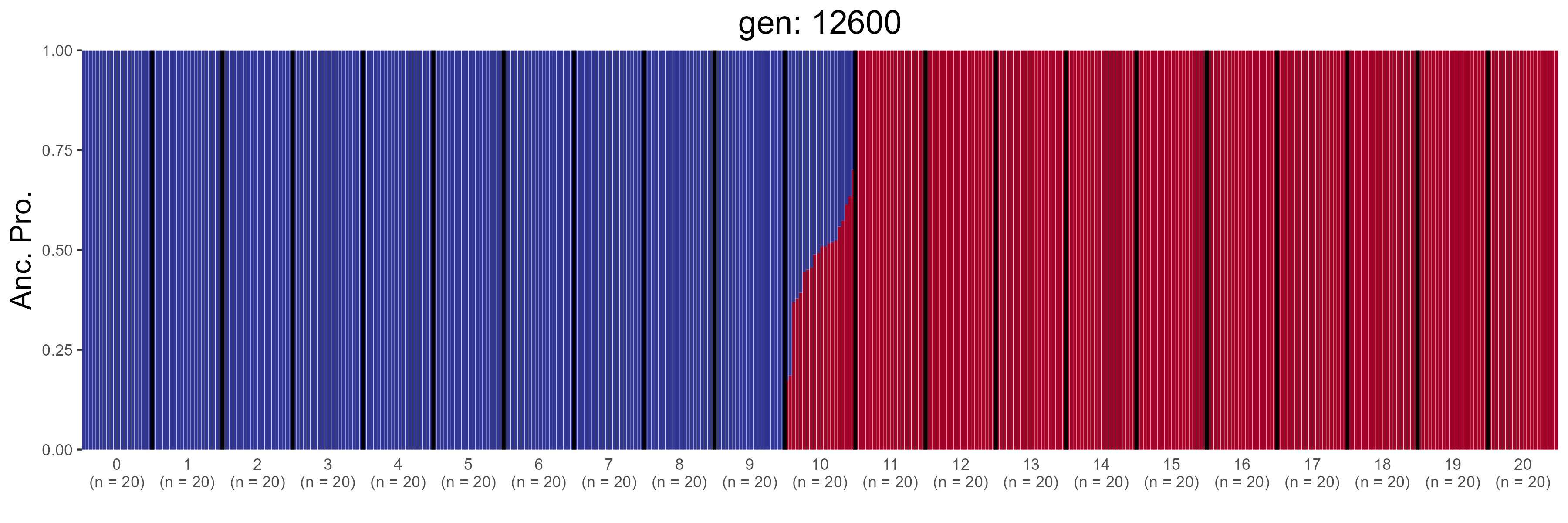

### GBC.12800.str.png

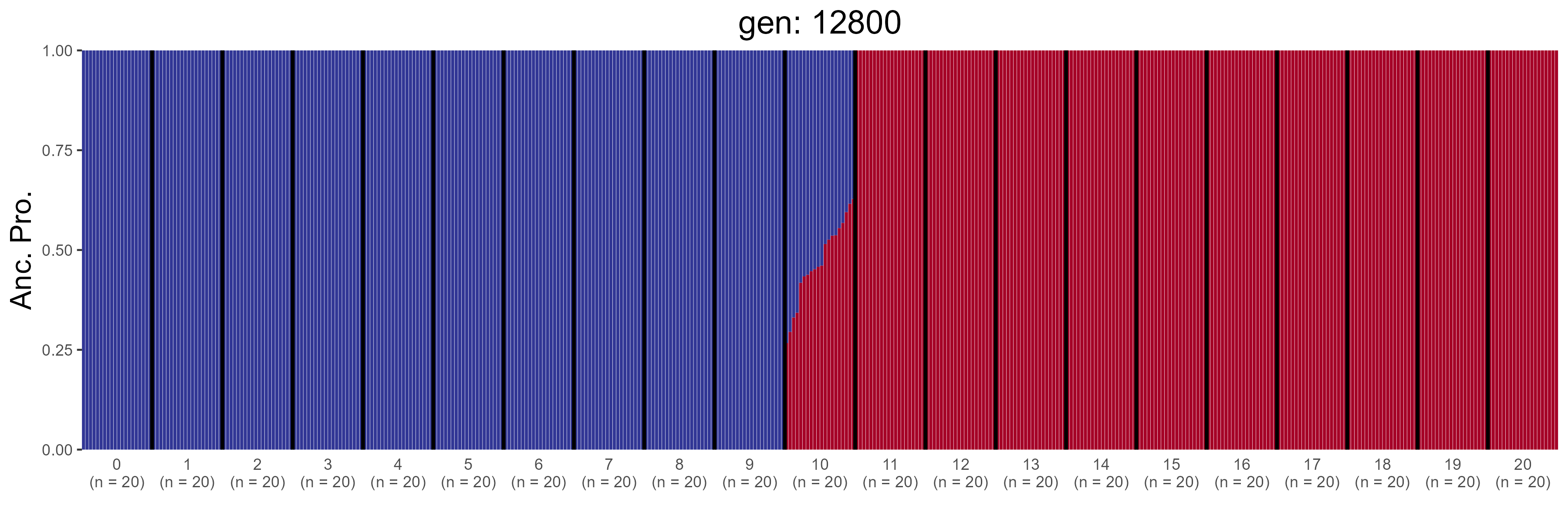

### GBC.13000.str.png

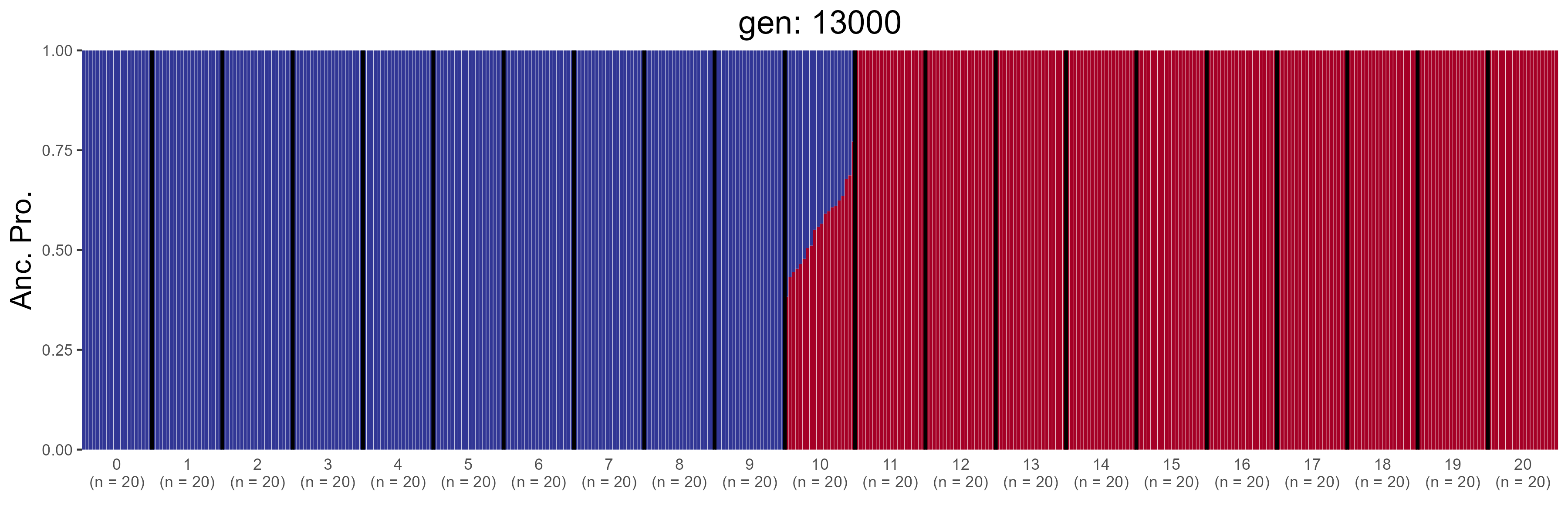

### GBC.13200.str.png

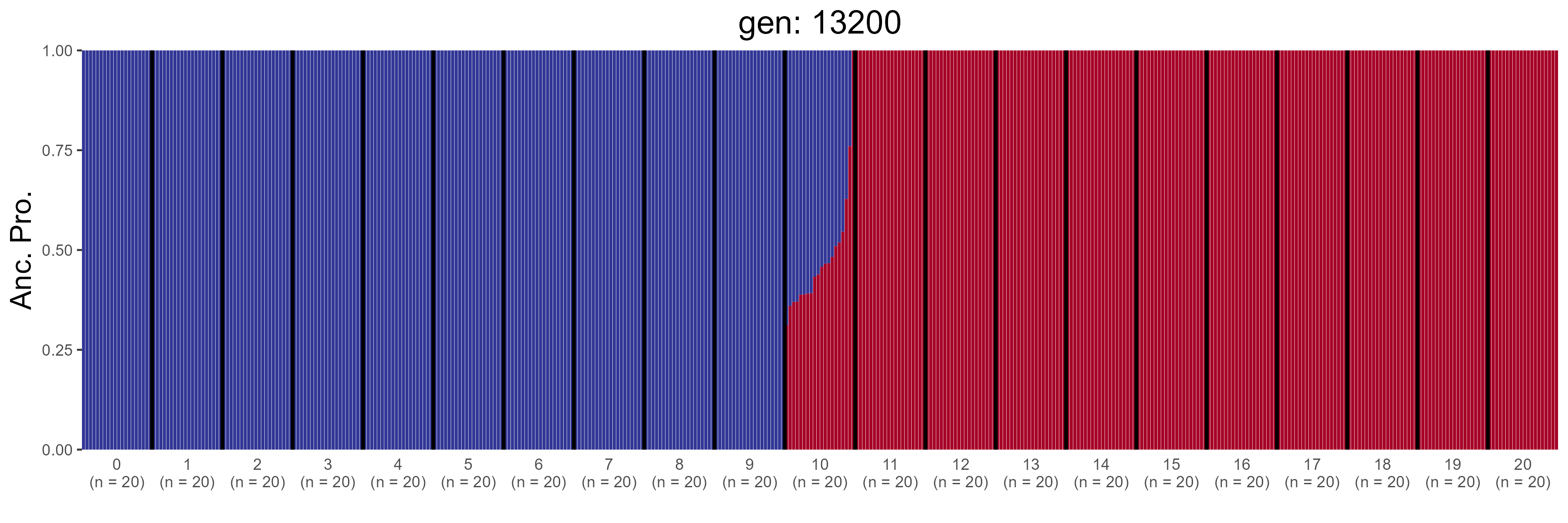

### GBC.13400.str.png

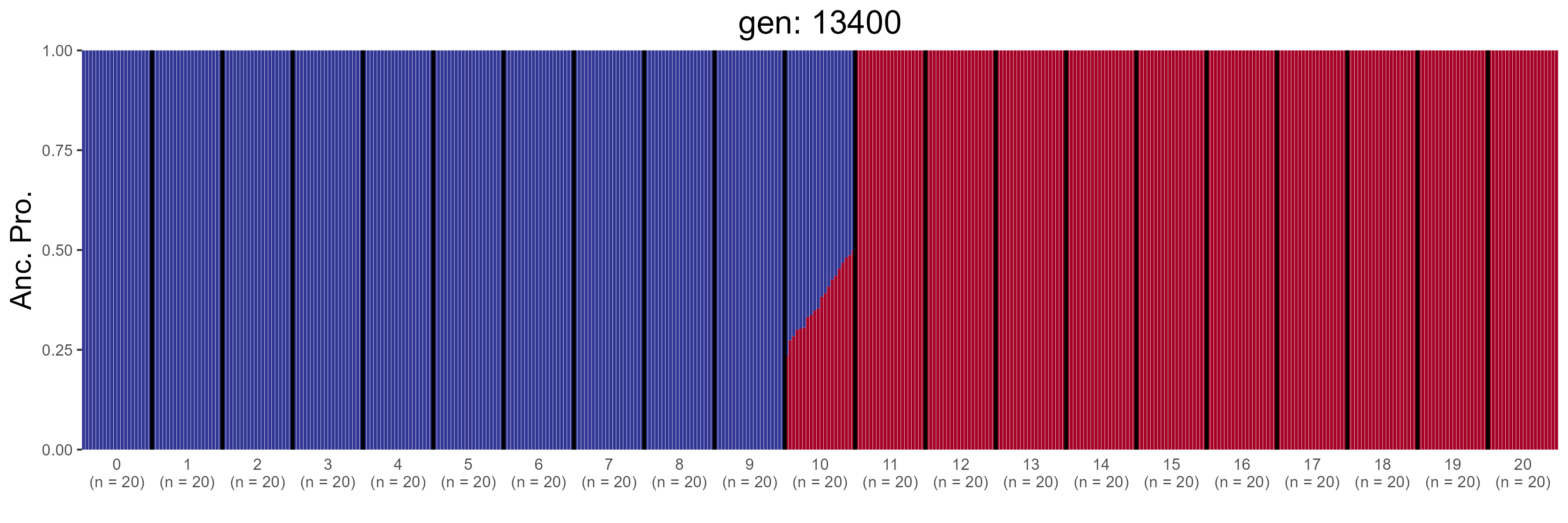

### GBC.13600.str.png

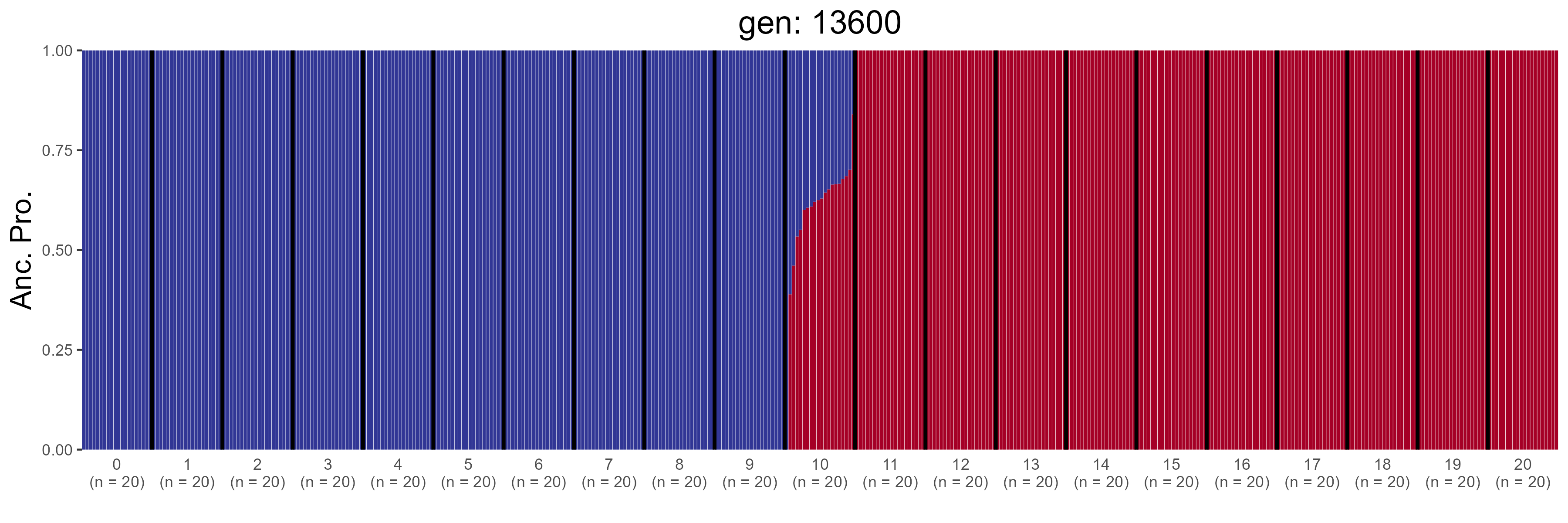

### GBC.13800.str.png

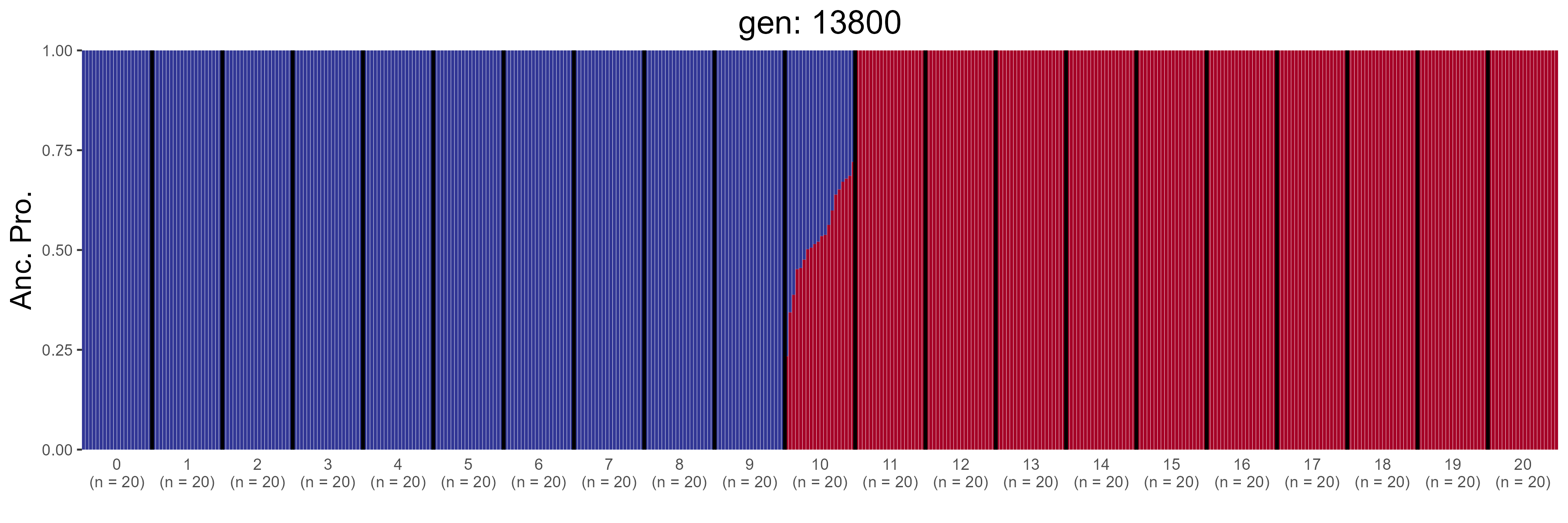

### GBC.14000.str.png

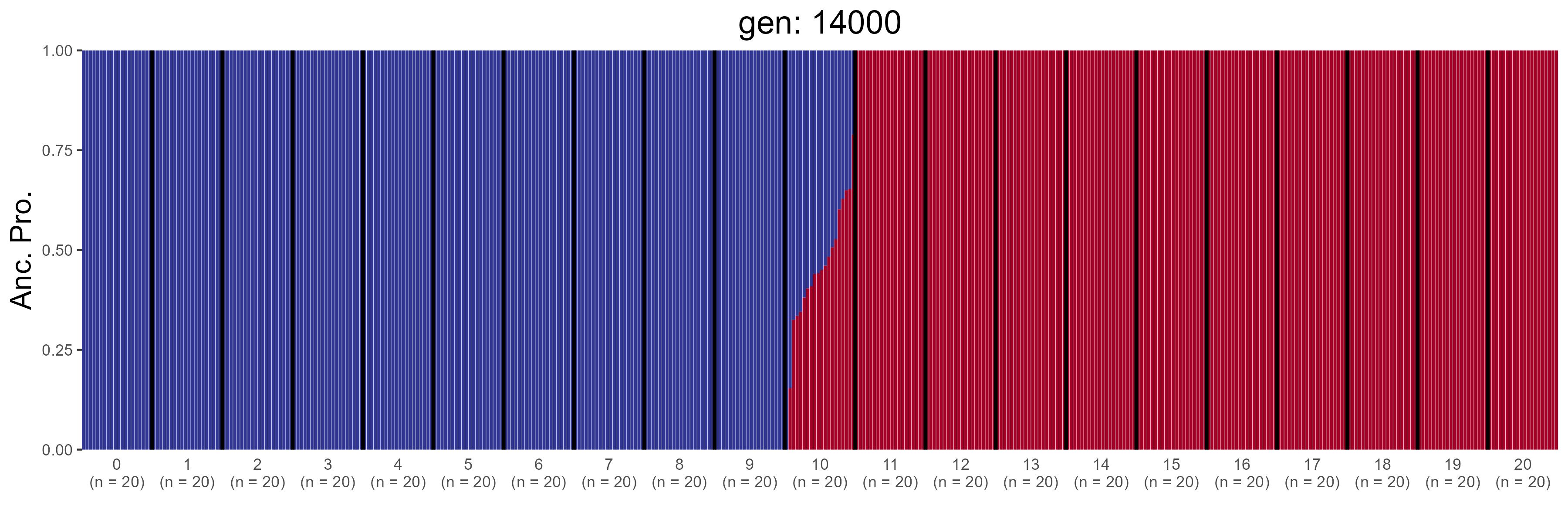

### GBC.14200.str.png

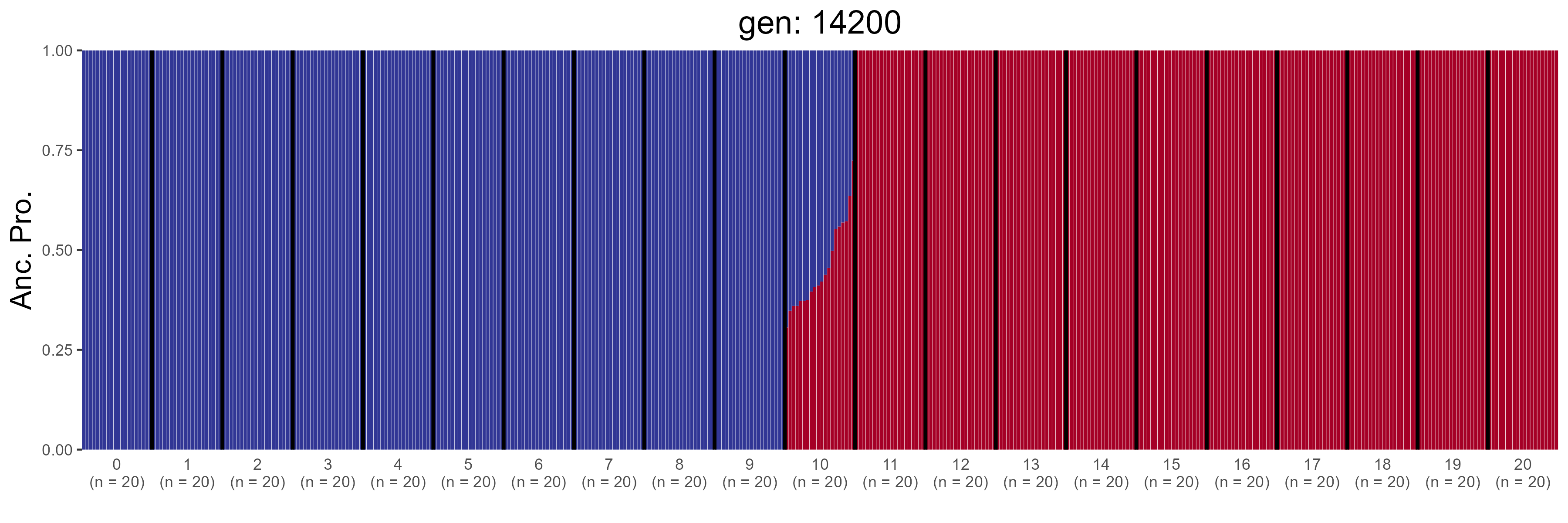

### GBC.14400.str.png

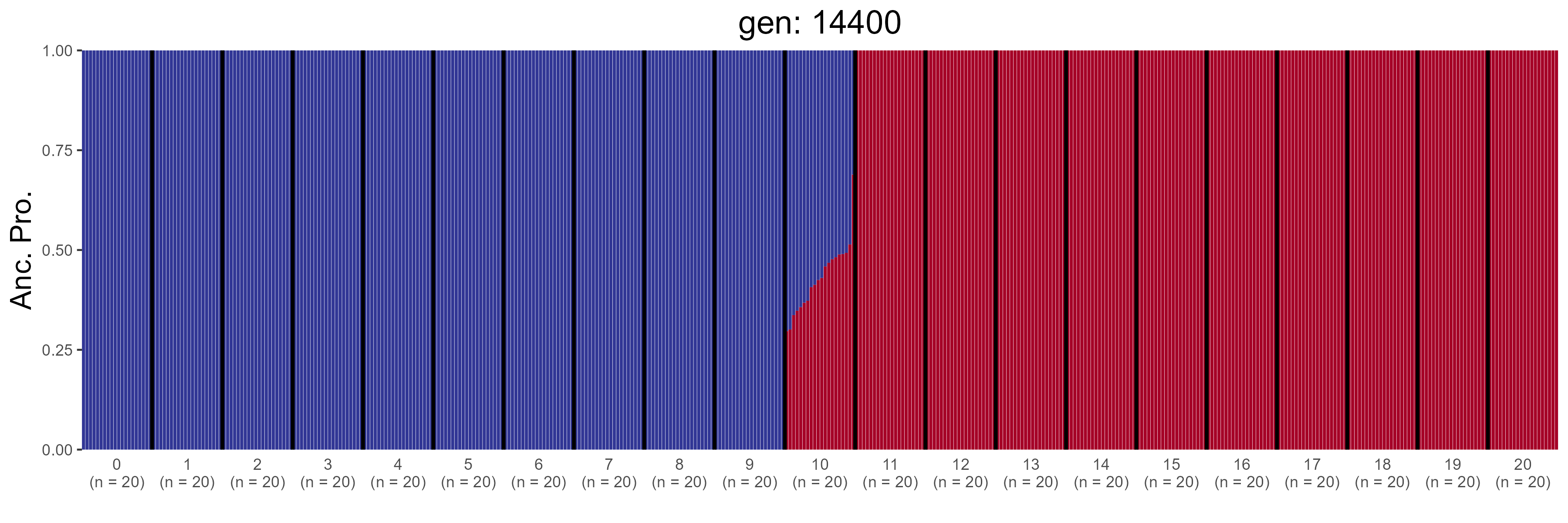

### GBC.14600.str.png

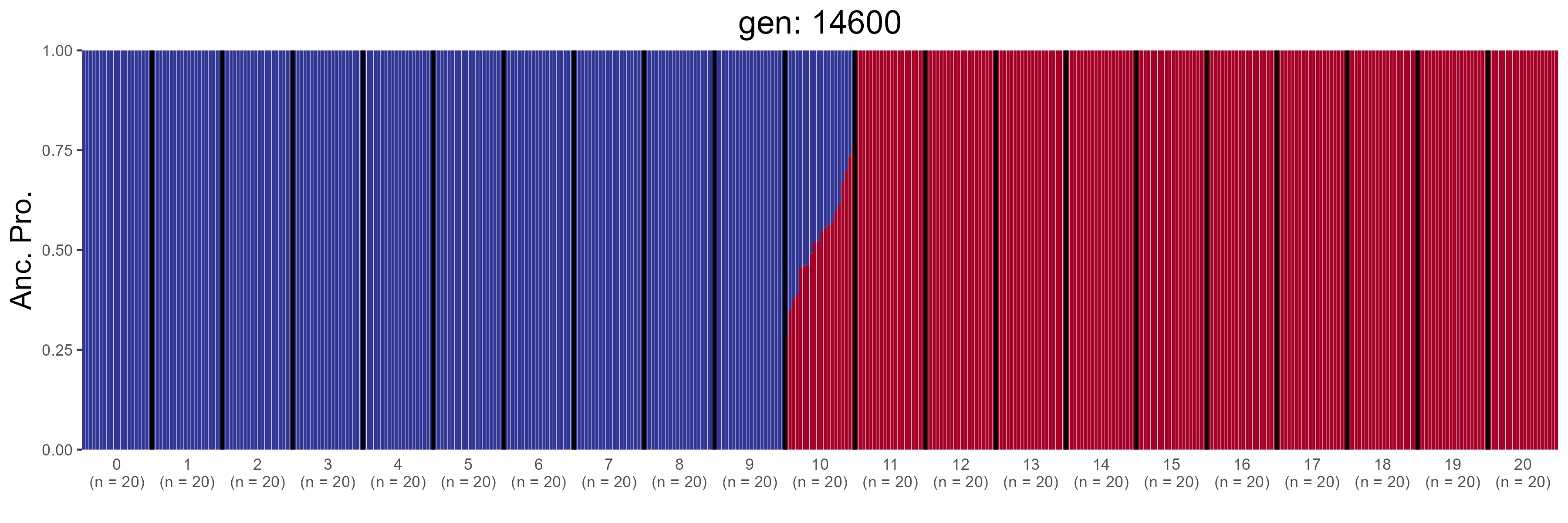

### GBC.14800.str.png

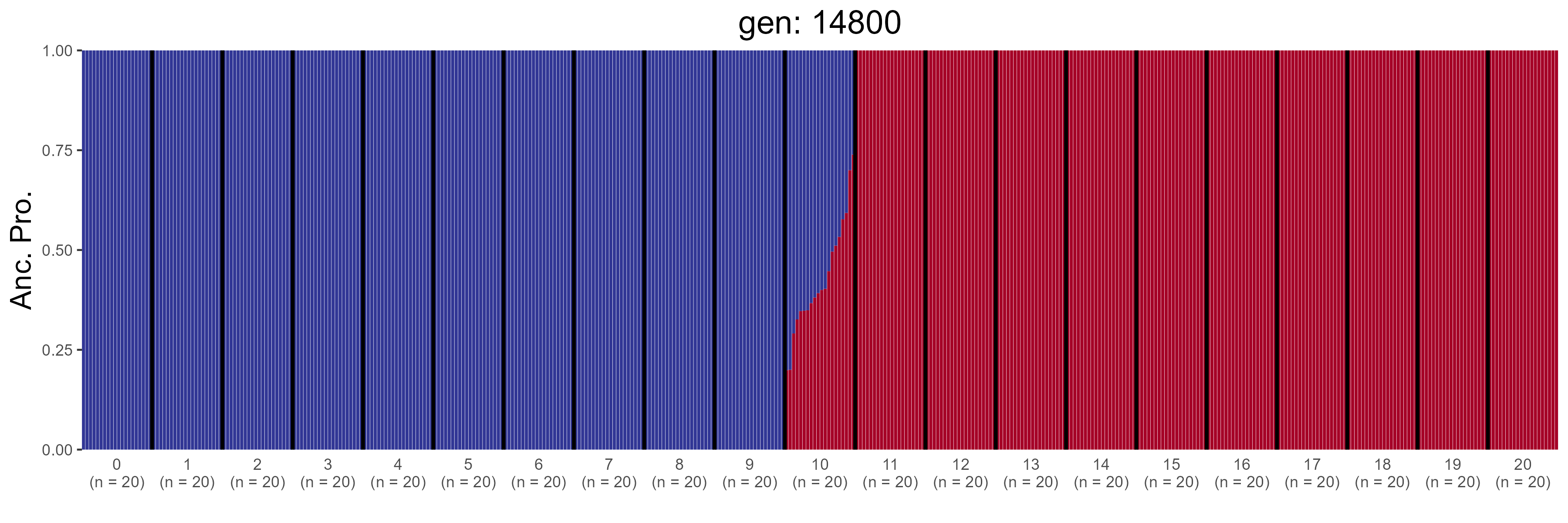

### GBC.15000.str.png

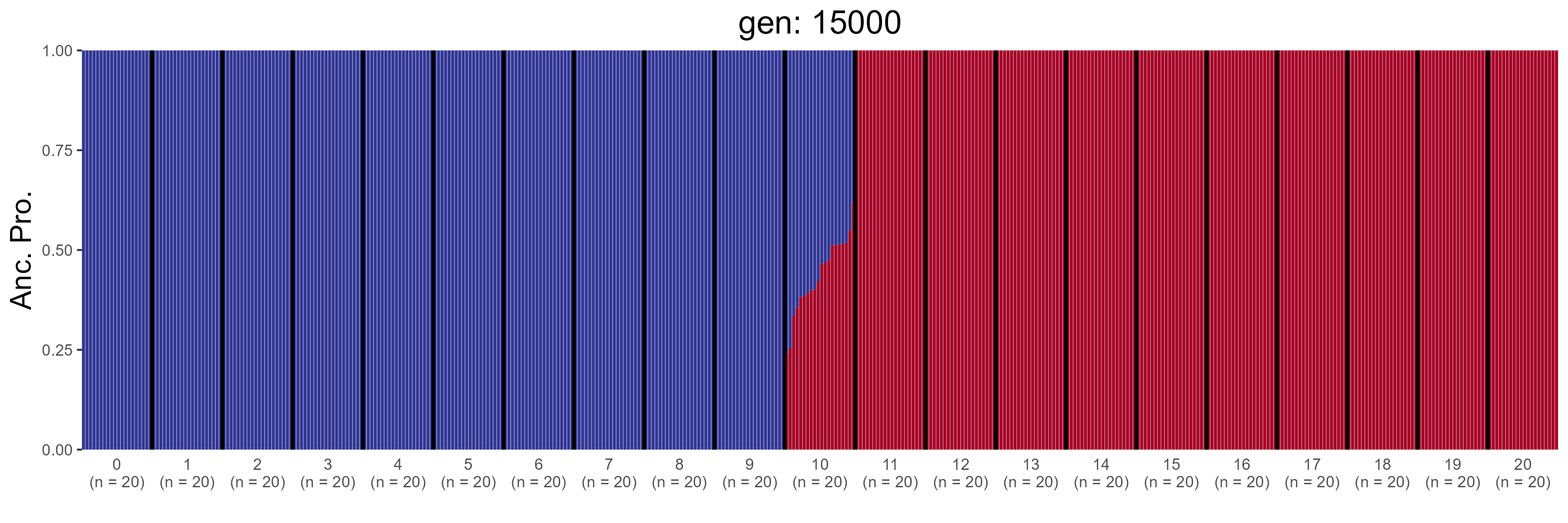

### GBC.15200.str.png

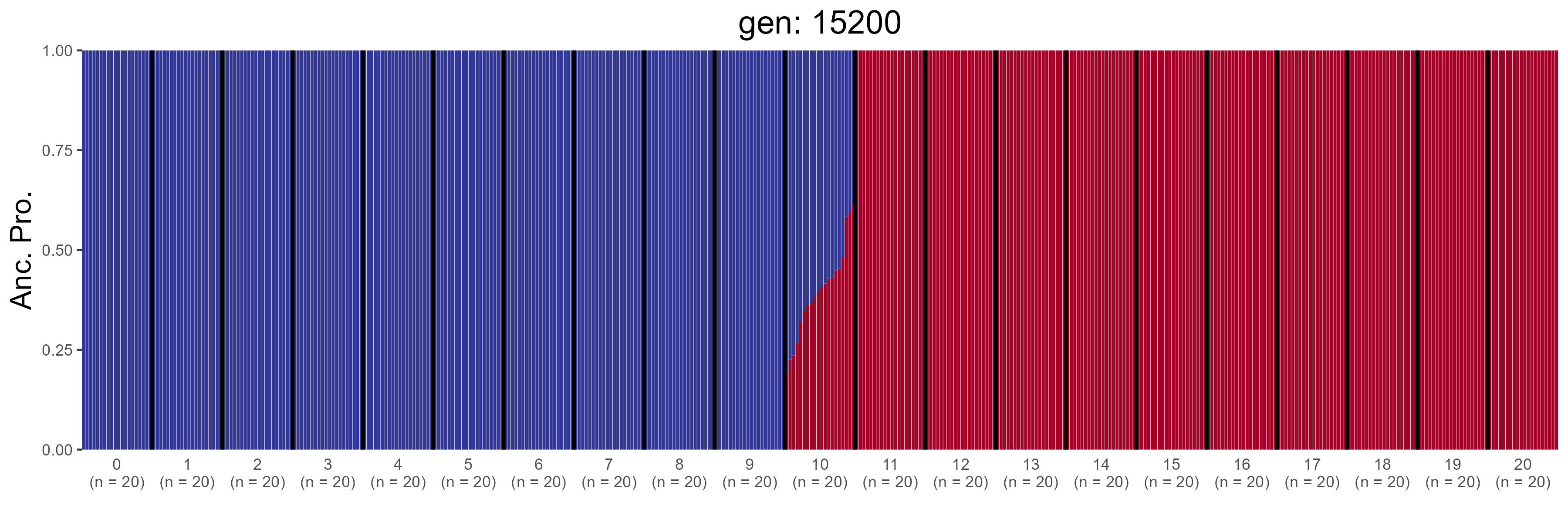

### GBC.15400.str.png

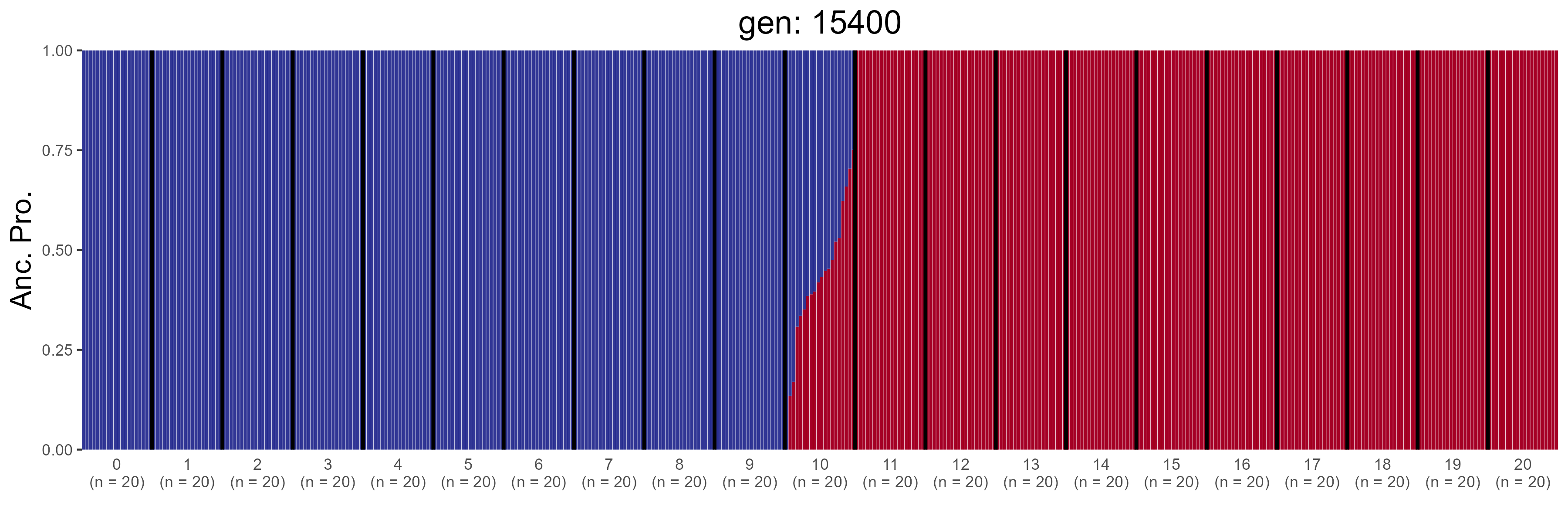

### GBC.15600.str.png

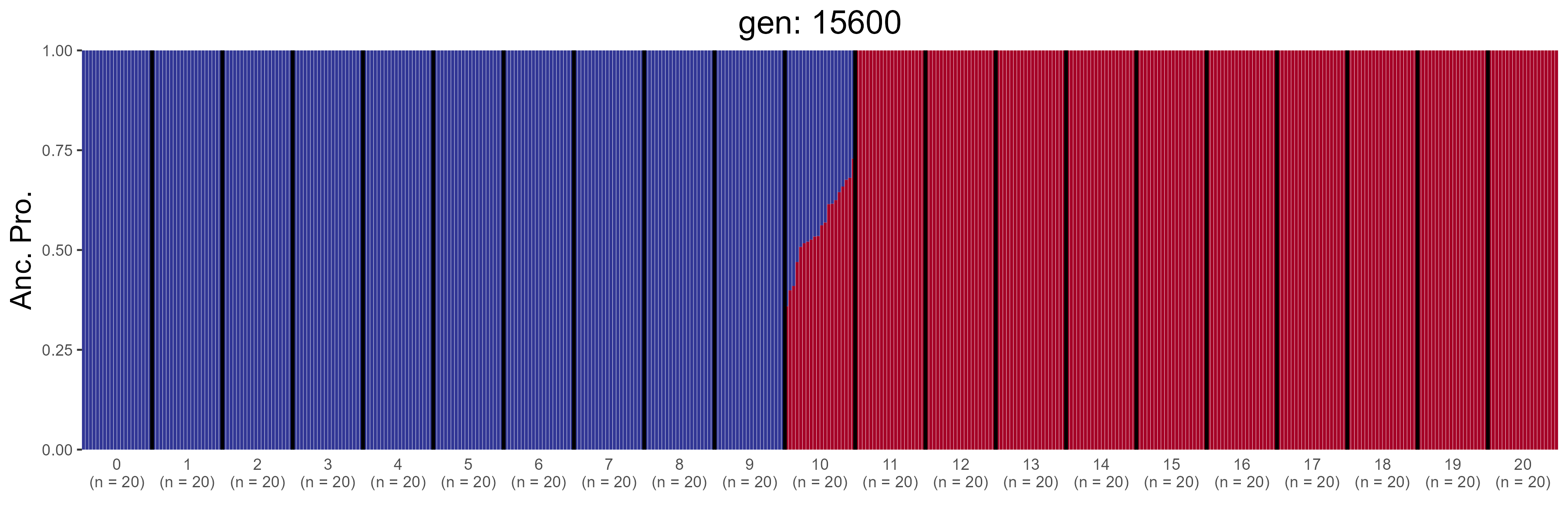

### GBC.15800.str.png

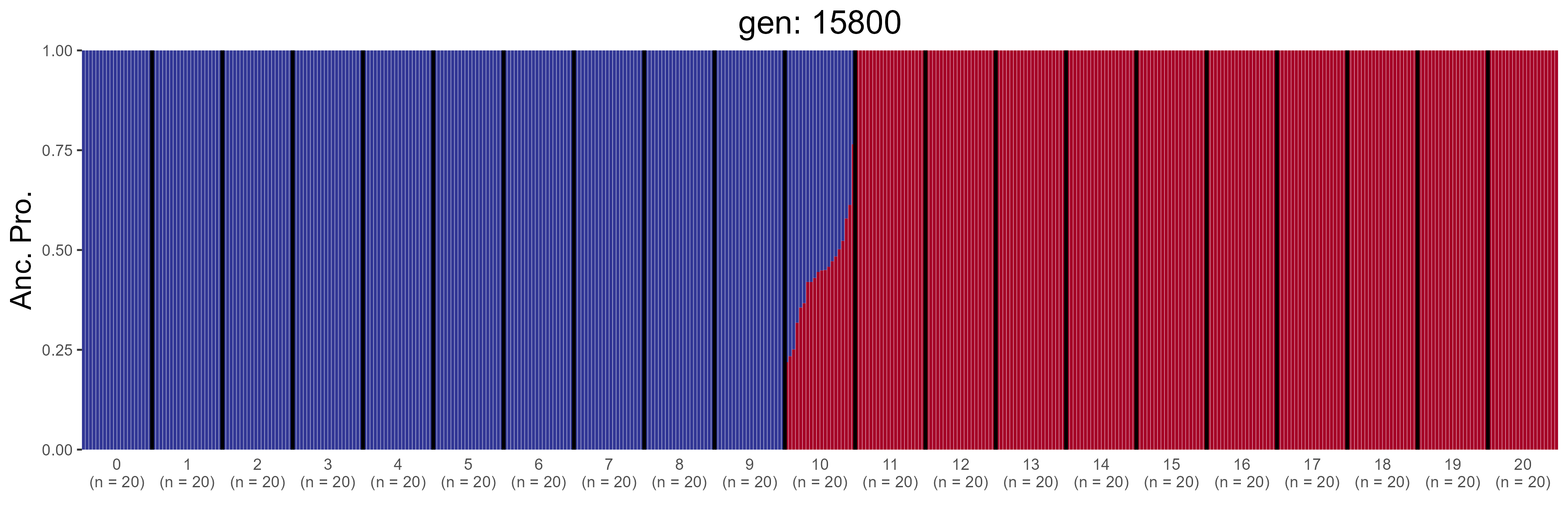

### GBC.16000.str.png

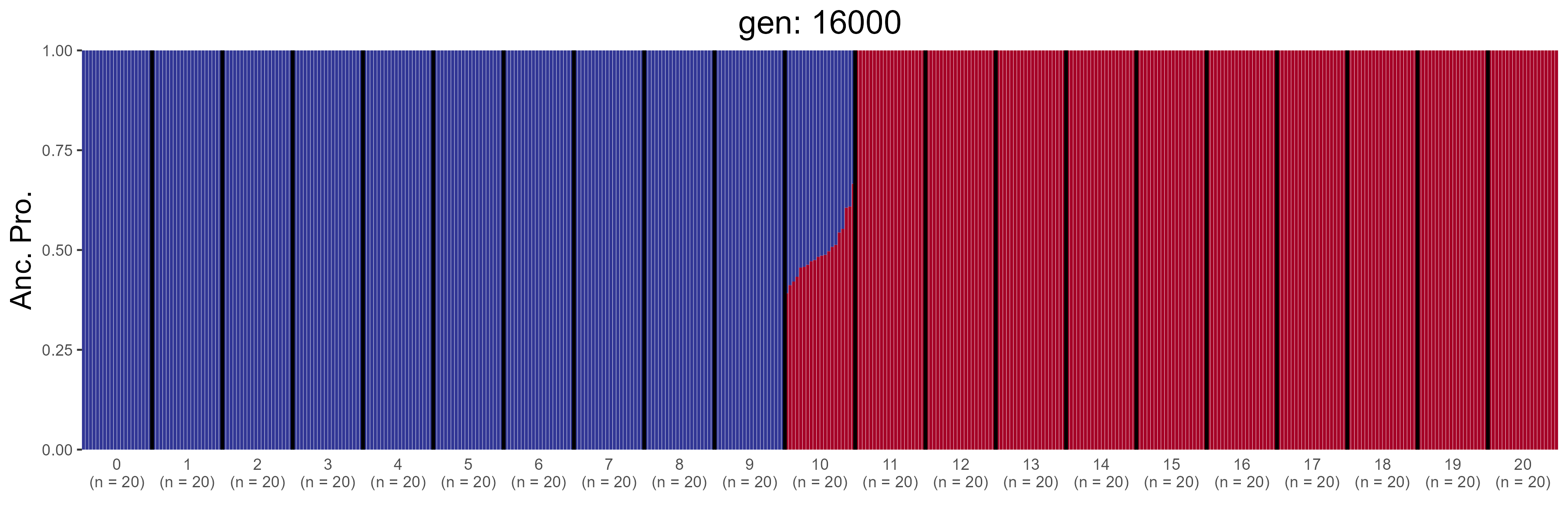

### GBC.16200.str.png

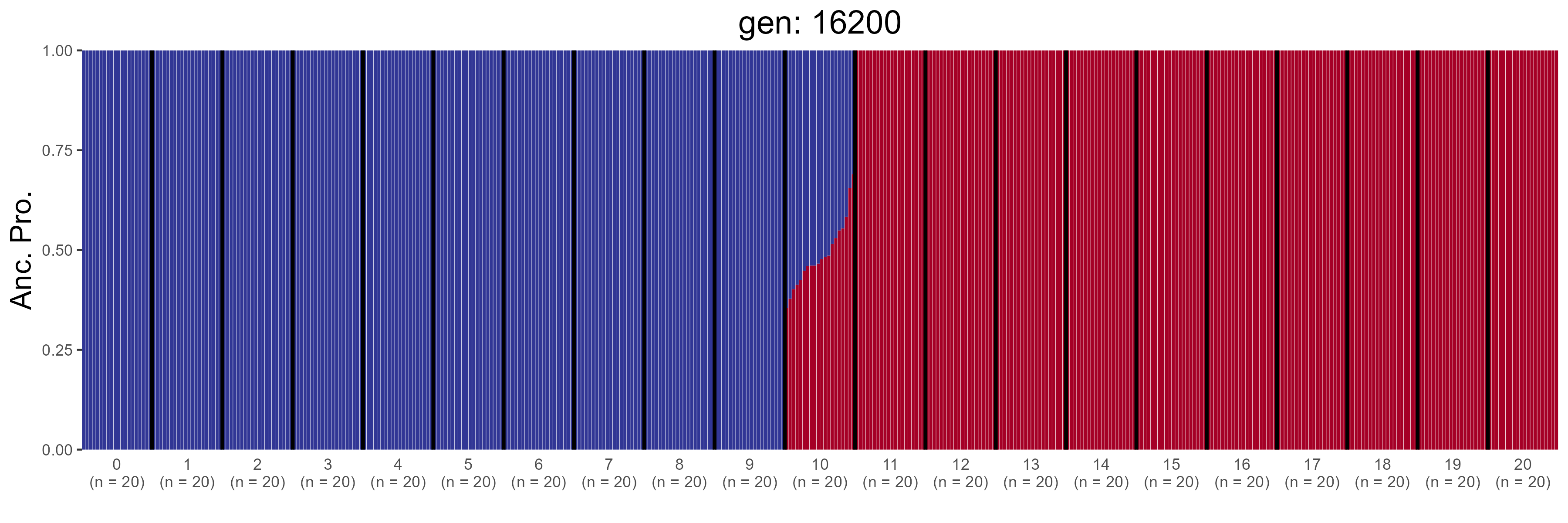

### GBC.16400.str.png

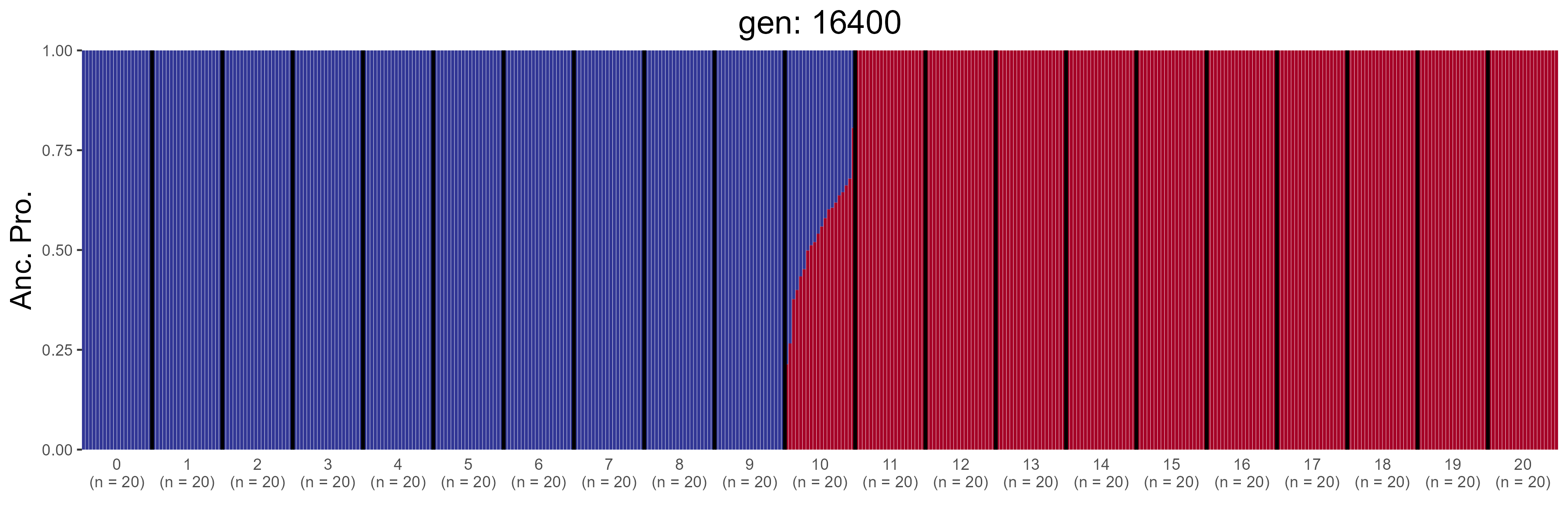

### GBC.16600.str.png

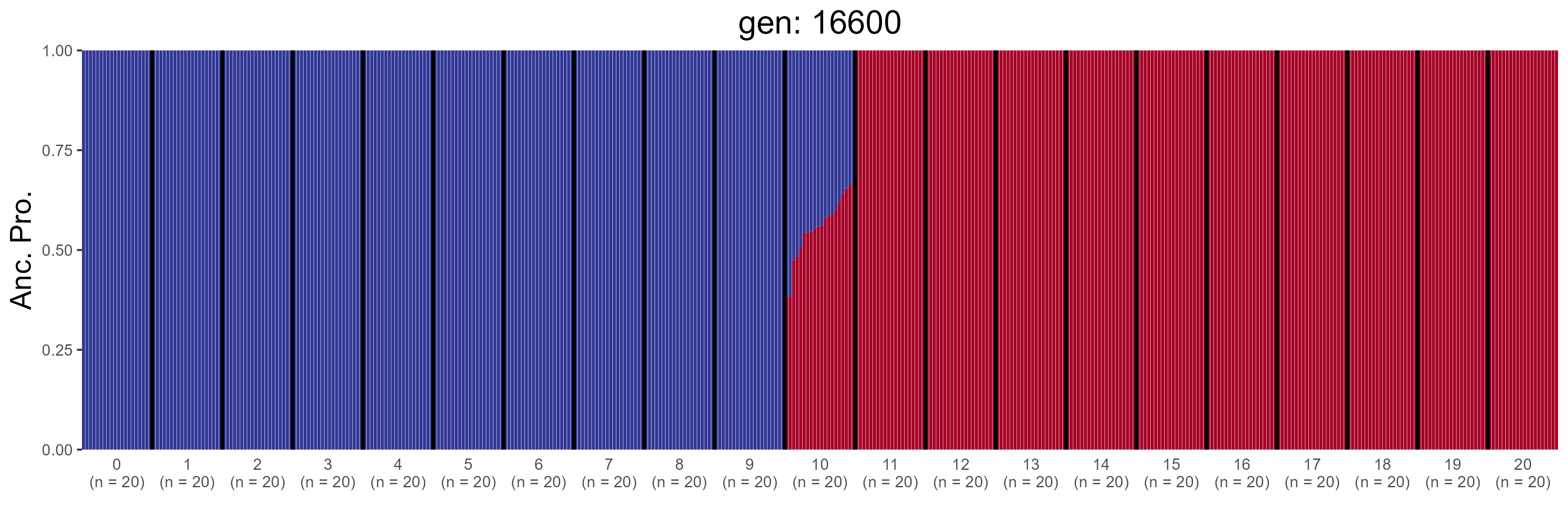

### GBC.16800.str.png

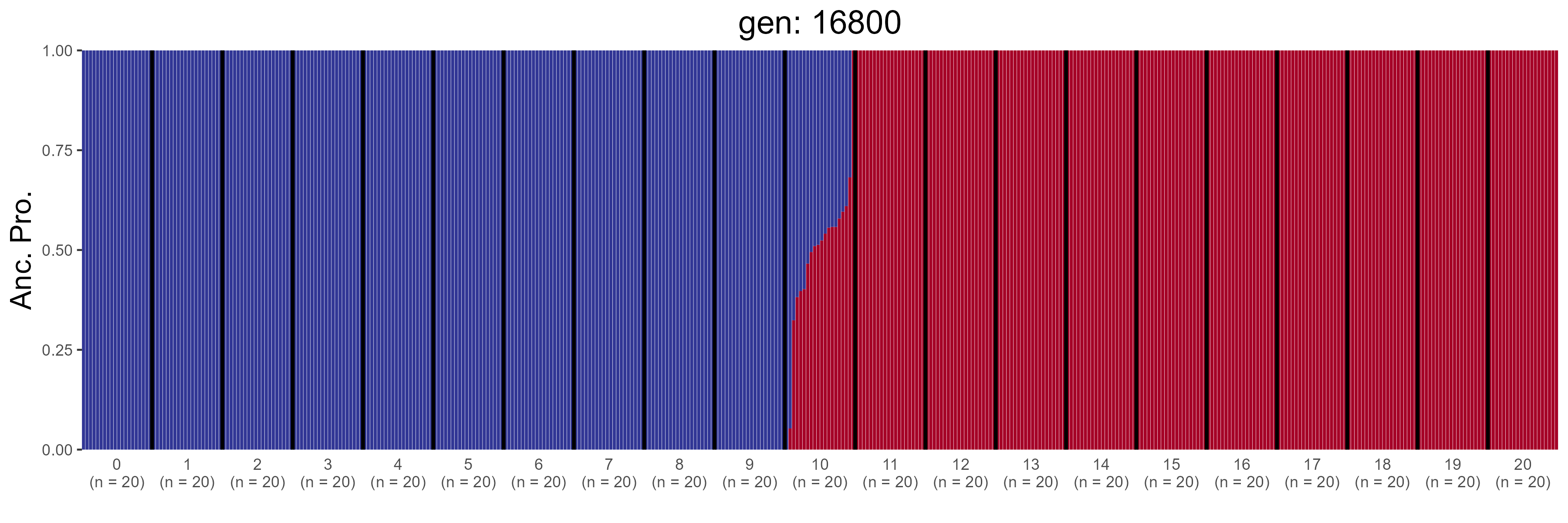

### GBC.17000.str.png

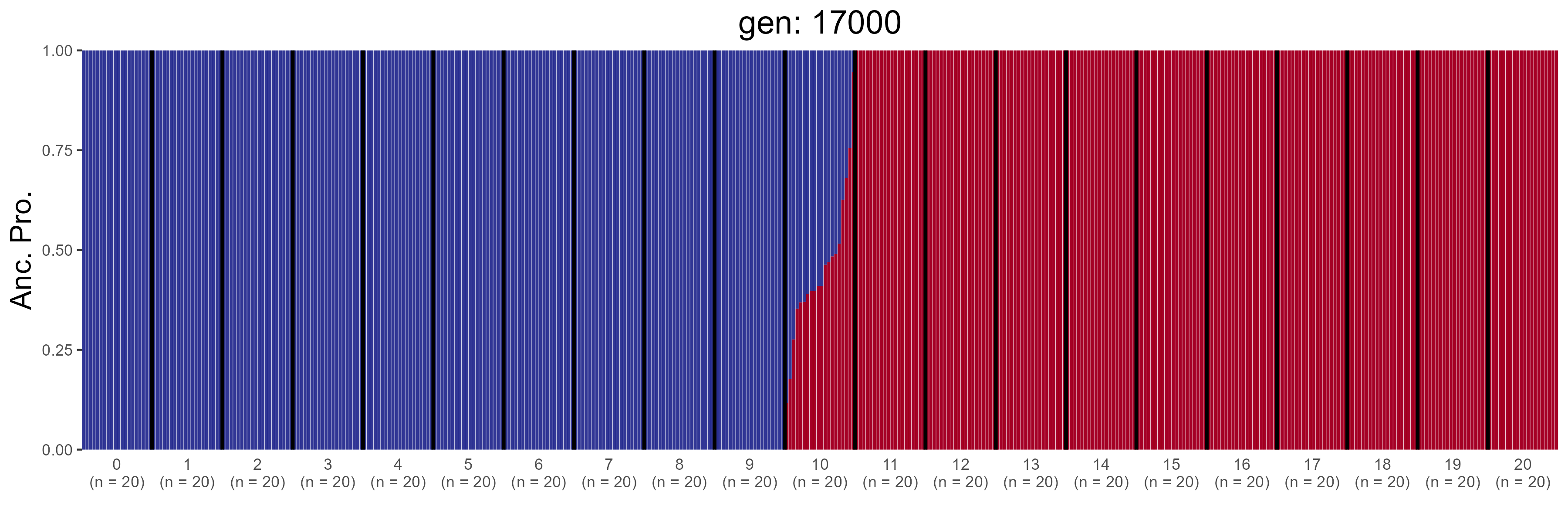

### GBC.17200.str.png

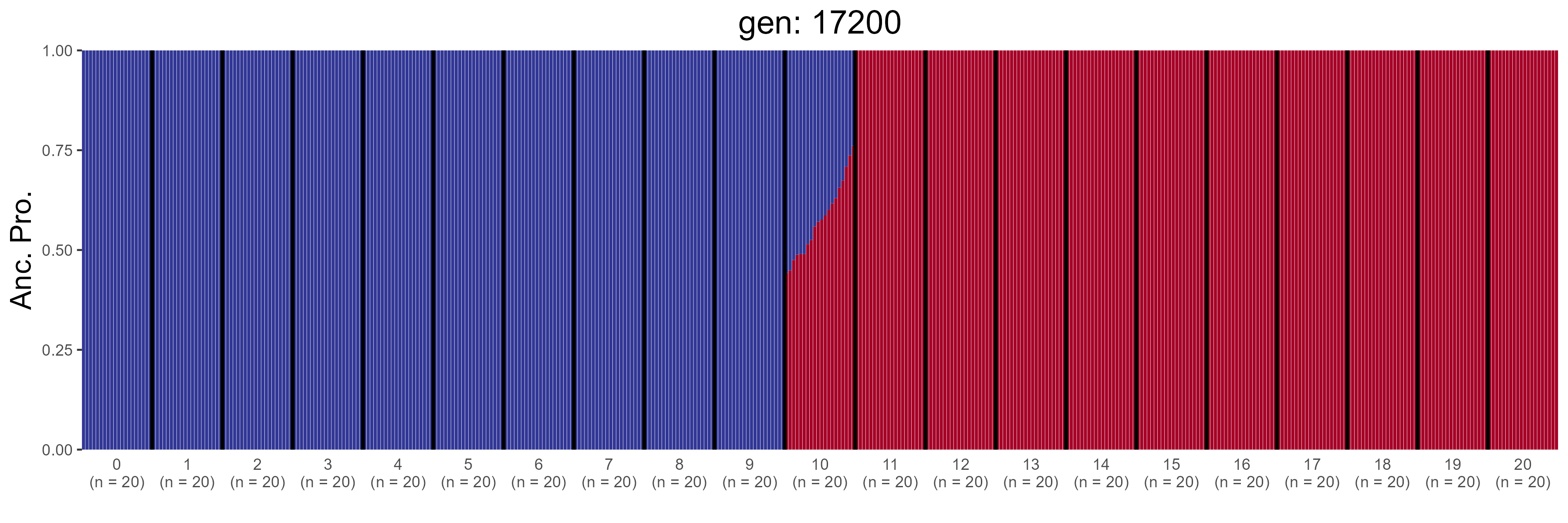

### GBC.17400.str.png

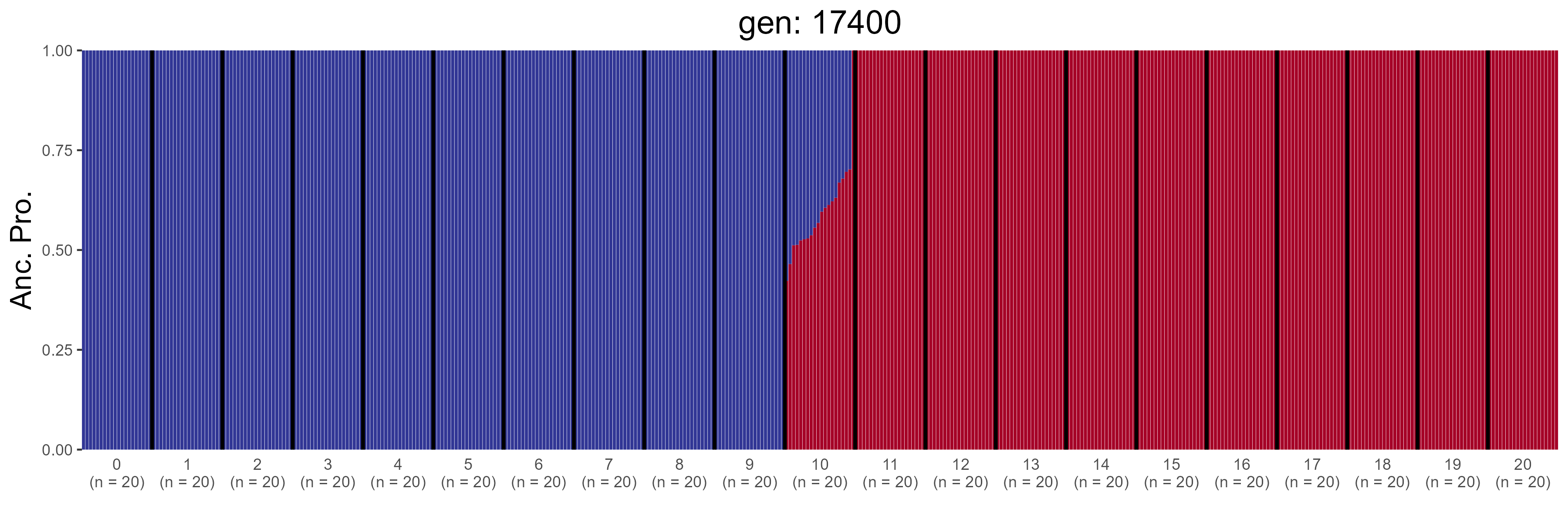

### GBC.17600.str.png

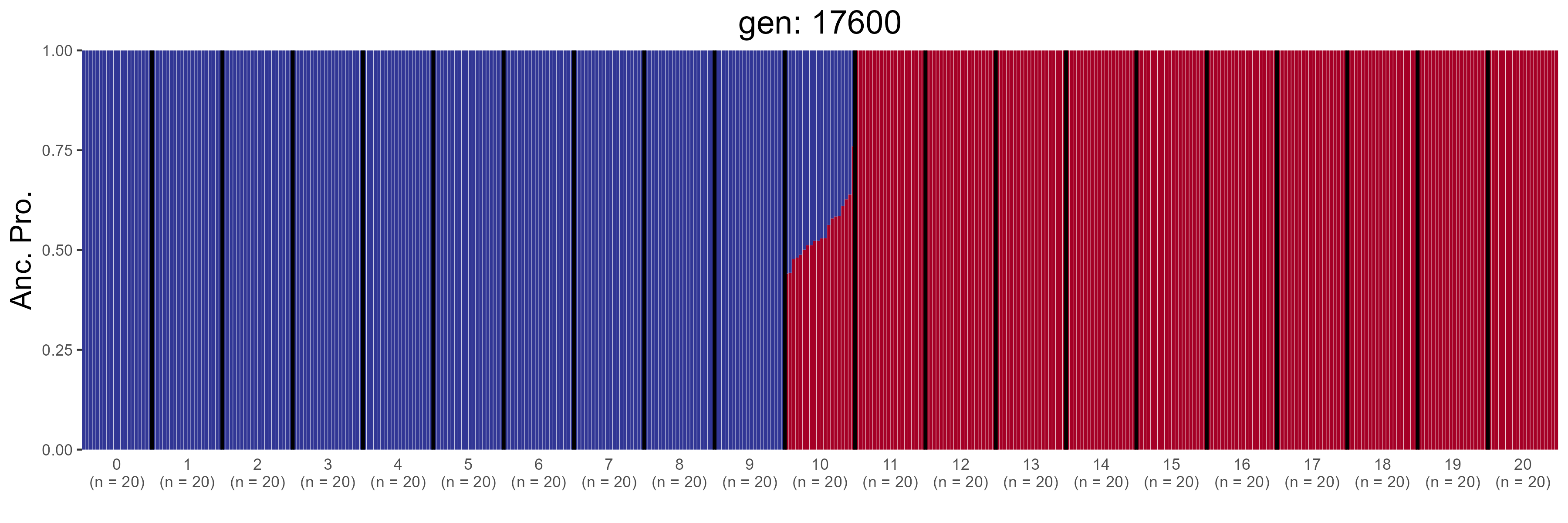

### GBC.17800.str.png

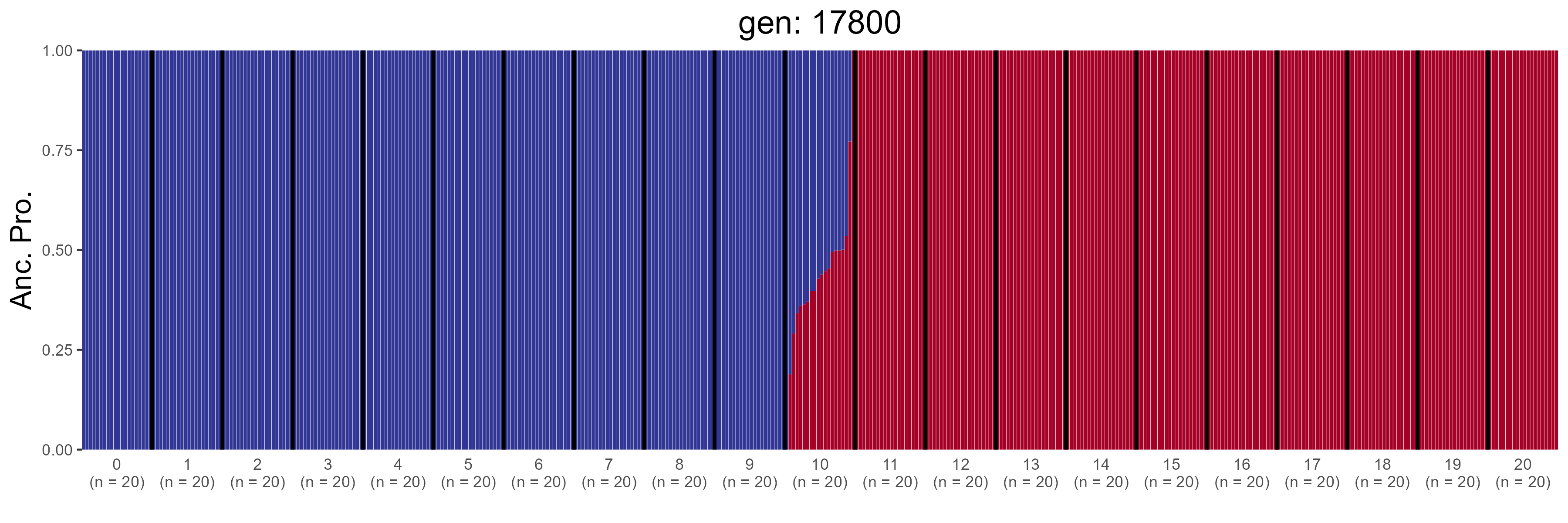

### GBC.18000.str.png

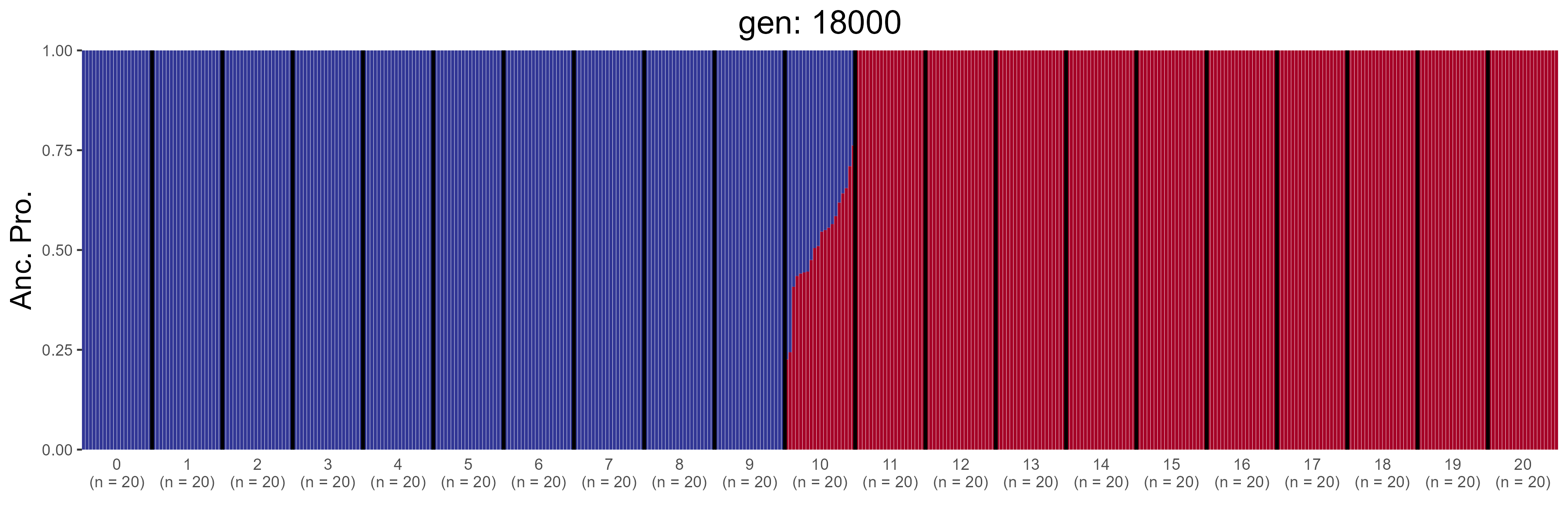

### GBC.18200.str.png

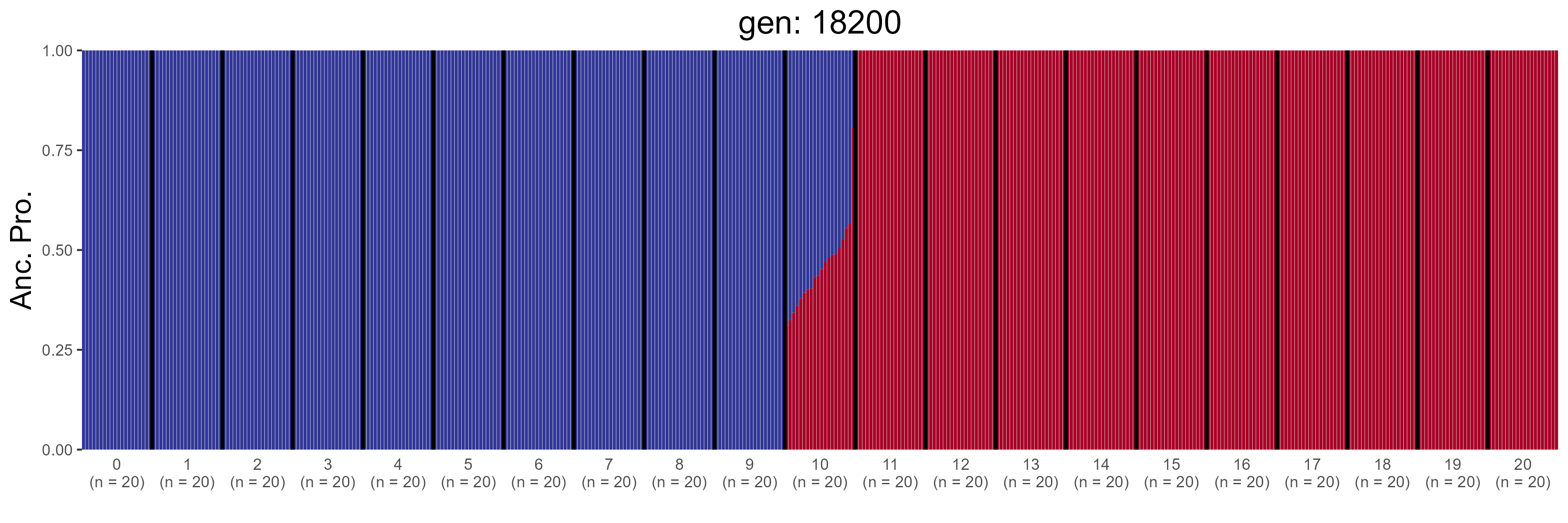

### GBC.18400.str.png

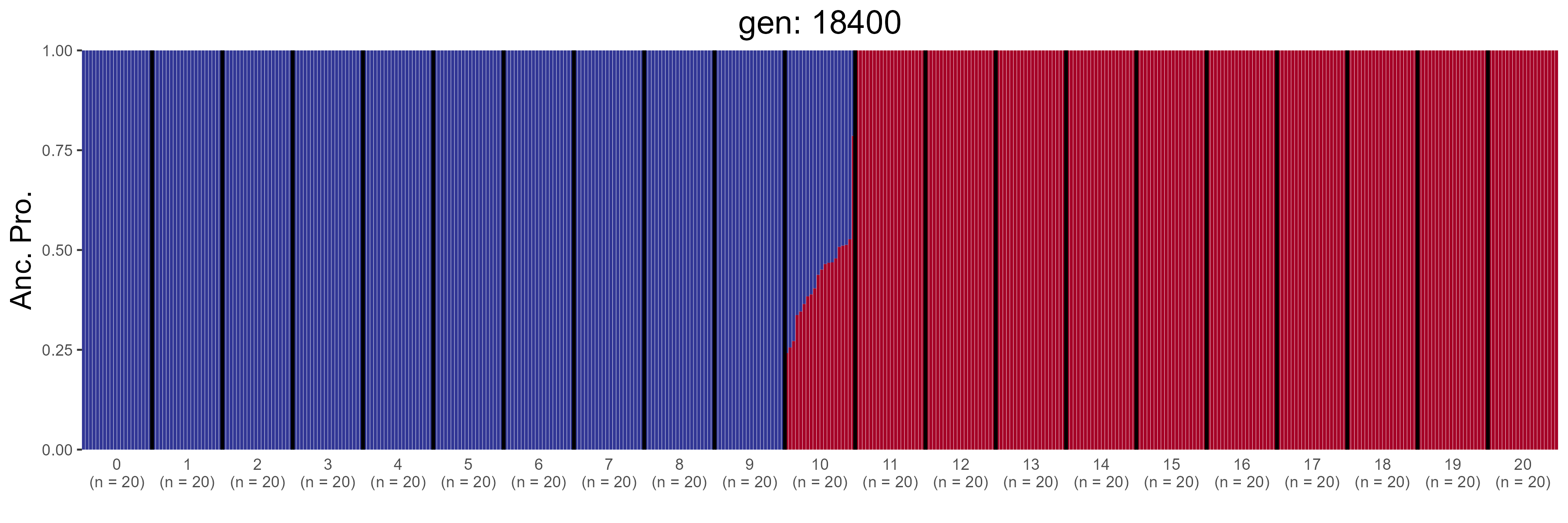

### GBC.18600.str.png

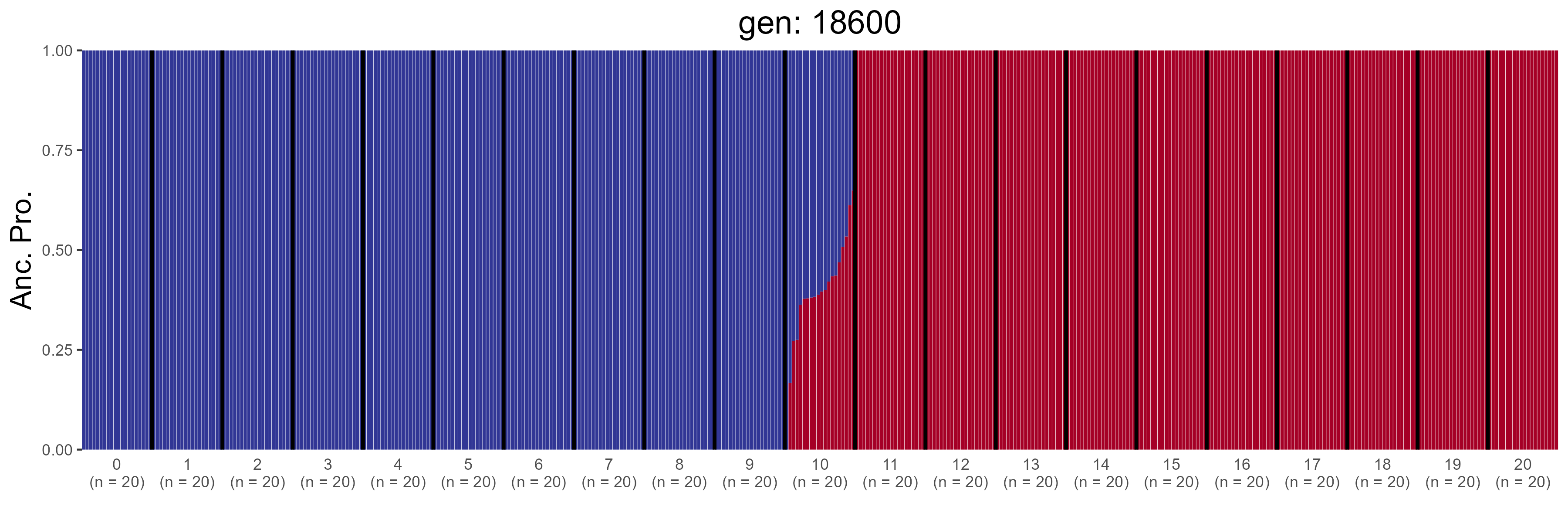

### GBC.18800.str.png

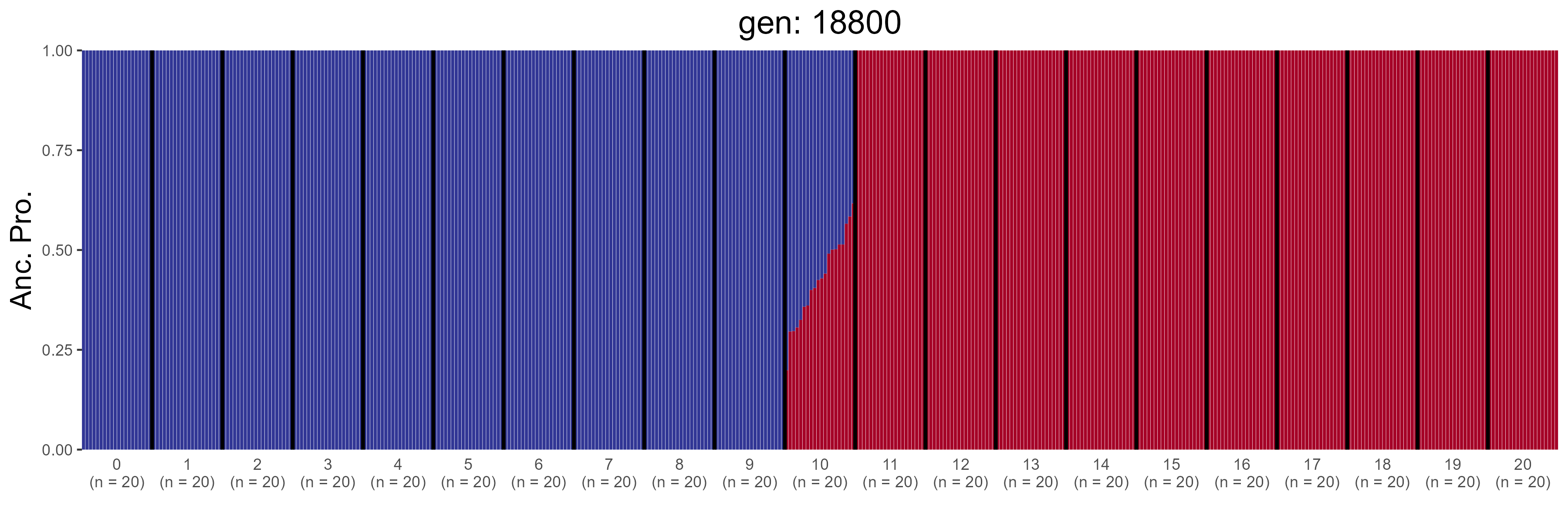

### GBC.19000.str.png

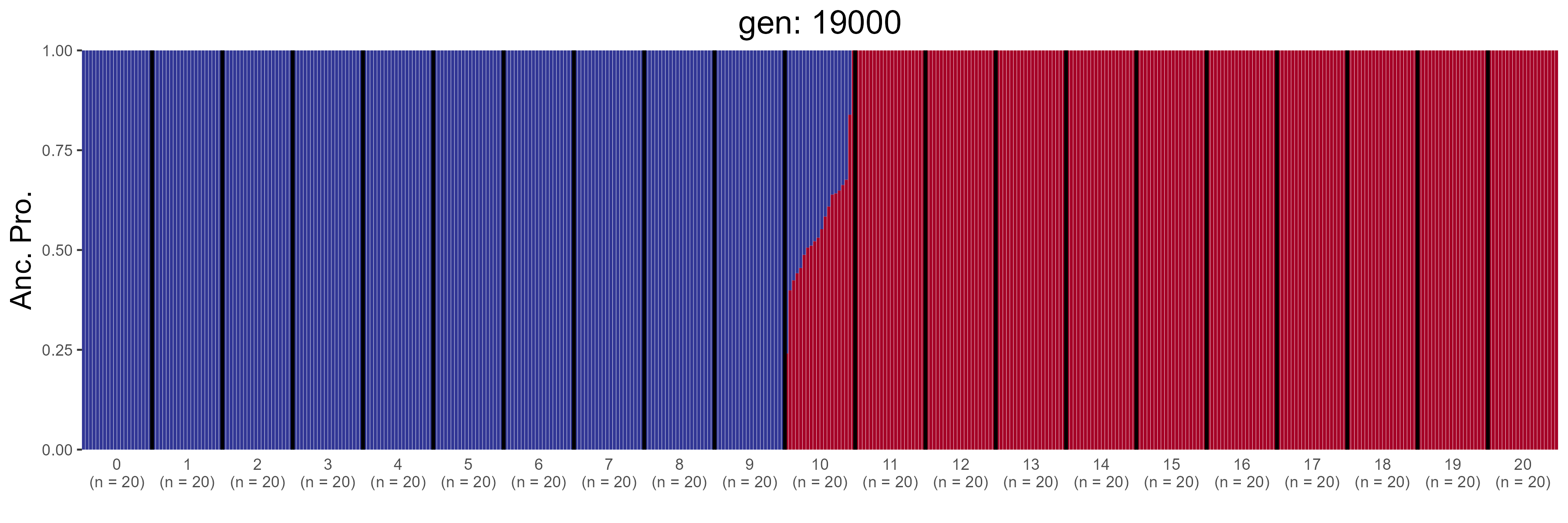

### IBD.0.str.png

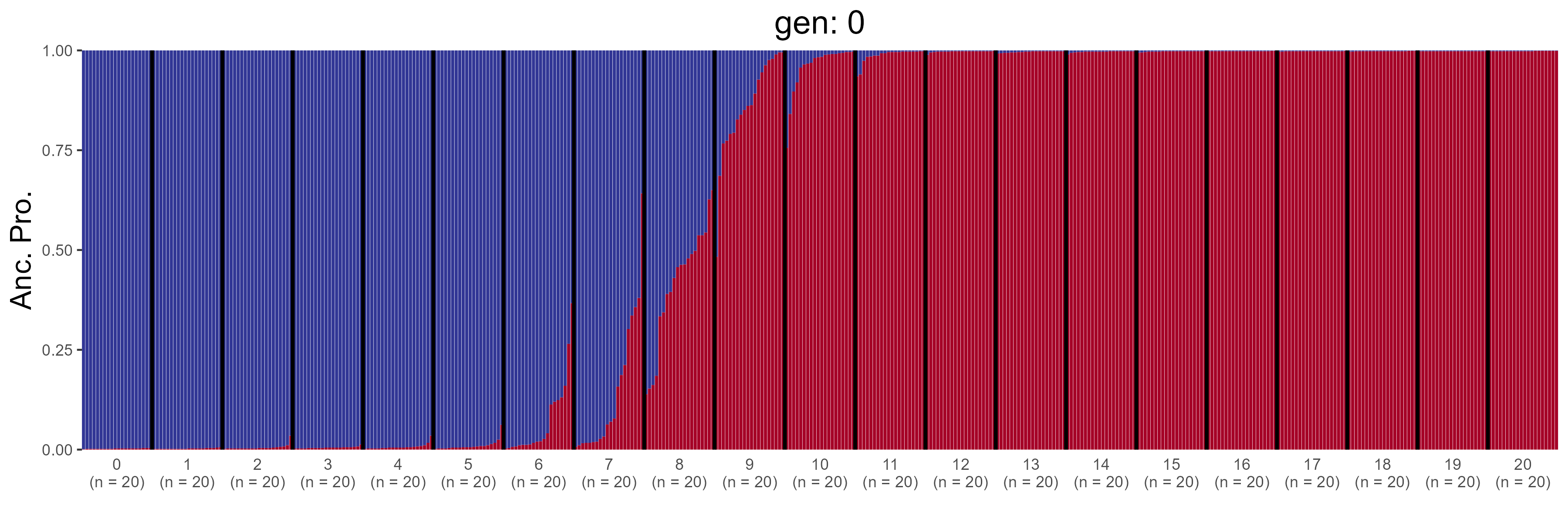

### IBD.1000.str.png

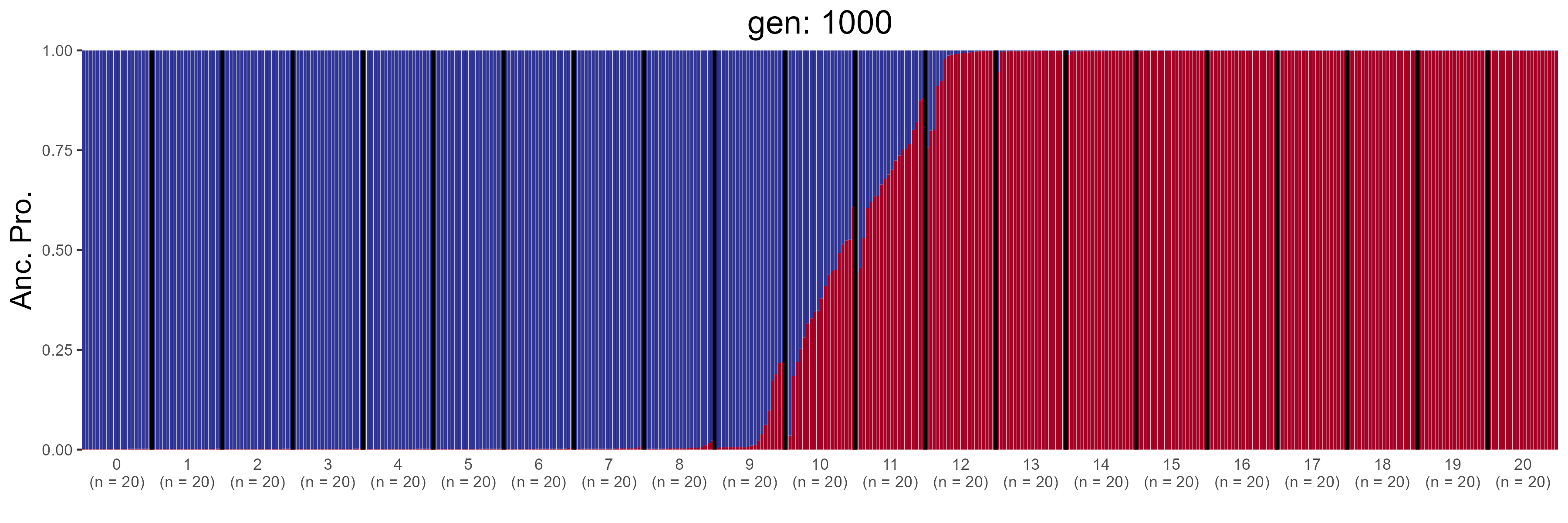

### IBD.10000.str.png

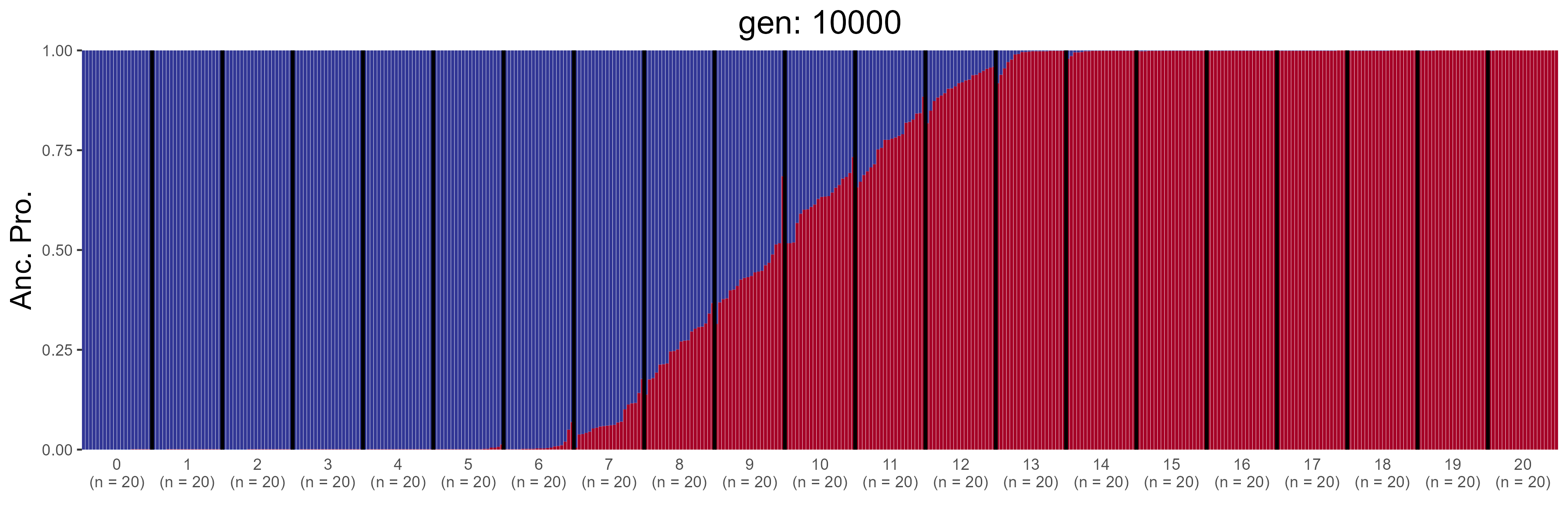

### IBD.10200.str.png

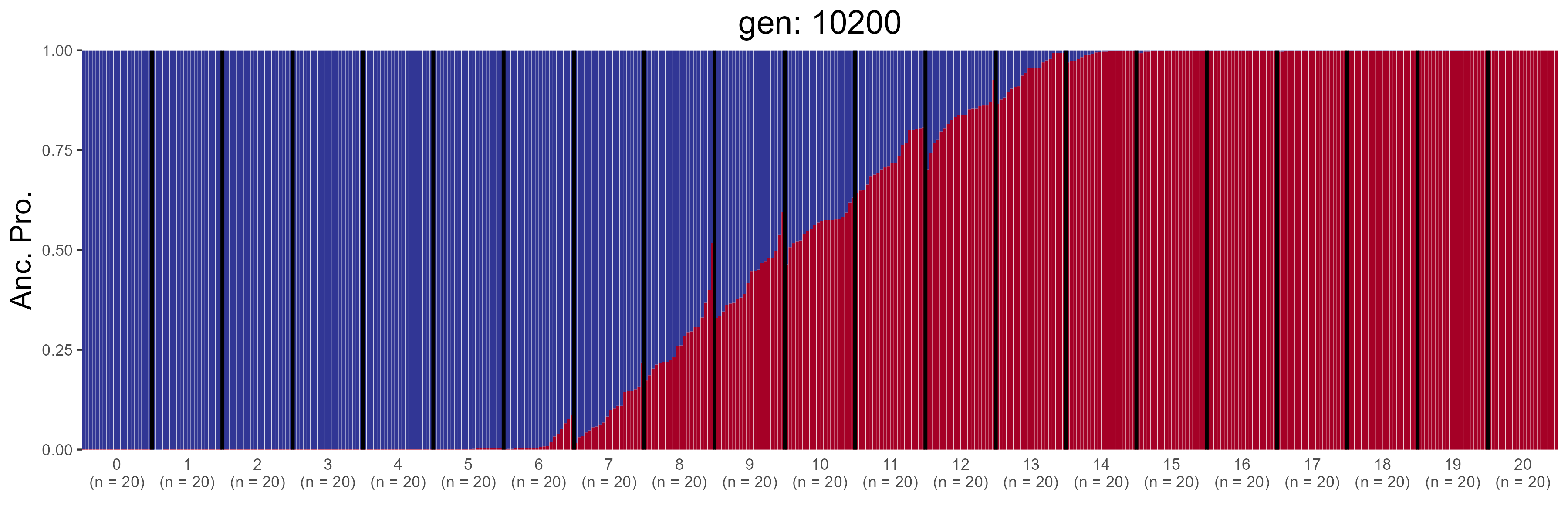
